# A Novel Approach for Accurate Sequence Assembly Using de Bruijn graphs

**DOI:** 10.1101/2024.05.29.596541

**Authors:** Cameron J. Prybol, Aeron T. Hammack, Euan A. Ashley, Michael P. Snyder

## Abstract

Sequence assembly methods are valuable for reconstructing genomes from shorter read fragments. Modern nucleic acid sequencing instruments produce quality scores associated with each reported base; however, these quality scores are not generally used as a core part of sequence assembly or alignment algorithms. Here, we leverage weighted de Bruijn graphs as graphical probability models representing the relative abundances and qualities of kmers within FASTQ-encoded observations. We then utilize these weighted de Bruijn graphs to identify alternate, higher-likelihood candidate sequences compared to the original observations, which are known to contain errors. By improving the original observations with these resampled paths, iteratively across increasing k-lengths, we can use this expectation-maximization approach to “polish” read sets from any sequencing technology according to the mutual information shared in the reads. We use this polishing approach to probabilistically correct simulated short- and long-read datasets of lower coverages and higher error rates than some algorithms can produce satisfactory assemblies for. We find that this approach corrects sequencing errors at rates that are able to produce error-free and nearly-error-free de Bruijn assembly graphs for simulated read-set challenges.

## 1 Introduction

The dominant paradigms in sequence assembly have evolved as developments in instrument technology, compute capacity, and algorithm capability have enabled new approaches. Short, highly accurate and low-cost reads are readily generated and used by most research groups [1]. Although short reads cannot bridge the repetitive regions of repeat-dense prokaryotic and eukaryotic genomes, their high accuracy enable reliable reconstruction of some prokaryotic genomes as well as most viral genomes and protein coding regions within genomes (exomes).

Long-read sequencing is available through a variety of commercially-available platforms that vary in accuracy, read length, and throughput. These longer reads have enabled researchers to reliably “close” circular genomes of prokaryotic organisms [2] and, more recently, to produce telomere-to-telomere [3] chromosomes of eukaryotic organisms. Combining multiple modalities of information, such as long-read sequencing, short-read sequencing, and physical proximity information such as Hi-C data, will generally produce the highest quality of assemblies [4–6]. Although long reads alone provide tremendous value in being able to span long repeats, long reads are more expensive and often fail to accurately report the length of those repeats — especially for homopolymeric and low-complexity repeats, leading to problems in downstream assemblies and outputs [7]. A self-contained, probability-based means of correcting random and systematic errors in FASTQ datasets, particularly long-read, low-fidelity datasets without the need of additional short-read data, would be an appealing solution for overcoming these limitations.

In developing our implementation of these algorithms, available within Mycelia (https://github.com/cjprybol/Mycelia), we sought to explore whether we could reuse a common data structure in short-read assemblers, the de Bruijn graph, as a probability model to perform error correction on long, error-rich reads in scenarios where current approaches fail due to some combination of sequence length, depth of coverage, and error rate. De Bruijn graphs, as used in sequence assembly, encode exact subsequences of *k* length (kmers) from the observed data and their exact *k −* 1 match and overlap between the suffix of parent nodes and the prefix of child nodes within the directed graph [8] [9]. This model enables existing graph libraries and graph analysis packages to be used for pathfinding, traversals, and graph simplification.

This error correction approach is analogous to the early spectral alignment correction method taken by the Euler assembler[10], updated to apply to noisy long reads and to utilize FASTQ quality scores to help further differentiate “solid” kmers that are likely to be valid and truly present relative to kmers that do not have enough supporting evidence in quantity, quality, or both to be considered valid. Short-read correction tools such as quake [11] and lighter [12] do utilize kmer quality information and offer faster correction methodologies through uses of statistical filters and algorithmic speedups, but it is not clear if they allow for insertion-deletion (indel) corrections, limiting their applicability toward modern 3rd generation data where indel errors are much more of a concern.

Salmela et al. [13] implemented a similar concept of using the mutual information in long reads to correct the reads, but relied on using overlap alignments between reads and required depths of coverage > 75×, at which point sufficient depth of coverage may exist to not need error correction. Furthermore, this is unique from the tf-idf approach taken by Canu, run-length encoding approach taken by shasta, minimizer approaches taken by miniasm, hifiasm, and mdbg, as well as the numerous alignment-based approaches compared in [14]. The most similar approach is Apollo, which has many overlapping aims to this proposal, although Apollo uses read alignments to an existing assembly to perform its correction rather than operating on raw reads directly [15].

Taken together, each individual component of our implementation has been implemented and proven to be useful by others at some point in time, but to the best of our knowledge, these features have not been combined to implement quality-aware, alignment-free error correction that is generalizable across technologies and data types.

Here, we implement a proof-of-concept workflow using these techniques and demonstrate that it rapidly resolves the majority of sequencing errors found in low-coverage, low-fidelity long-reads, producing perfect and near-perfect contigs from simulated challenge sets for which existing long-read assemblers fail to return any output at all.

Although the challenge sets utilized herein are test cases, they do highlight important limitations in valid edge cases where individuals are likely to want to produce an assembled output, but for which current best-in-class methods either fail to produce an output or produce an output with unclear accuracy expectations. The initial success of this approach is encouraging in that we find that this method, in conjunction with parameter searches and machine learning optimization, is likely to be applicable in more challenging and impactful conditions such as mixed-coverage metagenomic data, hybrid assembly of multiple data types, transcriptomics, and proteomics. To the best of our knowledge, no assembly package can handle all of these disparate data types and relative abundance conditions; however, many assembly packages that implement these capabilities rely on these shared features and data structures. The methods presented here are designed to be alphabet-agnostic and generalizable to other biological polymer assembly problems. They have been used to assemble RNA and amino-acid sequences (data not shown) via integration with existing software libraries in the Bio-Julia ecosystem https://github.com/BioJulia, and can be modified to handle arbitrary alphabets by utilizing fixed-length strings (known as n-grams but equivalent to their biological analog — k-mers), or other tokenized representations utilized across natural language processing and large language modeling applications [16]. When considering these alphabets, it is not necessarily appropriate to consider the quality of the reverse complement as has been done here with DNA, as RNA-seq data are often collected with strand specificity [17], and amino acids and natural (human) languages do not have reverse complements.

## 2 Results

### 2.1 Analysis of Kmer Frequency Power-Law Distributions in Reference Sequences

Inspired by the power-law distributions followed by natural languages known as Zipf’s law [18] [19] [20] [21], we examined whether there existed a k-length at which genomes and their genetic languages would also follow a power-law distribution. If so, this statistical property could be leveraged, either directly or indirectly, for the purposes of differentiating whether an observed kmer is more likely to be valid or an artifact.

In Figure 1 and Supplemental Figure 6, we counted the canonical kmers of the smallest reference quality genome from each major refseq category used in NCBI (virus, bacteria, archaea, fungi, plant, invertebrate, vertebrate-other, and vertebrate-mammalian), as well as the current human reference genome. We visualized the relationship between the number of times a kmer was observed and the number of kmers that were observed that many times. We found that while small genomes form stable log-log linear relationships in their kmer frequency spectra at k-lengths as short as 11, all evaluated genomes formed log-log linear relationships for *k ≥* 17.

**Fig. 1.**
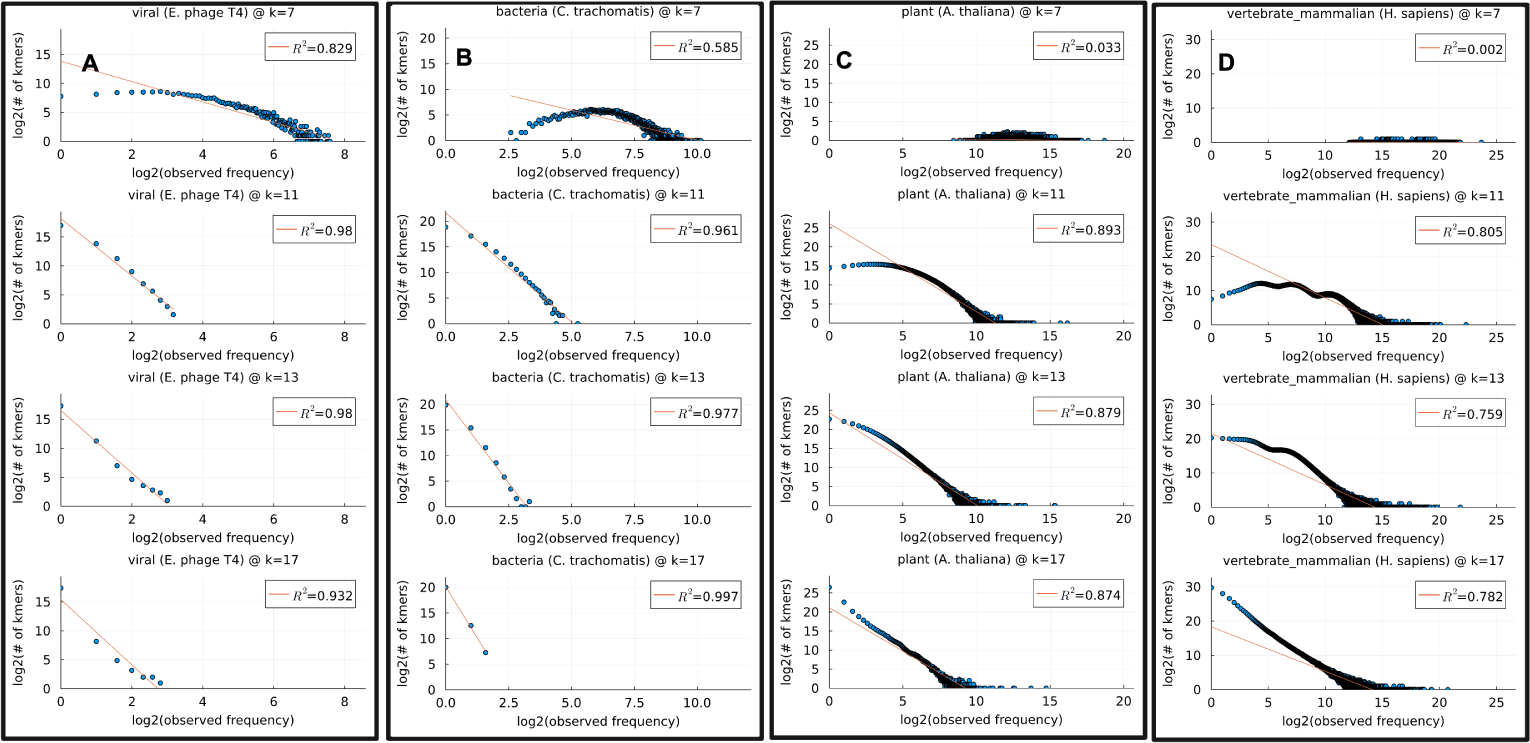
Identifying k-lengths at which log-log linear kmer frequency relationships appear. Genomes were selected across major taxonomic groups. Genomes form approximately log-log linear relationships between the kmer frequencies and the number of kmers at each frequency as the k-lengths increase. Beyond this initial value, the relationship holds until all kmers are unique. **A)** viruses **B)** bacteria **C)** plants **D)** human. A very long tail for the high-copy number kmers from repeats reduces the fit of the linear trendline relative to the other genomes shown.

### 2.2 Optimizing Kmer Lengths for Error Correction of Sequencing Data

Given our knowledge that above certain k-lengths, the kmers in genomes tend to form log-log linear frequency patterns that should be learnable by statistical models, we examined how to account for the additional noise incurred by the errors in the sequencing process. Although we could simulate the kmer frequency spectra at various error rates and learn from those observations, we took an alternate approach of trying to infer how large a k-value is needed for any given dataset to ensure that we do not saturate the total available kmer space. Our logic for this is made most clear by considering the extremes: Using a kmer length of 1, the kmer counts of a sequence yield an output equivalent to calculating %-GC, and using a k-length equal to the genome length, we will only have a single unique kmer to count. All of our ability to leverage kmer counts to differentiate true sequencing signal from artifact noise should come at the k-lengths between these two extremes.

Another feature to consider is the length at which kmers are expected, on average, to contain errors. If there is an error rate of 33%, then 1 in every 3 bases is expected to be an error. By *k* = 3, most kmers are expected to contain errors. For low-fidelity long reads in the realms of 10% error rates, we would not necessarily expect to see an error for k lengths of 7, however, for *k ≥* 11, sequencing artifacts in kmers would be expected. At higher accuracies and higher depths of coverage, larger k-lengths can be used without discarding too much information and still return an assembly - in fact, this is one of the core properties of analyzing sequencing data that short read assemblers leverage. When there are high error rates and low coverage, we likely need to use as short of a k-mer length as possible for the initial rounds of polishing. Because the number of erroneous kmers in a de Bruijn graph grows exponentially with k, correcting the majority of sequencing artifacts at k-lengths as short as possible is ideal to ensure that the computational complexity does not grow beyond what is reasonable to analyze.

For this work, we reuse the Michaelis-Menten equation [22], substituting standard reaction rate kinetics variables for our kmer variables, in both cases attempting to solve the saturation point of a system from a handful of samplings. In Figure 2, we use the canonical kmers for our counts, so theoretical maximum values are set to 4*^k^/*2, however, the same principles should apply to strand-specific data where complements may exist but should not necessarily be considered (RNA-seq) as well as alphabets without complement strands (e.g. amino acids).

**Fig. 2.**
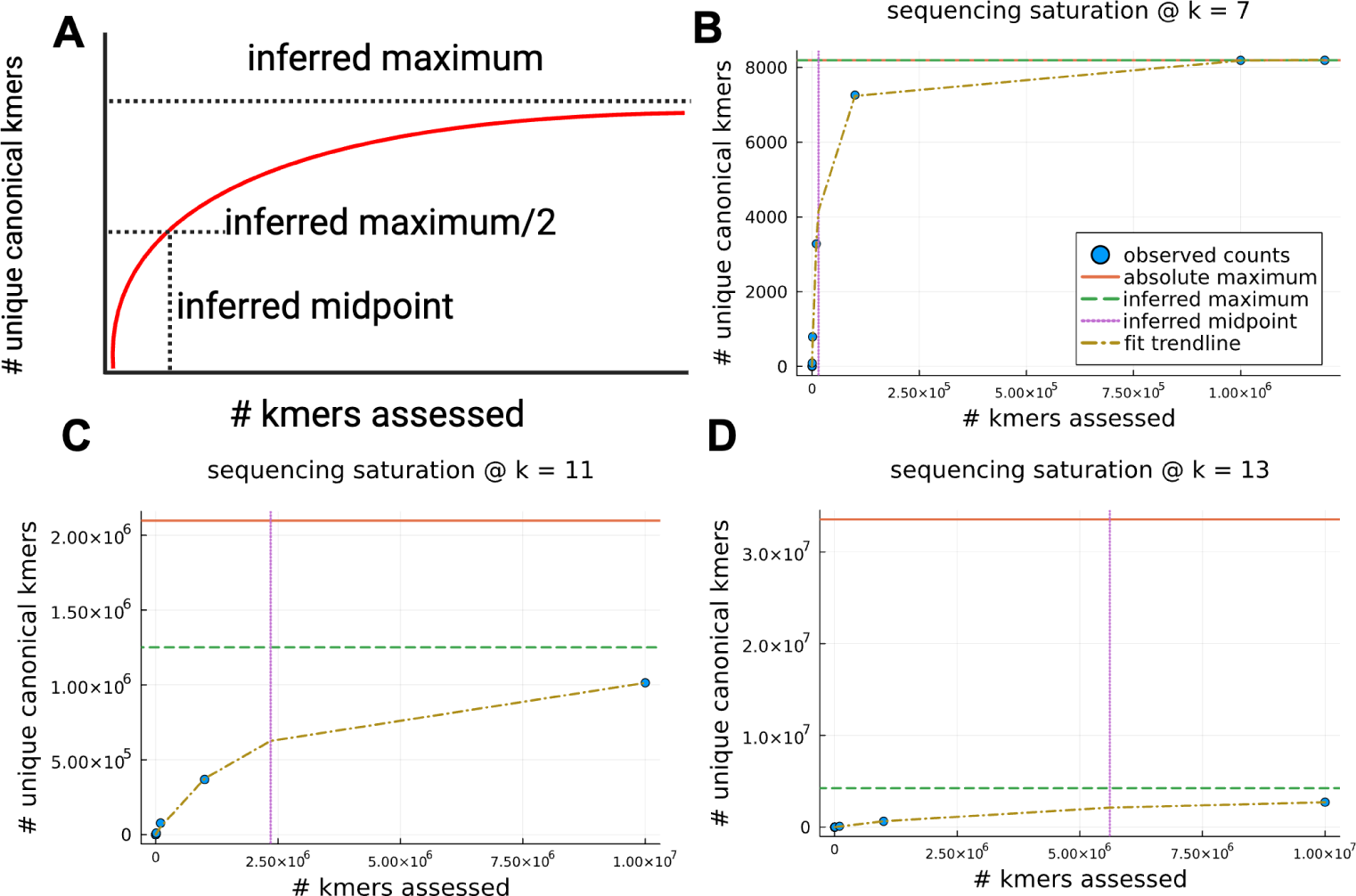
Estimating kmer saturation via Michaelis–Menten kinetics. A) A visual representation of the Michaelis-Menten equation, using variable substitution to show that the saturation limit can be estimated from the curve fit to observed data B) Evaluating canonical kmer saturation for a sample using k=7. The known upper limit of canonical kmers is quickly reached. C) Evaluation of the canonical kmer saturation for a sample using k=11. The known upper limit of canonical kmers is not estimated to be exceeded, but it is within a factor of 2, indicating that we may saturate all possible kmers with this dataset but it is unlikely. D) Evaluation of the canonical kmer saturation for a sample using k=13. We estimate that this sample will not reach saturation based on the kmers sampled so far. This is an appropriate kmer length to begin evaluating this sample.

### 2.3 Error Correction using Hidden Markov Models and Iterative Viterbi correction

With this information regarding kmer frequency distributions as they relate to real genomes and error profiles, we investigated whether de Bruijn graphs could be used as Markov models, and more specifically if, by extension, we could model the DNA sequencing process as a Hidden Markov Model (HMM) where the emission matrix of error probabilities is fit to the distribution of the actual error rates of the sequencing system used. This is similar to what Oxford Nanopore’s Medaka (https://github.com/nanoporetech/medaka) tool performs with neural networks trained on read pileups from alignments — using knowledge about the sequencing process to perform vendor and workflow-specific error correction to maximize the quality of the assembly outputs. If the sequencing process can be modeled as an HMM, then we should be able to use the Viterbi algorithm [23] to find the maximum likelihood path through the de Bruijn graph [24]. If we assume that this maximum likelihood path, which may or may not be the same as the actual path observed, is the actual sequence that went into the sequencer, then we can replace this higher likelihood sequence with the originally observed sequence greedily or in proportion to their relative likelihoods (1:10 if alternate path is 10x more likely).

If we perform this process iteratively, over increasingly long k-lengths, we are in effect performing an expectation maximization error correction based on the statistical patterns of the data. In Figure 3, we show a simulation of a short amplicon sequenced to *≈* 300x coverage with a 10% error rate. The initial bandage plot and kmer frequency spectra show lots of low-abundance kmers (represented as faint, whispy lines relative to the thick and colored kmers in the genome visualization) that are expected to be artifacts from sequencing errors. By using the kmer relative abundances to define state likelihoods, the relative frequencies of transitions between kmers to define transition likelihoods, and an estimate of the error rate (which one should be able to measure or determine from the instrument specifications) to define the emission matrix likelihoods, we were able to use the Viterbi algorithm to iteratively correct the majority of the sequencing errors to reconstruct a nearly error-free assembly directly from the longest unbranching paths formed by the error-corrected reads.

**Fig. 3.**
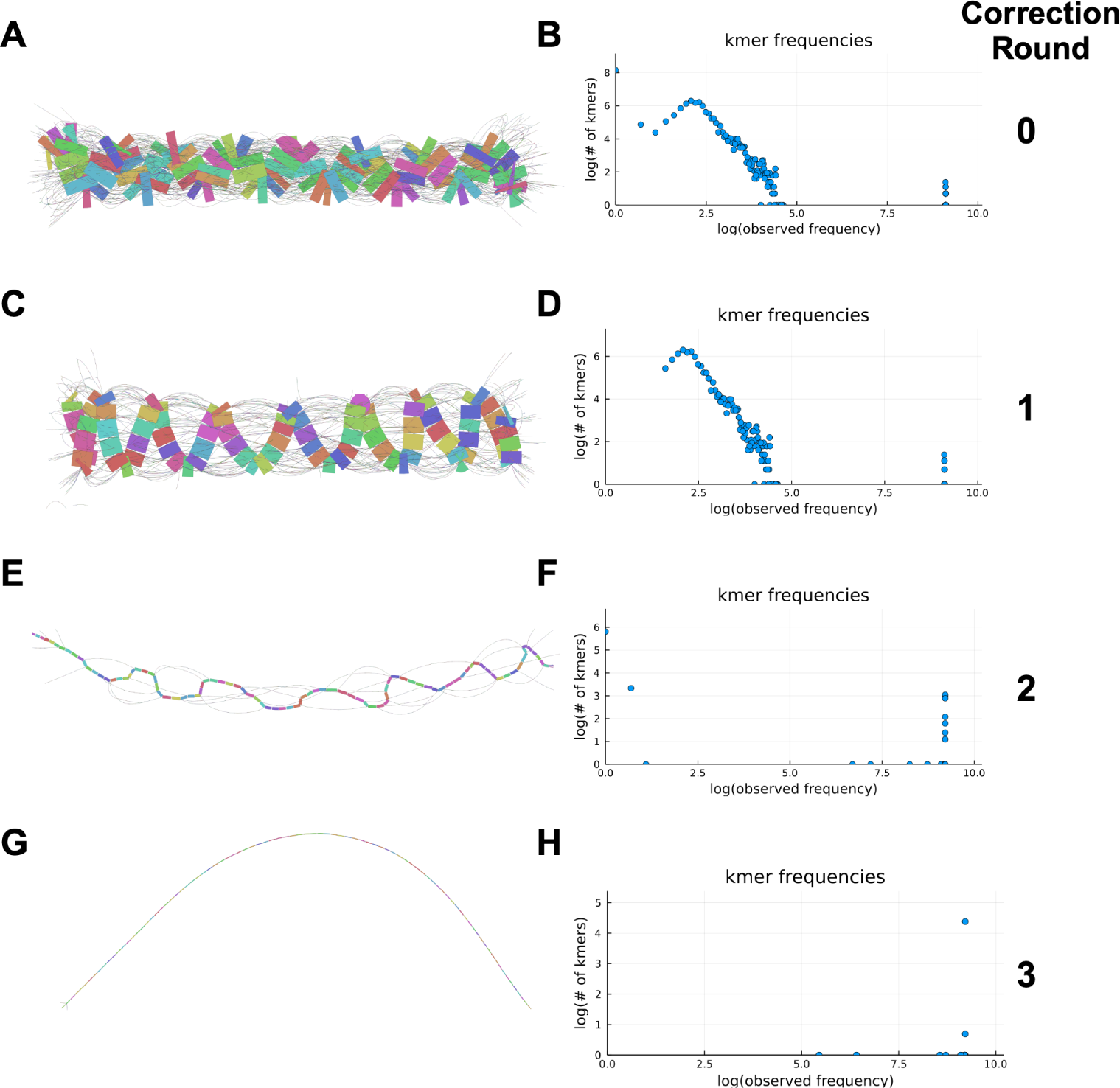
Iterative Viterbi correction reduces errors across iterative k-lengths. **A)** An uncorrected de Bruijn graph visualized with Bandage showing the diversity of low-coverage paths (faint lines) interconnected with the true genome (colored blocks) **B)** The canonical kmer frequency spectra for the uncorrected de Bruijn graph visualized in A. **C)** the de-Bruijn graph produced after 1 round of correction. **D)** The kmer frequency spectra after 1 round of correction. **E)** Same as C, but after the second round of correction. Note that as we lose relative coverage differences as errors are corrected to their maximum likelihood counterparts, the bands become thinner because they have fewer relative abundance differences as compared to the remaining low-coverage errors **F)** Same as D, but after the second round of correction. **G)** Same as C and E, but after the third round of correction. Very few errors remain now and a consensus assembly is obvious where it may not have been obvious initially **H)** Same as D and F, but after the third round of correction.

While this approach proved successful in our tests, the Viterbi algorithm is a very computationally intensive algorithm to compute when the Markov models and observation sequences have large numbers of states — both of which are true of genomics data. In an effort to carry forward the utility of a graph-based, probabilistic error-correction solution while avoiding the exhaustive calculations of the Viterbi algorithm, we sought to leverage the kmer counts and their quality scores to identify the kmers that are least supported in the dataset and only resample those paths. By avoiding the computations on the highest-likelihood regions of the data, we can focus our efforts on refining the regions of highest uncertainty in the raw data and the resulting de Bruijn graph.

### 2.4 Evaluating FASTQ Sequence Quality and Using Kmer-Based Approaches

In Figure 4, we show how the different ways of evaluating the quality of FASTQ sequences vary in their ability to reliably separate the signal from the noise. Although canonical kmer counts and mean FASTQ quality scores cannot easily differentiate the signal distribution of known true kmer states from known erroneous kmer states within this low-coverage, low-fidelity sample, when we share information across both strands and sum the cumulative quality of all bases observed in each kmer from either strand, we observe that our known true kmer population cleanly separates from the known artifact kmers such that one does not need sophisticated techniques to adequately set threshold boundaries that well differentiate likely error states from likely valid states. This is similar to the qmer count transformations used by [11].

**Fig. 4.**
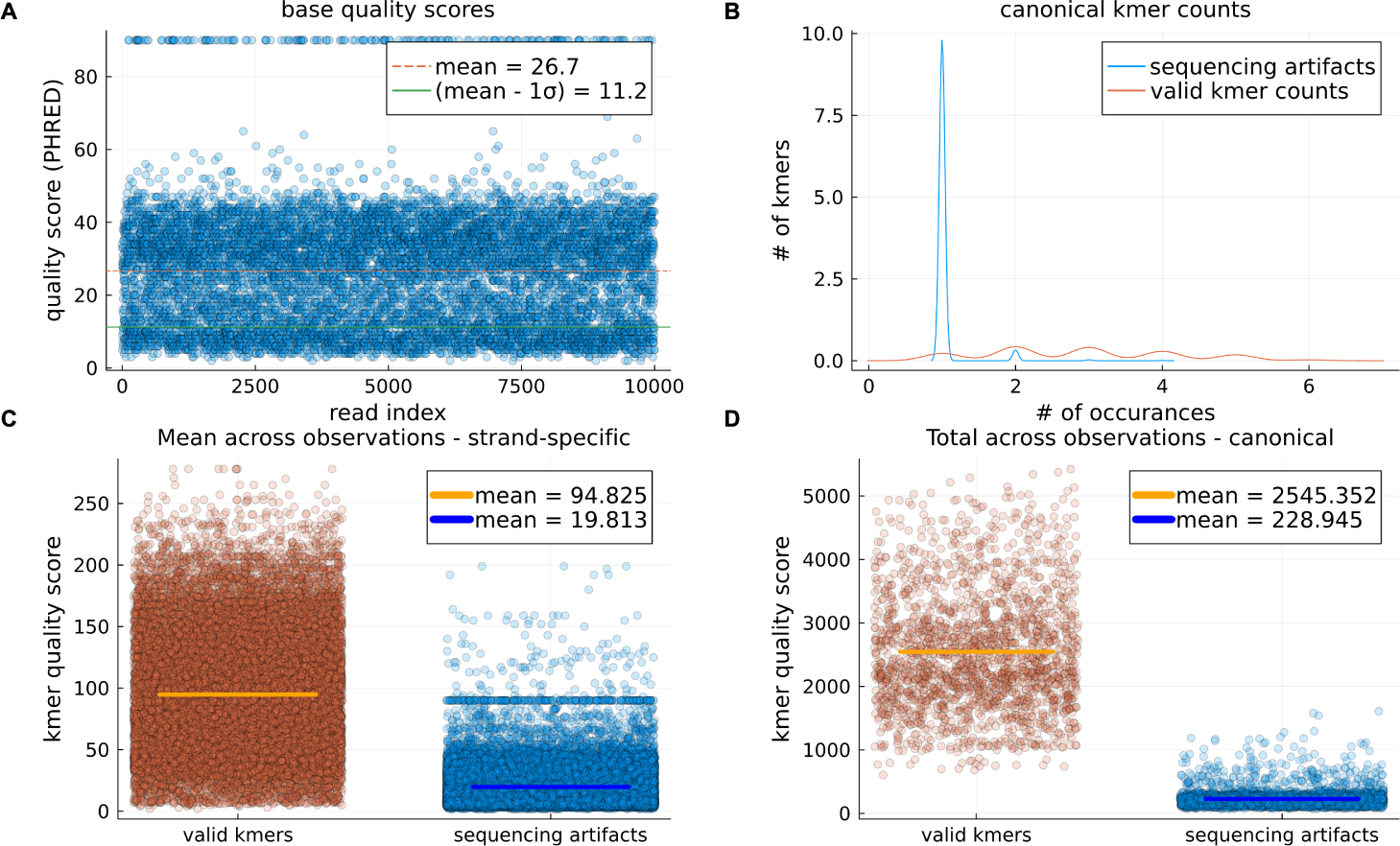
Integrating quality scores with kmer counts improves the separation between valid and artifact kmers over kmer counts alone. A) Base quality scores for a 10kb simulated genome sequenced at 10× depth of coverage using BadRead and the PacBio-hifi simulation model. B) Canonical kmer counts showing overlapping distributions that cannot allow us to readily separate valid kmers from artifact kmers based on coverage alone. C) If we sum the underlying FASTQ quality base scores of the individual kmers, and then average those sums for each kmer, we give ourselves additional dynamic range, but we do not improve our ability to distinguish erroneous vs valid kmers. D) Rather than averaging the sum of the underlying FASTQ base quality scores for each stand-specific kmer as in C, here we take the sum total of the underlying FASTQ base quality scores for kmers on either strand (we sum the weights of each stranded state while retaining the individual strands, rather than collapsing them into a single canonical state).

Given a set of kmers flagged as low quality according to the kmer quality values, or an analogous means of flagging states, we can build the de Bruijn graph from the observations and then identify candidate alternate paths, rank those candidate paths according to their relative likelihoods, and then replace our original observation with one of these candidate paths (which may be the same as the original) according to relative likelihoods. Performing this process iteratively over increasing k-lengths enables the correction of the majority of the most obvious errors at short k-lengths, preventing the computational requirements from expanding beyond what is tractable if we were to start at k-lengths that are larger than necessary.

### 2.5 Iterative FASTQ Polishing, Assembly Performance Evaluation, and Comparison with Existing Tools

In Figure 5, we iteratively polished the FASTQ observations with k-lengths of 11, 13, 17, 19, 23, 31, 53, and 89. At each round, a de Bruijn graph is created at the k-length, the kmers are assessed for their canonical cumulative quality scores, and then all kmers below the mean value are flagged for resampling. After performing this read correction, we assess the performance of this approach in its ability to support quality downstream assemblies by performing a simple coverage-based thresholding and longest path traversal of the connected components of the resulting de Bruijn graph, and report those walks as the assembly output. This is a rudimentary assembly method compared to what is performed by current best-in-class assemblers, which is intentional — [25–27] have shown in various ways that given a sufficiently long read length to span the longest exact repeats, we expect to easily reconstruct the genome if we can achieve error-free reads.

**Fig. 5.**
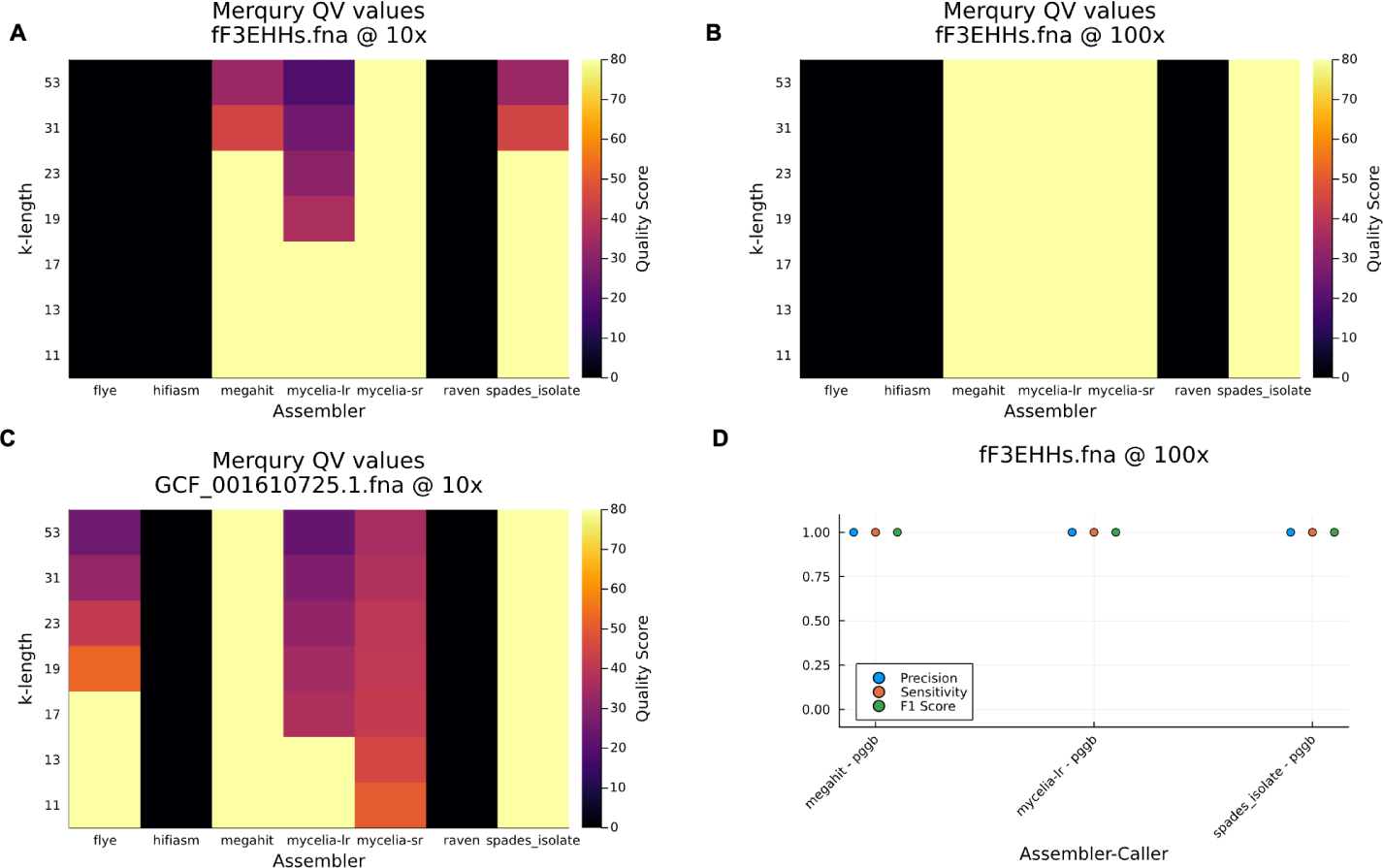
Mean-thresholded iterative de Bruijn error correction enables perfect and near-perfect reconstruction of low-coverage, low-fidelity long read data but alternate resampling thresholds required for longer genomes. A) the Merqury assembly quality values for various assemblers on a *≈* 1*kb* randomly simulated DNA sequence simulated using either ART (default parameters simulate Illumina HiSeq paired end reads) or BadRead (default parameters replicate moderate quality nanopore reads). Scores of 0, shown in black, represent missing outputs where the tool failed to produce any assembly. Scores of 80 are artificially capped - the original values were infinity, as the kmer spectra of the assembly were a perfect subset of the kmer spectra of the observed reads. We can see that the iterative correction implemented in Mycelia (see Code Availability in Methods) produced a perfect reconstruction from the input data, as reflected by the kmer compositions of the raw data and the assembly. All other methods either have some disagreement in the kmer spectra at longer k-lengths or fail to produce any output. B) Same as A, but with 100× coverages. All outputs produced had perfect Merqury quality values, and, as with A, all of the long-read assemblers failed to produce any output. C) with a RefSeq viral genome at 10× coverage, megaHIT produced a perfect Merqury quality value at all k lengths assessed. Flye is now generating output with longer input sequences and performs better than the naive mean-based resampling implemented by the correction algorithm. The correction algorithm is now actively falsifying the data as seen in that the short read assembly outputs fail to reflect the kmer spectra of the data at every k-length assessed. D) Direct pangenomic variant calling between *de novo* assemblies and original data shows perfect variant calling precision, sensitivity, and f1 scores for megaHIT, Spades, and the iterative correction approach on long-read data. Note that the tested tools (cactus-minigraph and PGGB) did not produce outputs for all assemblies attempted.

Neither hifiasm nor raven produced assemblies from these simulation sets - it is unclear whether the issue is primarily driven by the data quality (high error rates), data quantity (limited coverage), or size of the simulated sequences (short lengths).

The mean threshold resampling performed worse on the larger simulated sequence with short reads, indicating that starting at larger kmers and increasing the stringency for resampling with a better threshold would both likely be valuable to improve performance on short reads. Megahit produced quality outputs under all tested conditions, suggesting that the algorithm it uses may be worth further study and also that there may not be many substantive improvements left to be made algorithmically in the effective processing of short-read-only datasets.

While hifiasm and raven failed to produce outputs in our test cases, flye produced an output on the larger test set that was more accurate than what was produced from a simple traversal of Mycelia’s corrected reads, indicating that there are still refinements that are needed to make this probabilistic method competitive with best-in-class performance of existing approaches that have already been parameter optimized for real-world performance.

In summary, this novel approach leveraging weighted de Bruijn graphs and iterative error correction demonstrates the ability to accurately assemble genomes from low-coverage, high-error long reads than some current assemblers can analyze, and furthermore generates assembled contigs of comparable or higher quality than vendor-specific approaches while remaining vendor-agnostic.

**Fig. 6.**
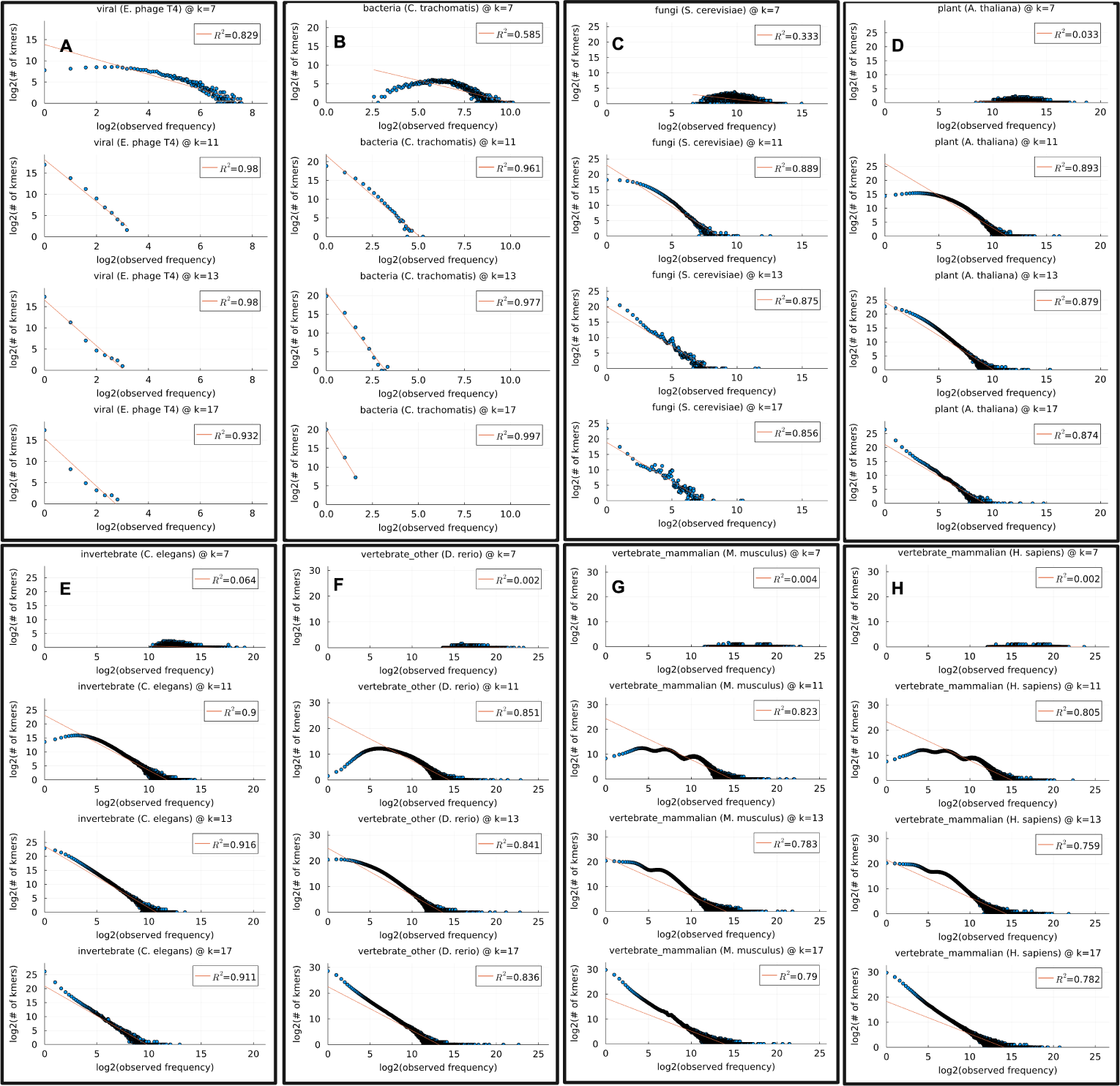
Identifying k-lengths at which log-log linear kmer frequency relationships appear. Genomes were selected across major taxonomic groups. Genomes form approximately log-log linear relationships between the kmer frequencies and the number of kmers at each frequency as the k-lengths increase. Beyond this initial value, the relationship holds until all kmers are unique. **A)** viruses **B)** bacteria **C)** fungi **D)** plants **E)** invertebrates **F)** vertebrates **G)** mammals **H)** human. A very long tail for the high-copy number kmers from repeats reduces the fit of the linear trendline relative to the other genomes shown.

## 3 Methods

### 3.1 NCBI Reference Genome Kmer Frequency Spectra

NCBI reference genomes were identified by downloading NCBI’s refseq assembly summary file (https://ftp.ncbi.nlm.nih.gov/genomes/refseq/assembly_summary_refseq.txt) and manually selecting a small reference quality genome in selected refseq groups (viral, bacteria, archaea, protozoa, fungi, plant, invertebrate, vertebrate other, vertebrate mammalian). Genomes were acquired using the NCBI datasets CLI tool, and then kmers were counted using JellyFish [28] using the --canonical flag for k-lengths of the prime numbers from 3 to 31, {3, 5, 7, 11, 13, 17, 19, 23, 29, 31}. For select genomes, we continued to increment k by primes until all kmers were unique as a means to spot check that the pattern continued to hold (data not shown). This approach became impractical beyond the genome sizes of bacteria, and for eukaryotic organisms we did not assess the consistency of the kmer saturation until we reached k lengths at which every kmer was unique.

### 3.2 Data Generation

The simulated reference sequences were generated using https://github.com/BioJulia/BioSequences.jl from the Julia package ecosystem [29]. NCBI reference genomes were identified by downloading the NCBI refseq assembly summary file (https://ftp.ncbi.nlm.nih.gov/genomes/refseq/assembly_summary_refseq.txt accessed on 20240121), subsetting to the records where

~~~
( genome rep = = ‘ f u l l ’ ) &&
( a s s e m b l y l e v e l in ( ‘ Chromosome ’, ‘ Complete Genome ’ ) ) &&
( r e f s e q c a t e g o r y in ( ‘ r e f e r e n c e genome ’, ‘ r e p r e s e n t a t i v e genome ’ )) &&
( genome s i ze > 10_000 )
~~~

Variants including substitutions, insertions, deletions, and inversions were simulated as follows. Each reference sequence was divided into 10 windows of equal length, and the type of variant was chosen such that substitutions were 10 times more likely than insertions, deletions, and/or inversions. The sizes of insertions, deletions, and inversions were chosen according to an exponential distribution set to

~~~
1/ s q r t ( window s i ze ) where window s i ze = 1/10 genome s i z e
~~~

Mutated versions of the reference sequences were created by updating the original sequences with the variants in the simulated VCFs using bcftools consensus.

Paired-end short reads were simulated using ART [30], with parameters `--seqSys HS25 --paired --len 150 --mflen 500 --sdev 10` and variable coverages of 10x and 100x. Long reads were simulated with coverages of 10x and 100x using Badread [31] with default parameters. Attempts were made to simulate single-ended long reads using NanoSim [32], but we were unable to get the software to run. We were able to run a NanoSim fork, NanoSim-h [33], however, the error profiles were out of date in relation to current sequencing technology and therefore were not used. Long reads were QC filtered and trimmed using https://github.com/rrwick/Filtlong with parameters `--min_mean_q 10 --keep_percent 90` and short reads were QC filtered and trimmed using Trim Galore [34] with default parameters.

### 3.3 De Bruijn Graph Construction and Data Model

De Bruijn graphs were constructed by parsing the FASTQ files using [35] and split into kmers using https://github.com/BioJulia/Kmers.jl using the appropriate strand-specific or canonical iteration utilities provided within. Graph data structures were constructed as directed graphs in the https://github.com/JuliaGraphs/MetaGraphs.jl package with key → value metadata information attached to nodes, edges, and graph, where appropriate. Graph traversals were performed using functions contained within Graphs.jl [36] and transformations between kmers and reconstructed sequences were performed using https://github.com/BioJulia/BioSequences.jl.

### 3.4 Viterbi Error Correction

Given a directed de Bruijn kmer graph, the shortest paths between states were indexed using the floyd warshall shortest paths function in Graphs.jl[36]. Edge probabilities were calculated on the basis of the relative likelihood of any given kmer going into each of the four possible downstream kmer states. The probabilities of these indexed shortest paths were calculated as the product of the relative edge likelihoods and indexed as well. The state probabilities were calculated as the relative likelihoods of each of the kmer states. During the evaluation of the initial kmer state, we calculate the edit distance between the observed initial kmer and each candidate kmer, and calculate the likelihoods of being in each state as *accuracy^n^ ^matches^ × error rate^n^ ^mismatches^*. The probabilities were transformed into log space to avoid underflow problems. Emission probabilities are calculated dynamically during the construction of the probability matrix used to keep track of the most likely path of hidden states that leads to each possible state at each time step, based on whether the considered path is a match, mismatch, insertion, or deletion relative to the observed path for each record kmer state being considered. This probability matrix is then used to find the maximum likelihood path through the HMM, known as the Viterbi path, which upon returning completes the Viterbi algorithm, a dynamic programming algorithm used to find the most likely sequence of hidden states in an HMM given a series of observed states [24]. When evaluating states, we consider the pre-indexed shortest path for the transition from the current state to each downstream state. The relative likelihoods of each candidate path are considered as the product of the relative likelihoods of the states and edge transitions on the path. We have a special case in how we handle transitions from a state to the same state. The transitions from a state to the same state are valid in de Bruijn graphs (AAAA → k=3 de Bruijn graph = *AAA* ⟲), but we want to be able to consider deletion events, which we model as remaining in the same node in the Viterbi backtracking matrix. We therefore consider a deletion event and a true state-state transition event (if such an event is supported by the raw data) and store the most likely of the two events in the Viterbi probaility matrix and backtracking table. The probabilities of deletions are modeled by *P* (*state*) × *P* (*deletions*). After performing these calculations stepwise through the observation, we identify the most likely path, which may or may not match the original path, and return it. Viterbi error correction code is documented in https://github.com/cjprybol/Eisenia.

### 3.5 Yen’s K Shortest Paths Correction

The intensiveness of the Viterbi algorithm calculations, particularly on large de Bruijn graphs that have an exponential growth on the runtime complexity of the Viterbi algorithm, motivated us to search for approximations that we believed would be able to give us most of the error-correction value provided by the Viterbi approach with the minimal amount of actual computation required. We recognize that considering every possible kmer against all other possible kmers does not take into account that most high-coverage kmers for high-quality data are valid and would not benefit the results if reconsidered. To address this, we investigated various visualizations of kmer counts, quality scores, and unbranching path scores, and ultimately arrived at the joint product of individual underlying kmer quality scores from the FASTQ file as the most useful metric to create maximum separation between high-coverage, high-quality “solid” kmers and low-coverage, low-quality “suspicious” kmers. We find this as a confirmation of the qmer score originally implemented by Quake [11]. We take a somewhat crude threshold of the mean kmer quality value, which causes us to resample some valid kmers but adequately covers all erroneous kmers in the unblinded simulations that we ran and evaluated. While more work is needed to optimize these settings, we found the model somewhat robust to threshold selection the initial evaluations, although considering more states than necessary is computationally wasteful and potentially counterproductive if we end up modifying correct observations into incorrect ones. The decision of whether the suspect stretch of the observation is left as is or corrected is based on the relative probabilities of the observed path as compared to the alternate paths identified by Yen’s algorithm for finding the K shortest paths [37] between the nearest upstream “solid” outbranching node and the nearest downstream “solid” inbranching node. If the same path is reselected, the underlying sequence and FASTQ scores of the FASTQ record are left intact. If an alternate path is selected, then the underlying sequence is replaced with the new sequence and the underlying FASTQ quality scores are resampled from the background distribution of quality scores. This process is performed on each read in a FASTQ file at a given k length, and then the modified reads are output. This process can be repeated as many times as desired, and we adopt the practice of increasing k every iteration rather than reanalyzing the same k length multiple times.

### 3.6 Assembly

Long-read and short-read assemblies were performed on the QC-trimmed and filtered reads, as correcting the most serious and systematic errors that are regularly prefiltered before assembly and read mapping is not an explicit goal of this work nor of other assembly and error-correction tools, to the best of our knowledge.

Short reads were assembled using `megaHIT` [38] [39] and `spades` [40] in the standard `--isolate` mode used for the assembly of isolate genomes. Long reads were assembled using `flye` [41] with the `--meta` [42] and `--nano-hq` parameters, `hifiasm` with the `--primary -l0` flags set as recommended for haploid isolates, and `raven` [43] was run using default parameters. The number of threads was set to use all available cores on the machine used to run the analysis, which was variable within the range of instance sizes available on the HPC systems used for the analysis.

### 3.7 Coverage-Thresholded Longest-Path Traversal

Our goal was to generate fasta contigs from the error-corrected reads as a means to determine whether the error correction could sufficiently correct enough errors to trivialize the assembly algorithm process to these steps. We also wanted to avoid any contributions made by error correction logic integrated into existing assemblers in our assessment of the accuracy, so while we could have simply fed our error correction reads into the other assembly methods tested, it would make it difficult to determine which tool was contributing what to the overall output quality. Therefore, we implemented a solution where we take the final set of corrected reads, build a de Bruijn graph, and walk the set of longest simple paths on each connected component to generate our assembled contigs.

### 3.8 Variant Calling

The variants were called by comparing the assembled output of each tool with the original unmodified reference genome (before insertion of the simulated variation and sequencing). This is meant to reflect the process by which many researchers perform reference-based variant calling, although this methodology first constructs a de novo assembly and compares that assembly against the reference genome rather than mapping the reads to the reference and then using a statistical method (bayesian, frequentist, neural network classifiers, etc.) to determine whether the differences between the reads and the reference are consistent enough to report a true variation call.

We ran both `PGGB` and `cactus-minigraph`. We had success with both pipelines, but for `cactus-minigraph`, only with reference genomes larger than those used for these simulations.

The precision, sensitivity, and F1 scores of the variation calls were evaluated using the eval command from RTG Tools [44].

### 3.9 Assembly Quality

As the data sets used for this work included a mix of simulated genomes (without regard to protein coding potential), gene-completeness-based genome accuracy tools such as [45] are not appropriate for all test cases. Furthermore, traditional N50-based genome contiguity scores are also not the target measure of interest, as our goal here is to produce the most accurate output, rather than the longest. While we directly compare the assembled outputs against the gold-standard references via the variant calling workflow, we wanted to evaluate what a reference-free quantitative score would report for these tools, as that would be useful in all situations, regardless of whether the genetic makeup of the input is known or unknown. We adopted the QV metric calculations from Merqury [46], computed directly from the JellyFish [28] kmer counts rather than by using Merqury directly.

### 3.10 Code Availability

Code is available at https://github.com/cjprybol/Mycelia and the notebooks and metadata specific to this work are within the https://github.com/cjprybol/Mycelia/tree/master/projects/variant-calling-benchmarking folder contained within that same repository.

## 4 Discussion

Given the algorithmic complexity of this approach relative to simpler and more efficient assembly graph cleaning procedures, the additional resource requirements of this methodology are unlikely to be justified in situations where high-accuracy approximations are sufficient. Furthermore, there is no guarantee that this algorithm will always produce the most accurate or the most useful genome output. It is designed to deliver the most likely sequence assembly given the data. Most sequence assemblers promise to offer only an output that is most consistent with the input data and their heuristic-based model of the world, which may or may not be documented and available to the researcher.

There are likely scenarios where sequence assembly would be appropriate, but the conditions defy the constraints and assumptions of this model. In situations where this is known to be true, the heuristics encoded in existing sequence assembly tools designed for those use cases are likely superior and should be used. In these situations, heuristic-based assemblers and neural network ‘black-box’ approaches will likely be superior to a state-model approach until we are able to learn the statistical patterns of those anomalies and adequately account for them.

The confidence scores obtained through this methodology apply to the genome, as well as to the variants, mutations, and alternate alleles that are detected. By calling variants by comparing the output assembly graphs directly against reference (pan)genomes using modern pangenome graph analysis tools such as VG [47] and ODGI [48], we eliminate the need to call variants based on read mapping, providing an independent and probabilistic means of evaluating the data toward equivalent objectives. By evaluating the expected impacts of the genome by genome differences, we can produce clinically meaningful variant reports in much the same way that we do now through workflows centered on mapping reads to a reference.

Another goal of this work was to develop an approach with a minimal set of parameters to tune. The list of “advanced” parameters that can be tuned in most read mapping and sequence assembly algorithms can be overwhelming to someone unfamiliar with how modifying each may affect the accuracy and interpretability of the results. Many of these parameters also suggest that the assemblers are doing more behind the scenes than their publications report. A core design philosophy around the approach presented here is to minimize the amount of user-provided information required to obtain a high-quality output (both to increase efficiency and to minimize user-introduced bias or error).

From the reads presented in a FASTQ file, we can gather two pieces of information reliably — the observed sequence and the confidence, at each observed base, of that sequence. FASTQ data files do not guarantee the availability of instrument type, and therefore we designed the approach to not require this, although if it is known, it can be provided.

In our efforts to produce ever more assemblies at ever greater accuracy, one must always consider the trade-offs of how much effort is put into generating data relative to how much effort is spent analyzing data. This study highlights the intricate nature of sequence assembly. We are fundamentally limited; sequencers do not directly reveal the true underlying DNA sequence. Instead, assembly algorithms must infer the original sequence from error-prone reads and their associated quality scores. The probabilistic approach implemented here underscores the value of both read-count and quality-score information in guiding this inference process.

By using information extracted directly from the raw data to evaluate read quality distributions, read length distributions, and kmer counts, we can build profiles of what kind of raw data distributions different types of dataset have and use that to automatically select the most appropriate parameters that have been identified by benchmarking on *in-silico* datasets.

## 5 Conclusion

Taken as a whole, this work demonstrates a novel, platform-agnostic, context-aware methodology for identifying high-confidence kmers and then using those confidence scores for probabilistic assemblies, which can then be used directly for variant calling without any read-mapping necessary.

Although the approach here is novel in many ways, it has not been studied to find optimal parameters for all common use cases. We present solutions found when correcting using the Viterbi algorithm and a Viterbi-inspired resampling method that perform comparably to or better than current state-of-the-art, but much work remains before this approach will be comparable to current best-in-class approaches on cutting-edge assembly problems. We believe that the approaches presented herein will rapidly gain ground as they are self-contained. This enables automated machine learning-based optimization of parameters and correction decisions within simulation and evaluation feedback loops. This feedback loop can be used by dynamic optimization programs to find ideal parameter sets for common use cases, and enable the software to auto-detect these common use cases based on the input data. This is all in the effort of helping make reference-ready genome assemblies more accessible to more individuals who may not have the skills or knowledge necessary to generate optimal quality assemblies on their own, in addition to providing a means of identifying whether a statistical approach to error correction and assembly will arrive at the same algorithms and heuristics that the community has identified and adopted based on acceptable results in practice.

### 5.1 Future Directions

Although we look forward to improving the algorithmic efficiency of the approach to enable it to scale to larger, more challenging genomic problems, it is likely that this approach will be best utilized in synergy with existing methodologies, rather than in replacement of them. Given the simplicity of kmer count-based analyses, it is appealing to continue to use them in high-coverage, high-accuracy contexts where they are currently working well. It would likely make sense to only consider quality scores when coverage is low enough that an obvious count-based separation cannot be found to separate the errors from the signal, although this distinction can be made on-the-fly within a tool rather than requiring entirely different tools to perform it. Furthermore, this separation by kmer quality scores can be done on the level of an entire assembly, or in the context of metagenomic work, high depth-of-coverage sequence bins can be assembled by existing, high-efficiency approaches with low depth-of-coverage sequence bins can be assembled using a more careful approach to maximize the utility of the limited read information as presented here.

Furthermore, because this approach is not vendor-specific in how it handles data, it can work as a hybrid assembler with mixed inputs of short and long reads, and we intend to explore this use case more. We look forward to obtaining high-quality pre-built models that apply to the most common use cases.

## Acknowledgements

Work at the Molecular Foundry was supported by the Office of Science, Office of Basic Energy Sciences, of the U.S. Department of Energy under Contract No. DE-AC02-05CH11231.

Some of the computing for this project was performed on the SCG3 Informatics Cluster. We would like to thank Stanford University and the Stanford Research Computing for providing computational resources and support that contributed to these research results.

Figures were prepared using BioRender under an academic license.

## Supplementary information

A detailed step-through of building a de Bruijn graph and then assessing the likelihood of an observation relative to that de Bruijn graph is shared in the supplemental materials within viterbi-correction.pdf. A detailed visualization of the stranded de Bruijn graphs created by this method, and the de-noising of the data and resulting assembly graph through the iterative error-correction procedure can be found within iterative-assembly.pdf.

## References

[1] Reuter, J.A., Spacek, D.V., Snyder, M.P.: High-throughput sequencing technologies. Molecular Cell 58(4), 586–597 (2015) 10.1016/j.molcel.2015.05.004

[2] Moss, E.L., Maghini, D.G., Bhatt, A.S.: Complete, closed bacterial genomes from microbiomes using nanopore sequencing. Nature Biotechnology 38(6), 701–707 (2020) 10.1038/s41587-020-0422-6. Accessed 2024-05-07

[3] Nurk, S., Koren, S., Rhie, A., Rautiainen, M., Bzikadze, A.V., Mikheenko, A., Vollger, M.R., Altemose, N., Uralsky, L., Gershman, A., Aganezov, S., Hoyt, S.J., Diekhans, M., Logsdon, G.A., Alonge, M., Antonarakis, S.E., Borchers, M., Bouffard, G.G., Brooks, S.Y., Caldas, G.V., Chen, N.-C., Cheng, H., Chin, C.-S., Chow, W., De Lima, L.G., Dishuck, P.C., Durbin, R., Dvorkina, T., Fiddes, I.T., Formenti, G., Fulton, R.S., Fungtammasan, A., Garrison, E., Grady, P.G.S., Graves-Lindsay, T.A., Hall, I.M., Hansen, N.F., Hartley, G.A., Haukness, M., Howe, K., Hunkapiller, M.W., Jain, C., Jain, M., Jarvis, E.D., Kerpedjiev, P., Kirsche, M., Kolmogorov, M., Korlach, J., Kremitzki, M., Li, H., Maduro, V.V., Marschall, T., McCartney, A.M., McDaniel, J., Miller, D.E., Mullikin, J.C., Myers, E.W., Olson, N.D., Paten, B., Peluso, P., Pevzner, P.A., Porubsky, D., Potapova, T., Rogaev, E.I., Rosenfeld, J.A., Salzberg, S.L., Schneider, V.A., Sedlazeck, F.J., Shafin, K., Shew, C.J., Shumate, A., Sims, Y., Smit, A.F.A., Soto, D.C., Sovíc, I., Storer, J.M., Streets, A., Sullivan, B.A., Thibaud-Nissen, F., Torrance, J., Wagner, J., Walenz, B.P., Wenger, A., Wood, J.M.D., Xiao, C., Yan, S.M., Young, A.C., Zarate, S., Surti, U., McCoy, R.C., Dennis, M.Y., Alexandrov, I.A., Gerton, J.L., O’Neill, R.J., Timp, W., Zook, J.M., Schatz, M.C., Eichler, E.E., Miga, K.H., Phillippy, A.M.: The complete sequence of a human genome. Science 376(6588), 44–53 (2022) 10.1126/science.abj6987. Accessed 2024-05-07

[4] Wick, R.R., Judd, L.M., Holt, K.E.: Assembling the perfect bacterial genome using Oxford Nanopore and Illumina sequencing. PLOS Computational Biology 19(3), 1010905 (2023) 10.1371/journal.pcbi.1010905. Accessed 2024-05-07

[5] Cheng, H., Concepcion, G.T., Feng, X., Zhang, H., Li, H.: Haplotype-resolved de novo assembly using phased assembly graphs with hifiasm. Nature Methods 18(2), 170–175 (2021) 10.1038/s41592-020-01056-5. Accessed 2024-03-24

[6] Nurk, S., Meleshko, D., Korobeynikov, A., Pevzner, P.A.: metaSPAdes: a new versatile metagenomic assembler. Genome Research 27(5), 824–834 (2017) 10.1101/gr.213959.116. Accessed 2024-05-07

[7] Watson, M., Warr, A.: Errors in long-read assemblies can critically affect protein prediction. Nature Biotechnology 37(2), 124–126 (2019) 10.1038/s41587-018-0004-z. Accessed 2024-03-24

[8] Zerbino, D.R., Birney, E.: Velvet: Algorithms for de novo short read assembly using de Bruijn graphs. Genome Research 18(5), 821–829 (2008) 10.1101/gr.074492.107. Accessed 2024-03-24

[9] Compeau, P.E.C., Pevzner, P.A., Tesler, G.: How to apply de Bruijn graphs to genome assembly. Nat. Biotechnol. 29(11), 987–991 (2011) 10.1038/nbt.2023

[10] Pevzner, P.A., Tang, H., Waterman, M.S.: An Eulerian path approach to DNA fragment assembly. Proceedings of the National Academy of Sciences 98(17), 9748–9753 (2001) 10.1073/pnas.171285098. Accessed 2024-05-07

[11] Kelley, D.R., Schatz, M.C., Salzberg, S.L.: Quake: quality-aware detection and correction of sequencing errors. Genome Biology 11(11), 116 (2010) 10.1186/gb-2010-11-11-r116. Accessed 2024-05-07

[12] Song, L., Florea, L., Langmead, B.: Lighter: fast and memory-efficient sequencing error correction without counting. Genome Biology 15(11), 509 (2014) 10.1186/s13059-014-0509-9. Accessed 2024-03-24

[13] Salmela, L., Walve, R., Rivals, E., Ukkonen, E.: Accurate self-correction of errors in long reads using de Bruijn graphs. Bioinformatics 33(6), 799–806 (2017) 10.1093/bioinformatics/btw321. Accessed 2024-04-04

[14] Lee, J.Y., Kong, M., Oh, J., Lim, J., Chung, S.H., Kim, J.-M., Kim, J.-S., Kim, K.-H., Yoo, J.-C., Kwak, W.: Comparative evaluation of Nanopore polishing tools for microbial genome assembly and polishing strategies for downstream analysis. Scientific Reports 11(1), 20740 (2021) 10.1038/s41598-021-00178-w. Accessed 2024-05-07

[15] Firtina, C., Kim, J.S., Alser, M., Senol Cali, D., Cicek, A.E., Alkan, C., Mutlu, O.: Apollo: a sequencing-technology-independent, scalable and accurate assembly polishing algorithm. Bioinformatics 36(12), 3669–3679 (2020) 10.1093/bioinformatics/btaa179. Accessed 2024-05-07

[16] Mullen, L.A., Benoit, K., Keyes, O., Selivanov, D., Arnold, J.: Fast, Consistent Tokenization of Natural Language Text. Journal of Open Source Software 3(23), 655 (2018) 10.21105/joss.00655. Accessed 2024-05-29

[17] Wang, Z., Gerstein, M., Snyder, M.: RNA-Seq: a revolutionary tool for transcriptomics. Nature Reviews Genetics 10(1), 57–63 (2009) 10.1038/nrg2484. Accessed 2024-05-29

[18] Bender, M.L., Gill, P.: The Genetic Code and Zipf’s Law. Current Anthropology 27(3), 280–283 (1986) 10.1086/203436. Accessed 2024-05-23

[19] Furusawa, C., Kaneko, K.: Zipf’s Law in Gene Expression. Physical Review Letters 90(8), 088102 (2003) 10.1103/PhysRevLett.90.088102. Accessed 2024-05-23

[20] Wang, J.D., Liu, H.-C., Tsai, J.J.P., Ng, K.-L.: Scaling Behavior of Maximal Repeat Distributions in Genomic Sequences:. International Journal of Cognitive Informatics and Natural Intelligence 2(3), 31–42 (2008) 10.4018/jcini.2008070103. Accessed 2024-05-23

[21] Kalankesh, L.R., Stevens, R., Brass, A.: The language of gene ontology: a Zipf’s law analysis. BMC Bioinformatics 13(1), 127 (2012) 10.1186/1471-2105-13-127. Accessed 2024-05-23

[22] Michaelis, L., Menten, M.L., Johnson, K.A., Goody, R.S.: The original Michaelis constant: translation of the 1913 Michaelis-Menten paper. Biochemistry 50(39), 8264–8269 (2011) 10.1021/bi201284u

[23] Viterbi, A.: Error bounds for convolutional codes and an asymptotically optimum decoding algorithm. IEEE Trans. Inf. Theory 13(2), 260–269 (1967) 10.1109/TIT.1967.1054010. Publisher: IEEE

[24] Rabiner, L.R.: A tutorial on hidden Markov models and selected applications in speech recognition. Proceedings of the IEEE 77(2), 257–286 (1989) 10.1109/5.18626. Accessed 2024-03-24

[25] Bresler, G., Bresler, M., Tse, D.: Optimal assembly for high throughput shotgun sequencing. BMC Bioinformatics 14 Suppl 5(Suppl 5), 18 (2013) 10.1186/1471-2105-14-S5-S18

[26] Shomorony, I., Courtade, T., Tse, D.: Do Read Errors Matter for Genome Assembly? Technical Report arXiv:1501.06194, arXiv (January 2015). arXiv:1501.06194 [cs, math, q-bio] type: article. http://arxiv.org/abs/1501.06194 Accessed 2022-07-08

[27] Kingsford, C., Schatz, M.C., Pop, M.: Assembly complexity of prokaryotic genomes using short reads. BMC Bioinformatics 11(1), 21 (2010) 10.1186/1471-2105-11-21. Accessed 2024-05-07

[28] Marçais, G., Kingsford, C.: A fast, lock-free approach for efficient parallel counting of occurrences of *k*-mers. Bioinformatics 27(6), 764–770 (2011) 10.1093/bioinformatics/btr011. Accessed 2024-05-13

[29] Bezanson, J., Edelman, A., Karpinski, S., Shah, V.B.: Julia: A Fresh Approach to Numerical Computing. SIAM Review 59(1), 65–98 (2017) 10.1137/141000671. Accessed 2024-05-10

[30] Huang, W., Li, L., Myers, J.R., Marth, G.T.: ART: a next-generation sequencing read simulator. Bioinformatics 28(4), 593–594 (2012) 10.1093/bioinformatics/btr708. Accessed 2024-05-10

[31] Wick, R.: Badread: simulation of error-prone long reads. Journal of Open Source Software 4(36), 1316 (2019) 10.21105/joss.01316. Accessed 2024-05-10

[32] Yang, C., Chu, J., Warren, R.L., Birol, I.: NanoSim: nanopore sequence read simulator based on statistical characterization. GigaScience 6(4) (2017) 10.1093/gigascience/gix010. Accessed 2024-05-14

[33] Břinda, K., Yang, C., Chu, J., Linthorst, J., Franus, W.: karel-brinda/NanoSim-H: NanoSim-H 1.1.0.4. [object Object] (2018). 10.5281/ZENODO.1341250. https://zenodo.org/record/1341250 Accessed 2024-05-10

[34] Andrews, S., Krueger, F., Segonds-Pichon, A., Biggins, L., et al.: Trim Galore. Trim Galore

[35] Jakob Nybo Nissen, Nicholas Bauer, Kevin Bonham, Ciarán O’Mara, Harry Scholes, Thomas Poulsen, Tim Holy, blaiseli, Dehann Fourie: BioJulia/FASTX.jl: v2.1.5. [object Object] (2024). 10.5281/ZENODO.3361839. https://zenodo.org/doi/10.5281/zenodo.3361839 Accessed 2024-05-14

[36] Fairbanks, J., Besançon, M., Simon, S., Hoffiman, J., Eubank, N., Karpinski, S.: JuliaGraphs/Graphs.jl: an optimized graphs package for the Julia programming language (2021). https://github.com/JuliaGraphs/Graphs.jl/

[37] Yen, J.Y.: An algorithm for finding shortest routes from all source nodes to a given destination in general networks. Quarterly of Applied Mathematics 27(4), 526–530 (1970) 10.1090/qam/253822. Accessed 2024-05-29

[38] Li, D., Liu, C.-M., Luo, R., Sadakane, K., Lam, T.-W.: MEGAHIT: an ultra-fast single-node solution for large and complex metagenomics assembly via succinct *de Bruijn* graph. Bioinformatics 31(10), 1674–1676 (2015) 10.1093/bioinformatics/btv033. Accessed 2024-03-24

[39] Li, D., Luo, R., Liu, C.-M., Leung, C.-M., Ting, H.-F., Sadakane, K., Yamashita, H., Lam, T.-W.: MEGAHIT v1.0: A fast and scalable metagenome assembler driven by advanced methodologies and community practices. Methods 102, 3–11 (2016) 10.1016/j.ymeth.2016.02.020

[40] Bankevich, A., Nurk, S., Antipov, D., Gurevich, A.A., Dvorkin, M., Kulikov, A.S., Lesin, V.M., Nikolenko, S.I., Pham, S., Prjibelski, A.D., Pyshkin, A.V., Sirotkin, A.V., Vyahhi, N., Tesler, G., Alekseyev, M.A., Pevzner, P.A.: SPAdes: A New Genome Assembly Algorithm and Its Applications to Single-Cell Sequencing. Journal of Computational Biology 19(5), 455–477 (2012) 10.1089/cmb.2012.0021. Accessed 2024-03-24

[41] Kolmogorov, M., Yuan, J., Lin, Y., Pevzner, P.A.: Assembly of long, error-prone reads using repeat graphs. Nature Biotechnology 37(5), 540–546 (2019) 10.1038/s41587-019-0072-8. Accessed 2024-03-24

[42] Kolmogorov, M., Bickhart, D.M., Behsaz, B., Gurevich, A., Rayko, M., Shin, S.B., Kuhn, K., Yuan, J., Polevikov, E., Smith, T.P.L., Pevzner, P.A.: metaFlye: scalable long-read metagenome assembly using repeat graphs. Nature Methods 17(11), 1103–1110 (2020) 10.1038/s41592-020-00971-x. Accessed 2024-05-14

[43] Vaser, R., Sĭkić, M.: Time- and memory-efficient genome assembly with Raven. Nature Computational Science 1(5), 332–336 (2021) 10.1038/s43588-021-00073-4. Accessed 2024-03-24

[44] Cleary, J.G., Braithwaite, R., Gaastra, K., Hilbush, B.S., Inglis, S., Irvine, S.A., Jackson, A., Littin, R., Rathod, M., Ware, D., Zook, J.M., Trigg, L., De La Vega, F.M.: Comparing Variant Call Files for Performance Benchmarking of Next-Generation Sequencing Variant Calling Pipelines (2015). 10.1101/023754. http://biorxiv.org/lookup/doi/10.1101/023754 Accessed 2024-05-14

[45] Simão, F.A., Waterhouse, R.M., Ioannidis, P., Kriventseva, E.V., Zdobnov, E.M.: BUSCO: assessing genome assembly and annotation completeness with single-copy orthologs. Bioinformatics 31(19), 3210–3212 (2015) 10.1093/bioinformatics/btv351. Accessed 2024-03-24

[46] Rhie, A., Walenz, B.P., Koren, S., Phillippy, A.M.: Merqury: reference-free quality, completeness, and phasing assessment for genome assemblies. Genome Biology 21(1), 245 (2020) 10.1186/s13059-020-02134-9. Accessed 2024-03-24

[47] Garrison, E., Sirén, J., Novak, A.M., Hickey, G., Eizenga, J.M., Dawson, E.T., Jones, W., Garg, S., Markello, C., Lin, M.F., Paten, B., Durbin, R.: Variation graph toolkit improves read mapping by representing genetic variation in the reference. Nat. Biotechnol. 36(9), 875–879 (2018) 10.1038/nbt.4227

[48] Guarracino, A., Heumos, S., Nahnsen, S., Prins, P., Garrison, E.: ODGI: understanding pangenome graphs. Bioinformatics, 308 (2022) 10.1093/bioinformatics/btac308. Accessed 2022-06-19

