## Supplemental Material 1 for "A Novel Approach for Accurate Sequence Assembly Using de Bruijn graphs"

The following example shows the full detail walkthrough calculations for utilizing the viterbi algorithm to determine the most likely path for a given sequence/read to take through an assembly graph. In this very simple example, the key is showing how even when a relatively high error rate is assumed, no corrections are made (true negative)

In [1]:

```
using Eisenia
using Random
using Dates
```

```
┌ Info: Recompiling stale cache file /Users/Cameron/.julia/compiled/v1.1/Eisenia.ji for Eisenia [top-level]
└ @ Base loading.jl:1184
```

In [2]:

```
L = 10
Random.seed!(L)
reference_sequence = randdnaseq(L)
reference_sequence_id = randstring(Int(round(log10(length(L)))+3))
reference_FASTA_record = FASTA.Record(reference_sequence_id, reference_sequence)
```

Out[2]:

```
BioSequences.FASTA.Record:
   identifier: Apx
  description: <missing>
     sequence: ACCAAACTAT
```

In [3]:

```
error_rate = 0.15
observations = [reference_FASTA_record]
```

Out[3]:

```
1-element Array{BioSequences.FASTA.Record,1}:
 BioSequences.FASTA.Record:
   identifier: Apx
  description: <missing>
     sequence: ACCAAACTAT
```

In [4]:

```
k = 1
canonical_kmers = collect(keys(Eisenia.count_canonical_kmers(observations, k)))
stranded_kmer_graph = Eisenia.build_stranded_kmer_graph(canonical_kmers, observations)
filename = reference_sequence_id * "." * replace(string(Dates.now()), ':' => '.') * ".svg"
Eisenia.plot_stranded_kmer_graph(stranded_kmer_graph, filename=filename)
HTML("""
<image src="$filename" width=50%>
""")
```

Out[4]:

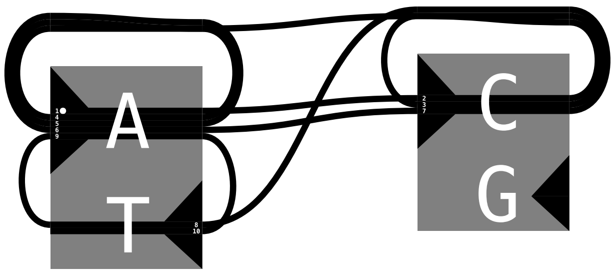


In [5]:

```
Eisenia.viterbi_maximum_likelihood_traversals(stranded_kmer_graph, error_rate = error_rate, verbosity="debug");
```

```
computing kmer counts...
computing kmer state likelihoods...
STATE LIKELIHOODS:
        kmer    count   likelihood
        A       5       0.5
        C       3       0.3
        G       0       0.0
        T       2       0.2
finding shortest paths between kmers...
        1       1       [1, 1]
        1       2       [1, 2]
        1       3       Int64[]
        1       4       [1, 4]
        2       1       [2, 1]
        2       2       [2, 2]
        2       3       Int64[]
        2       4       [2, 4]
        3       1       Int64[]
        3       2       Int64[]
        3       3       Int64[]
        3       4       Int64[]
        4       1       [4, 1]
        4       2       [4, 1, 2]
        4       3       Int64[]
        4       4       Int64[]
finding viterbi maximum likelihood paths for observed sequences...

evaluating sequence 1 of 1
        considering path state 1
                observed kmer A
                Initial state log likelihoods:
                        4-element Array{Float64,1}:
                           -0.16251892949777494
                           -2.4079456086518722
                         -Inf
                           -2.8134107167600364
        considering path state 2
                observed base C
                kmer log likelihoods
                        4×10 Array{Float64,2}:
                           -0.162519    -2.75279  0.0  0.0  0.0  0.0  0.0  0.0  0.0  0.0
                           -2.40795     -1.52901  0.0  0.0  0.0  0.0  0.0  0.0  0.0  0.0
                         -Inf         -Inf        0.0  0.0  0.0  0.0  0.0  0.0  0.0  0.0
                           -2.81341     -3.66908  0.0  0.0  0.0  0.0  0.0  0.0  0.0  0.0
                arrival paths
                        4×10 Array{Array{Int64,1},2}:
                         []  [1, 1]  #undef  #undef  #undef  #undef  #undef  #undef  #undef  #undef
                         []  [1, 2]  #undef  #undef  #undef  #undef  #undef  #undef  #undef  #undef
                         []  []      #undef  #undef  #undef  #undef  #undef  #undef  #undef  #undef
                         []  [1, 4]  #undef  #undef  #undef  #undef  #undef  #undef  #undef  #undef
        considering path state 3
                observed base C
                kmer log likelihoods
                        4×10 Array{Float64,2}:
                           -0.162519    -2.75279    -4.11928  0.0  0.0  0.0  0.0  0.0  0.0  0.0
                           -2.40795     -1.52901    -2.8955   0.0  0.0  0.0  0.0  0.0  0.0  0.0
                         -Inf         -Inf        -Inf        0.0  0.0  0.0  0.0  0.0  0.0  0.0
                           -2.81341     -3.66908    -5.03557  0.0  0.0  0.0  0.0  0.0  0.0  0.0
                arrival paths
                        4×10 Array{Array{Int64,1},2}:
                         []  [1, 1]  [2, 1]  #undef  #undef  #undef  #undef  #undef  #undef  #undef
                         []  [1, 2]  [2, 2]  #undef  #undef  #undef  #undef  #undef  #undef  #undef
                         []  []      []      #undef  #undef  #undef  #undef  #undef  #undef  #undef
                         []  [1, 4]  [2, 4]  #undef  #undef  #undef  #undef  #undef  #undef  #undef
        considering path state 4
                observed base A
                kmer log likelihoods
                        4×10 Array{Float64,2}:
                           -0.162519    -2.75279    -4.11928    -3.75117  0.0  0.0  0.0  0.0  0.0  0.0
                           -2.40795     -1.52901    -2.8955     -5.9966   0.0  0.0  0.0  0.0  0.0  0.0
                         -Inf         -Inf        -Inf        -Inf        0.0  0.0  0.0  0.0  0.0  0.0
                           -2.81341     -3.66908    -5.03557    -6.40206  0.0  0.0  0.0  0.0  0.0  0.0
                arrival paths
                        4×10 Array{Array{Int64,1},2}:
                         []  [1, 1]  [2, 1]  [2, 1]  #undef  #undef  #undef  #undef  #undef  #undef
                         []  [1, 2]  [2, 2]  [2, 2]  #undef  #undef  #undef  #undef  #undef  #undef
                         []  []      []      []      #undef  #undef  #undef  #undef  #undef  #undef
                         []  [1, 4]  [2, 4]  [2, 4]  #undef  #undef  #undef  #undef  #undef  #undef
        considering path state 5
                observed base A
                kmer log likelihoods
                        4×10 Array{Float64,2}:
                           -0.162519    -2.75279    -4.11928    -3.75117    -4.60683  0.0  0.0  0.0  0.0  0.0
                           -2.40795     -1.52901    -2.8955     -5.9966     -6.85226  0.0  0.0  0.0  0.0  0.0
                         -Inf         -Inf        -Inf        -Inf        -Inf        0.0  0.0  0.0  0.0  0.0
                           -2.81341     -3.66908    -5.03557    -6.40206    -7.25773  0.0  0.0  0.0  0.0  0.0
                arrival paths
                        4×10 Array{Array{Int64,1},2}:
                         []  [1, 1]  [2, 1]  [2, 1]  [1, 1]  #undef  #undef  #undef  #undef  #undef
                         []  [1, 2]  [2, 2]  [2, 2]  [1, 2]  #undef  #undef  #undef  #undef  #undef
                         []  []      []      []      []      #undef  #undef  #undef  #undef  #undef
                         []  [1, 4]  [2, 4]  [2, 4]  [1, 4]  #undef  #undef  #undef  #undef  #undef
        considering path state 6
                observed base A
                kmer log likelihoods
                        4×10 Array{Float64,2}:
                           -0.162519    -2.75279    -4.11928    -3.75117    -4.60683    -5.4625   0.0  0.0  0.0  0.0
                           -2.40795     -1.52901    -2.8955     -5.9966     -6.85226    -7.70793  0.0  0.0  0.0  0.0
                         -Inf         -Inf        -Inf        -Inf        -Inf        -Inf        0.0  0.0  0.0  0.0
                           -2.81341     -3.66908    -5.03557    -6.40206    -7.25773    -8.11339  0.0  0.0  0.0  0.0
                arrival paths
                        4×10 Array{Array{Int64,1},2}:
                         []  [1, 1]  [2, 1]  [2, 1]  [1, 1]  [1, 1]  #undef  #undef  #undef  #undef
                         []  [1, 2]  [2, 2]  [2, 2]  [1, 2]  [1, 2]  #undef  #undef  #undef  #undef
                         []  []      []      []      []      []      #undef  #undef  #undef  #undef
                         []  [1, 4]  [2, 4]  [2, 4]  [1, 4]  [1, 4]  #undef  #undef  #undef  #undef
        considering path state 7
```

```
		observed base C
		kmer log likelihoods
			4×10 Array{Float64,2}:
			  -0.162519   -2.75279   -4.11928   -3.75117   -4.60683   -5.4625    -8.05277  0.0  0.0  0.0
			  -2.40795    -1.52901   -2.8955    -5.9966    -6.85226   -7.70793   -6.82899  0.0  0.0  0.0
			 -Inf        -Inf       -Inf       -Inf       -Inf       -Inf       -Inf       0.0  0.0  0.0
			  -2.81341    -3.66908   -5.03557   -6.40206   -7.25773   -8.11339   -8.96906  0.0  0.0  0.0
		arrival paths
			4×10 Array{Array{Int64,1},2}:
			 []  [1, 1]  [2, 1]  [2, 1]  [1, 1]  [1, 1]  [1, 1]  #undef  #undef  #undef
			 []  [1, 2]  [2, 2]  [2, 2]  [1, 2]  [1, 2]  [1, 2]  #undef  #undef  #undef
			 []  []      []      []      []      []      []      #undef  #undef  #undef
			 []  [1, 4]  [2, 4]  [2, 4]  [1, 4]  [1, 4]  [1, 4]  #undef  #undef  #undef
	considering path state 8
		observed base T
		kmer log likelihoods
			4×10 Array{Float64,2}:
			  -0.162519   -2.75279   -4.11928   -3.75117   -4.60683   -5.4625    -8.05277   -9.41926  0.0  0.0
			  -2.40795    -1.52901   -2.8955    -5.9966    -6.85226   -7.70793   -6.82899   -9.93009  0.0  0.0
			 -Inf        -Inf       -Inf       -Inf       -Inf       -Inf       -Inf       -Inf       0.0  0.0
			  -2.81341    -3.66908   -5.03557   -6.40206   -7.25773   -8.11339   -8.96906   -8.60095  0.0  0.0
		arrival paths
			4×10 Array{Array{Int64,1},2}:
			 []  [1, 1]  [2, 1]  [2, 1]  [1, 1]  [1, 1]  [1, 1]  [2, 1]  #undef  #undef
			 []  [1, 2]  [2, 2]  [2, 2]  [1, 2]  [1, 2]  [1, 2]  [2, 2]  #undef  #undef
			 []  []      []      []      []      []      []      []      #undef  #undef
			 []  [1, 4]  [2, 4]  [2, 4]  [1, 4]  [1, 4]  [1, 4]  [2, 4]  #undef  #undef
	considering path state 9
		observed base A
		kmer log likelihoods
			4×10 Array{Float64,2}:
			  -0.162519   -2.75279   -4.11928   -3.75117   -4.60683   -5.4625    -8.05277   -9.41926   -9.45662  0.0
			  -2.40795    -1.52901   -2.8955    -5.9966    -6.85226   -7.70793   -6.82899   -9.93009  -12.5204   0.0
			 -Inf        -Inf       -Inf       -Inf       -Inf       -Inf       -Inf       -Inf       -Inf       0.0
			  -2.81341    -3.66908   -5.03557   -6.40206   -7.25773   -8.11339   -8.96906   -8.60095  -12.9258   0.0
		arrival paths
			4×10 Array{Array{Int64,1},2}:
			 []  [1, 1]  [2, 1]  [2, 1]  [1, 1]  [1, 1]  [1, 1]  [2, 1]  [4, 1]  #undef
			 []  [1, 2]  [2, 2]  [2, 2]  [1, 2]  [1, 2]  [1, 2]  [2, 2]  [1, 2]  #undef
			 []  []      []      []      []      []      []      []      []      #undef
			 []  [1, 4]  [2, 4]  [2, 4]  [1, 4]  [1, 4]  [1, 4]  [2, 4]  [1, 4]  #undef
	considering path state 10
		observed base T
		kmer log likelihoods
			4×10 Array{Float64,2}:
			  -0.162519   -2.75279   -4.11928   -3.75117   -4.60683   -5.4625    -8.05277   -9.41926   -9.45662  -12.0469
			  -2.40795    -1.52901   -2.8955    -5.9966    -6.85226   -7.70793   -6.82899   -9.93009  -12.5204   -12.5577
			 -Inf        -Inf       -Inf       -Inf       -Inf       -Inf       -Inf       -Inf       -Inf       -Inf
			  -2.81341    -3.66908   -5.03557   -6.40206   -7.25773   -8.11339   -8.96906   -8.60095  -12.9258   -11.2286
		arrival paths
			4×10 Array{Array{Int64,1},2}:
			 []  [1, 1]  [2, 1]  [2, 1]  [1, 1]  [1, 1]  [1, 1]  [2, 1]  [4, 1]  [1, 1]
			 []  [1, 2]  [2, 2]  [2, 2]  [1, 2]  [1, 2]  [1, 2]  [2, 2]  [1, 2]  [1, 2]
			 []  []      []      []      []      []      []      []      []      []
			 []  [1, 4]  [2, 4]  [2, 4]  [1, 4]  [1, 4]  [1, 4]  [2, 4]  [1, 4]  [1, 4]

	Inputs for viterbi maximum likelihood traversal evaluation:
		kmer log likelihoods
			4×10 Array{Float64,2}:
			  -0.162519   -2.75279   -4.11928   -3.75117   -4.60683   -5.4625    -8.05277   -9.41926   -9.45662  -12.0469
			  -2.40795    -1.52901   -2.8955    -5.9966    -6.85226   -7.70793   -6.82899   -9.93009  -12.5204   -12.5577
			 -Inf        -Inf       -Inf       -Inf       -Inf       -Inf       -Inf       -Inf       -Inf       -Inf
			  -2.81341    -3.66908   -5.03557   -6.40206   -7.25773   -8.11339   -8.96906   -8.60095  -12.9258   -11.2286
		kmer arrival paths
			4×10 Array{Array{Int64,1},2}:
			 []  [1, 1]  [2, 1]  [2, 1]  [1, 1]  [1, 1]  [1, 1]  [2, 1]  [4, 1]  [1, 1]
			 []  [1, 2]  [2, 2]  [2, 2]  [1, 2]  [1, 2]  [1, 2]  [2, 2]  [1, 2]  [1, 2]
			 []  []      []      []      []      []      []      []      []      []
			 []  [1, 4]  [2, 4]  [2, 4]  [1, 4]  [1, 4]  [1, 4]  [2, 4]  [1, 4]  [1, 4]
		edit distances
			4×10 Array{Int64,2}:
			 0  1  1  0  0  0  1  1  0  1
			 1  0  0  1  1  1  0  1  2  1
			 1  0  0  0  0  0  0  0  0  0
			 1  1  1  1  1  1  1  0  2  0
	observed sequence                 ACCAAACTAT
	maximum likelihood sequence       ACCAAACTAT
	maximum likelihood edit distance  0

DATASET STATISTICS:
	assumed error rate    15.0%
	total bases observed  10
	total edits accepted  0
	inferred error rate   0.0%
```
