## Supplemental Material 2 for "A Novel Approach for Accurate Sequence Assembly Using de Bruijn graphs"

### Iterative Assembly

The following notebook shows some small examples of the assembly approach that I have been developing that I call iterative assembly. The approach works by first building a de-bruijn assembly graph of the sequenced DNA. The approach then utilizes that assembly graph as a probablistic model to perform error correction via the viterbi algorithm on the reads in the dataset to reduce error rates and improve assembly accuracy. By correcting most errors using relatively short kmers, and then progressively working up to longer kmers that help resolve more repeats, it is possible to avoid a common issue during the genome assembly process where most kmers in the dataset are erroneous, as well as to overcome the sequencing accuracy limitations of long read nanopore sequencers.

It may be helpful to first start by looking at the initial assembly graph figure, which is laid out using a breadth-first search traversal. The deeper the graph, the more complexity (e.g. errors) are present. As the errors are removed during the iterative correction process, the graph flattens out and widens into a relatively easy to resolve assembly graph with few, if any, remaing errors.

In these examples, I am assuming the error rates to be 1/(k+1) (the default when the user does not provide a known error rate). Note the inferred error rate is generally closer to the true error rate than the assumed error rate, suggesting it is robust to imperfect settings.

In my experiments, I have found that iteratively assembling and correcting at a given k until no more corrections are performed and then incrementing k is the most effective way to remove errors. When two consecutive iterations do not produce any error corrections, the process can be halted and a final genome assembly produced (variants, coverage depths, closest known species, and other attributes can also be produced at this point).

In [1]:
```
using Eisenia
using Random
using Dates
```

# L10

In [2]:
```
L = 10
Random.seed!(L)
reference_sequence = randdnaseq(L)
reference_sequence_id = randstring(Int(round(log10(length(L)))+3))
reference_FASTA_record = FASTA.Record(reference_sequence_id, reference_sequence)
```

Out[2]:
```
BioSequences.FASTA.Record:
   identifier: Apx
  description: <missing>
     sequence: ACCAAACTAT
```

In [3]:
```
reverse_complement(reference_sequence)
```

Out[3]:
```
10nt DNA Sequence:
ATAGTTTGGT
```

In [4]:
```
error_rate = 0.15
n_sequences = 100
observations = [Eisenia.observe(reference_FASTA_record, error_rate=error_rate) for i in 1:n_sequences]
```

```
Out[4]: 100-element Array{BioSequences.FASTA.Record,1}:
         BioSequences.FASTA.Record:
           identifier: RO08
          description: <missing>
             sequence: ATCCAAACTA
         BioSequences.FASTA.Record:
           identifier: EUNI
          description: <missing>
             sequence: ATAGTTTTGGT
         BioSequences.FASTA.Record:
           identifier: vMBU
          description: <missing>
             sequence: ATAGTTTGG
         BioSequences.FASTA.Record:
           identifier: bbqN
          description: <missing>
             sequence: ACCAAACTAACT
         BioSequences.FASTA.Record:
           identifier: R0Fc
          description: <missing>
             sequence: ACCCAAACTAT
         BioSequences.FASTA.Record:
           identifier: kTEg
          description: <missing>
             sequence: ACCAACCAT
         BioSequences.FASTA.Record:
           identifier: j2mA
          description: <missing>
             sequence: ACCAACTAT
         BioSequences.FASTA.Record:
           identifier: nQi9
          description: <missing>
             sequence: ACCAACTAT
         BioSequences.FASTA.Record:
           identifier: SOR3
          description: <missing>
             sequence: AGTAGTTTGTGT
         BioSequences.FASTA.Record:
           identifier: 3N4E
          description: <missing>
             sequence: ATAGTTTGGT
         BioSequences.FASTA.Record:
           identifier: YIc7
          description: <missing>
             sequence: ATAGTTTGGT
         BioSequences.FASTA.Record:
           identifier: jC90
          description: <missing>
             sequence: AGCAACTAT
         BioSequences.FASTA.Record:
           identifier: Ddcm
          description: <missing>
             sequence: ACCTAACTAT
         ⋮
         BioSequences.FASTA.Record:
           identifier: JhfN
          description: <missing>
             sequence: ACCAAACTAT
         BioSequences.FASTA.Record:
           identifier: ijZ0
          description: <missing>
             sequence: ATAGTTTGGT
         BioSequences.FASTA.Record:
           identifier: pVdC
          description: <missing>
             sequence: ACCACACTAT
         BioSequences.FASTA.Record:
           identifier: eav3
          description: <missing>
             sequence: ACCCAAACTAT
         BioSequences.FASTA.Record:
           identifier: ebvn
          description: <missing>
             sequence: ACCCAACAT
         BioSequences.FASTA.Record:
           identifier: bIyC
          description: <missing>
             sequence: ACCTAACAT
         BioSequences.FASTA.Record:
           identifier: TEIx
          description: <missing>
             sequence: ACACAAACTAT
         BioSequences.FASTA.Record:
           identifier: 1hpo
          description: <missing>
             sequence: ACCAACTAT
         BioSequences.FASTA.Record:
           identifier: w9bv
          description: <missing>
             sequence: CATAGTTTGGT
         BioSequences.FASTA.Record:
           identifier: 1TLN
          description: <missing>
             sequence: ATAGTTTGGT
         BioSequences.FASTA.Record:
           identifier: wj9i
          description: <missing>
             sequence: ATAGTTTGGT
         BioSequences.FASTA.Record:
```

```
identifier: nYeF
description: <missing>
   sequence: ATAGTTTGGT
```

#### L10 starting @ k=5

In [5]:

```
k = 5
canonical_kmers = collect(keys(Eisenia.count_canonical_kmers(observations, k)))
stranded_kmer_graph = Eisenia.build_stranded_kmer_graph(canonical_kmers, observations)
filename = reference_sequence_id * "." * replace(string(Dates.now()), ':' => '.') * ".svg"
Eisenia.plot_stranded_kmer_graph(stranded_kmer_graph, filename=filename)
HTML("""
<image src="$filename" width=50%>
""")
```

Out[5]:

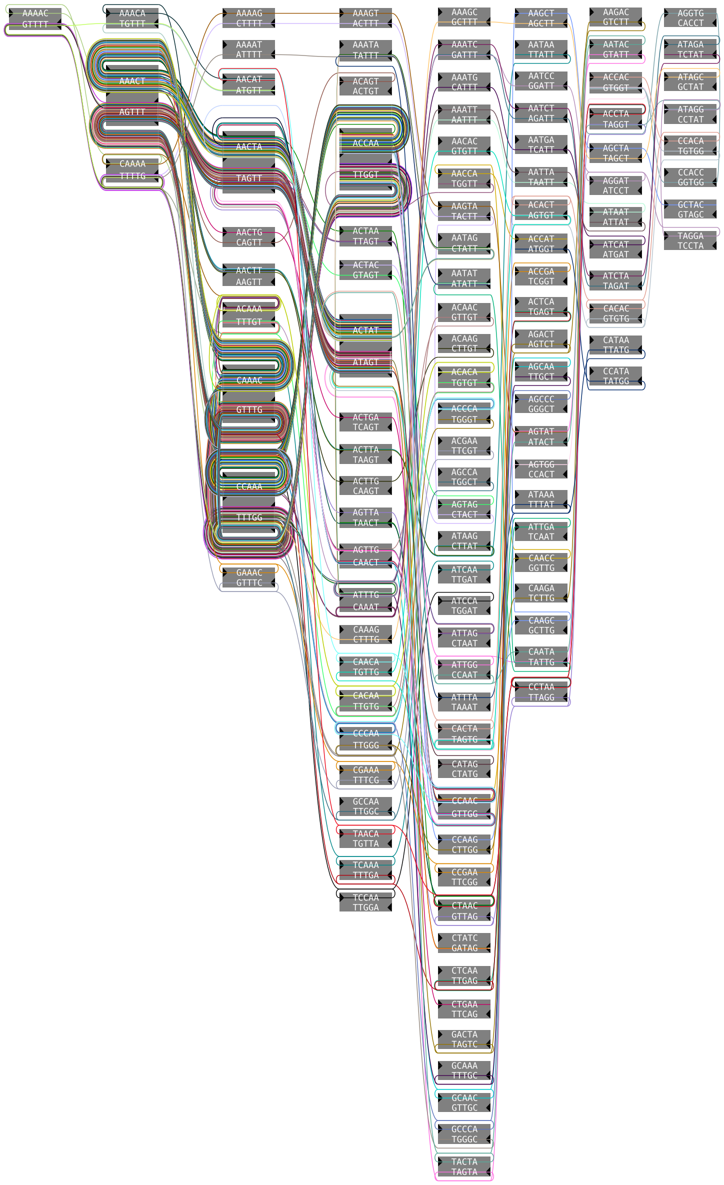


In [6]:

```
maximum_likelihood_observations = Eisenia.viterbi_maximum_likelihood_traversals(stranded_kmer_graph, verbosity="reads");
```

```
computing kmer counts...
computing kmer state likelihoods...
finding shortest paths between kmers...
finding viterbi maximum likelihood paths for observed sequences...

evaluating sequence 1 of 100
        observed sequence                 ATCCAAACTA
        maximum likelihood sequence       ACCAAACTA
        maximum likelihood edit distance  3

evaluating sequence 2 of 100
        observed sequence                 ATAGTTTTGGT
        maximum likelihood sequence       ATAGTTTGGT
        maximum likelihood edit distance  1

evaluating sequence 3 of 100
        observed sequence                 ATAGTTTGG
        maximum likelihood sequence       ATAGTTTGG
        maximum likelihood edit distance  0

evaluating sequence 4 of 100
        observed sequence                 ACCAAACTAACT
        maximum likelihood sequence       ACCAAACTAT
        maximum likelihood edit distance  3

evaluating sequence 5 of 100
        observed sequence                 ACCCAAACTAT
        maximum likelihood sequence       ACCAAACTAT
        maximum likelihood edit distance  2

evaluating sequence 6 of 100
        observed sequence                 ACCAACCAT
        maximum likelihood sequence       ACCAAACTAT
        maximum likelihood edit distance  2

evaluating sequence 7 of 100
        observed sequence                 ACCAACTAT
        maximum likelihood sequence       ACCAAACTAT
        maximum likelihood edit distance  1

evaluating sequence 8 of 100
        observed sequence                 ACCAACTAT
        maximum likelihood sequence       ACCAAACTAT
        maximum likelihood edit distance  1

evaluating sequence 9 of 100
        observed sequence                 AGTAGTTTGTGT
        maximum likelihood sequence       ATAGTTTGGT
        maximum likelihood edit distance  4

evaluating sequence 10 of 100
        observed sequence                 ATAGTTTGGT
        maximum likelihood sequence       ATAGTTTGGT
        maximum likelihood edit distance  0

evaluating sequence 11 of 100
        observed sequence                 ATAGTTTGGT
        maximum likelihood sequence       ATAGTTTGGT
        maximum likelihood edit distance  0

evaluating sequence 12 of 100
        observed sequence                 AGCAACTAT
        maximum likelihood sequence       ACCAAACTAT
        maximum likelihood edit distance  2

evaluating sequence 13 of 100
        observed sequence                 ACCTAACTAT
        maximum likelihood sequence       ACCAAACTAT
        maximum likelihood edit distance  1

evaluating sequence 14 of 100
        observed sequence                 ATAGTTTGGG
        maximum likelihood sequence       ATAGTTTGGT
        maximum likelihood edit distance  1

evaluating sequence 15 of 100
        observed sequence                 ACTACTTTTGGT
        maximum likelihood sequence       ATAGTTTGGT
        maximum likelihood edit distance  4

evaluating sequence 16 of 100
        observed sequence                 TAGTTTTGT
        maximum likelihood sequence       TAGTTTGGT
        maximum likelihood edit distance  1

evaluating sequence 17 of 100
        observed sequence                 ACCAAACTAT
        maximum likelihood sequence       ACCAAACTAT
        maximum likelihood edit distance  0

evaluating sequence 18 of 100
        observed sequence                 ACCAAACTAT
        maximum likelihood sequence       ACCAAACTAT
        maximum likelihood edit distance  0

evaluating sequence 19 of 100
        observed sequence                 ACAAACTAT
        maximum likelihood sequence       ACAAACTAT
        maximum likelihood edit distance  0
```

```
evaluating sequence 20 of 100
        observed sequence                 ACCAAACTAT
        maximum likelihood sequence       ACCAAACTAT
        maximum likelihood edit distance  0

evaluating sequence 21 of 100
        observed sequence                 ATAGTTTGGT
        maximum likelihood sequence       ATAGTTTGGT
        maximum likelihood edit distance  0

evaluating sequence 22 of 100
        observed sequence                 ACCAAACTT
        maximum likelihood sequence       ACCAAACT
        maximum likelihood edit distance  1

evaluating sequence 23 of 100
        observed sequence                 ATCAAACTTATT
        maximum likelihood sequence       ACCAAACTAT
        maximum likelihood edit distance  3

evaluating sequence 24 of 100
        observed sequence                 ACCAAACTAT
        maximum likelihood sequence       ACCAAACTAT
        maximum likelihood edit distance  0

evaluating sequence 25 of 100
        observed sequence                 ATAGTTTTGGT
        maximum likelihood sequence       ATAGTTTGGT
        maximum likelihood edit distance  1

evaluating sequence 26 of 100
        observed sequence                 ATAGTTTGGT
        maximum likelihood sequence       ATAGTTTGGT
        maximum likelihood edit distance  0

evaluating sequence 27 of 100
        observed sequence                 ATAATTTGGT
        maximum likelihood sequence       ATAGTTTGGT
        maximum likelihood edit distance  1

evaluating sequence 28 of 100
        observed sequence                 AACTTGT
        maximum likelihood sequence       AAACTAT
        maximum likelihood edit distance  3

evaluating sequence 29 of 100
        observed sequence                 ATAGTTTGGT
        maximum likelihood sequence       ATAGTTTGGT
        maximum likelihood edit distance  0

evaluating sequence 30 of 100
        observed sequence                 ATAGTATTGGT
        maximum likelihood sequence       ATAGTTTGGT
        maximum likelihood edit distance  2

evaluating sequence 31 of 100
        observed sequence                 ATATTGAGT
        maximum likelihood sequence       TAGTTTGGT
        maximum likelihood edit distance  3

evaluating sequence 32 of 100
        observed sequence                 ACAGTTTGGT
        maximum likelihood sequence       ATAGTTTGGT
        maximum likelihood edit distance  1

evaluating sequence 33 of 100
        observed sequence                 ATTAGTTTGGTT
        maximum likelihood sequence       ATAGTTTGGT
        maximum likelihood edit distance  3

evaluating sequence 34 of 100
        observed sequence                 ACCAAACTT
        maximum likelihood sequence       ACCAAACT
        maximum likelihood edit distance  1

evaluating sequence 35 of 100
        observed sequence                 ACCAAACTAT
        maximum likelihood sequence       ACCAAACTAT
        maximum likelihood edit distance  0

evaluating sequence 36 of 100
        observed sequence                 ATAGTGGT
        maximum likelihood sequence       TAGTTTGGT
        maximum likelihood edit distance  2

evaluating sequence 37 of 100
        observed sequence                 CTATTTGGT
        maximum likelihood sequence       TAGTTTGGT
        maximum likelihood edit distance  1

evaluating sequence 38 of 100
        observed sequence                 ACCAACTAT
        maximum likelihood sequence       ACCAAACTAT
        maximum likelihood edit distance  1

evaluating sequence 39 of 100
        observed sequence                 ATAGTTAGGT
        maximum likelihood sequence       ATAGTTTGGT
```

```
        maximum likelihood edit distance  1

evaluating sequence 40 of 100
        observed sequence                 GATAGTTTGGT
        maximum likelihood sequence       GATAGTTTGGT
        maximum likelihood edit distance  0

evaluating sequence 41 of 100
        observed sequence                 ATAGTTTGT
        maximum likelihood sequence       ATAGTTTGGT
        maximum likelihood edit distance  1

evaluating sequence 42 of 100
        observed sequence                 ACAAACTAT
        maximum likelihood sequence       ACAAACTAT
        maximum likelihood edit distance  0

evaluating sequence 43 of 100
        observed sequence                 ACCAAACTAT
        maximum likelihood sequence       ACCAAACTAT
        maximum likelihood edit distance  0

evaluating sequence 44 of 100
        observed sequence                 ACCAAACTAT
        maximum likelihood sequence       ACCAAACTAT
        maximum likelihood edit distance  0

evaluating sequence 45 of 100
        observed sequence                 ACCAAACAT
        maximum likelihood sequence       ACCAAACTAT
        maximum likelihood edit distance  1

evaluating sequence 46 of 100
        observed sequence                 ATAGTTTGGT
        maximum likelihood sequence       ATAGTTTGGT
        maximum likelihood edit distance  0

evaluating sequence 47 of 100
        observed sequence                 CCAAACT
        maximum likelihood sequence       CCAAACT
        maximum likelihood edit distance  0

evaluating sequence 48 of 100
        observed sequence                 ATAGGTGGT
        maximum likelihood sequence       ATAGTTTGGT
        maximum likelihood edit distance  2

evaluating sequence 49 of 100
        observed sequence                 ATAGTTGTGT
        maximum likelihood sequence       ATAGTTTGGT
        maximum likelihood edit distance  2

evaluating sequence 50 of 100
        observed sequence                 ATAGTTTGAGT
        maximum likelihood sequence       ATAGTTTGGT
        maximum likelihood edit distance  2

evaluating sequence 51 of 100
        observed sequence                 ACCGAAACTAT
        maximum likelihood sequence       ACCAAACTAT
        maximum likelihood edit distance  2

evaluating sequence 52 of 100
        observed sequence                 ACAAACTAT
        maximum likelihood sequence       ACAAACTAT
        maximum likelihood edit distance  0

evaluating sequence 53 of 100
        observed sequence                 ACCAAAACTAT
        maximum likelihood sequence       ACCAAACTAT
        maximum likelihood edit distance  1

evaluating sequence 54 of 100
        observed sequence                 AGTTTCGT
        maximum likelihood sequence       AGTTTGGT
        maximum likelihood edit distance  1

evaluating sequence 55 of 100
        observed sequence                 ATTAGTTTGGT
        maximum likelihood sequence       ATAGTTTGGT
        maximum likelihood edit distance  2

evaluating sequence 56 of 100
        observed sequence                 ATAGTTGGT
        maximum likelihood sequence       ATAGTTTGGT
        maximum likelihood edit distance  1

evaluating sequence 57 of 100
        observed sequence                 ACAAAAGTAT
        maximum likelihood sequence       ACCAAACTAT
        maximum likelihood edit distance  2

evaluating sequence 58 of 100
        observed sequence                 ATAGTGTGGT
        maximum likelihood sequence       ATAGTTTGGT
        maximum likelihood edit distance  1

evaluating sequence 59 of 100
        observed sequence                 ACCAAACTAT
```

```
        maximum likelihood sequence       ACCAAACTAT
        maximum likelihood edit distance  0

evaluating sequence 60 of 100
        observed sequence                 ACCAAACTAT
        maximum likelihood sequence       ACCAAACTAT
        maximum likelihood edit distance  0

evaluating sequence 61 of 100
        observed sequence                 ATCATTTGCT
        maximum likelihood sequence       ATAGTTTGGT
        maximum likelihood edit distance  3

evaluating sequence 62 of 100
        observed sequence                 ATATTTGGT
        maximum likelihood sequence       TAGTTTGGT
        maximum likelihood edit distance  1

evaluating sequence 63 of 100
        observed sequence                 ATAGTTGGT
        maximum likelihood sequence       ATAGTTTGGT
        maximum likelihood edit distance  1

evaluating sequence 64 of 100
        observed sequence                 ACCAAGCTAC
        maximum likelihood sequence       ACCAAACTAT
        maximum likelihood edit distance  2

evaluating sequence 65 of 100
        observed sequence                 ATAGATTTGGCT
        maximum likelihood sequence       ATAGTTTGGT
        maximum likelihood edit distance  4

evaluating sequence 66 of 100
        observed sequence                 ATAGTTTGGT
        maximum likelihood sequence       ATAGTTTGGT
        maximum likelihood edit distance  0

evaluating sequence 67 of 100
        observed sequence                 ACCAAACTAT
        maximum likelihood sequence       ACCAAACTAT
        maximum likelihood edit distance  0

evaluating sequence 68 of 100
        observed sequence                 GATAGTTTGGT
        maximum likelihood sequence       GATAGTTTGGT
        maximum likelihood edit distance  0

evaluating sequence 69 of 100
        observed sequence                 ATAGGATTTGT
        maximum likelihood sequence       ATAGTTTGGT
        maximum likelihood edit distance  4

evaluating sequence 70 of 100
        observed sequence                 CCAAACTGAA
        maximum likelihood sequence       CCAAACTAT
        maximum likelihood edit distance  3

evaluating sequence 71 of 100
        observed sequence                 ACCAAATTAT
        maximum likelihood sequence       ACCAAACTAT
        maximum likelihood edit distance  1

evaluating sequence 72 of 100
        observed sequence                 ATAGTGTTGGT
        maximum likelihood sequence       ATAGTTTGGT
        maximum likelihood edit distance  2

evaluating sequence 73 of 100
        observed sequence                 ACCAAACTAT
        maximum likelihood sequence       ACCAAACTAT
        maximum likelihood edit distance  0

evaluating sequence 74 of 100
        observed sequence                 ATAGTTTGGT
        maximum likelihood sequence       ATAGTTTGGT
        maximum likelihood edit distance  0

evaluating sequence 75 of 100
        observed sequence                 ACACAAACTATT
        maximum likelihood sequence       ACCAAACTAT
        maximum likelihood edit distance  4

evaluating sequence 76 of 100
        observed sequence                 ACCAAACTA
        maximum likelihood sequence       ACCAAACTA
        maximum likelihood edit distance  0

evaluating sequence 77 of 100
        observed sequence                 ATAGTCTTGGGT
        maximum likelihood sequence       ATAGTTTGGT
        maximum likelihood edit distance  3

evaluating sequence 78 of 100
        observed sequence                 ACCAAATCTAT
        maximum likelihood sequence       ACCAAACTAT
        maximum likelihood edit distance  2

evaluating sequence 79 of 100
```

```
        observed sequence                 ATGTTTTGGT
        maximum likelihood sequence       ATAGTTTGGT
        maximum likelihood edit distance  2

evaluating sequence 80 of 100
        observed sequence                 ATAGCTTTGGT
        maximum likelihood sequence       ATAGTTTGGT
        maximum likelihood edit distance  2

evaluating sequence 81 of 100
        observed sequence                 ATGTTTGGT
        maximum likelihood sequence       TAGTTTGGT
        maximum likelihood edit distance  2

evaluating sequence 82 of 100
        observed sequence                 ACCAAACTAT
        maximum likelihood sequence       ACCAAACTAT
        maximum likelihood edit distance  0

evaluating sequence 83 of 100
        observed sequence                 ATAGTTGGT
        maximum likelihood sequence       ATAGTTTGGT
        maximum likelihood edit distance  1

evaluating sequence 84 of 100
        observed sequence                 AGCCCAAACTT
        maximum likelihood sequence       ACCAAACT
        maximum likelihood edit distance  6

evaluating sequence 85 of 100
        observed sequence                 ATATTTATGGT
        maximum likelihood sequence       ATAGTTTGGT
        maximum likelihood edit distance  3

evaluating sequence 86 of 100
        observed sequence                 ACTATTTTGGGC
        maximum likelihood sequence       ATAGTTTGGT
        maximum likelihood edit distance  4

evaluating sequence 87 of 100
        observed sequence                 ACCAATACTATT
        maximum likelihood sequence       ACCAAACTAT
        maximum likelihood edit distance  3

evaluating sequence 88 of 100
        observed sequence                 ACCAAACTTAT
        maximum likelihood sequence       ACCAAACTAT
        maximum likelihood edit distance  1

evaluating sequence 89 of 100
        observed sequence                 ACCAAACTAT
        maximum likelihood sequence       ACCAAACTAT
        maximum likelihood edit distance  0

evaluating sequence 90 of 100
        observed sequence                 ATAGTTTGGT
        maximum likelihood sequence       ATAGTTTGGT
        maximum likelihood edit distance  0

evaluating sequence 91 of 100
        observed sequence                 ACCACACTAT
        maximum likelihood sequence       ACCAAACTAT
        maximum likelihood edit distance  1

evaluating sequence 92 of 100
        observed sequence                 ACCCAAACTAT
        maximum likelihood sequence       ACCAAACTAT
        maximum likelihood edit distance  2

evaluating sequence 93 of 100
        observed sequence                 ACCCAACAT
        maximum likelihood sequence       ACCAAACTAT
        maximum likelihood edit distance  2

evaluating sequence 94 of 100
        observed sequence                 ACCTAACAT
        maximum likelihood sequence       ACCAAACTAT
        maximum likelihood edit distance  2

evaluating sequence 95 of 100
        observed sequence                 ACACAAACTAT
        maximum likelihood sequence       ACCAAACTAT
        maximum likelihood edit distance  3

evaluating sequence 96 of 100
        observed sequence                 ACCAACTAT
        maximum likelihood sequence       ACCAAACTAT
        maximum likelihood edit distance  1

evaluating sequence 97 of 100
        observed sequence                 CATAGTTTGGT
        maximum likelihood sequence       ATAGTTTGGT
        maximum likelihood edit distance  2

evaluating sequence 98 of 100
        observed sequence                 ATAGTTTGGT
        maximum likelihood sequence       ATAGTTTGGT
        maximum likelihood edit distance  0
```

```
evaluating sequence 99 of 100
        observed sequence                 ATAGTTTGGT
        maximum likelihood sequence       ATAGTTTGGT
        maximum likelihood edit distance  0

evaluating sequence 100 of 100
        observed sequence                 ATAGTTTGGT
        maximum likelihood sequence       ATAGTTTGGT
        maximum likelihood edit distance  0

DATASET STATISTICS:
        assumed error rate    16.666666666666664%
        total bases observed  1004
        total edits accepted  135
        inferred error rate   13.44621513944223%
```

In [7]:

```
canonical_kmers = collect(keys(Eisenia.count_canonical_kmers(maximum_likelihood_observations, k)))
stranded_kmer_graph = Eisenia.build_stranded_kmer_graph(canonical_kmers, maximum_likelihood_observations)
filename = reference_sequence_id * "." * replace(string(Dates.now()), ':' => '.') * ".svg"
Eisenia.plot_stranded_kmer_graph(stranded_kmer_graph, filename=filename)
HTML("""
<image src="$filename" width=50%>
""")
```

Out[7]:

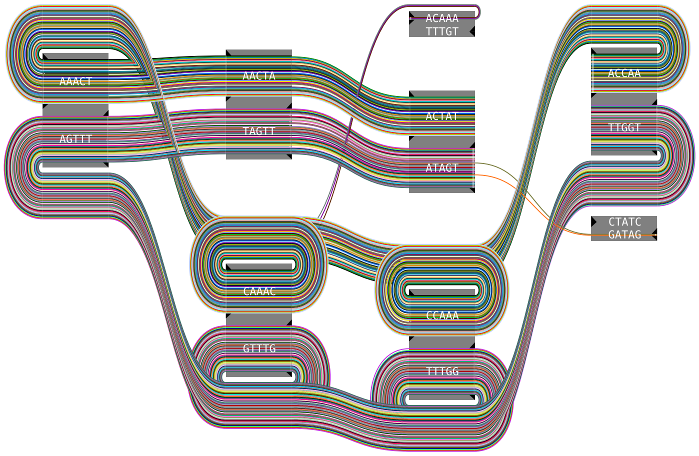


In [8]:

```
maximum_likelihood_observations = Eisenia.viterbi_maximum_likelihood_traversals(stranded_kmer_graph, error_rate = error_rate, verbosity="reads");
```

```
computing kmer counts...
computing kmer state likelihoods...
finding shortest paths between kmers...
finding viterbi maximum likelihood paths for observed sequences...

evaluating sequence 1 of 100
        observed sequence                 ACCAAACTA
        maximum likelihood sequence       ACCAAACTA
        maximum likelihood edit distance  0

evaluating sequence 2 of 100
        observed sequence                 ATAGTTTGGT
        maximum likelihood sequence       ATAGTTTGGT
        maximum likelihood edit distance  0

evaluating sequence 3 of 100
        observed sequence                 ATAGTTTGG
        maximum likelihood sequence       ATAGTTTGG
        maximum likelihood edit distance  0

evaluating sequence 4 of 100
        observed sequence                 ACCAAACTAT
        maximum likelihood sequence       ACCAAACTAT
        maximum likelihood edit distance  0

evaluating sequence 5 of 100
        observed sequence                 ACCAAACTAT
        maximum likelihood sequence       ACCAAACTAT
        maximum likelihood edit distance  0

evaluating sequence 6 of 100
        observed sequence                 ACCAAACTAT
        maximum likelihood sequence       ACCAAACTAT
        maximum likelihood edit distance  0

evaluating sequence 7 of 100
        observed sequence                 ACCAAACTAT
        maximum likelihood sequence       ACCAAACTAT
        maximum likelihood edit distance  0

evaluating sequence 8 of 100
        observed sequence                 ACCAAACTAT
        maximum likelihood sequence       ACCAAACTAT
        maximum likelihood edit distance  0

evaluating sequence 9 of 100
        observed sequence                 ATAGTTTGGT
        maximum likelihood sequence       ATAGTTTGGT
        maximum likelihood edit distance  0

evaluating sequence 10 of 100
        observed sequence                 ATAGTTTGGT
        maximum likelihood sequence       ATAGTTTGGT
        maximum likelihood edit distance  0

evaluating sequence 11 of 100
        observed sequence                 ATAGTTTGGT
        maximum likelihood sequence       ATAGTTTGGT
        maximum likelihood edit distance  0

evaluating sequence 12 of 100
        observed sequence                 ACCAAACTAT
        maximum likelihood sequence       ACCAAACTAT
        maximum likelihood edit distance  0

evaluating sequence 13 of 100
        observed sequence                 ACCAAACTAT
        maximum likelihood sequence       ACCAAACTAT
        maximum likelihood edit distance  0

evaluating sequence 14 of 100
        observed sequence                 ATAGTTTGGT
        maximum likelihood sequence       ATAGTTTGGT
        maximum likelihood edit distance  0

evaluating sequence 15 of 100
        observed sequence                 ATAGTTTGGT
        maximum likelihood sequence       ATAGTTTGGT
        maximum likelihood edit distance  0

evaluating sequence 16 of 100
        observed sequence                 TAGTTTGGT
        maximum likelihood sequence       TAGTTTGGT
        maximum likelihood edit distance  0

evaluating sequence 17 of 100
        observed sequence                 ACCAAACTAT
        maximum likelihood sequence       ACCAAACTAT
        maximum likelihood edit distance  0

evaluating sequence 18 of 100
        observed sequence                 ACCAAACTAT
        maximum likelihood sequence       ACCAAACTAT
        maximum likelihood edit distance  0

evaluating sequence 19 of 100
        observed sequence                 ACAAACTAT
        maximum likelihood sequence       CCAAACTAT
        maximum likelihood edit distance  1
```

```
evaluating sequence 20 of 100
        observed sequence                 ACCAAACTAT
        maximum likelihood sequence       ACCAAACTAT
        maximum likelihood edit distance  0

evaluating sequence 21 of 100
        observed sequence                 ATAGTTTGGT
        maximum likelihood sequence       ATAGTTTGGT
        maximum likelihood edit distance  0

evaluating sequence 22 of 100
        observed sequence                 ACCAAACT
        maximum likelihood sequence       ACCAAACT
        maximum likelihood edit distance  0

evaluating sequence 23 of 100
        observed sequence                 ACCAAACTAT
        maximum likelihood sequence       ACCAAACTAT
        maximum likelihood edit distance  0

evaluating sequence 24 of 100
        observed sequence                 ACCAAACTAT
        maximum likelihood sequence       ACCAAACTAT
        maximum likelihood edit distance  0

evaluating sequence 25 of 100
        observed sequence                 ATAGTTTGGT
        maximum likelihood sequence       ATAGTTTGGT
        maximum likelihood edit distance  0

evaluating sequence 26 of 100
        observed sequence                 ATAGTTTGGT
        maximum likelihood sequence       ATAGTTTGGT
        maximum likelihood edit distance  0

evaluating sequence 27 of 100
        observed sequence                 ATAGTTTGGT
        maximum likelihood sequence       ATAGTTTGGT
        maximum likelihood edit distance  0

evaluating sequence 28 of 100
        observed sequence                 AAACTAT
        maximum likelihood sequence       AAACTAT
        maximum likelihood edit distance  0

evaluating sequence 29 of 100
        observed sequence                 ATAGTTTGGT
        maximum likelihood sequence       ATAGTTTGGT
        maximum likelihood edit distance  0

evaluating sequence 30 of 100
        observed sequence                 ATAGTTTGGT
        maximum likelihood sequence       ATAGTTTGGT
        maximum likelihood edit distance  0

evaluating sequence 31 of 100
        observed sequence                 TAGTTTGGT
        maximum likelihood sequence       TAGTTTGGT
        maximum likelihood edit distance  0

evaluating sequence 32 of 100
        observed sequence                 ATAGTTTGGT
        maximum likelihood sequence       ATAGTTTGGT
        maximum likelihood edit distance  0

evaluating sequence 33 of 100
        observed sequence                 ATAGTTTGGT
        maximum likelihood sequence       ATAGTTTGGT
        maximum likelihood edit distance  0

evaluating sequence 34 of 100
        observed sequence                 ACCAAACT
        maximum likelihood sequence       ACCAAACT
        maximum likelihood edit distance  0

evaluating sequence 35 of 100
        observed sequence                 ACCAAACTAT
        maximum likelihood sequence       ACCAAACTAT
        maximum likelihood edit distance  0

evaluating sequence 36 of 100
        observed sequence                 TAGTTTGGT
        maximum likelihood sequence       TAGTTTGGT
        maximum likelihood edit distance  0

evaluating sequence 37 of 100
        observed sequence                 TAGTTTGGT
        maximum likelihood sequence       TAGTTTGGT
        maximum likelihood edit distance  0

evaluating sequence 38 of 100
        observed sequence                 ACCAAACTAT
        maximum likelihood sequence       ACCAAACTAT
        maximum likelihood edit distance  0

evaluating sequence 39 of 100
        observed sequence                 ATAGTTTGGT
        maximum likelihood sequence       ATAGTTTGGT
```

```
        maximum likelihood edit distance  0

evaluating sequence 40 of 100
        observed sequence                 GATAGTTTGGT
        maximum likelihood sequence       GATAGTTTGGT
        maximum likelihood edit distance  0

evaluating sequence 41 of 100
        observed sequence                 ATAGTTTGGT
        maximum likelihood sequence       ATAGTTTGGT
        maximum likelihood edit distance  0

evaluating sequence 42 of 100
        observed sequence                 ACAAACTAT
        maximum likelihood sequence       CCAAACTAT
        maximum likelihood edit distance  1

evaluating sequence 43 of 100
        observed sequence                 ACCAAACTAT
        maximum likelihood sequence       ACCAAACTAT
        maximum likelihood edit distance  0

evaluating sequence 44 of 100
        observed sequence                 ACCAAACTAT
        maximum likelihood sequence       ACCAAACTAT
        maximum likelihood edit distance  0

evaluating sequence 45 of 100
        observed sequence                 ACCAAACTAT
        maximum likelihood sequence       ACCAAACTAT
        maximum likelihood edit distance  0

evaluating sequence 46 of 100
        observed sequence                 ATAGTTTGGT
        maximum likelihood sequence       ATAGTTTGGT
        maximum likelihood edit distance  0

evaluating sequence 47 of 100
        observed sequence                 CCAAACT
        maximum likelihood sequence       CCAAACT
        maximum likelihood edit distance  0

evaluating sequence 48 of 100
        observed sequence                 ATAGTTTGGT
        maximum likelihood sequence       ATAGTTTGGT
        maximum likelihood edit distance  0

evaluating sequence 49 of 100
        observed sequence                 ATAGTTTGGT
        maximum likelihood sequence       ATAGTTTGGT
        maximum likelihood edit distance  0

evaluating sequence 50 of 100
        observed sequence                 ATAGTTTGGT
        maximum likelihood sequence       ATAGTTTGGT
        maximum likelihood edit distance  0

evaluating sequence 51 of 100
        observed sequence                 ACCAAACTAT
        maximum likelihood sequence       ACCAAACTAT
        maximum likelihood edit distance  0

evaluating sequence 52 of 100
        observed sequence                 ACAAACTAT
        maximum likelihood sequence       CCAAACTAT
        maximum likelihood edit distance  1

evaluating sequence 53 of 100
        observed sequence                 ACCAAACTAT
        maximum likelihood sequence       ACCAAACTAT
        maximum likelihood edit distance  0

evaluating sequence 54 of 100
        observed sequence                 AGTTTGGT
        maximum likelihood sequence       AGTTTGGT
        maximum likelihood edit distance  0

evaluating sequence 55 of 100
        observed sequence                 ATAGTTTGGT
        maximum likelihood sequence       ATAGTTTGGT
        maximum likelihood edit distance  0

evaluating sequence 56 of 100
        observed sequence                 ATAGTTTGGT
        maximum likelihood sequence       ATAGTTTGGT
        maximum likelihood edit distance  0

evaluating sequence 57 of 100
        observed sequence                 ACCAAACTAT
        maximum likelihood sequence       ACCAAACTAT
        maximum likelihood edit distance  0

evaluating sequence 58 of 100
        observed sequence                 ATAGTTTGGT
        maximum likelihood sequence       ATAGTTTGGT
        maximum likelihood edit distance  0

evaluating sequence 59 of 100
        observed sequence                 ACCAAACTAT
```

```
        maximum likelihood sequence       ACCAAACTAT
        maximum likelihood edit distance  0

evaluating sequence 60 of 100
        observed sequence                 ACCAAACTAT
        maximum likelihood sequence       ACCAAACTAT
        maximum likelihood edit distance  0

evaluating sequence 61 of 100
        observed sequence                 ATAGTTTGGT
        maximum likelihood sequence       ATAGTTTGGT
        maximum likelihood edit distance  0

evaluating sequence 62 of 100
        observed sequence                 TAGTTTGGT
        maximum likelihood sequence       TAGTTTGGT
        maximum likelihood edit distance  0

evaluating sequence 63 of 100
        observed sequence                 ATAGTTTGGT
        maximum likelihood sequence       ATAGTTTGGT
        maximum likelihood edit distance  0

evaluating sequence 64 of 100
        observed sequence                 ACCAAACTAT
        maximum likelihood sequence       ACCAAACTAT
        maximum likelihood edit distance  0

evaluating sequence 65 of 100
        observed sequence                 ATAGTTTGGT
        maximum likelihood sequence       ATAGTTTGGT
        maximum likelihood edit distance  0

evaluating sequence 66 of 100
        observed sequence                 ATAGTTTGGT
        maximum likelihood sequence       ATAGTTTGGT
        maximum likelihood edit distance  0

evaluating sequence 67 of 100
        observed sequence                 ACCAAACTAT
        maximum likelihood sequence       ACCAAACTAT
        maximum likelihood edit distance  0

evaluating sequence 68 of 100
        observed sequence                 GATAGTTTGGT
        maximum likelihood sequence       GATAGTTTGGT
        maximum likelihood edit distance  0

evaluating sequence 69 of 100
        observed sequence                 ATAGTTTGGT
        maximum likelihood sequence       ATAGTTTGGT
        maximum likelihood edit distance  0

evaluating sequence 70 of 100
        observed sequence                 CCAAACTAT
        maximum likelihood sequence       CCAAACTAT
        maximum likelihood edit distance  0

evaluating sequence 71 of 100
        observed sequence                 ACCAAACTAT
        maximum likelihood sequence       ACCAAACTAT
        maximum likelihood edit distance  0

evaluating sequence 72 of 100
        observed sequence                 ATAGTTTGGT
        maximum likelihood sequence       ATAGTTTGGT
        maximum likelihood edit distance  0

evaluating sequence 73 of 100
        observed sequence                 ACCAAACTAT
        maximum likelihood sequence       ACCAAACTAT
        maximum likelihood edit distance  0

evaluating sequence 74 of 100
        observed sequence                 ATAGTTTGGT
        maximum likelihood sequence       ATAGTTTGGT
        maximum likelihood edit distance  0

evaluating sequence 75 of 100
        observed sequence                 ACCAAACTAT
        maximum likelihood sequence       ACCAAACTAT
        maximum likelihood edit distance  0

evaluating sequence 76 of 100
        observed sequence                 ACCAAACTA
        maximum likelihood sequence       ACCAAACTA
        maximum likelihood edit distance  0

evaluating sequence 77 of 100
        observed sequence                 ATAGTTTGGT
        maximum likelihood sequence       ATAGTTTGGT
        maximum likelihood edit distance  0

evaluating sequence 78 of 100
        observed sequence                 ACCAAACTAT
        maximum likelihood sequence       ACCAAACTAT
        maximum likelihood edit distance  0

evaluating sequence 79 of 100
```

```
        observed sequence                 ATAGTTTGGT
        maximum likelihood sequence       ATAGTTTGGT
        maximum likelihood edit distance  0

evaluating sequence 80 of 100
        observed sequence                 ATAGTTTGGT
        maximum likelihood sequence       ATAGTTTGGT
        maximum likelihood edit distance  0

evaluating sequence 81 of 100
        observed sequence                 TAGTTTGGT
        maximum likelihood sequence       TAGTTTGGT
        maximum likelihood edit distance  0

evaluating sequence 82 of 100
        observed sequence                 ACCAAACTAT
        maximum likelihood sequence       ACCAAACTAT
        maximum likelihood edit distance  0

evaluating sequence 83 of 100
        observed sequence                 ATAGTTTGGT
        maximum likelihood sequence       ATAGTTTGGT
        maximum likelihood edit distance  0

evaluating sequence 84 of 100
        observed sequence                 ACCAAACT
        maximum likelihood sequence       ACCAAACT
        maximum likelihood edit distance  0

evaluating sequence 85 of 100
        observed sequence                 ATAGTTTGGT
        maximum likelihood sequence       ATAGTTTGGT
        maximum likelihood edit distance  0

evaluating sequence 86 of 100
        observed sequence                 ATAGTTTGGT
        maximum likelihood sequence       ATAGTTTGGT
        maximum likelihood edit distance  0

evaluating sequence 87 of 100
        observed sequence                 ACCAAACTAT
        maximum likelihood sequence       ACCAAACTAT
        maximum likelihood edit distance  0

evaluating sequence 88 of 100
        observed sequence                 ACCAAACTAT
        maximum likelihood sequence       ACCAAACTAT
        maximum likelihood edit distance  0

evaluating sequence 89 of 100
        observed sequence                 ACCAAACTAT
        maximum likelihood sequence       ACCAAACTAT
        maximum likelihood edit distance  0

evaluating sequence 90 of 100
        observed sequence                 ATAGTTTGGT
        maximum likelihood sequence       ATAGTTTGGT
        maximum likelihood edit distance  0

evaluating sequence 91 of 100
        observed sequence                 ACCAAACTAT
        maximum likelihood sequence       ACCAAACTAT
        maximum likelihood edit distance  0

evaluating sequence 92 of 100
        observed sequence                 ACCAAACTAT
        maximum likelihood sequence       ACCAAACTAT
        maximum likelihood edit distance  0

evaluating sequence 93 of 100
        observed sequence                 ACCAAACTAT
        maximum likelihood sequence       ACCAAACTAT
        maximum likelihood edit distance  0

evaluating sequence 94 of 100
        observed sequence                 ACCAAACTAT
        maximum likelihood sequence       ACCAAACTAT
        maximum likelihood edit distance  0

evaluating sequence 95 of 100
        observed sequence                 ACCAAACTAT
        maximum likelihood sequence       ACCAAACTAT
        maximum likelihood edit distance  0

evaluating sequence 96 of 100
        observed sequence                 ACCAAACTAT
        maximum likelihood sequence       ACCAAACTAT
        maximum likelihood edit distance  0

evaluating sequence 97 of 100
        observed sequence                 ATAGTTTGGT
        maximum likelihood sequence       ATAGTTTGGT
        maximum likelihood edit distance  0

evaluating sequence 98 of 100
        observed sequence                 ATAGTTTGGT
        maximum likelihood sequence       ATAGTTTGGT
        maximum likelihood edit distance  0
```

```
evaluating sequence 99 of 100
        observed sequence                 ATAGTTTGGT
        maximum likelihood sequence       ATAGTTTGGT
        maximum likelihood edit distance  0

evaluating sequence 100 of 100
        observed sequence                 ATAGTTTGGT
        maximum likelihood sequence       ATAGTTTGGT
        maximum likelihood edit distance  0

DATASET STATISTICS:
        assumed error rate    15.0%
        total bases observed  975
        total edits accepted  3
        inferred error rate   0.3076923076923077%
```

In [9]:

```
canonical_kmers = collect(keys(Eisenia.count_canonical_kmers(maximum_likelihood_observations, k)))
stranded_kmer_graph = Eisenia.build_stranded_kmer_graph(canonical_kmers, maximum_likelihood_observations)
filename = reference_sequence_id * "." * replace(string(Dates.now()), ':' => '.') * ".svg"
Eisenia.plot_stranded_kmer_graph(stranded_kmer_graph, filename=filename)
HTML("""
<image src="$filename" width=50%>
""")
```

Out[9]:

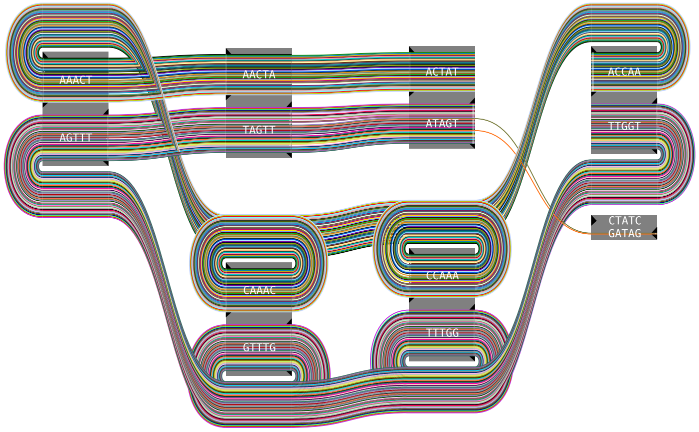


In [10]:

```
maximum_likelihood_observations = Eisenia.viterbi_maximum_likelihood_traversals(stranded_kmer_graph, error_rate = error_rate, verbosity="reads");
```

```
computing kmer counts...
computing kmer state likelihoods...
finding shortest paths between kmers...
finding viterbi maximum likelihood paths for observed sequences...

evaluating sequence 1 of 100
        observed sequence                 ACCAAACTA
        maximum likelihood sequence       ACCAAACTA
        maximum likelihood edit distance  0

evaluating sequence 2 of 100
        observed sequence                 ATAGTTTGGT
        maximum likelihood sequence       ATAGTTTGGT
        maximum likelihood edit distance  0

evaluating sequence 3 of 100
        observed sequence                 ATAGTTTGG
        maximum likelihood sequence       ATAGTTTGG
        maximum likelihood edit distance  0

evaluating sequence 4 of 100
        observed sequence                 ACCAAACTAT
        maximum likelihood sequence       ACCAAACTAT
        maximum likelihood edit distance  0

evaluating sequence 5 of 100
        observed sequence                 ACCAAACTAT
        maximum likelihood sequence       ACCAAACTAT
        maximum likelihood edit distance  0

evaluating sequence 6 of 100
        observed sequence                 ACCAAACTAT
        maximum likelihood sequence       ACCAAACTAT
        maximum likelihood edit distance  0

evaluating sequence 7 of 100
        observed sequence                 ACCAAACTAT
        maximum likelihood sequence       ACCAAACTAT
        maximum likelihood edit distance  0

evaluating sequence 8 of 100
        observed sequence                 ACCAAACTAT
        maximum likelihood sequence       ACCAAACTAT
        maximum likelihood edit distance  0

evaluating sequence 9 of 100
        observed sequence                 ATAGTTTGGT
        maximum likelihood sequence       ATAGTTTGGT
        maximum likelihood edit distance  0

evaluating sequence 10 of 100
        observed sequence                 ATAGTTTGGT
        maximum likelihood sequence       ATAGTTTGGT
        maximum likelihood edit distance  0

evaluating sequence 11 of 100
        observed sequence                 ATAGTTTGGT
        maximum likelihood sequence       ATAGTTTGGT
        maximum likelihood edit distance  0

evaluating sequence 12 of 100
        observed sequence                 ACCAAACTAT
        maximum likelihood sequence       ACCAAACTAT
        maximum likelihood edit distance  0

evaluating sequence 13 of 100
        observed sequence                 ACCAAACTAT
        maximum likelihood sequence       ACCAAACTAT
        maximum likelihood edit distance  0

evaluating sequence 14 of 100
        observed sequence                 ATAGTTTGGT
        maximum likelihood sequence       ATAGTTTGGT
        maximum likelihood edit distance  0

evaluating sequence 15 of 100
        observed sequence                 ATAGTTTGGT
        maximum likelihood sequence       ATAGTTTGGT
        maximum likelihood edit distance  0

evaluating sequence 16 of 100
        observed sequence                 TAGTTTGGT
        maximum likelihood sequence       TAGTTTGGT
        maximum likelihood edit distance  0

evaluating sequence 17 of 100
        observed sequence                 ACCAAACTAT
        maximum likelihood sequence       ACCAAACTAT
        maximum likelihood edit distance  0

evaluating sequence 18 of 100
        observed sequence                 ACCAAACTAT
        maximum likelihood sequence       ACCAAACTAT
        maximum likelihood edit distance  0

evaluating sequence 19 of 100
        observed sequence                 CCAAACTAT
        maximum likelihood sequence       CCAAACTAT
        maximum likelihood edit distance  0
```

```
evaluating sequence 20 of 100
        observed sequence                 ACCAAACTAT
        maximum likelihood sequence       ACCAAACTAT
        maximum likelihood edit distance  0

evaluating sequence 21 of 100
        observed sequence                 ATAGTTTGGT
        maximum likelihood sequence       ATAGTTTGGT
        maximum likelihood edit distance  0

evaluating sequence 22 of 100
        observed sequence                 ACCAAACT
        maximum likelihood sequence       ACCAAACT
        maximum likelihood edit distance  0

evaluating sequence 23 of 100
        observed sequence                 ACCAAACTAT
        maximum likelihood sequence       ACCAAACTAT
        maximum likelihood edit distance  0

evaluating sequence 24 of 100
        observed sequence                 ACCAAACTAT
        maximum likelihood sequence       ACCAAACTAT
        maximum likelihood edit distance  0

evaluating sequence 25 of 100
        observed sequence                 ATAGTTTGGT
        maximum likelihood sequence       ATAGTTTGGT
        maximum likelihood edit distance  0

evaluating sequence 26 of 100
        observed sequence                 ATAGTTTGGT
        maximum likelihood sequence       ATAGTTTGGT
        maximum likelihood edit distance  0

evaluating sequence 27 of 100
        observed sequence                 ATAGTTTGGT
        maximum likelihood sequence       ATAGTTTGGT
        maximum likelihood edit distance  0

evaluating sequence 28 of 100
        observed sequence                 AAACTAT
        maximum likelihood sequence       AAACTAT
        maximum likelihood edit distance  0

evaluating sequence 29 of 100
        observed sequence                 ATAGTTTGGT
        maximum likelihood sequence       ATAGTTTGGT
        maximum likelihood edit distance  0

evaluating sequence 30 of 100
        observed sequence                 ATAGTTTGGT
        maximum likelihood sequence       ATAGTTTGGT
        maximum likelihood edit distance  0

evaluating sequence 31 of 100
        observed sequence                 TAGTTTGGT
        maximum likelihood sequence       TAGTTTGGT
        maximum likelihood edit distance  0

evaluating sequence 32 of 100
        observed sequence                 ATAGTTTGGT
        maximum likelihood sequence       ATAGTTTGGT
        maximum likelihood edit distance  0

evaluating sequence 33 of 100
        observed sequence                 ATAGTTTGGT
        maximum likelihood sequence       ATAGTTTGGT
        maximum likelihood edit distance  0

evaluating sequence 34 of 100
        observed sequence                 ACCAAACT
        maximum likelihood sequence       ACCAAACT
        maximum likelihood edit distance  0

evaluating sequence 35 of 100
        observed sequence                 ACCAAACTAT
        maximum likelihood sequence       ACCAAACTAT
        maximum likelihood edit distance  0

evaluating sequence 36 of 100
        observed sequence                 TAGTTTGGT
        maximum likelihood sequence       TAGTTTGGT
        maximum likelihood edit distance  0

evaluating sequence 37 of 100
        observed sequence                 TAGTTTGGT
        maximum likelihood sequence       TAGTTTGGT
        maximum likelihood edit distance  0

evaluating sequence 38 of 100
        observed sequence                 ACCAAACTAT
        maximum likelihood sequence       ACCAAACTAT
        maximum likelihood edit distance  0

evaluating sequence 39 of 100
        observed sequence                 ATAGTTTGGT
        maximum likelihood sequence       ATAGTTTGGT
```

```
        maximum likelihood edit distance  0

evaluating sequence 40 of 100
        observed sequence                 GATAGTTTGGT
        maximum likelihood sequence       GATAGTTTGGT
        maximum likelihood edit distance  0

evaluating sequence 41 of 100
        observed sequence                 ATAGTTTGGT
        maximum likelihood sequence       ATAGTTTGGT
        maximum likelihood edit distance  0

evaluating sequence 42 of 100
        observed sequence                 CCAAACTAT
        maximum likelihood sequence       CCAAACTAT
        maximum likelihood edit distance  0

evaluating sequence 43 of 100
        observed sequence                 ACCAAACTAT
        maximum likelihood sequence       ACCAAACTAT
        maximum likelihood edit distance  0

evaluating sequence 44 of 100
        observed sequence                 ACCAAACTAT
        maximum likelihood sequence       ACCAAACTAT
        maximum likelihood edit distance  0

evaluating sequence 45 of 100
        observed sequence                 ACCAAACTAT
        maximum likelihood sequence       ACCAAACTAT
        maximum likelihood edit distance  0

evaluating sequence 46 of 100
        observed sequence                 ATAGTTTGGT
        maximum likelihood sequence       ATAGTTTGGT
        maximum likelihood edit distance  0

evaluating sequence 47 of 100
        observed sequence                 CCAAACT
        maximum likelihood sequence       CCAAACT
        maximum likelihood edit distance  0

evaluating sequence 48 of 100
        observed sequence                 ATAGTTTGGT
        maximum likelihood sequence       ATAGTTTGGT
        maximum likelihood edit distance  0

evaluating sequence 49 of 100
        observed sequence                 ATAGTTTGGT
        maximum likelihood sequence       ATAGTTTGGT
        maximum likelihood edit distance  0

evaluating sequence 50 of 100
        observed sequence                 ATAGTTTGGT
        maximum likelihood sequence       ATAGTTTGGT
        maximum likelihood edit distance  0

evaluating sequence 51 of 100
        observed sequence                 ACCAAACTAT
        maximum likelihood sequence       ACCAAACTAT
        maximum likelihood edit distance  0

evaluating sequence 52 of 100
        observed sequence                 CCAAACTAT
        maximum likelihood sequence       CCAAACTAT
        maximum likelihood edit distance  0

evaluating sequence 53 of 100
        observed sequence                 ACCAAACTAT
        maximum likelihood sequence       ACCAAACTAT
        maximum likelihood edit distance  0

evaluating sequence 54 of 100
        observed sequence                 AGTTTGGT
        maximum likelihood sequence       AGTTTGGT
        maximum likelihood edit distance  0

evaluating sequence 55 of 100
        observed sequence                 ATAGTTTGGT
        maximum likelihood sequence       ATAGTTTGGT
        maximum likelihood edit distance  0

evaluating sequence 56 of 100
        observed sequence                 ATAGTTTGGT
        maximum likelihood sequence       ATAGTTTGGT
        maximum likelihood edit distance  0

evaluating sequence 57 of 100
        observed sequence                 ACCAAACTAT
        maximum likelihood sequence       ACCAAACTAT
        maximum likelihood edit distance  0

evaluating sequence 58 of 100
        observed sequence                 ATAGTTTGGT
        maximum likelihood sequence       ATAGTTTGGT
        maximum likelihood edit distance  0

evaluating sequence 59 of 100
        observed sequence                 ACCAAACTAT
```

```
        maximum likelihood sequence       ACCAAACTAT
        maximum likelihood edit distance  0

evaluating sequence 60 of 100
        observed sequence                 ACCAAACTAT
        maximum likelihood sequence       ACCAAACTAT
        maximum likelihood edit distance  0

evaluating sequence 61 of 100
        observed sequence                 ATAGTTTGGT
        maximum likelihood sequence       ATAGTTTGGT
        maximum likelihood edit distance  0

evaluating sequence 62 of 100
        observed sequence                 TAGTTTGGT
        maximum likelihood sequence       TAGTTTGGT
        maximum likelihood edit distance  0

evaluating sequence 63 of 100
        observed sequence                 ATAGTTTGGT
        maximum likelihood sequence       ATAGTTTGGT
        maximum likelihood edit distance  0

evaluating sequence 64 of 100
        observed sequence                 ACCAAACTAT
        maximum likelihood sequence       ACCAAACTAT
        maximum likelihood edit distance  0

evaluating sequence 65 of 100
        observed sequence                 ATAGTTTGGT
        maximum likelihood sequence       ATAGTTTGGT
        maximum likelihood edit distance  0

evaluating sequence 66 of 100
        observed sequence                 ATAGTTTGGT
        maximum likelihood sequence       ATAGTTTGGT
        maximum likelihood edit distance  0

evaluating sequence 67 of 100
        observed sequence                 ACCAAACTAT
        maximum likelihood sequence       ACCAAACTAT
        maximum likelihood edit distance  0

evaluating sequence 68 of 100
        observed sequence                 GATAGTTTGGT
        maximum likelihood sequence       GATAGTTTGGT
        maximum likelihood edit distance  0

evaluating sequence 69 of 100
        observed sequence                 ATAGTTTGGT
        maximum likelihood sequence       ATAGTTTGGT
        maximum likelihood edit distance  0

evaluating sequence 70 of 100
        observed sequence                 CCAAACTAT
        maximum likelihood sequence       CCAAACTAT
        maximum likelihood edit distance  0

evaluating sequence 71 of 100
        observed sequence                 ACCAAACTAT
        maximum likelihood sequence       ACCAAACTAT
        maximum likelihood edit distance  0

evaluating sequence 72 of 100
        observed sequence                 ATAGTTTGGT
        maximum likelihood sequence       ATAGTTTGGT
        maximum likelihood edit distance  0

evaluating sequence 73 of 100
        observed sequence                 ACCAAACTAT
        maximum likelihood sequence       ACCAAACTAT
        maximum likelihood edit distance  0

evaluating sequence 74 of 100
        observed sequence                 ATAGTTTGGT
        maximum likelihood sequence       ATAGTTTGGT
        maximum likelihood edit distance  0

evaluating sequence 75 of 100
        observed sequence                 ACCAAACTAT
        maximum likelihood sequence       ACCAAACTAT
        maximum likelihood edit distance  0

evaluating sequence 76 of 100
        observed sequence                 ACCAAACTA
        maximum likelihood sequence       ACCAAACTA
        maximum likelihood edit distance  0

evaluating sequence 77 of 100
        observed sequence                 ATAGTTTGGT
        maximum likelihood sequence       ATAGTTTGGT
        maximum likelihood edit distance  0

evaluating sequence 78 of 100
        observed sequence                 ACCAAACTAT
        maximum likelihood sequence       ACCAAACTAT
        maximum likelihood edit distance  0

evaluating sequence 79 of 100
```

```
        observed sequence                 ATAGTTTGGT
        maximum likelihood sequence       ATAGTTTGGT
        maximum likelihood edit distance  0

evaluating sequence 80 of 100
        observed sequence                 ATAGTTTGGT
        maximum likelihood sequence       ATAGTTTGGT
        maximum likelihood edit distance  0

evaluating sequence 81 of 100
        observed sequence                 TAGTTTGGT
        maximum likelihood sequence       TAGTTTGGT
        maximum likelihood edit distance  0

evaluating sequence 82 of 100
        observed sequence                 ACCAAACTAT
        maximum likelihood sequence       ACCAAACTAT
        maximum likelihood edit distance  0

evaluating sequence 83 of 100
        observed sequence                 ATAGTTTGGT
        maximum likelihood sequence       ATAGTTTGGT
        maximum likelihood edit distance  0

evaluating sequence 84 of 100
        observed sequence                 ACCAAACT
        maximum likelihood sequence       ACCAAACT
        maximum likelihood edit distance  0

evaluating sequence 85 of 100
        observed sequence                 ATAGTTTGGT
        maximum likelihood sequence       ATAGTTTGGT
        maximum likelihood edit distance  0

evaluating sequence 86 of 100
        observed sequence                 ATAGTTTGGT
        maximum likelihood sequence       ATAGTTTGGT
        maximum likelihood edit distance  0

evaluating sequence 87 of 100
        observed sequence                 ACCAAACTAT
        maximum likelihood sequence       ACCAAACTAT
        maximum likelihood edit distance  0

evaluating sequence 88 of 100
        observed sequence                 ACCAAACTAT
        maximum likelihood sequence       ACCAAACTAT
        maximum likelihood edit distance  0

evaluating sequence 89 of 100
        observed sequence                 ACCAAACTAT
        maximum likelihood sequence       ACCAAACTAT
        maximum likelihood edit distance  0

evaluating sequence 90 of 100
        observed sequence                 ATAGTTTGGT
        maximum likelihood sequence       ATAGTTTGGT
        maximum likelihood edit distance  0

evaluating sequence 91 of 100
        observed sequence                 ACCAAACTAT
        maximum likelihood sequence       ACCAAACTAT
        maximum likelihood edit distance  0

evaluating sequence 92 of 100
        observed sequence                 ACCAAACTAT
        maximum likelihood sequence       ACCAAACTAT
        maximum likelihood edit distance  0

evaluating sequence 93 of 100
        observed sequence                 ACCAAACTAT
        maximum likelihood sequence       ACCAAACTAT
        maximum likelihood edit distance  0

evaluating sequence 94 of 100
        observed sequence                 ACCAAACTAT
        maximum likelihood sequence       ACCAAACTAT
        maximum likelihood edit distance  0

evaluating sequence 95 of 100
        observed sequence                 ACCAAACTAT
        maximum likelihood sequence       ACCAAACTAT
        maximum likelihood edit distance  0

evaluating sequence 96 of 100
        observed sequence                 ACCAAACTAT
        maximum likelihood sequence       ACCAAACTAT
        maximum likelihood edit distance  0

evaluating sequence 97 of 100
        observed sequence                 ATAGTTTGGT
        maximum likelihood sequence       ATAGTTTGGT
        maximum likelihood edit distance  0

evaluating sequence 98 of 100
        observed sequence                 ATAGTTTGGT
        maximum likelihood sequence       ATAGTTTGGT
        maximum likelihood edit distance  0
```

```
evaluating sequence 99 of 100
        observed sequence                  ATAGTTTGGT
        maximum likelihood sequence        ATAGTTTGGT
        maximum likelihood edit distance  0

evaluating sequence 100 of 100
        observed sequence                  ATAGTTTGGT
        maximum likelihood sequence        ATAGTTTGGT
        maximum likelihood edit distance  0

DATASET STATISTICS:
        assumed error rate    15.0%
        total bases observed  975
        total edits accepted  0
        inferred error rate   0.0%
```

In [11]:

```
k = 7
canonical_kmers = collect(keys(Eisenia.count_canonical_kmers(maximum_likelihood_observations, k)))
stranded_kmer_graph = Eisenia.build_stranded_kmer_graph(canonical_kmers, maximum_likelihood_observations)
filename = reference_sequence_id * "." * replace(string(Dates.now()), ':' => '.') * ".svg"
Eisenia.plot_stranded_kmer_graph(stranded_kmer_graph, filename=filename)
HTML("""
<image src="$filename" width=50%>
""")
```

Out[11]:

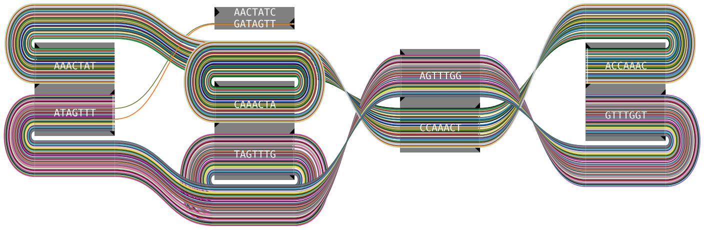


In [12]:

```
maximum_likelihood_observations = Eisenia.viterbi_maximum_likelihood_traversals(stranded_kmer_graph, error_rate = error_rate, verbosity="reads");
```

```
computing kmer counts...
computing kmer state likelihoods...
finding shortest paths between kmers...
finding viterbi maximum likelihood paths for observed sequences...

evaluating sequence 1 of 100
        observed sequence                 ACCAAACTA
        maximum likelihood sequence       ACCAAACTA
        maximum likelihood edit distance  0

evaluating sequence 2 of 100
        observed sequence                 ATAGTTTGGT
        maximum likelihood sequence       ATAGTTTGGT
        maximum likelihood edit distance  0

evaluating sequence 3 of 100
        observed sequence                 ATAGTTTGG
        maximum likelihood sequence       ATAGTTTGG
        maximum likelihood edit distance  0

evaluating sequence 4 of 100
        observed sequence                 ACCAAACTAT
        maximum likelihood sequence       ACCAAACTAT
        maximum likelihood edit distance  0

evaluating sequence 5 of 100
        observed sequence                 ACCAAACTAT
        maximum likelihood sequence       ACCAAACTAT
        maximum likelihood edit distance  0

evaluating sequence 6 of 100
        observed sequence                 ACCAAACTAT
        maximum likelihood sequence       ACCAAACTAT
        maximum likelihood edit distance  0

evaluating sequence 7 of 100
        observed sequence                 ACCAAACTAT
        maximum likelihood sequence       ACCAAACTAT
        maximum likelihood edit distance  0

evaluating sequence 8 of 100
        observed sequence                 ACCAAACTAT
        maximum likelihood sequence       ACCAAACTAT
        maximum likelihood edit distance  0

evaluating sequence 9 of 100
        observed sequence                 ATAGTTTGGT
        maximum likelihood sequence       ATAGTTTGGT
        maximum likelihood edit distance  0

evaluating sequence 10 of 100
        observed sequence                 ATAGTTTGGT
        maximum likelihood sequence       ATAGTTTGGT
        maximum likelihood edit distance  0

evaluating sequence 11 of 100
        observed sequence                 ATAGTTTGGT
        maximum likelihood sequence       ATAGTTTGGT
        maximum likelihood edit distance  0

evaluating sequence 12 of 100
        observed sequence                 ACCAAACTAT
        maximum likelihood sequence       ACCAAACTAT
        maximum likelihood edit distance  0

evaluating sequence 13 of 100
        observed sequence                 ACCAAACTAT
        maximum likelihood sequence       ACCAAACTAT
        maximum likelihood edit distance  0

evaluating sequence 14 of 100
        observed sequence                 ATAGTTTGGT
        maximum likelihood sequence       ATAGTTTGGT
        maximum likelihood edit distance  0

evaluating sequence 15 of 100
        observed sequence                 ATAGTTTGGT
        maximum likelihood sequence       ATAGTTTGGT
        maximum likelihood edit distance  0

evaluating sequence 16 of 100
        observed sequence                 TAGTTTGGT
        maximum likelihood sequence       TAGTTTGGT
        maximum likelihood edit distance  0

evaluating sequence 17 of 100
        observed sequence                 ACCAAACTAT
        maximum likelihood sequence       ACCAAACTAT
        maximum likelihood edit distance  0

evaluating sequence 18 of 100
        observed sequence                 ACCAAACTAT
        maximum likelihood sequence       ACCAAACTAT
        maximum likelihood edit distance  0

evaluating sequence 19 of 100
        observed sequence                 CCAAACTAT
        maximum likelihood sequence       CCAAACTAT
        maximum likelihood edit distance  0
```

```
evaluating sequence 20 of 100
        observed sequence                 ACCAAACTAT
        maximum likelihood sequence       ACCAAACTAT
        maximum likelihood edit distance  0

evaluating sequence 21 of 100
        observed sequence                 ATAGTTTGGT
        maximum likelihood sequence       ATAGTTTGGT
        maximum likelihood edit distance  0

evaluating sequence 22 of 100
        observed sequence                 ACCAAACT
        maximum likelihood sequence       ACCAAACT
        maximum likelihood edit distance  0

evaluating sequence 23 of 100
        observed sequence                 ACCAAACTAT
        maximum likelihood sequence       ACCAAACTAT
        maximum likelihood edit distance  0

evaluating sequence 24 of 100
        observed sequence                 ACCAAACTAT
        maximum likelihood sequence       ACCAAACTAT
        maximum likelihood edit distance  0

evaluating sequence 25 of 100
        observed sequence                 ATAGTTTGGT
        maximum likelihood sequence       ATAGTTTGGT
        maximum likelihood edit distance  0

evaluating sequence 26 of 100
        observed sequence                 ATAGTTTGGT
        maximum likelihood sequence       ATAGTTTGGT
        maximum likelihood edit distance  0

evaluating sequence 27 of 100
        observed sequence                 ATAGTTTGGT
        maximum likelihood sequence       ATAGTTTGGT
        maximum likelihood edit distance  0

evaluating sequence 28 of 100
        observed sequence                 AAACTAT
        maximum likelihood sequence       AAACTAT
        maximum likelihood edit distance  0

evaluating sequence 29 of 100
        observed sequence                 ATAGTTTGGT
        maximum likelihood sequence       ATAGTTTGGT
        maximum likelihood edit distance  0

evaluating sequence 30 of 100
        observed sequence                 ATAGTTTGGT
        maximum likelihood sequence       ATAGTTTGGT
        maximum likelihood edit distance  0

evaluating sequence 31 of 100
        observed sequence                 TAGTTTGGT
        maximum likelihood sequence       TAGTTTGGT
        maximum likelihood edit distance  0

evaluating sequence 32 of 100
        observed sequence                 ATAGTTTGGT
        maximum likelihood sequence       ATAGTTTGGT
        maximum likelihood edit distance  0

evaluating sequence 33 of 100
        observed sequence                 ATAGTTTGGT
        maximum likelihood sequence       ATAGTTTGGT
        maximum likelihood edit distance  0

evaluating sequence 34 of 100
        observed sequence                 ACCAAACT
        maximum likelihood sequence       ACCAAACT
        maximum likelihood edit distance  0

evaluating sequence 35 of 100
        observed sequence                 ACCAAACTAT
        maximum likelihood sequence       ACCAAACTAT
        maximum likelihood edit distance  0

evaluating sequence 36 of 100
        observed sequence                 TAGTTTGGT
        maximum likelihood sequence       TAGTTTGGT
        maximum likelihood edit distance  0

evaluating sequence 37 of 100
        observed sequence                 TAGTTTGGT
        maximum likelihood sequence       TAGTTTGGT
        maximum likelihood edit distance  0

evaluating sequence 38 of 100
        observed sequence                 ACCAAACTAT
        maximum likelihood sequence       ACCAAACTAT
        maximum likelihood edit distance  0

evaluating sequence 39 of 100
        observed sequence                 ATAGTTTGGT
        maximum likelihood sequence       ATAGTTTGGT
```

```
        maximum likelihood edit distance  0

evaluating sequence 40 of 100
        observed sequence                 GATAGTTTGGT
        maximum likelihood sequence       GATAGTTTGGT
        maximum likelihood edit distance  0

evaluating sequence 41 of 100
        observed sequence                 ATAGTTTGGT
        maximum likelihood sequence       ATAGTTTGGT
        maximum likelihood edit distance  0

evaluating sequence 42 of 100
        observed sequence                 CCAAACTAT
        maximum likelihood sequence       CCAAACTAT
        maximum likelihood edit distance  0

evaluating sequence 43 of 100
        observed sequence                 ACCAAACTAT
        maximum likelihood sequence       ACCAAACTAT
        maximum likelihood edit distance  0

evaluating sequence 44 of 100
        observed sequence                 ACCAAACTAT
        maximum likelihood sequence       ACCAAACTAT
        maximum likelihood edit distance  0

evaluating sequence 45 of 100
        observed sequence                 ACCAAACTAT
        maximum likelihood sequence       ACCAAACTAT
        maximum likelihood edit distance  0

evaluating sequence 46 of 100
        observed sequence                 ATAGTTTGGT
        maximum likelihood sequence       ATAGTTTGGT
        maximum likelihood edit distance  0

evaluating sequence 47 of 100
        observed sequence                 CCAAACT
        maximum likelihood sequence       CCAAACT
        maximum likelihood edit distance  0

evaluating sequence 48 of 100
        observed sequence                 ATAGTTTGGT
        maximum likelihood sequence       ATAGTTTGGT
        maximum likelihood edit distance  0

evaluating sequence 49 of 100
        observed sequence                 ATAGTTTGGT
        maximum likelihood sequence       ATAGTTTGGT
        maximum likelihood edit distance  0

evaluating sequence 50 of 100
        observed sequence                 ATAGTTTGGT
        maximum likelihood sequence       ATAGTTTGGT
        maximum likelihood edit distance  0

evaluating sequence 51 of 100
        observed sequence                 ACCAAACTAT
        maximum likelihood sequence       ACCAAACTAT
        maximum likelihood edit distance  0

evaluating sequence 52 of 100
        observed sequence                 CCAAACTAT
        maximum likelihood sequence       CCAAACTAT
        maximum likelihood edit distance  0

evaluating sequence 53 of 100
        observed sequence                 ACCAAACTAT
        maximum likelihood sequence       ACCAAACTAT
        maximum likelihood edit distance  0

evaluating sequence 54 of 100
        observed sequence                 AGTTTGGT
        maximum likelihood sequence       AGTTTGGT
        maximum likelihood edit distance  0

evaluating sequence 55 of 100
        observed sequence                 ATAGTTTGGT
        maximum likelihood sequence       ATAGTTTGGT
        maximum likelihood edit distance  0

evaluating sequence 56 of 100
        observed sequence                 ATAGTTTGGT
        maximum likelihood sequence       ATAGTTTGGT
        maximum likelihood edit distance  0

evaluating sequence 57 of 100
        observed sequence                 ACCAAACTAT
        maximum likelihood sequence       ACCAAACTAT
        maximum likelihood edit distance  0

evaluating sequence 58 of 100
        observed sequence                 ATAGTTTGGT
        maximum likelihood sequence       ATAGTTTGGT
        maximum likelihood edit distance  0

evaluating sequence 59 of 100
        observed sequence                 ACCAAACTAT
```

```
        maximum likelihood sequence       ACCAAACTAT
        maximum likelihood edit distance  0

evaluating sequence 60 of 100
        observed sequence                 ACCAAACTAT
        maximum likelihood sequence       ACCAAACTAT
        maximum likelihood edit distance  0

evaluating sequence 61 of 100
        observed sequence                 ATAGTTTGGT
        maximum likelihood sequence       ATAGTTTGGT
        maximum likelihood edit distance  0

evaluating sequence 62 of 100
        observed sequence                 TAGTTTGGT
        maximum likelihood sequence       TAGTTTGGT
        maximum likelihood edit distance  0

evaluating sequence 63 of 100
        observed sequence                 ATAGTTTGGT
        maximum likelihood sequence       ATAGTTTGGT
        maximum likelihood edit distance  0

evaluating sequence 64 of 100
        observed sequence                 ACCAAACTAT
        maximum likelihood sequence       ACCAAACTAT
        maximum likelihood edit distance  0

evaluating sequence 65 of 100
        observed sequence                 ATAGTTTGGT
        maximum likelihood sequence       ATAGTTTGGT
        maximum likelihood edit distance  0

evaluating sequence 66 of 100
        observed sequence                 ATAGTTTGGT
        maximum likelihood sequence       ATAGTTTGGT
        maximum likelihood edit distance  0

evaluating sequence 67 of 100
        observed sequence                 ACCAAACTAT
        maximum likelihood sequence       ACCAAACTAT
        maximum likelihood edit distance  0

evaluating sequence 68 of 100
        observed sequence                 GATAGTTTGGT
        maximum likelihood sequence       GATAGTTTGGT
        maximum likelihood edit distance  0

evaluating sequence 69 of 100
        observed sequence                 ATAGTTTGGT
        maximum likelihood sequence       ATAGTTTGGT
        maximum likelihood edit distance  0

evaluating sequence 70 of 100
        observed sequence                 CCAAACTAT
        maximum likelihood sequence       CCAAACTAT
        maximum likelihood edit distance  0

evaluating sequence 71 of 100
        observed sequence                 ACCAAACTAT
        maximum likelihood sequence       ACCAAACTAT
        maximum likelihood edit distance  0

evaluating sequence 72 of 100
        observed sequence                 ATAGTTTGGT
        maximum likelihood sequence       ATAGTTTGGT
        maximum likelihood edit distance  0

evaluating sequence 73 of 100
        observed sequence                 ACCAAACTAT
        maximum likelihood sequence       ACCAAACTAT
        maximum likelihood edit distance  0

evaluating sequence 74 of 100
        observed sequence                 ATAGTTTGGT
        maximum likelihood sequence       ATAGTTTGGT
        maximum likelihood edit distance  0

evaluating sequence 75 of 100
        observed sequence                 ACCAAACTAT
        maximum likelihood sequence       ACCAAACTAT
        maximum likelihood edit distance  0

evaluating sequence 76 of 100
        observed sequence                 ACCAAACTA
        maximum likelihood sequence       ACCAAACTA
        maximum likelihood edit distance  0

evaluating sequence 77 of 100
        observed sequence                 ATAGTTTGGT
        maximum likelihood sequence       ATAGTTTGGT
        maximum likelihood edit distance  0

evaluating sequence 78 of 100
        observed sequence                 ACCAAACTAT
        maximum likelihood sequence       ACCAAACTAT
        maximum likelihood edit distance  0

evaluating sequence 79 of 100
```

```
        observed sequence                 ATAGTTTGGT
        maximum likelihood sequence       ATAGTTTGGT
        maximum likelihood edit distance  0

evaluating sequence 80 of 100
        observed sequence                 ATAGTTTGGT
        maximum likelihood sequence       ATAGTTTGGT
        maximum likelihood edit distance  0

evaluating sequence 81 of 100
        observed sequence                 TAGTTTGGT
        maximum likelihood sequence       TAGTTTGGT
        maximum likelihood edit distance  0

evaluating sequence 82 of 100
        observed sequence                 ACCAAACTAT
        maximum likelihood sequence       ACCAAACTAT
        maximum likelihood edit distance  0

evaluating sequence 83 of 100
        observed sequence                 ATAGTTTGGT
        maximum likelihood sequence       ATAGTTTGGT
        maximum likelihood edit distance  0

evaluating sequence 84 of 100
        observed sequence                 ACCAAACT
        maximum likelihood sequence       ACCAAACT
        maximum likelihood edit distance  0

evaluating sequence 85 of 100
        observed sequence                 ATAGTTTGGT
        maximum likelihood sequence       ATAGTTTGGT
        maximum likelihood edit distance  0

evaluating sequence 86 of 100
        observed sequence                 ATAGTTTGGT
        maximum likelihood sequence       ATAGTTTGGT
        maximum likelihood edit distance  0

evaluating sequence 87 of 100
        observed sequence                 ACCAAACTAT
        maximum likelihood sequence       ACCAAACTAT
        maximum likelihood edit distance  0

evaluating sequence 88 of 100
        observed sequence                 ACCAAACTAT
        maximum likelihood sequence       ACCAAACTAT
        maximum likelihood edit distance  0

evaluating sequence 89 of 100
        observed sequence                 ACCAAACTAT
        maximum likelihood sequence       ACCAAACTAT
        maximum likelihood edit distance  0

evaluating sequence 90 of 100
        observed sequence                 ATAGTTTGGT
        maximum likelihood sequence       ATAGTTTGGT
        maximum likelihood edit distance  0

evaluating sequence 91 of 100
        observed sequence                 ACCAAACTAT
        maximum likelihood sequence       ACCAAACTAT
        maximum likelihood edit distance  0

evaluating sequence 92 of 100
        observed sequence                 ACCAAACTAT
        maximum likelihood sequence       ACCAAACTAT
        maximum likelihood edit distance  0

evaluating sequence 93 of 100
        observed sequence                 ACCAAACTAT
        maximum likelihood sequence       ACCAAACTAT
        maximum likelihood edit distance  0

evaluating sequence 94 of 100
        observed sequence                 ACCAAACTAT
        maximum likelihood sequence       ACCAAACTAT
        maximum likelihood edit distance  0

evaluating sequence 95 of 100
        observed sequence                 ACCAAACTAT
        maximum likelihood sequence       ACCAAACTAT
        maximum likelihood edit distance  0

evaluating sequence 96 of 100
        observed sequence                 ACCAAACTAT
        maximum likelihood sequence       ACCAAACTAT
        maximum likelihood edit distance  0

evaluating sequence 97 of 100
        observed sequence                 ATAGTTTGGT
        maximum likelihood sequence       ATAGTTTGGT
        maximum likelihood edit distance  0

evaluating sequence 98 of 100
        observed sequence                 ATAGTTTGGT
        maximum likelihood sequence       ATAGTTTGGT
        maximum likelihood edit distance  0
```

```
evaluating sequence 99 of 100
	observed sequence                 ATAGTTTGGT
	maximum likelihood sequence       ATAGTTTGGT
	maximum likelihood edit distance  0

evaluating sequence 100 of 100
	observed sequence                 ATAGTTTGGT
	maximum likelihood sequence       ATAGTTTGGT
	maximum likelihood edit distance  0

DATASET STATISTICS:
	assumed error rate    15.0%
	total bases observed  975
	total edits accepted  0
	inferred error rate   0.0%
```

## L20

In [13]:
```
L = 20
Random.seed!(L)
reference_sequence = randdnaseq(L)
reference_sequence_id = randstring(Int(round(log10(length(L)))+3))
reference_FASTA_record = FASTA.Record(reference_sequence_id, reference_sequence)
```

Out[13]:
```
BioSequences.FASTA.Record:
   identifier: Bg3
  description: <missing>
     sequence: CTGCAAGGTCGAATCCGGTC
```

In [14]:
```
error_rate = 0.15
n_sequences = 100
observations = [Eisenia.observe(reference_FASTA_record, error_rate=error_rate) for i in 1:n_sequences]
```

```
Out[14]: 100-element Array{BioSequences.FASTA.Record,1}:
 BioSequences.FASTA.Record:
   identifier: UClF
  description: <missing>
     sequence: GACCGGATTCGGCCTGGGAG
 BioSequences.FASTA.Record:
   identifier: 6u5g
  description: <missing>
     sequence: CTGCCCAGGGTCGGAATCCGGTAC
 BioSequences.FASTA.Record:
   identifier: CghJ
  description: <missing>
     sequence: GATCCGGATTCGATCCTTGCAG
 BioSequences.FASTA.Record:
   identifier: WVAh
  description: <missing>
     sequence: ACCGAGATATCGACCTTGCAG
 BioSequences.FASTA.Record:
   identifier: yBd8
  description: <missing>
     sequence: GACCGGATTCCGATGGCAG
 BioSequences.FASTA.Record:
   identifier: Bxw5
  description: <missing>
     sequence: GAACCGGGTTCGACCTTGCAG
 BioSequences.FASTA.Record:
   identifier: XFoi
  description: <missing>
     sequence: CTGCAGGTTCGAATACCGGTC
 BioSequences.FASTA.Record:
   identifier: RfvP
  description: <missing>
     sequence: CTGCAAGGTCGAATCCGGTC
 BioSequences.FASTA.Record:
   identifier: 9sYB
  description: <missing>
     sequence: GCCCGGATTCGCCTTTGCAG
 BioSequences.FASTA.Record:
   identifier: fR8h
  description: <missing>
     sequence: CTGCAAGGTCGAATCCGGTC
 BioSequences.FASTA.Record:
   identifier: KA0V
  description: <missing>
     sequence: GACCGGAGGAACCTTGCATG
 BioSequences.FASTA.Record:
   identifier: yD6T
  description: <missing>
     sequence: CTGCAAGGTCCATCCGGC
 BioSequences.FASTA.Record:
   identifier: 91tv
  description: <missing>
     sequence: TACCGGATTCGACCTTGCAG
 ⋮
 BioSequences.FASTA.Record:
   identifier: kdBM
  description: <missing>
     sequence: CTAGCAGGTCGAATCCGGTC
 BioSequences.FASTA.Record:
   identifier: 81FM
  description: <missing>
     sequence: GACCGGAGTACGACCTTCGCAAG
 BioSequences.FASTA.Record:
   identifier: hsIV
  description: <missing>
     sequence: CTGTCAAGGTCGAATCCGGTC
 BioSequences.FASTA.Record:
   identifier: Ltb5
  description: <missing>
     sequence: CTGCCAAGGCGAATCCGGC
 BioSequences.FASTA.Record:
   identifier: vZaj
  description: <missing>
     sequence: GACCTGGATTCGACCTTGCAG
 BioSequences.FASTA.Record:
   identifier: j1db
  description: <missing>
     sequence: GACCGGATTCGACCTTGCAG
 BioSequences.FASTA.Record:
   identifier: BkDQ
  description: <missing>
     sequence: CTGAAAGGTCCCAATCCGGTC
 BioSequences.FASTA.Record:
   identifier: 1IcU
  description: <missing>
     sequence: TGACGAAGGTCAAATCCGGTC
 BioSequences.FASTA.Record:
   identifier: 081o
  description: <missing>
     sequence: GTGCACAGATCCGAATCCACGC
 BioSequences.FASTA.Record:
   identifier: ZWqs
  description: <missing>
     sequence: CTGCAAGGTCGATCCCGTTC
 BioSequences.FASTA.Record:
   identifier: RUED
  description: <missing>
     sequence: CTGCACGGCGAATCCGGT
 BioSequences.FASTA.Record:
```

```
  identifier: iVVA
 description: <missing>
    sequence: GACCGGATTAGACCTTGCAG
```

#### L20 starting @ K=7

In [15]:

```
k = 7
canonical_kmers = collect(keys(Eisenia.count_canonical_kmers(observations, k)))
stranded_kmer_graph = Eisenia.build_stranded_kmer_graph(canonical_kmers, observations)
filename = reference_sequence_id * "." * replace(string(Dates.now()), ':' => '.') * ".svg"
Eisenia.plot_stranded_kmer_graph(stranded_kmer_graph, filename=filename)
HTML("""
<image src="$filename" width=50%>
""")
```

Out[15]:

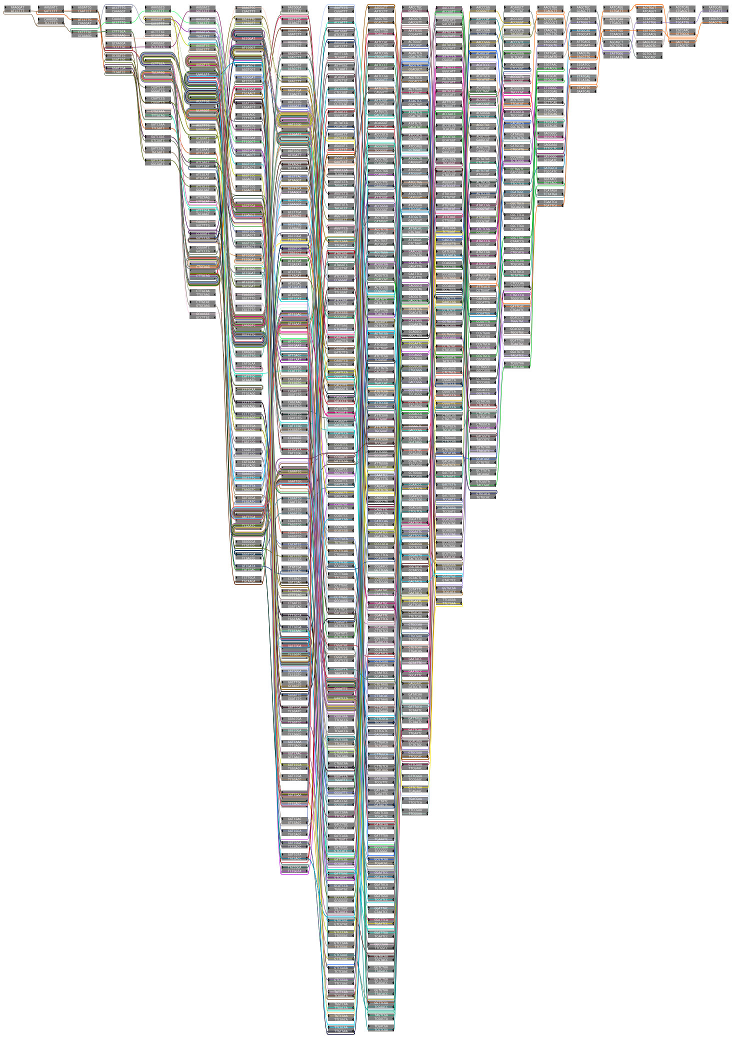

In [16]:

```
maximum_likelihood_observations = Eisenia.viterbi_maximum_likelihood_traversals(stranded_kmer_graph, verbosity="reads");
```

```
computing kmer counts...
computing kmer state likelihoods...
finding shortest paths between kmers...
finding viterbi maximum likelihood paths for observed sequences...

evaluating sequence 1 of 100
        observed sequence                 GACCGGATTCGGCCTGGGAG
        maximum likelihood sequence       GACCGGATTCGACCTTGCAG
        maximum likelihood edit distance  3

evaluating sequence 2 of 100
        observed sequence                 CTGCCCAGGGTCGGAATCCGGTAC
        maximum likelihood sequence       CCTGCAAGGTCGAATCCGGTC
        maximum likelihood edit distance  6

evaluating sequence 3 of 100
        observed sequence                 GATCCGGATTCGATCCTTGCAG
        maximum likelihood sequence       GACCGGATTCGACCTTGCAG
        maximum likelihood edit distance  4

evaluating sequence 4 of 100
        observed sequence                 ACCGAGATATCGACCTTGCAG
        maximum likelihood sequence       GACCGGATTCGACCTTGCAG
        maximum likelihood edit distance  3

evaluating sequence 5 of 100
        observed sequence                 GACCGGATTCCGATGGCAG
        maximum likelihood sequence       GACCGGATTCGACCTTGCAG
        maximum likelihood edit distance  4

evaluating sequence 6 of 100
        observed sequence                 GAACCGGGTTCGACCTTGCAG
        maximum likelihood sequence       GACCGGATTCGACCTTGCAG
        maximum likelihood edit distance  3

evaluating sequence 7 of 100
        observed sequence                 CTGCAGGTTCGAATACCGGTC
        maximum likelihood sequence       CTGCAAGGTCGAATCCGGTC
        maximum likelihood edit distance  4

evaluating sequence 8 of 100
        observed sequence                 CTGCAAGGTCGAATCCGGTC
        maximum likelihood sequence       CTGCAAGGTCGAATCCGGTC
        maximum likelihood edit distance  0

evaluating sequence 9 of 100
        observed sequence                 GCCCGGATTCGCCTTTGCAG
        maximum likelihood sequence       GACCGGATTCGACCTTGCAG
        maximum likelihood edit distance  3

evaluating sequence 10 of 100
        observed sequence                 CTGCAAGGTCGAATCCGGTC
        maximum likelihood sequence       CTGCAAGGTCGAATCCGGTC
        maximum likelihood edit distance  0

evaluating sequence 11 of 100
        observed sequence                 GACCGGAGGAACCTTGCATG
        maximum likelihood sequence       CCGGATTCGACCTTGCATG
        maximum likelihood edit distance  4

evaluating sequence 12 of 100
        observed sequence                 CTGCAAGGTCCATCCGGC
        maximum likelihood sequence       CTGCAAGGTCGAATCCGGT
        maximum likelihood edit distance  3

evaluating sequence 13 of 100
        observed sequence                 TACCGGATTCGACCTTGCAG
        maximum likelihood sequence       GACCGGATTCGACCTTGCAG
        maximum likelihood edit distance  1

evaluating sequence 14 of 100
        observed sequence                 TCAAGCGTCGAATCCCGGTC
        maximum likelihood sequence       TGCAAGGTCGAATCCGGTC
        maximum likelihood edit distance  2

evaluating sequence 15 of 100
        observed sequence                 GAACGGATTACATCCCTGCAG
        maximum likelihood sequence       GACCGGATTCGACCTTGCAG
        maximum likelihood edit distance  6

evaluating sequence 16 of 100
        observed sequence                 ACCGGATTGACCTTGCAA
        maximum likelihood sequence       ACCGGATTCGACCTTGCA
        maximum likelihood edit distance  2

evaluating sequence 17 of 100
        observed sequence                 CCTGCAAGGTCGAATCCTGGGC
        maximum likelihood sequence       CCTGCAAGGTCGAATCCGGTC
        maximum likelihood edit distance  3

evaluating sequence 18 of 100
        observed sequence                 CTGCAAGGTCGAATCCGGTC
        maximum likelihood sequence       CTGCAAGGTCGAATCCGGTC
        maximum likelihood edit distance  0

evaluating sequence 19 of 100
        observed sequence                 CTGCAAGATCCGAATCCGGGTC
        maximum likelihood sequence       CTGCAAGGTCGAATCCGGTC
        maximum likelihood edit distance  3
```

```
evaluating sequence 20 of 100
        observed sequence                  GAGCCGGATCCGACCTTGCCAG
        maximum likelihood sequence        GACCGGATTCGACCTTGCAG
        maximum likelihood edit distance  4

evaluating sequence 21 of 100
        observed sequence                  GACCGGATTCAACTTGCAG
        maximum likelihood sequence        GACCGGATTCGACCTTGCAG
        maximum likelihood edit distance  2

evaluating sequence 22 of 100
        observed sequence                  GACCGGATTGACCTTGGCA
        maximum likelihood sequence        GACCGGATTCGACCTTGCA
        maximum likelihood edit distance  2

evaluating sequence 23 of 100
        observed sequence                  TTTCCTCGTCGAACCCGTC
        maximum likelihood sequence        CTGCAAGGTCGAATCCGGTC
        maximum likelihood edit distance  7

evaluating sequence 24 of 100
        observed sequence                  GACCGGATTCGACCTTGCCAG
        maximum likelihood sequence        GACCGGATTCGACCTTGCAG
        maximum likelihood edit distance  1

evaluating sequence 25 of 100
        observed sequence                  ACGGATTCGACCTTGCAG
        maximum likelihood sequence        ACGGATTCGACCTTGCAG
        maximum likelihood edit distance  0

evaluating sequence 26 of 100
        observed sequence                  CTGACGTCAATGCGTGC
        maximum likelihood sequence        CAAGGTCGAATCCGGTC
        maximum likelihood edit distance  7

evaluating sequence 27 of 100
        observed sequence                  GACCGGTATTCGACCTTGCATG
        maximum likelihood sequence        GACCGGATTCGACCTTGCATG
        maximum likelihood edit distance  2

evaluating sequence 28 of 100
        observed sequence                  CTGCAAGGTCGAATCCGGGTC
        maximum likelihood sequence        CTGCAAGGTCGAATCCGGTC
        maximum likelihood edit distance  1

evaluating sequence 29 of 100
        observed sequence                  GCACCGGATTCGGACCTTGCAG
        maximum likelihood sequence        GACCGGATTCGACCTTGCAG
        maximum likelihood edit distance  3

evaluating sequence 30 of 100
        observed sequence                  GACCGGATTCGACCTTGCATG
        maximum likelihood sequence        GACCGGATTCGACCTTGCATG
        maximum likelihood edit distance  0

evaluating sequence 31 of 100
        observed sequence                  CTGTATAGTCGAATCCGGTC
        maximum likelihood sequence        CTGCAAGGTCGAATCCGGTC
        maximum likelihood edit distance  3

evaluating sequence 32 of 100
        observed sequence                  CTTCAAGGTCGAATCCGTC
        maximum likelihood sequence        CTGCAAGGTCGAATCCGGTC
        maximum likelihood edit distance  2

evaluating sequence 33 of 100
        observed sequence                  CGACAAGGTCAACCGGTC
        maximum likelihood sequence        CTGCAAGGTCGAATCCGGTC
        maximum likelihood edit distance  4

evaluating sequence 34 of 100
        observed sequence                  GAACGGATTCGACCTTGTCAG
        maximum likelihood sequence        GACCGGATTCGACCTTGCAG
        maximum likelihood edit distance  3

evaluating sequence 35 of 100
        observed sequence                  GACCGGATTCGACCTTGCAG
        maximum likelihood sequence        GACCGGATTCGACCTTGCAG
        maximum likelihood edit distance  0

evaluating sequence 36 of 100
        observed sequence                  CTGCAAGGCCGAATCCGGGTC
        maximum likelihood sequence        CTGCAAGGTCGAATCCGGTC
        maximum likelihood edit distance  2

evaluating sequence 37 of 100
        observed sequence                  CCTGCAAGGTCGAATCCCTGGTC
        maximum likelihood sequence        CCTGCAAGGTCGAATCCGGTC
        maximum likelihood edit distance  3

evaluating sequence 38 of 100
        observed sequence                  GATCGGATTCGACCTTGCGAAG
        maximum likelihood sequence        GACCGGATTCGACCTTGCAG
        maximum likelihood edit distance  4

evaluating sequence 39 of 100
        observed sequence                  CTGCATTGGGTGCGAATCCGGTC
        maximum likelihood sequence        CCTGCAAGGTCGAATCCGGTC
```

```
        maximum likelihood edit distance  5

evaluating sequence 40 of 100
        observed sequence                 CTGTAAGGTCGAATACGGGTC
        maximum likelihood sequence       CTGCAAGGTCGAATCCGGTC
        maximum likelihood edit distance  3

evaluating sequence 41 of 100
        observed sequence                 TGACCCGGATTCGACCTCTGCG
        maximum likelihood sequence       TGACCCGGATTCGACCTTGCA
        maximum likelihood edit distance  3

evaluating sequence 42 of 100
        observed sequence                 CTGAAAGGTCAATCTCGGGC
        maximum likelihood sequence       CTGCAAGGTCGAATCCGGTC
        maximum likelihood edit distance  5

evaluating sequence 43 of 100
        observed sequence                 GATCGGATTCGACCTTGCAG
        maximum likelihood sequence       GACCGGATTCGACCTTGCAG
        maximum likelihood edit distance  1

evaluating sequence 44 of 100
        observed sequence                 GTGCACAGCGTCGAGATCCGGTC
        maximum likelihood sequence       CCTGCAAGGTCGAATCCGGTC
        maximum likelihood edit distance  6

evaluating sequence 45 of 100
        observed sequence                 CTTCCAGGTCGAATCCGGTC
        maximum likelihood sequence       CTGCAAGGTCGAATCCGGTC
        maximum likelihood edit distance  2

evaluating sequence 46 of 100
        observed sequence                 GGCCAGGATTCGACGCTTCAG
        maximum likelihood sequence       GGACCGGATTCGACCTTGCAG
        maximum likelihood edit distance  5

evaluating sequence 47 of 100
        observed sequence                 GATCCGGATTCGACGCTGCAG
        maximum likelihood sequence       GACCGGATTCGACCTTGCAG
        maximum likelihood edit distance  4

evaluating sequence 48 of 100
        observed sequence                 GACCGGTTCGACTCTATGAG
        maximum likelihood sequence       ACCGGATTCGACCTTGCAG
        maximum likelihood edit distance  6

evaluating sequence 49 of 100
        observed sequence                 CCTGCAAGTCGAATTCGGTC
        maximum likelihood sequence       CTGCAAGGTCGAATCCGGTC
        maximum likelihood edit distance  2

evaluating sequence 50 of 100
        observed sequence                 GACCGGATACGACCCTGTACG
        maximum likelihood sequence       GACCGGATTCGACCTTGCAG
        maximum likelihood edit distance  4

evaluating sequence 51 of 100
        observed sequence                 CTGCAAGGTCGAATCCGGTC
        maximum likelihood sequence       CTGCAAGGTCGAATCCGGTC
        maximum likelihood edit distance  0

evaluating sequence 52 of 100
        observed sequence                 TAGCAGCTTGTATCCGGTC
        maximum likelihood sequence       TGCAAGGTCGAATCCGGTC
        maximum likelihood edit distance  4

evaluating sequence 53 of 100
        observed sequence                 GACGGATCGACCTTGCAG
        maximum likelihood sequence       ACGGATTCGACCTTGCAG
        maximum likelihood edit distance  1

evaluating sequence 54 of 100
        observed sequence                 GACCGGATTCGACCTTTGCAG
        maximum likelihood sequence       GACCGGATTCGACCTTGCAG
        maximum likelihood edit distance  1

evaluating sequence 55 of 100
        observed sequence                 GACCGGGATTCGACCTTGCAG
        maximum likelihood sequence       GACCGGATTCGACCTTGCAG
        maximum likelihood edit distance  2

evaluating sequence 56 of 100
        observed sequence                 CTGCATAGGTCGGAATCCGGTC
        maximum likelihood sequence       CCTGCAAGGTCGAATCCGGTC
        maximum likelihood edit distance  2

evaluating sequence 57 of 100
        observed sequence                 GAAGGATTCGCACCTTCAG
        maximum likelihood sequence       ACCGGATTCGACCTTGCAG
        maximum likelihood edit distance  5

evaluating sequence 58 of 100
        observed sequence                 CTGCAAGGTCGAATCCGGTC
        maximum likelihood sequence       CTGCAAGGTCGAATCCGGTC
        maximum likelihood edit distance  0

evaluating sequence 59 of 100
        observed sequence                 GACCGGTTACCGACCGTGCAG
```

```
        maximum likelihood sequence       GACCGGATTCGACCTTGCAG
        maximum likelihood edit distance  6

evaluating sequence 60 of 100
        observed sequence                 GACGGTACGACCTTGCCAG
        maximum likelihood sequence       ACGGATTCGACCTTGCAG
        maximum likelihood edit distance  3

evaluating sequence 61 of 100
        observed sequence                 GACCGGATCGACCTTGCAG
        maximum likelihood sequence       ACCGGATTCGACCTTGCAG
        maximum likelihood edit distance  1

evaluating sequence 62 of 100
        observed sequence                 CTGCAAAGTCGAATCCGGTC
        maximum likelihood sequence       CTGCAAGGTCGAATCCGGTC
        maximum likelihood edit distance  1

evaluating sequence 63 of 100
        observed sequence                 CTGCAAGGTCGAATGCCCGTC
        maximum likelihood sequence       CTGCAAGGTCGAATCCGGTC
        maximum likelihood edit distance  3

evaluating sequence 64 of 100
        observed sequence                 GACGGGATTCGACCTTACAC
        maximum likelihood sequence       GACCGGATTCGACCTTGCAG
        maximum likelihood edit distance  3

evaluating sequence 65 of 100
        observed sequence                 CTGCGGGTCGAATCGGTC
        maximum likelihood sequence       TGCAAGGTCGAATCCGGTC
        maximum likelihood edit distance  3

evaluating sequence 66 of 100
        observed sequence                 ACTGCAGGGTCGAATCCGGTC
        maximum likelihood sequence       CTGCAAGGTCGAATCCGGTC
        maximum likelihood edit distance  2

evaluating sequence 67 of 100
        observed sequence                 CTGCAAGGTTCTGAATCCGGCTC
        maximum likelihood sequence       CTGCAAGGTCGAATCCGGTC
        maximum likelihood edit distance  5

evaluating sequence 68 of 100
        observed sequence                 GGACCTGATTCACGTTGGCG
        maximum likelihood sequence       GGACCGGATTCGACCTTGCA
        maximum likelihood edit distance  5

evaluating sequence 69 of 100
        observed sequence                 TGAAGGTCGAATCCGGTC
        maximum likelihood sequence       GCAAGGTCGAATCCGGTC
        maximum likelihood edit distance  2

evaluating sequence 70 of 100
        observed sequence                 CTGCAAGGTCGAATGCGTC
        maximum likelihood sequence       CTGCAAGGTCGAATCCGGTC
        maximum likelihood edit distance  2

evaluating sequence 71 of 100
        observed sequence                 CCTGCAAGGTCTGATCCGGTC
        maximum likelihood sequence       CCTGCAAGGTCGAATCCGGTC
        maximum likelihood edit distance  2

evaluating sequence 72 of 100
        observed sequence                 GACCCGGATTGACCATTTGCAG
        maximum likelihood sequence       GACCGGATTCGACCTTGCAG
        maximum likelihood edit distance  7

evaluating sequence 73 of 100
        observed sequence                 GACCGGATGCGATCCTTTGCAG
        maximum likelihood sequence       GACCGGATTCGACCTTGCAG
        maximum likelihood edit distance  4

evaluating sequence 74 of 100
        observed sequence                 CTGACAAGGTCGAGATCCGGTC
        maximum likelihood sequence       CCTGCAAGGTCGAATCCGGTC
        maximum likelihood edit distance  3

evaluating sequence 75 of 100
        observed sequence                 GACCGGATTCGACCTTTCAG
        maximum likelihood sequence       GACCGGATTCGACCTTGCAG
        maximum likelihood edit distance  1

evaluating sequence 76 of 100
        observed sequence                 GACCGTGAATTGACCTTGCTG
        maximum likelihood sequence       GGACCGGATTCGACCTTGCAG
        maximum likelihood edit distance  4

evaluating sequence 77 of 100
        observed sequence                 CTGCAAGTCGAATCCGGTC
        maximum likelihood sequence       CTGCAAGGTCGAATCCGGTC
        maximum likelihood edit distance  1

evaluating sequence 78 of 100
        observed sequence                 GGATCGGCATTCGGTCCTTGCAG
        maximum likelihood sequence       GGACCGGATTCGACCTTGCAG
        maximum likelihood edit distance  5

evaluating sequence 79 of 100
```

```
        observed sequence                 CTCGCAAGGTCGAATCCGGTG
        maximum likelihood sequence       CTGCAAGGTCGAATCCGGTC
        maximum likelihood edit distance  3

evaluating sequence 80 of 100
        observed sequence                 CTGCATGTCGAATCCGGTC
        maximum likelihood sequence       CTGCAAGGTCGAATCCGGTC
        maximum likelihood edit distance  2

evaluating sequence 81 of 100
        observed sequence                 CCTGCAAGGTTCGGAATCCGTC
        maximum likelihood sequence       CCTGCAAGGTCGAATCCGGTC
        maximum likelihood edit distance  3

evaluating sequence 82 of 100
        observed sequence                 GGACCGGATTCGACCTTGCAG
        maximum likelihood sequence       GGACCGGATTCGACCTTGCAG
        maximum likelihood edit distance  0

evaluating sequence 83 of 100
        observed sequence                 GACCGGTATTCGACCTTGCAG
        maximum likelihood sequence       GACCGGATTCGACCTTGCAG
        maximum likelihood edit distance  2

evaluating sequence 84 of 100
        observed sequence                 CTGCAAGGTCGCATCCAGTC
        maximum likelihood sequence       CTGCAAGGTCGAATCCGGTC
        maximum likelihood edit distance  2

evaluating sequence 85 of 100
        observed sequence                 CTGCGAAGGGTCGATAGTCCGGTTC
        maximum likelihood sequence       CCTGCAAGGTCGAATCCGGTC
        maximum likelihood edit distance  7

evaluating sequence 86 of 100
        observed sequence                 CGTGCAAGGTCGACCGTTC
        maximum likelihood sequence       CCTGCAAGGTCGAATCCGGTC
        maximum likelihood edit distance  4

evaluating sequence 87 of 100
        observed sequence                 GGCCGGATTACACCTTGCAG
        maximum likelihood sequence       GACCGGATTCGACCTTGCAG
        maximum likelihood edit distance  3

evaluating sequence 88 of 100
        observed sequence                 GACGGATCGCCCCGCAG
        maximum likelihood sequence       ACGGATTCGACCTTGCAG
        maximum likelihood edit distance  4

evaluating sequence 89 of 100
        observed sequence                 CTAGCAGGTCGAATCCGGTC
        maximum likelihood sequence       CTGCAAGGTCGAATCCGGTC
        maximum likelihood edit distance  1

evaluating sequence 90 of 100
        observed sequence                 GACCGGAGTACGACCTTCGCAAG
        maximum likelihood sequence       GACCGGATTCGACCTTGCAG
        maximum likelihood edit distance  6

evaluating sequence 91 of 100
        observed sequence                 CTGTCAAGGTCGAATCCGGTC
        maximum likelihood sequence       CCTGCAAGGTCGAATCCGGTC
        maximum likelihood edit distance  1

evaluating sequence 92 of 100
        observed sequence                 CTGCCAAGGCGAATCCGGC
        maximum likelihood sequence       CCTGCAAGGTCGAATCCGGT
        maximum likelihood edit distance  3

evaluating sequence 93 of 100
        observed sequence                 GACCTGGATTCGACCTTGCAG
        maximum likelihood sequence       GGACCGGATTCGACCTTGCAG
        maximum likelihood edit distance  1

evaluating sequence 94 of 100
        observed sequence                 GACCGGATTCGACCTTGCAG
        maximum likelihood sequence       GACCGGATTCGACCTTGCAG
        maximum likelihood edit distance  0

evaluating sequence 95 of 100
        observed sequence                 CTGAAAGGTCCCAATCCGGTC
        maximum likelihood sequence       CTGCAAGGTCGAATCCGGTC
        maximum likelihood edit distance  3

evaluating sequence 96 of 100
        observed sequence                 TGACGAAGGTCAAATCCGGTC
        maximum likelihood sequence       CCTGCAAGGTCGAATCCGGTC
        maximum likelihood edit distance  3

evaluating sequence 97 of 100
        observed sequence                 GTGCACAGATCCGAATCCACGC
        maximum likelihood sequence       CCTGCAAGGTCGAATCCGGTC
        maximum likelihood edit distance  7

evaluating sequence 98 of 100
        observed sequence                 CTGCAAGGTCGATCCCGTTC
        maximum likelihood sequence       CTGCAAGGTCGAATCCGGTC
        maximum likelihood edit distance  3
```

```
evaluating sequence 99 of 100
	observed sequence                 CTGCACGGCGAATCCGGT
	maximum likelihood sequence       CTGCAAGGTCGAATCCGGT
	maximum likelihood edit distance  2

evaluating sequence 100 of 100
	observed sequence                 GACCGGATTAGACCTTGCAG
	maximum likelihood sequence       GACCGGATTCGACCTTGCAG
	maximum likelihood edit distance  1

DATASET STATISTICS:
	assumed error rate    12.5%
	total bases observed  2047
	total edits accepted  289
	inferred error rate   14.118221787982414%
```

In [17]:

```
canonical_kmers = collect(keys(Eisenia.count_canonical_kmers(maximum_likelihood_observations, k)))
stranded_kmer_graph = Eisenia.build_stranded_kmer_graph(canonical_kmers, maximum_likelihood_observations)
filename = reference_sequence_id * "." * replace(string(Dates.now()), ':' => '.') * ".svg"
Eisenia.plot_stranded_kmer_graph(stranded_kmer_graph, filename=filename)
HTML("""
<image src="$filename" width=50%>
""")
```

Out[17]:

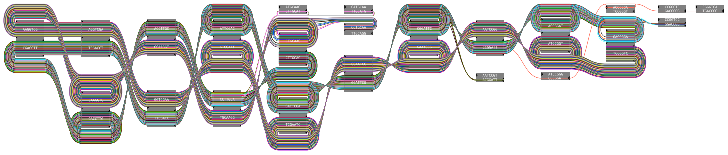

In [18]:

```
maximum_likelihood_observations = Eisenia.viterbi_maximum_likelihood_traversals(stranded_kmer_graph, verbosity="reads");
```

```
computing kmer counts...
computing kmer state likelihoods...
finding shortest paths between kmers...
finding viterbi maximum likelihood paths for observed sequences...

evaluating sequence 1 of 100
        observed sequence                 GACCGGATTCGACCTTGCAG
        maximum likelihood sequence       GACCGGATTCGACCTTGCAG
        maximum likelihood edit distance  0

evaluating sequence 2 of 100
        observed sequence                 CCTGCAAGGTCGAATCCGGTC
        maximum likelihood sequence       CCTGCAAGGTCGAATCCGGTC
        maximum likelihood edit distance  0

evaluating sequence 3 of 100
        observed sequence                 GACCGGATTCGACCTTGCAG
        maximum likelihood sequence       GACCGGATTCGACCTTGCAG
        maximum likelihood edit distance  0

evaluating sequence 4 of 100
        observed sequence                 GACCGGATTCGACCTTGCAG
        maximum likelihood sequence       GACCGGATTCGACCTTGCAG
        maximum likelihood edit distance  0

evaluating sequence 5 of 100
        observed sequence                 GACCGGATTCGACCTTGCAG
        maximum likelihood sequence       GACCGGATTCGACCTTGCAG
        maximum likelihood edit distance  0

evaluating sequence 6 of 100
        observed sequence                 GACCGGATTCGACCTTGCAG
        maximum likelihood sequence       GACCGGATTCGACCTTGCAG
        maximum likelihood edit distance  0

evaluating sequence 7 of 100
        observed sequence                 CTGCAAGGTCGAATCCGGTC
        maximum likelihood sequence       CTGCAAGGTCGAATCCGGTC
        maximum likelihood edit distance  0

evaluating sequence 8 of 100
        observed sequence                 CTGCAAGGTCGAATCCGGTC
        maximum likelihood sequence       CTGCAAGGTCGAATCCGGTC
        maximum likelihood edit distance  0

evaluating sequence 9 of 100
        observed sequence                 GACCGGATTCGACCTTGCAG
        maximum likelihood sequence       GACCGGATTCGACCTTGCAG
        maximum likelihood edit distance  0

evaluating sequence 10 of 100
        observed sequence                 CTGCAAGGTCGAATCCGGTC
        maximum likelihood sequence       CTGCAAGGTCGAATCCGGTC
        maximum likelihood edit distance  0

evaluating sequence 11 of 100
        observed sequence                 CCGGATTCGACCTTGCATG
        maximum likelihood sequence       CCGGATTCGACCTTGCAG
        maximum likelihood edit distance  1

evaluating sequence 12 of 100
        observed sequence                 CTGCAAGGTCGAATCCGGT
        maximum likelihood sequence       CTGCAAGGTCGAATCCGGT
        maximum likelihood edit distance  0

evaluating sequence 13 of 100
        observed sequence                 GACCGGATTCGACCTTGCAG
        maximum likelihood sequence       GACCGGATTCGACCTTGCAG
        maximum likelihood edit distance  0

evaluating sequence 14 of 100
        observed sequence                 TGCAAGGTCGAATCCGGTC
        maximum likelihood sequence       TGCAAGGTCGAATCCGGTC
        maximum likelihood edit distance  0

evaluating sequence 15 of 100
        observed sequence                 GACCGGATTCGACCTTGCAG
        maximum likelihood sequence       GACCGGATTCGACCTTGCAG
        maximum likelihood edit distance  0

evaluating sequence 16 of 100
        observed sequence                 ACCGGATTCGACCTTGCA
        maximum likelihood sequence       ACCGGATTCGACCTTGCA
        maximum likelihood edit distance  0

evaluating sequence 17 of 100
        observed sequence                 CCTGCAAGGTCGAATCCGGTC
        maximum likelihood sequence       CCTGCAAGGTCGAATCCGGTC
        maximum likelihood edit distance  0

evaluating sequence 18 of 100
        observed sequence                 CTGCAAGGTCGAATCCGGTC
        maximum likelihood sequence       CTGCAAGGTCGAATCCGGTC
        maximum likelihood edit distance  0

evaluating sequence 19 of 100
        observed sequence                 CTGCAAGGTCGAATCCGGTC
        maximum likelihood sequence       CTGCAAGGTCGAATCCGGTC
        maximum likelihood edit distance  0
```

```
evaluating sequence 20 of 100
        observed sequence                 GACCGGATTCGACCTTGCAG
        maximum likelihood sequence       GACCGGATTCGACCTTGCAG
        maximum likelihood edit distance  0

evaluating sequence 21 of 100
        observed sequence                 GACCGGATTCGACCTTGCAG
        maximum likelihood sequence       GACCGGATTCGACCTTGCAG
        maximum likelihood edit distance  0

evaluating sequence 22 of 100
        observed sequence                 GACCGGATTCGACCTTGCA
        maximum likelihood sequence       GACCGGATTCGACCTTGCA
        maximum likelihood edit distance  0

evaluating sequence 23 of 100
        observed sequence                 CTGCAAGGTCGAATCCGGTC
        maximum likelihood sequence       CTGCAAGGTCGAATCCGGTC
        maximum likelihood edit distance  0

evaluating sequence 24 of 100
        observed sequence                 GACCGGATTCGACCTTGCAG
        maximum likelihood sequence       GACCGGATTCGACCTTGCAG
        maximum likelihood edit distance  0

evaluating sequence 25 of 100
        observed sequence                 ACGGATTCGACCTTGCAG
        maximum likelihood sequence       CCGGATTCGACCTTGCAG
        maximum likelihood edit distance  1

evaluating sequence 26 of 100
        observed sequence                 CAAGGTCGAATCCGGTC
        maximum likelihood sequence       CAAGGTCGAATCCGGTC
        maximum likelihood edit distance  0

evaluating sequence 27 of 100
        observed sequence                 GACCGGATTCGACCTTGCATG
        maximum likelihood sequence       GACCGGATTCGACCTTGCAG
        maximum likelihood edit distance  1

evaluating sequence 28 of 100
        observed sequence                 CTGCAAGGTCGAATCCGGTC
        maximum likelihood sequence       CTGCAAGGTCGAATCCGGTC
        maximum likelihood edit distance  0

evaluating sequence 29 of 100
        observed sequence                 GACCGGATTCGACCTTGCAG
        maximum likelihood sequence       GACCGGATTCGACCTTGCAG
        maximum likelihood edit distance  0

evaluating sequence 30 of 100
        observed sequence                 GACCGGATTCGACCTTGCATG
        maximum likelihood sequence       GACCGGATTCGACCTTGCAG
        maximum likelihood edit distance  1

evaluating sequence 31 of 100
        observed sequence                 CTGCAAGGTCGAATCCGGTC
        maximum likelihood sequence       CTGCAAGGTCGAATCCGGTC
        maximum likelihood edit distance  0

evaluating sequence 32 of 100
        observed sequence                 CTGCAAGGTCGAATCCGGTC
        maximum likelihood sequence       CTGCAAGGTCGAATCCGGTC
        maximum likelihood edit distance  0

evaluating sequence 33 of 100
        observed sequence                 CTGCAAGGTCGAATCCGGTC
        maximum likelihood sequence       CTGCAAGGTCGAATCCGGTC
        maximum likelihood edit distance  0

evaluating sequence 34 of 100
        observed sequence                 GACCGGATTCGACCTTGCAG
        maximum likelihood sequence       GACCGGATTCGACCTTGCAG
        maximum likelihood edit distance  0

evaluating sequence 35 of 100
        observed sequence                 GACCGGATTCGACCTTGCAG
        maximum likelihood sequence       GACCGGATTCGACCTTGCAG
        maximum likelihood edit distance  0

evaluating sequence 36 of 100
        observed sequence                 CTGCAAGGTCGAATCCGGTC
        maximum likelihood sequence       CTGCAAGGTCGAATCCGGTC
        maximum likelihood edit distance  0

evaluating sequence 37 of 100
        observed sequence                 CCTGCAAGGTCGAATCCGGTC
        maximum likelihood sequence       CCTGCAAGGTCGAATCCGGTC
        maximum likelihood edit distance  0

evaluating sequence 38 of 100
        observed sequence                 GACCGGATTCGACCTTGCAG
        maximum likelihood sequence       GACCGGATTCGACCTTGCAG
        maximum likelihood edit distance  0

evaluating sequence 39 of 100
        observed sequence                 CCTGCAAGGTCGAATCCGGTC
        maximum likelihood sequence       CCTGCAAGGTCGAATCCGGTC
```

```
        maximum likelihood edit distance  0

evaluating sequence 40 of 100
        observed sequence                 CTGCAAGGTCGAATCCGGTC
        maximum likelihood sequence       CTGCAAGGTCGAATCCGGTC
        maximum likelihood edit distance  0

evaluating sequence 41 of 100
        observed sequence                 TGACCCGGATTCGACCTTGCA
        maximum likelihood sequence       GGACCGGATTCGACCTTGCA
        maximum likelihood edit distance  3

evaluating sequence 42 of 100
        observed sequence                 CTGCAAGGTCGAATCCGGTC
        maximum likelihood sequence       CTGCAAGGTCGAATCCGGTC
        maximum likelihood edit distance  0

evaluating sequence 43 of 100
        observed sequence                 GACCGGATTCGACCTTGCAG
        maximum likelihood sequence       GACCGGATTCGACCTTGCAG
        maximum likelihood edit distance  0

evaluating sequence 44 of 100
        observed sequence                 CCTGCAAGGTCGAATCCGGTC
        maximum likelihood sequence       CCTGCAAGGTCGAATCCGGTC
        maximum likelihood edit distance  0

evaluating sequence 45 of 100
        observed sequence                 CTGCAAGGTCGAATCCGGTC
        maximum likelihood sequence       CTGCAAGGTCGAATCCGGTC
        maximum likelihood edit distance  0

evaluating sequence 46 of 100
        observed sequence                 GGACCGGATTCGACCTTGCAG
        maximum likelihood sequence       GGACCGGATTCGACCTTGCAG
        maximum likelihood edit distance  0

evaluating sequence 47 of 100
        observed sequence                 GACCGGATTCGACCTTGCAG
        maximum likelihood sequence       GACCGGATTCGACCTTGCAG
        maximum likelihood edit distance  0

evaluating sequence 48 of 100
        observed sequence                 ACCGGATTCGACCTTGCAG
        maximum likelihood sequence       ACCGGATTCGACCTTGCAG
        maximum likelihood edit distance  0

evaluating sequence 49 of 100
        observed sequence                 CTGCAAGGTCGAATCCGGTC
        maximum likelihood sequence       CTGCAAGGTCGAATCCGGTC
        maximum likelihood edit distance  0

evaluating sequence 50 of 100
        observed sequence                 GACCGGATTCGACCTTGCAG
        maximum likelihood sequence       GACCGGATTCGACCTTGCAG
        maximum likelihood edit distance  0

evaluating sequence 51 of 100
        observed sequence                 CTGCAAGGTCGAATCCGGTC
        maximum likelihood sequence       CTGCAAGGTCGAATCCGGTC
        maximum likelihood edit distance  0

evaluating sequence 52 of 100
        observed sequence                 TGCAAGGTCGAATCCGGTC
        maximum likelihood sequence       TGCAAGGTCGAATCCGGTC
        maximum likelihood edit distance  0

evaluating sequence 53 of 100
        observed sequence                 ACGGATTCGACCTTGCAG
        maximum likelihood sequence       CCGGATTCGACCTTGCAG
        maximum likelihood edit distance  1

evaluating sequence 54 of 100
        observed sequence                 GACCGGATTCGACCTTGCAG
        maximum likelihood sequence       GACCGGATTCGACCTTGCAG
        maximum likelihood edit distance  0

evaluating sequence 55 of 100
        observed sequence                 GACCGGATTCGACCTTGCAG
        maximum likelihood sequence       GACCGGATTCGACCTTGCAG
        maximum likelihood edit distance  0

evaluating sequence 56 of 100
        observed sequence                 CCTGCAAGGTCGAATCCGGTC
        maximum likelihood sequence       CCTGCAAGGTCGAATCCGGTC
        maximum likelihood edit distance  0

evaluating sequence 57 of 100
        observed sequence                 ACCGGATTCGACCTTGCAG
        maximum likelihood sequence       ACCGGATTCGACCTTGCAG
        maximum likelihood edit distance  0

evaluating sequence 58 of 100
        observed sequence                 CTGCAAGGTCGAATCCGGTC
        maximum likelihood sequence       CTGCAAGGTCGAATCCGGTC
        maximum likelihood edit distance  0

evaluating sequence 59 of 100
        observed sequence                 GACCGGATTCGACCTTGCAG
```

```
        maximum likelihood sequence       GACCGGATTCGACCTTGCAG
        maximum likelihood edit distance  0

evaluating sequence 60 of 100
        observed sequence                 ACGGATTCGACCTTGCAG
        maximum likelihood sequence       CCGGATTCGACCTTGCAG
        maximum likelihood edit distance  1

evaluating sequence 61 of 100
        observed sequence                 ACCGGATTCGACCTTGCAG
        maximum likelihood sequence       ACCGGATTCGACCTTGCAG
        maximum likelihood edit distance  0

evaluating sequence 62 of 100
        observed sequence                 CTGCAAGGTCGAATCCGGTC
        maximum likelihood sequence       CTGCAAGGTCGAATCCGGTC
        maximum likelihood edit distance  0

evaluating sequence 63 of 100
        observed sequence                 CTGCAAGGTCGAATCCGGTC
        maximum likelihood sequence       CTGCAAGGTCGAATCCGGTC
        maximum likelihood edit distance  0

evaluating sequence 64 of 100
        observed sequence                 GACCGGATTCGACCTTGCAG
        maximum likelihood sequence       GACCGGATTCGACCTTGCAG
        maximum likelihood edit distance  0

evaluating sequence 65 of 100
        observed sequence                 TGCAAGGTCGAATCCGGTC
        maximum likelihood sequence       TGCAAGGTCGAATCCGGTC
        maximum likelihood edit distance  0

evaluating sequence 66 of 100
        observed sequence                 CTGCAAGGTCGAATCCGGTC
        maximum likelihood sequence       CTGCAAGGTCGAATCCGGTC
        maximum likelihood edit distance  0

evaluating sequence 67 of 100
        observed sequence                 CTGCAAGGTCGAATCCGGTC
        maximum likelihood sequence       CTGCAAGGTCGAATCCGGTC
        maximum likelihood edit distance  0

evaluating sequence 68 of 100
        observed sequence                 GGACCGGATTCGACCTTGCA
        maximum likelihood sequence       GGACCGGATTCGACCTTGCA
        maximum likelihood edit distance  0

evaluating sequence 69 of 100
        observed sequence                 GCAAGGTCGAATCCGGTC
        maximum likelihood sequence       GCAAGGTCGAATCCGGTC
        maximum likelihood edit distance  0

evaluating sequence 70 of 100
        observed sequence                 CTGCAAGGTCGAATCCGGTC
        maximum likelihood sequence       CTGCAAGGTCGAATCCGGTC
        maximum likelihood edit distance  0

evaluating sequence 71 of 100
        observed sequence                 CCTGCAAGGTCGAATCCGGTC
        maximum likelihood sequence       CCTGCAAGGTCGAATCCGGTC
        maximum likelihood edit distance  0

evaluating sequence 72 of 100
        observed sequence                 GACCGGATTCGACCTTGCAG
        maximum likelihood sequence       GACCGGATTCGACCTTGCAG
        maximum likelihood edit distance  0

evaluating sequence 73 of 100
        observed sequence                 GACCGGATTCGACCTTGCAG
        maximum likelihood sequence       GACCGGATTCGACCTTGCAG
        maximum likelihood edit distance  0

evaluating sequence 74 of 100
        observed sequence                 CCTGCAAGGTCGAATCCGGTC
        maximum likelihood sequence       CCTGCAAGGTCGAATCCGGTC
        maximum likelihood edit distance  0

evaluating sequence 75 of 100
        observed sequence                 GACCGGATTCGACCTTGCAG
        maximum likelihood sequence       GACCGGATTCGACCTTGCAG
        maximum likelihood edit distance  0

evaluating sequence 76 of 100
        observed sequence                 GGACCGGATTCGACCTTGCAG
        maximum likelihood sequence       GGACCGGATTCGACCTTGCAG
        maximum likelihood edit distance  0

evaluating sequence 77 of 100
        observed sequence                 CTGCAAGGTCGAATCCGGTC
        maximum likelihood sequence       CTGCAAGGTCGAATCCGGTC
        maximum likelihood edit distance  0

evaluating sequence 78 of 100
        observed sequence                 GGACCGGATTCGACCTTGCAG
        maximum likelihood sequence       GGACCGGATTCGACCTTGCAG
        maximum likelihood edit distance  0

evaluating sequence 79 of 100
```

```
        observed sequence                 CTGCAAGGTCGAATCCGGTC
        maximum likelihood sequence       CTGCAAGGTCGAATCCGGTC
        maximum likelihood edit distance  0

evaluating sequence 80 of 100
        observed sequence                 CTGCAAGGTCGAATCCGGTC
        maximum likelihood sequence       CTGCAAGGTCGAATCCGGTC
        maximum likelihood edit distance  0

evaluating sequence 81 of 100
        observed sequence                 CCTGCAAGGTCGAATCCGGTC
        maximum likelihood sequence       CCTGCAAGGTCGAATCCGGTC
        maximum likelihood edit distance  0

evaluating sequence 82 of 100
        observed sequence                 GGACCGGATTCGACCTTGCAG
        maximum likelihood sequence       GGACCGGATTCGACCTTGCAG
        maximum likelihood edit distance  0

evaluating sequence 83 of 100
        observed sequence                 GACCGGATTCGACCTTGCAG
        maximum likelihood sequence       GACCGGATTCGACCTTGCAG
        maximum likelihood edit distance  0

evaluating sequence 84 of 100
        observed sequence                 CTGCAAGGTCGAATCCGGTC
        maximum likelihood sequence       CTGCAAGGTCGAATCCGGTC
        maximum likelihood edit distance  0

evaluating sequence 85 of 100
        observed sequence                 CCTGCAAGGTCGAATCCGGTC
        maximum likelihood sequence       CCTGCAAGGTCGAATCCGGTC
        maximum likelihood edit distance  0

evaluating sequence 86 of 100
        observed sequence                 CCTGCAAGGTCGAATCCGGTC
        maximum likelihood sequence       CCTGCAAGGTCGAATCCGGTC
        maximum likelihood edit distance  0

evaluating sequence 87 of 100
        observed sequence                 GACCGGATTCGACCTTGCAG
        maximum likelihood sequence       GACCGGATTCGACCTTGCAG
        maximum likelihood edit distance  0

evaluating sequence 88 of 100
        observed sequence                 ACGGATTCGACCTTGCAG
        maximum likelihood sequence       CCGGATTCGACCTTGCAG
        maximum likelihood edit distance  1

evaluating sequence 89 of 100
        observed sequence                 CTGCAAGGTCGAATCCGGTC
        maximum likelihood sequence       CTGCAAGGTCGAATCCGGTC
        maximum likelihood edit distance  0

evaluating sequence 90 of 100
        observed sequence                 GACCGGATTCGACCTTGCAG
        maximum likelihood sequence       GACCGGATTCGACCTTGCAG
        maximum likelihood edit distance  0

evaluating sequence 91 of 100
        observed sequence                 CCTGCAAGGTCGAATCCGGTC
        maximum likelihood sequence       CCTGCAAGGTCGAATCCGGTC
        maximum likelihood edit distance  0

evaluating sequence 92 of 100
        observed sequence                 CCTGCAAGGTCGAATCCGGT
        maximum likelihood sequence       CCTGCAAGGTCGAATCCGGT
        maximum likelihood edit distance  0

evaluating sequence 93 of 100
        observed sequence                 GGACCGGATTCGACCTTGCAG
        maximum likelihood sequence       GGACCGGATTCGACCTTGCAG
        maximum likelihood edit distance  0

evaluating sequence 94 of 100
        observed sequence                 GACCGGATTCGACCTTGCAG
        maximum likelihood sequence       GACCGGATTCGACCTTGCAG
        maximum likelihood edit distance  0

evaluating sequence 95 of 100
        observed sequence                 CTGCAAGGTCGAATCCGGTC
        maximum likelihood sequence       CTGCAAGGTCGAATCCGGTC
        maximum likelihood edit distance  0

evaluating sequence 96 of 100
        observed sequence                 CCTGCAAGGTCGAATCCGGTC
        maximum likelihood sequence       CCTGCAAGGTCGAATCCGGTC
        maximum likelihood edit distance  0

evaluating sequence 97 of 100
        observed sequence                 CCTGCAAGGTCGAATCCGGTC
        maximum likelihood sequence       CCTGCAAGGTCGAATCCGGTC
        maximum likelihood edit distance  0

evaluating sequence 98 of 100
        observed sequence                 CTGCAAGGTCGAATCCGGTC
        maximum likelihood sequence       CTGCAAGGTCGAATCCGGTC
        maximum likelihood edit distance  0
```

```
evaluating sequence 99 of 100
	observed sequence                 CTGCAAGGTCGAATCCGGT
	maximum likelihood sequence       CTGCAAGGTCGAATCCGGT
	maximum likelihood edit distance  0

evaluating sequence 100 of 100
	observed sequence                 GACCGGATTCGACCTTGCAG
	maximum likelihood sequence       GACCGGATTCGACCTTGCAG
	maximum likelihood edit distance  0

DATASET STATISTICS:
	assumed error rate    12.5%
	total bases observed  1997
	total edits accepted  10
	inferred error rate   0.5007511266900351%
```

In [19]:

```
canonical_kmers = collect(keys(Eisenia.count_canonical_kmers(maximum_likelihood_observations, k)))
stranded_kmer_graph = Eisenia.build_stranded_kmer_graph(canonical_kmers, maximum_likelihood_observations)
filename = reference_sequence_id * "." * replace(string(Dates.now()), ':' => '.') * ".svg"
Eisenia.plot_stranded_kmer_graph(stranded_kmer_graph, filename=filename)
HTML("""
<image src="$filename" width=50%>
""")
```

Out[19]:

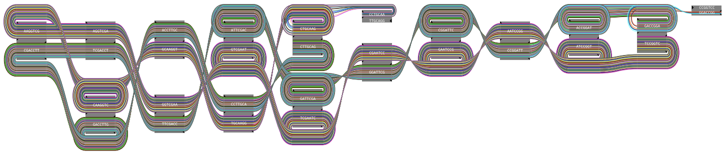

In [20]:

```
maximum_likelihood_observations = Eisenia.viterbi_maximum_likelihood_traversals(stranded_kmer_graph, verbosity="reads");
```

```
computing kmer counts...
computing kmer state likelihoods...
finding shortest paths between kmers...
finding viterbi maximum likelihood paths for observed sequences...

evaluating sequence 1 of 100
        observed sequence                 GACCGGATTCGACCTTGCAG
        maximum likelihood sequence       GACCGGATTCGACCTTGCAG
        maximum likelihood edit distance  0

evaluating sequence 2 of 100
        observed sequence                 CCTGCAAGGTCGAATCCGGTC
        maximum likelihood sequence       CCTGCAAGGTCGAATCCGGTC
        maximum likelihood edit distance  0

evaluating sequence 3 of 100
        observed sequence                 GACCGGATTCGACCTTGCAG
        maximum likelihood sequence       GACCGGATTCGACCTTGCAG
        maximum likelihood edit distance  0

evaluating sequence 4 of 100
        observed sequence                 GACCGGATTCGACCTTGCAG
        maximum likelihood sequence       GACCGGATTCGACCTTGCAG
        maximum likelihood edit distance  0

evaluating sequence 5 of 100
        observed sequence                 GACCGGATTCGACCTTGCAG
        maximum likelihood sequence       GACCGGATTCGACCTTGCAG
        maximum likelihood edit distance  0

evaluating sequence 6 of 100
        observed sequence                 GACCGGATTCGACCTTGCAG
        maximum likelihood sequence       GACCGGATTCGACCTTGCAG
        maximum likelihood edit distance  0

evaluating sequence 7 of 100
        observed sequence                 CTGCAAGGTCGAATCCGGTC
        maximum likelihood sequence       CTGCAAGGTCGAATCCGGTC
        maximum likelihood edit distance  0

evaluating sequence 8 of 100
        observed sequence                 CTGCAAGGTCGAATCCGGTC
        maximum likelihood sequence       CTGCAAGGTCGAATCCGGTC
        maximum likelihood edit distance  0

evaluating sequence 9 of 100
        observed sequence                 GACCGGATTCGACCTTGCAG
        maximum likelihood sequence       GACCGGATTCGACCTTGCAG
        maximum likelihood edit distance  0

evaluating sequence 10 of 100
        observed sequence                 CTGCAAGGTCGAATCCGGTC
        maximum likelihood sequence       CTGCAAGGTCGAATCCGGTC
        maximum likelihood edit distance  0

evaluating sequence 11 of 100
        observed sequence                 CCGGATTCGACCTTGCAG
        maximum likelihood sequence       CCGGATTCGACCTTGCAG
        maximum likelihood edit distance  0

evaluating sequence 12 of 100
        observed sequence                 CTGCAAGGTCGAATCCGGT
        maximum likelihood sequence       CTGCAAGGTCGAATCCGGT
        maximum likelihood edit distance  0

evaluating sequence 13 of 100
        observed sequence                 GACCGGATTCGACCTTGCAG
        maximum likelihood sequence       GACCGGATTCGACCTTGCAG
        maximum likelihood edit distance  0

evaluating sequence 14 of 100
        observed sequence                 TGCAAGGTCGAATCCGGTC
        maximum likelihood sequence       TGCAAGGTCGAATCCGGTC
        maximum likelihood edit distance  0

evaluating sequence 15 of 100
        observed sequence                 GACCGGATTCGACCTTGCAG
        maximum likelihood sequence       GACCGGATTCGACCTTGCAG
        maximum likelihood edit distance  0

evaluating sequence 16 of 100
        observed sequence                 ACCGGATTCGACCTTGCA
        maximum likelihood sequence       ACCGGATTCGACCTTGCA
        maximum likelihood edit distance  0

evaluating sequence 17 of 100
        observed sequence                 CCTGCAAGGTCGAATCCGGTC
        maximum likelihood sequence       CCTGCAAGGTCGAATCCGGTC
        maximum likelihood edit distance  0

evaluating sequence 18 of 100
        observed sequence                 CTGCAAGGTCGAATCCGGTC
        maximum likelihood sequence       CTGCAAGGTCGAATCCGGTC
        maximum likelihood edit distance  0

evaluating sequence 19 of 100
        observed sequence                 CTGCAAGGTCGAATCCGGTC
        maximum likelihood sequence       CTGCAAGGTCGAATCCGGTC
        maximum likelihood edit distance  0
```

```
evaluating sequence 20 of 100
        observed sequence                 GACCGGATTCGACCTTGCAG
        maximum likelihood sequence       GACCGGATTCGACCTTGCAG
        maximum likelihood edit distance  0

evaluating sequence 21 of 100
        observed sequence                 GACCGGATTCGACCTTGCAG
        maximum likelihood sequence       GACCGGATTCGACCTTGCAG
        maximum likelihood edit distance  0

evaluating sequence 22 of 100
        observed sequence                 GACCGGATTCGACCTTGCA
        maximum likelihood sequence       GACCGGATTCGACCTTGCA
        maximum likelihood edit distance  0

evaluating sequence 23 of 100
        observed sequence                 CTGCAAGGTCGAATCCGGTC
        maximum likelihood sequence       CTGCAAGGTCGAATCCGGTC
        maximum likelihood edit distance  0

evaluating sequence 24 of 100
        observed sequence                 GACCGGATTCGACCTTGCAG
        maximum likelihood sequence       GACCGGATTCGACCTTGCAG
        maximum likelihood edit distance  0

evaluating sequence 25 of 100
        observed sequence                 CCGGATTCGACCTTGCAG
        maximum likelihood sequence       CCGGATTCGACCTTGCAG
        maximum likelihood edit distance  0

evaluating sequence 26 of 100
        observed sequence                 CAAGGTCGAATCCGGTC
        maximum likelihood sequence       CAAGGTCGAATCCGGTC
        maximum likelihood edit distance  0

evaluating sequence 27 of 100
        observed sequence                 GACCGGATTCGACCTTGCAG
        maximum likelihood sequence       GACCGGATTCGACCTTGCAG
        maximum likelihood edit distance  0

evaluating sequence 28 of 100
        observed sequence                 CTGCAAGGTCGAATCCGGTC
        maximum likelihood sequence       CTGCAAGGTCGAATCCGGTC
        maximum likelihood edit distance  0

evaluating sequence 29 of 100
        observed sequence                 GACCGGATTCGACCTTGCAG
        maximum likelihood sequence       GACCGGATTCGACCTTGCAG
        maximum likelihood edit distance  0

evaluating sequence 30 of 100
        observed sequence                 GACCGGATTCGACCTTGCAG
        maximum likelihood sequence       GACCGGATTCGACCTTGCAG
        maximum likelihood edit distance  0

evaluating sequence 31 of 100
        observed sequence                 CTGCAAGGTCGAATCCGGTC
        maximum likelihood sequence       CTGCAAGGTCGAATCCGGTC
        maximum likelihood edit distance  0

evaluating sequence 32 of 100
        observed sequence                 CTGCAAGGTCGAATCCGGTC
        maximum likelihood sequence       CTGCAAGGTCGAATCCGGTC
        maximum likelihood edit distance  0

evaluating sequence 33 of 100
        observed sequence                 CTGCAAGGTCGAATCCGGTC
        maximum likelihood sequence       CTGCAAGGTCGAATCCGGTC
        maximum likelihood edit distance  0

evaluating sequence 34 of 100
        observed sequence                 GACCGGATTCGACCTTGCAG
        maximum likelihood sequence       GACCGGATTCGACCTTGCAG
        maximum likelihood edit distance  0

evaluating sequence 35 of 100
        observed sequence                 GACCGGATTCGACCTTGCAG
        maximum likelihood sequence       GACCGGATTCGACCTTGCAG
        maximum likelihood edit distance  0

evaluating sequence 36 of 100
        observed sequence                 CTGCAAGGTCGAATCCGGTC
        maximum likelihood sequence       CTGCAAGGTCGAATCCGGTC
        maximum likelihood edit distance  0

evaluating sequence 37 of 100
        observed sequence                 CCTGCAAGGTCGAATCCGGTC
        maximum likelihood sequence       CCTGCAAGGTCGAATCCGGTC
        maximum likelihood edit distance  0

evaluating sequence 38 of 100
        observed sequence                 GACCGGATTCGACCTTGCAG
        maximum likelihood sequence       GACCGGATTCGACCTTGCAG
        maximum likelihood edit distance  0

evaluating sequence 39 of 100
        observed sequence                 CCTGCAAGGTCGAATCCGGTC
        maximum likelihood sequence       CCTGCAAGGTCGAATCCGGTC
```

```
        maximum likelihood edit distance  0

evaluating sequence 40 of 100
        observed sequence                 CTGCAAGGTCGAATCCGGTC
        maximum likelihood sequence       CTGCAAGGTCGAATCCGGTC
        maximum likelihood edit distance  0

evaluating sequence 41 of 100
        observed sequence                 GGACCGGATTCGACCTTGCA
        maximum likelihood sequence       GGACCGGATTCGACCTTGCA
        maximum likelihood edit distance  0

evaluating sequence 42 of 100
        observed sequence                 CTGCAAGGTCGAATCCGGTC
        maximum likelihood sequence       CTGCAAGGTCGAATCCGGTC
        maximum likelihood edit distance  0

evaluating sequence 43 of 100
        observed sequence                 GACCGGATTCGACCTTGCAG
        maximum likelihood sequence       GACCGGATTCGACCTTGCAG
        maximum likelihood edit distance  0

evaluating sequence 44 of 100
        observed sequence                 CCTGCAAGGTCGAATCCGGTC
        maximum likelihood sequence       CCTGCAAGGTCGAATCCGGTC
        maximum likelihood edit distance  0

evaluating sequence 45 of 100
        observed sequence                 CTGCAAGGTCGAATCCGGTC
        maximum likelihood sequence       CTGCAAGGTCGAATCCGGTC
        maximum likelihood edit distance  0

evaluating sequence 46 of 100
        observed sequence                 GGACCGGATTCGACCTTGCAG
        maximum likelihood sequence       GGACCGGATTCGACCTTGCAG
        maximum likelihood edit distance  0

evaluating sequence 47 of 100
        observed sequence                 GACCGGATTCGACCTTGCAG
        maximum likelihood sequence       GACCGGATTCGACCTTGCAG
        maximum likelihood edit distance  0

evaluating sequence 48 of 100
        observed sequence                 ACCGGATTCGACCTTGCAG
        maximum likelihood sequence       ACCGGATTCGACCTTGCAG
        maximum likelihood edit distance  0

evaluating sequence 49 of 100
        observed sequence                 CTGCAAGGTCGAATCCGGTC
        maximum likelihood sequence       CTGCAAGGTCGAATCCGGTC
        maximum likelihood edit distance  0

evaluating sequence 50 of 100
        observed sequence                 GACCGGATTCGACCTTGCAG
        maximum likelihood sequence       GACCGGATTCGACCTTGCAG
        maximum likelihood edit distance  0

evaluating sequence 51 of 100
        observed sequence                 CTGCAAGGTCGAATCCGGTC
        maximum likelihood sequence       CTGCAAGGTCGAATCCGGTC
        maximum likelihood edit distance  0

evaluating sequence 52 of 100
        observed sequence                 TGCAAGGTCGAATCCGGTC
        maximum likelihood sequence       TGCAAGGTCGAATCCGGTC
        maximum likelihood edit distance  0

evaluating sequence 53 of 100
        observed sequence                 CCGGATTCGACCTTGCAG
        maximum likelihood sequence       CCGGATTCGACCTTGCAG
        maximum likelihood edit distance  0

evaluating sequence 54 of 100
        observed sequence                 GACCGGATTCGACCTTGCAG
        maximum likelihood sequence       GACCGGATTCGACCTTGCAG
        maximum likelihood edit distance  0

evaluating sequence 55 of 100
        observed sequence                 GACCGGATTCGACCTTGCAG
        maximum likelihood sequence       GACCGGATTCGACCTTGCAG
        maximum likelihood edit distance  0

evaluating sequence 56 of 100
        observed sequence                 CCTGCAAGGTCGAATCCGGTC
        maximum likelihood sequence       CCTGCAAGGTCGAATCCGGTC
        maximum likelihood edit distance  0

evaluating sequence 57 of 100
        observed sequence                 ACCGGATTCGACCTTGCAG
        maximum likelihood sequence       ACCGGATTCGACCTTGCAG
        maximum likelihood edit distance  0

evaluating sequence 58 of 100
        observed sequence                 CTGCAAGGTCGAATCCGGTC
        maximum likelihood sequence       CTGCAAGGTCGAATCCGGTC
        maximum likelihood edit distance  0

evaluating sequence 59 of 100
        observed sequence                 GACCGGATTCGACCTTGCAG
```

```
        maximum likelihood sequence       GACCGGATTCGACCTTGCAG
        maximum likelihood edit distance  0

evaluating sequence 60 of 100
        observed sequence                 CCGGATTCGACCTTGCAG
        maximum likelihood sequence       CCGGATTCGACCTTGCAG
        maximum likelihood edit distance  0

evaluating sequence 61 of 100
        observed sequence                 ACCGGATTCGACCTTGCAG
        maximum likelihood sequence       ACCGGATTCGACCTTGCAG
        maximum likelihood edit distance  0

evaluating sequence 62 of 100
        observed sequence                 CTGCAAGGTCGAATCCGGTC
        maximum likelihood sequence       CTGCAAGGTCGAATCCGGTC
        maximum likelihood edit distance  0

evaluating sequence 63 of 100
        observed sequence                 CTGCAAGGTCGAATCCGGTC
        maximum likelihood sequence       CTGCAAGGTCGAATCCGGTC
        maximum likelihood edit distance  0

evaluating sequence 64 of 100
        observed sequence                 GACCGGATTCGACCTTGCAG
        maximum likelihood sequence       GACCGGATTCGACCTTGCAG
        maximum likelihood edit distance  0

evaluating sequence 65 of 100
        observed sequence                 TGCAAGGTCGAATCCGGTC
        maximum likelihood sequence       TGCAAGGTCGAATCCGGTC
        maximum likelihood edit distance  0

evaluating sequence 66 of 100
        observed sequence                 CTGCAAGGTCGAATCCGGTC
        maximum likelihood sequence       CTGCAAGGTCGAATCCGGTC
        maximum likelihood edit distance  0

evaluating sequence 67 of 100
        observed sequence                 CTGCAAGGTCGAATCCGGTC
        maximum likelihood sequence       CTGCAAGGTCGAATCCGGTC
        maximum likelihood edit distance  0

evaluating sequence 68 of 100
        observed sequence                 GGACCGGATTCGACCTTGCA
        maximum likelihood sequence       GGACCGGATTCGACCTTGCA
        maximum likelihood edit distance  0

evaluating sequence 69 of 100
        observed sequence                 GCAAGGTCGAATCCGGTC
        maximum likelihood sequence       GCAAGGTCGAATCCGGTC
        maximum likelihood edit distance  0

evaluating sequence 70 of 100
        observed sequence                 CTGCAAGGTCGAATCCGGTC
        maximum likelihood sequence       CTGCAAGGTCGAATCCGGTC
        maximum likelihood edit distance  0

evaluating sequence 71 of 100
        observed sequence                 CCTGCAAGGTCGAATCCGGTC
        maximum likelihood sequence       CCTGCAAGGTCGAATCCGGTC
        maximum likelihood edit distance  0

evaluating sequence 72 of 100
        observed sequence                 GACCGGATTCGACCTTGCAG
        maximum likelihood sequence       GACCGGATTCGACCTTGCAG
        maximum likelihood edit distance  0

evaluating sequence 73 of 100
        observed sequence                 GACCGGATTCGACCTTGCAG
        maximum likelihood sequence       GACCGGATTCGACCTTGCAG
        maximum likelihood edit distance  0

evaluating sequence 74 of 100
        observed sequence                 CCTGCAAGGTCGAATCCGGTC
        maximum likelihood sequence       CCTGCAAGGTCGAATCCGGTC
        maximum likelihood edit distance  0

evaluating sequence 75 of 100
        observed sequence                 GACCGGATTCGACCTTGCAG
        maximum likelihood sequence       GACCGGATTCGACCTTGCAG
        maximum likelihood edit distance  0

evaluating sequence 76 of 100
        observed sequence                 GGACCGGATTCGACCTTGCAG
        maximum likelihood sequence       GGACCGGATTCGACCTTGCAG
        maximum likelihood edit distance  0

evaluating sequence 77 of 100
        observed sequence                 CTGCAAGGTCGAATCCGGTC
        maximum likelihood sequence       CTGCAAGGTCGAATCCGGTC
        maximum likelihood edit distance  0

evaluating sequence 78 of 100
        observed sequence                 GGACCGGATTCGACCTTGCAG
        maximum likelihood sequence       GGACCGGATTCGACCTTGCAG
        maximum likelihood edit distance  0

evaluating sequence 79 of 100
```

```
        observed sequence                 CTGCAAGGTCGAATCCGGTC
        maximum likelihood sequence       CTGCAAGGTCGAATCCGGTC
        maximum likelihood edit distance  0

evaluating sequence 80 of 100
        observed sequence                 CTGCAAGGTCGAATCCGGTC
        maximum likelihood sequence       CTGCAAGGTCGAATCCGGTC
        maximum likelihood edit distance  0

evaluating sequence 81 of 100
        observed sequence                 CCTGCAAGGTCGAATCCGGTC
        maximum likelihood sequence       CCTGCAAGGTCGAATCCGGTC
        maximum likelihood edit distance  0

evaluating sequence 82 of 100
        observed sequence                 GGACCGGATTCGACCTTGCAG
        maximum likelihood sequence       GGACCGGATTCGACCTTGCAG
        maximum likelihood edit distance  0

evaluating sequence 83 of 100
        observed sequence                 GACCGGATTCGACCTTGCAG
        maximum likelihood sequence       GACCGGATTCGACCTTGCAG
        maximum likelihood edit distance  0

evaluating sequence 84 of 100
        observed sequence                 CTGCAAGGTCGAATCCGGTC
        maximum likelihood sequence       CTGCAAGGTCGAATCCGGTC
        maximum likelihood edit distance  0

evaluating sequence 85 of 100
        observed sequence                 CCTGCAAGGTCGAATCCGGTC
        maximum likelihood sequence       CCTGCAAGGTCGAATCCGGTC
        maximum likelihood edit distance  0

evaluating sequence 86 of 100
        observed sequence                 CCTGCAAGGTCGAATCCGGTC
        maximum likelihood sequence       CCTGCAAGGTCGAATCCGGTC
        maximum likelihood edit distance  0

evaluating sequence 87 of 100
        observed sequence                 GACCGGATTCGACCTTGCAG
        maximum likelihood sequence       GACCGGATTCGACCTTGCAG
        maximum likelihood edit distance  0

evaluating sequence 88 of 100
        observed sequence                 CCGGATTCGACCTTGCAG
        maximum likelihood sequence       CCGGATTCGACCTTGCAG
        maximum likelihood edit distance  0

evaluating sequence 89 of 100
        observed sequence                 CTGCAAGGTCGAATCCGGTC
        maximum likelihood sequence       CTGCAAGGTCGAATCCGGTC
        maximum likelihood edit distance  0

evaluating sequence 90 of 100
        observed sequence                 GACCGGATTCGACCTTGCAG
        maximum likelihood sequence       GACCGGATTCGACCTTGCAG
        maximum likelihood edit distance  0

evaluating sequence 91 of 100
        observed sequence                 CCTGCAAGGTCGAATCCGGTC
        maximum likelihood sequence       CCTGCAAGGTCGAATCCGGTC
        maximum likelihood edit distance  0

evaluating sequence 92 of 100
        observed sequence                 CCTGCAAGGTCGAATCCGGT
        maximum likelihood sequence       CCTGCAAGGTCGAATCCGGT
        maximum likelihood edit distance  0

evaluating sequence 93 of 100
        observed sequence                 GGACCGGATTCGACCTTGCAG
        maximum likelihood sequence       GGACCGGATTCGACCTTGCAG
        maximum likelihood edit distance  0

evaluating sequence 94 of 100
        observed sequence                 GACCGGATTCGACCTTGCAG
        maximum likelihood sequence       GACCGGATTCGACCTTGCAG
        maximum likelihood edit distance  0

evaluating sequence 95 of 100
        observed sequence                 CTGCAAGGTCGAATCCGGTC
        maximum likelihood sequence       CTGCAAGGTCGAATCCGGTC
        maximum likelihood edit distance  0

evaluating sequence 96 of 100
        observed sequence                 CCTGCAAGGTCGAATCCGGTC
        maximum likelihood sequence       CCTGCAAGGTCGAATCCGGTC
        maximum likelihood edit distance  0

evaluating sequence 97 of 100
        observed sequence                 CCTGCAAGGTCGAATCCGGTC
        maximum likelihood sequence       CCTGCAAGGTCGAATCCGGTC
        maximum likelihood edit distance  0

evaluating sequence 98 of 100
        observed sequence                 CTGCAAGGTCGAATCCGGTC
        maximum likelihood sequence       CTGCAAGGTCGAATCCGGTC
        maximum likelihood edit distance  0
```

```
evaluating sequence 99 of 100
        observed sequence                  CTGCAAGGTCGAATCCGGT
        maximum likelihood sequence        CTGCAAGGTCGAATCCGGT
        maximum likelihood edit distance  0

evaluating sequence 100 of 100
        observed sequence                  GACCGGATTCGACCTTGCAG
        maximum likelihood sequence        GACCGGATTCGACCTTGCAG
        maximum likelihood edit distance  0

DATASET STATISTICS:
        assumed error rate    12.5%
        total bases observed  1993
        total edits accepted  0
        inferred error rate   0.0%
```

In [21]:

```
k = 11
canonical_kmers = collect(keys(Eisenia.count_canonical_kmers(maximum_likelihood_observations, k)))
stranded_kmer_graph = Eisenia.build_stranded_kmer_graph(canonical_kmers, maximum_likelihood_observations)
filename = reference_sequence_id * "." * replace(string(Dates.now()), ':' => '.') * ".svg"
Eisenia.plot_stranded_kmer_graph(stranded_kmer_graph, filename=filename)
HTML("""
<image src="$filename" width=50%>
""")
```

Out[21]:

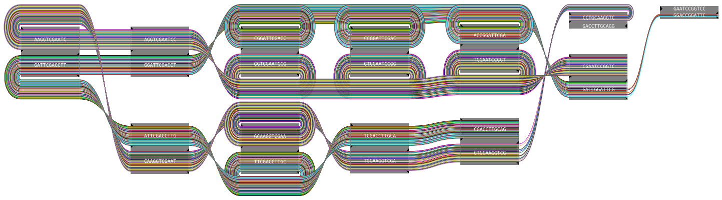

In [22]:

```
maximum_likelihood_observations = Eisenia.viterbi_maximum_likelihood_traversals(stranded_kmer_graph, verbosity="reads");
```

```
computing kmer counts...
computing kmer state likelihoods...
finding shortest paths between kmers...
finding viterbi maximum likelihood paths for observed sequences...

evaluating sequence 1 of 100
        observed sequence                 GACCGGATTCGACCTTGCAG
        maximum likelihood sequence       GACCGGATTCGACCTTGCAG
        maximum likelihood edit distance  0

evaluating sequence 2 of 100
        observed sequence                 CCTGCAAGGTCGAATCCGGTC
        maximum likelihood sequence       CCTGCAAGGTCGAATCCGGTC
        maximum likelihood edit distance  0

evaluating sequence 3 of 100
        observed sequence                 GACCGGATTCGACCTTGCAG
        maximum likelihood sequence       GACCGGATTCGACCTTGCAG
        maximum likelihood edit distance  0

evaluating sequence 4 of 100
        observed sequence                 GACCGGATTCGACCTTGCAG
        maximum likelihood sequence       GACCGGATTCGACCTTGCAG
        maximum likelihood edit distance  0

evaluating sequence 5 of 100
        observed sequence                 GACCGGATTCGACCTTGCAG
        maximum likelihood sequence       GACCGGATTCGACCTTGCAG
        maximum likelihood edit distance  0

evaluating sequence 6 of 100
        observed sequence                 GACCGGATTCGACCTTGCAG
        maximum likelihood sequence       GACCGGATTCGACCTTGCAG
        maximum likelihood edit distance  0

evaluating sequence 7 of 100
        observed sequence                 CTGCAAGGTCGAATCCGGTC
        maximum likelihood sequence       CTGCAAGGTCGAATCCGGTC
        maximum likelihood edit distance  0

evaluating sequence 8 of 100
        observed sequence                 CTGCAAGGTCGAATCCGGTC
        maximum likelihood sequence       CTGCAAGGTCGAATCCGGTC
        maximum likelihood edit distance  0

evaluating sequence 9 of 100
        observed sequence                 GACCGGATTCGACCTTGCAG
        maximum likelihood sequence       GACCGGATTCGACCTTGCAG
        maximum likelihood edit distance  0

evaluating sequence 10 of 100
        observed sequence                 CTGCAAGGTCGAATCCGGTC
        maximum likelihood sequence       CTGCAAGGTCGAATCCGGTC
        maximum likelihood edit distance  0

evaluating sequence 11 of 100
        observed sequence                 CCGGATTCGACCTTGCAG
        maximum likelihood sequence       CCGGATTCGACCTTGCAG
        maximum likelihood edit distance  0

evaluating sequence 12 of 100
        observed sequence                 CTGCAAGGTCGAATCCGGT
        maximum likelihood sequence       CTGCAAGGTCGAATCCGGT
        maximum likelihood edit distance  0

evaluating sequence 13 of 100
        observed sequence                 GACCGGATTCGACCTTGCAG
        maximum likelihood sequence       GACCGGATTCGACCTTGCAG
        maximum likelihood edit distance  0

evaluating sequence 14 of 100
        observed sequence                 TGCAAGGTCGAATCCGGTC
        maximum likelihood sequence       TGCAAGGTCGAATCCGGTC
        maximum likelihood edit distance  0

evaluating sequence 15 of 100
        observed sequence                 GACCGGATTCGACCTTGCAG
        maximum likelihood sequence       GACCGGATTCGACCTTGCAG
        maximum likelihood edit distance  0

evaluating sequence 16 of 100
        observed sequence                 ACCGGATTCGACCTTGCA
        maximum likelihood sequence       ACCGGATTCGACCTTGCA
        maximum likelihood edit distance  0

evaluating sequence 17 of 100
        observed sequence                 CCTGCAAGGTCGAATCCGGTC
        maximum likelihood sequence       CCTGCAAGGTCGAATCCGGTC
        maximum likelihood edit distance  0

evaluating sequence 18 of 100
        observed sequence                 CTGCAAGGTCGAATCCGGTC
        maximum likelihood sequence       CTGCAAGGTCGAATCCGGTC
        maximum likelihood edit distance  0

evaluating sequence 19 of 100
        observed sequence                 CTGCAAGGTCGAATCCGGTC
        maximum likelihood sequence       CTGCAAGGTCGAATCCGGTC
        maximum likelihood edit distance  0
```

```
evaluating sequence 20 of 100
        observed sequence                 GACCGGATTCGACCTTGCAG
        maximum likelihood sequence       GACCGGATTCGACCTTGCAG
        maximum likelihood edit distance  0

evaluating sequence 21 of 100
        observed sequence                 GACCGGATTCGACCTTGCAG
        maximum likelihood sequence       GACCGGATTCGACCTTGCAG
        maximum likelihood edit distance  0

evaluating sequence 22 of 100
        observed sequence                 GACCGGATTCGACCTTGCA
        maximum likelihood sequence       GACCGGATTCGACCTTGCA
        maximum likelihood edit distance  0

evaluating sequence 23 of 100
        observed sequence                 CTGCAAGGTCGAATCCGGTC
        maximum likelihood sequence       CTGCAAGGTCGAATCCGGTC
        maximum likelihood edit distance  0

evaluating sequence 24 of 100
        observed sequence                 GACCGGATTCGACCTTGCAG
        maximum likelihood sequence       GACCGGATTCGACCTTGCAG
        maximum likelihood edit distance  0

evaluating sequence 25 of 100
        observed sequence                 CCGGATTCGACCTTGCAG
        maximum likelihood sequence       CCGGATTCGACCTTGCAG
        maximum likelihood edit distance  0

evaluating sequence 26 of 100
        observed sequence                 CAAGGTCGAATCCGGTC
        maximum likelihood sequence       CAAGGTCGAATCCGGTC
        maximum likelihood edit distance  0

evaluating sequence 27 of 100
        observed sequence                 GACCGGATTCGACCTTGCAG
        maximum likelihood sequence       GACCGGATTCGACCTTGCAG
        maximum likelihood edit distance  0

evaluating sequence 28 of 100
        observed sequence                 CTGCAAGGTCGAATCCGGTC
        maximum likelihood sequence       CTGCAAGGTCGAATCCGGTC
        maximum likelihood edit distance  0

evaluating sequence 29 of 100
        observed sequence                 GACCGGATTCGACCTTGCAG
        maximum likelihood sequence       GACCGGATTCGACCTTGCAG
        maximum likelihood edit distance  0

evaluating sequence 30 of 100
        observed sequence                 GACCGGATTCGACCTTGCAG
        maximum likelihood sequence       GACCGGATTCGACCTTGCAG
        maximum likelihood edit distance  0

evaluating sequence 31 of 100
        observed sequence                 CTGCAAGGTCGAATCCGGTC
        maximum likelihood sequence       CTGCAAGGTCGAATCCGGTC
        maximum likelihood edit distance  0

evaluating sequence 32 of 100
        observed sequence                 CTGCAAGGTCGAATCCGGTC
        maximum likelihood sequence       CTGCAAGGTCGAATCCGGTC
        maximum likelihood edit distance  0

evaluating sequence 33 of 100
        observed sequence                 CTGCAAGGTCGAATCCGGTC
        maximum likelihood sequence       CTGCAAGGTCGAATCCGGTC
        maximum likelihood edit distance  0

evaluating sequence 34 of 100
        observed sequence                 GACCGGATTCGACCTTGCAG
        maximum likelihood sequence       GACCGGATTCGACCTTGCAG
        maximum likelihood edit distance  0

evaluating sequence 35 of 100
        observed sequence                 GACCGGATTCGACCTTGCAG
        maximum likelihood sequence       GACCGGATTCGACCTTGCAG
        maximum likelihood edit distance  0

evaluating sequence 36 of 100
        observed sequence                 CTGCAAGGTCGAATCCGGTC
        maximum likelihood sequence       CTGCAAGGTCGAATCCGGTC
        maximum likelihood edit distance  0

evaluating sequence 37 of 100
        observed sequence                 CCTGCAAGGTCGAATCCGGTC
        maximum likelihood sequence       CCTGCAAGGTCGAATCCGGTC
        maximum likelihood edit distance  0

evaluating sequence 38 of 100
        observed sequence                 GACCGGATTCGACCTTGCAG
        maximum likelihood sequence       GACCGGATTCGACCTTGCAG
        maximum likelihood edit distance  0

evaluating sequence 39 of 100
        observed sequence                 CCTGCAAGGTCGAATCCGGTC
        maximum likelihood sequence       CCTGCAAGGTCGAATCCGGTC
```

```
        maximum likelihood edit distance  0

evaluating sequence 40 of 100
        observed sequence                 CTGCAAGGTCGAATCCGGTC
        maximum likelihood sequence       CTGCAAGGTCGAATCCGGTC
        maximum likelihood edit distance  0

evaluating sequence 41 of 100
        observed sequence                 GGACCGGATTCGACCTTGCA
        maximum likelihood sequence       GGACCGGATTCGACCTTGCA
        maximum likelihood edit distance  0

evaluating sequence 42 of 100
        observed sequence                 CTGCAAGGTCGAATCCGGTC
        maximum likelihood sequence       CTGCAAGGTCGAATCCGGTC
        maximum likelihood edit distance  0

evaluating sequence 43 of 100
        observed sequence                 GACCGGATTCGACCTTGCAG
        maximum likelihood sequence       GACCGGATTCGACCTTGCAG
        maximum likelihood edit distance  0

evaluating sequence 44 of 100
        observed sequence                 CCTGCAAGGTCGAATCCGGTC
        maximum likelihood sequence       CCTGCAAGGTCGAATCCGGTC
        maximum likelihood edit distance  0

evaluating sequence 45 of 100
        observed sequence                 CTGCAAGGTCGAATCCGGTC
        maximum likelihood sequence       CTGCAAGGTCGAATCCGGTC
        maximum likelihood edit distance  0

evaluating sequence 46 of 100
        observed sequence                 GGACCGGATTCGACCTTGCAG
        maximum likelihood sequence       GGACCGGATTCGACCTTGCAG
        maximum likelihood edit distance  0

evaluating sequence 47 of 100
        observed sequence                 GACCGGATTCGACCTTGCAG
        maximum likelihood sequence       GACCGGATTCGACCTTGCAG
        maximum likelihood edit distance  0

evaluating sequence 48 of 100
        observed sequence                 ACCGGATTCGACCTTGCAG
        maximum likelihood sequence       ACCGGATTCGACCTTGCAG
        maximum likelihood edit distance  0

evaluating sequence 49 of 100
        observed sequence                 CTGCAAGGTCGAATCCGGTC
        maximum likelihood sequence       CTGCAAGGTCGAATCCGGTC
        maximum likelihood edit distance  0

evaluating sequence 50 of 100
        observed sequence                 GACCGGATTCGACCTTGCAG
        maximum likelihood sequence       GACCGGATTCGACCTTGCAG
        maximum likelihood edit distance  0

evaluating sequence 51 of 100
        observed sequence                 CTGCAAGGTCGAATCCGGTC
        maximum likelihood sequence       CTGCAAGGTCGAATCCGGTC
        maximum likelihood edit distance  0

evaluating sequence 52 of 100
        observed sequence                 TGCAAGGTCGAATCCGGTC
        maximum likelihood sequence       TGCAAGGTCGAATCCGGTC
        maximum likelihood edit distance  0

evaluating sequence 53 of 100
        observed sequence                 CCGGATTCGACCTTGCAG
        maximum likelihood sequence       CCGGATTCGACCTTGCAG
        maximum likelihood edit distance  0

evaluating sequence 54 of 100
        observed sequence                 GACCGGATTCGACCTTGCAG
        maximum likelihood sequence       GACCGGATTCGACCTTGCAG
        maximum likelihood edit distance  0

evaluating sequence 55 of 100
        observed sequence                 GACCGGATTCGACCTTGCAG
        maximum likelihood sequence       GACCGGATTCGACCTTGCAG
        maximum likelihood edit distance  0

evaluating sequence 56 of 100
        observed sequence                 CCTGCAAGGTCGAATCCGGTC
        maximum likelihood sequence       CCTGCAAGGTCGAATCCGGTC
        maximum likelihood edit distance  0

evaluating sequence 57 of 100
        observed sequence                 ACCGGATTCGACCTTGCAG
        maximum likelihood sequence       ACCGGATTCGACCTTGCAG
        maximum likelihood edit distance  0

evaluating sequence 58 of 100
        observed sequence                 CTGCAAGGTCGAATCCGGTC
        maximum likelihood sequence       CTGCAAGGTCGAATCCGGTC
        maximum likelihood edit distance  0

evaluating sequence 59 of 100
        observed sequence                 GACCGGATTCGACCTTGCAG
```

```
        maximum likelihood sequence       GACCGGATTCGACCTTGCAG
        maximum likelihood edit distance  0

evaluating sequence 60 of 100
        observed sequence                 CCGGATTCGACCTTGCAG
        maximum likelihood sequence       CCGGATTCGACCTTGCAG
        maximum likelihood edit distance  0

evaluating sequence 61 of 100
        observed sequence                 ACCGGATTCGACCTTGCAG
        maximum likelihood sequence       ACCGGATTCGACCTTGCAG
        maximum likelihood edit distance  0

evaluating sequence 62 of 100
        observed sequence                 CTGCAAGGTCGAATCCGGTC
        maximum likelihood sequence       CTGCAAGGTCGAATCCGGTC
        maximum likelihood edit distance  0

evaluating sequence 63 of 100
        observed sequence                 CTGCAAGGTCGAATCCGGTC
        maximum likelihood sequence       CTGCAAGGTCGAATCCGGTC
        maximum likelihood edit distance  0

evaluating sequence 64 of 100
        observed sequence                 GACCGGATTCGACCTTGCAG
        maximum likelihood sequence       GACCGGATTCGACCTTGCAG
        maximum likelihood edit distance  0

evaluating sequence 65 of 100
        observed sequence                 TGCAAGGTCGAATCCGGTC
        maximum likelihood sequence       TGCAAGGTCGAATCCGGTC
        maximum likelihood edit distance  0

evaluating sequence 66 of 100
        observed sequence                 CTGCAAGGTCGAATCCGGTC
        maximum likelihood sequence       CTGCAAGGTCGAATCCGGTC
        maximum likelihood edit distance  0

evaluating sequence 67 of 100
        observed sequence                 CTGCAAGGTCGAATCCGGTC
        maximum likelihood sequence       CTGCAAGGTCGAATCCGGTC
        maximum likelihood edit distance  0

evaluating sequence 68 of 100
        observed sequence                 GGACCGGATTCGACCTTGCA
        maximum likelihood sequence       GGACCGGATTCGACCTTGCA
        maximum likelihood edit distance  0

evaluating sequence 69 of 100
        observed sequence                 GCAAGGTCGAATCCGGTC
        maximum likelihood sequence       GCAAGGTCGAATCCGGTC
        maximum likelihood edit distance  0

evaluating sequence 70 of 100
        observed sequence                 CTGCAAGGTCGAATCCGGTC
        maximum likelihood sequence       CTGCAAGGTCGAATCCGGTC
        maximum likelihood edit distance  0

evaluating sequence 71 of 100
        observed sequence                 CCTGCAAGGTCGAATCCGGTC
        maximum likelihood sequence       CCTGCAAGGTCGAATCCGGTC
        maximum likelihood edit distance  0

evaluating sequence 72 of 100
        observed sequence                 GACCGGATTCGACCTTGCAG
        maximum likelihood sequence       GACCGGATTCGACCTTGCAG
        maximum likelihood edit distance  0

evaluating sequence 73 of 100
        observed sequence                 GACCGGATTCGACCTTGCAG
        maximum likelihood sequence       GACCGGATTCGACCTTGCAG
        maximum likelihood edit distance  0

evaluating sequence 74 of 100
        observed sequence                 CCTGCAAGGTCGAATCCGGTC
        maximum likelihood sequence       CCTGCAAGGTCGAATCCGGTC
        maximum likelihood edit distance  0

evaluating sequence 75 of 100
        observed sequence                 GACCGGATTCGACCTTGCAG
        maximum likelihood sequence       GACCGGATTCGACCTTGCAG
        maximum likelihood edit distance  0

evaluating sequence 76 of 100
        observed sequence                 GGACCGGATTCGACCTTGCAG
        maximum likelihood sequence       GGACCGGATTCGACCTTGCAG
        maximum likelihood edit distance  0

evaluating sequence 77 of 100
        observed sequence                 CTGCAAGGTCGAATCCGGTC
        maximum likelihood sequence       CTGCAAGGTCGAATCCGGTC
        maximum likelihood edit distance  0

evaluating sequence 78 of 100
        observed sequence                 GGACCGGATTCGACCTTGCAG
        maximum likelihood sequence       GGACCGGATTCGACCTTGCAG
        maximum likelihood edit distance  0

evaluating sequence 79 of 100
```

```
        observed sequence                 CTGCAAGGTCGAATCCGGTC
        maximum likelihood sequence       CTGCAAGGTCGAATCCGGTC
        maximum likelihood edit distance  0

evaluating sequence 80 of 100
        observed sequence                 CTGCAAGGTCGAATCCGGTC
        maximum likelihood sequence       CTGCAAGGTCGAATCCGGTC
        maximum likelihood edit distance  0

evaluating sequence 81 of 100
        observed sequence                 CCTGCAAGGTCGAATCCGGTC
        maximum likelihood sequence       CCTGCAAGGTCGAATCCGGTC
        maximum likelihood edit distance  0

evaluating sequence 82 of 100
        observed sequence                 GGACCGGATTCGACCTTGCAG
        maximum likelihood sequence       GGACCGGATTCGACCTTGCAG
        maximum likelihood edit distance  0

evaluating sequence 83 of 100
        observed sequence                 GACCGGATTCGACCTTGCAG
        maximum likelihood sequence       GACCGGATTCGACCTTGCAG
        maximum likelihood edit distance  0

evaluating sequence 84 of 100
        observed sequence                 CTGCAAGGTCGAATCCGGTC
        maximum likelihood sequence       CTGCAAGGTCGAATCCGGTC
        maximum likelihood edit distance  0

evaluating sequence 85 of 100
        observed sequence                 CCTGCAAGGTCGAATCCGGTC
        maximum likelihood sequence       CCTGCAAGGTCGAATCCGGTC
        maximum likelihood edit distance  0

evaluating sequence 86 of 100
        observed sequence                 CCTGCAAGGTCGAATCCGGTC
        maximum likelihood sequence       CCTGCAAGGTCGAATCCGGTC
        maximum likelihood edit distance  0

evaluating sequence 87 of 100
        observed sequence                 GACCGGATTCGACCTTGCAG
        maximum likelihood sequence       GACCGGATTCGACCTTGCAG
        maximum likelihood edit distance  0

evaluating sequence 88 of 100
        observed sequence                 CCGGATTCGACCTTGCAG
        maximum likelihood sequence       CCGGATTCGACCTTGCAG
        maximum likelihood edit distance  0

evaluating sequence 89 of 100
        observed sequence                 CTGCAAGGTCGAATCCGGTC
        maximum likelihood sequence       CTGCAAGGTCGAATCCGGTC
        maximum likelihood edit distance  0

evaluating sequence 90 of 100
        observed sequence                 GACCGGATTCGACCTTGCAG
        maximum likelihood sequence       GACCGGATTCGACCTTGCAG
        maximum likelihood edit distance  0

evaluating sequence 91 of 100
        observed sequence                 CCTGCAAGGTCGAATCCGGTC
        maximum likelihood sequence       CCTGCAAGGTCGAATCCGGTC
        maximum likelihood edit distance  0

evaluating sequence 92 of 100
        observed sequence                 CCTGCAAGGTCGAATCCGGT
        maximum likelihood sequence       CCTGCAAGGTCGAATCCGGT
        maximum likelihood edit distance  0

evaluating sequence 93 of 100
        observed sequence                 GGACCGGATTCGACCTTGCAG
        maximum likelihood sequence       GGACCGGATTCGACCTTGCAG
        maximum likelihood edit distance  0

evaluating sequence 94 of 100
        observed sequence                 GACCGGATTCGACCTTGCAG
        maximum likelihood sequence       GACCGGATTCGACCTTGCAG
        maximum likelihood edit distance  0

evaluating sequence 95 of 100
        observed sequence                 CTGCAAGGTCGAATCCGGTC
        maximum likelihood sequence       CTGCAAGGTCGAATCCGGTC
        maximum likelihood edit distance  0

evaluating sequence 96 of 100
        observed sequence                 CCTGCAAGGTCGAATCCGGTC
        maximum likelihood sequence       CCTGCAAGGTCGAATCCGGTC
        maximum likelihood edit distance  0

evaluating sequence 97 of 100
        observed sequence                 CCTGCAAGGTCGAATCCGGTC
        maximum likelihood sequence       CCTGCAAGGTCGAATCCGGTC
        maximum likelihood edit distance  0

evaluating sequence 98 of 100
        observed sequence                 CTGCAAGGTCGAATCCGGTC
        maximum likelihood sequence       CTGCAAGGTCGAATCCGGTC
        maximum likelihood edit distance  0
```

```
evaluating sequence 99 of 100
        observed sequence                  CTGCAAGGTCGAATCCGGT
        maximum likelihood sequence        CTGCAAGGTCGAATCCGGT
        maximum likelihood edit distance  0

evaluating sequence 100 of 100
        observed sequence                  GACCGGATTCGACCTTGCAG
        maximum likelihood sequence        GACCGGATTCGACCTTGCAG
        maximum likelihood edit distance  0

DATASET STATISTICS:
        assumed error rate    8.333333333333332%
        total bases observed  1993
        total edits accepted  0
        inferred error rate   0.0%
```

In [23]:

```
k = 13
canonical_kmers = collect(keys(Eisenia.count_canonical_kmers(maximum_likelihood_observations, k)))
stranded_kmer_graph = Eisenia.build_stranded_kmer_graph(canonical_kmers, maximum_likelihood_observations)
filename = reference_sequence_id * "." * replace(string(Dates.now()), ':' => '.') * ".svg"
Eisenia.plot_stranded_kmer_graph(stranded_kmer_graph, filename=filename)
HTML("""
<image src="$filename" width=50%>
""")
```

Out[23]:

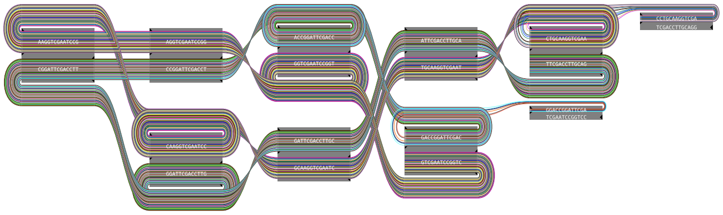

In [24]:

```
maximum_likelihood_observations = Eisenia.viterbi_maximum_likelihood_traversals(stranded_kmer_graph, verbosity="reads");
```

```
computing kmer counts...
computing kmer state likelihoods...
finding shortest paths between kmers...
finding viterbi maximum likelihood paths for observed sequences...

evaluating sequence 1 of 100
        observed sequence                 GACCGGATTCGACCTTGCAG
        maximum likelihood sequence       GACCGGATTCGACCTTGCAG
        maximum likelihood edit distance  0

evaluating sequence 2 of 100
        observed sequence                 CCTGCAAGGTCGAATCCGGTC
        maximum likelihood sequence       CCTGCAAGGTCGAATCCGGTC
        maximum likelihood edit distance  0

evaluating sequence 3 of 100
        observed sequence                 GACCGGATTCGACCTTGCAG
        maximum likelihood sequence       GACCGGATTCGACCTTGCAG
        maximum likelihood edit distance  0

evaluating sequence 4 of 100
        observed sequence                 GACCGGATTCGACCTTGCAG
        maximum likelihood sequence       GACCGGATTCGACCTTGCAG
        maximum likelihood edit distance  0

evaluating sequence 5 of 100
        observed sequence                 GACCGGATTCGACCTTGCAG
        maximum likelihood sequence       GACCGGATTCGACCTTGCAG
        maximum likelihood edit distance  0

evaluating sequence 6 of 100
        observed sequence                 GACCGGATTCGACCTTGCAG
        maximum likelihood sequence       GACCGGATTCGACCTTGCAG
        maximum likelihood edit distance  0

evaluating sequence 7 of 100
        observed sequence                 CTGCAAGGTCGAATCCGGTC
        maximum likelihood sequence       CTGCAAGGTCGAATCCGGTC
        maximum likelihood edit distance  0

evaluating sequence 8 of 100
        observed sequence                 CTGCAAGGTCGAATCCGGTC
        maximum likelihood sequence       CTGCAAGGTCGAATCCGGTC
        maximum likelihood edit distance  0

evaluating sequence 9 of 100
        observed sequence                 GACCGGATTCGACCTTGCAG
        maximum likelihood sequence       GACCGGATTCGACCTTGCAG
        maximum likelihood edit distance  0

evaluating sequence 10 of 100
        observed sequence                 CTGCAAGGTCGAATCCGGTC
        maximum likelihood sequence       CTGCAAGGTCGAATCCGGTC
        maximum likelihood edit distance  0

evaluating sequence 11 of 100
        observed sequence                 CCGGATTCGACCTTGCAG
        maximum likelihood sequence       CCGGATTCGACCTTGCAG
        maximum likelihood edit distance  0

evaluating sequence 12 of 100
        observed sequence                 CTGCAAGGTCGAATCCGGT
        maximum likelihood sequence       CTGCAAGGTCGAATCCGGT
        maximum likelihood edit distance  0

evaluating sequence 13 of 100
        observed sequence                 GACCGGATTCGACCTTGCAG
        maximum likelihood sequence       GACCGGATTCGACCTTGCAG
        maximum likelihood edit distance  0

evaluating sequence 14 of 100
        observed sequence                 TGCAAGGTCGAATCCGGTC
        maximum likelihood sequence       TGCAAGGTCGAATCCGGTC
        maximum likelihood edit distance  0

evaluating sequence 15 of 100
        observed sequence                 GACCGGATTCGACCTTGCAG
        maximum likelihood sequence       GACCGGATTCGACCTTGCAG
        maximum likelihood edit distance  0

evaluating sequence 16 of 100
        observed sequence                 ACCGGATTCGACCTTGCA
        maximum likelihood sequence       ACCGGATTCGACCTTGCA
        maximum likelihood edit distance  0

evaluating sequence 17 of 100
        observed sequence                 CCTGCAAGGTCGAATCCGGTC
        maximum likelihood sequence       CCTGCAAGGTCGAATCCGGTC
        maximum likelihood edit distance  0

evaluating sequence 18 of 100
        observed sequence                 CTGCAAGGTCGAATCCGGTC
        maximum likelihood sequence       CTGCAAGGTCGAATCCGGTC
        maximum likelihood edit distance  0

evaluating sequence 19 of 100
        observed sequence                 CTGCAAGGTCGAATCCGGTC
        maximum likelihood sequence       CTGCAAGGTCGAATCCGGTC
        maximum likelihood edit distance  0
```

```
evaluating sequence 20 of 100
        observed sequence                 GACCGGATTCGACCTTGCAG
        maximum likelihood sequence       GACCGGATTCGACCTTGCAG
        maximum likelihood edit distance  0

evaluating sequence 21 of 100
        observed sequence                 GACCGGATTCGACCTTGCAG
        maximum likelihood sequence       GACCGGATTCGACCTTGCAG
        maximum likelihood edit distance  0

evaluating sequence 22 of 100
        observed sequence                 GACCGGATTCGACCTTGCA
        maximum likelihood sequence       GACCGGATTCGACCTTGCA
        maximum likelihood edit distance  0

evaluating sequence 23 of 100
        observed sequence                 CTGCAAGGTCGAATCCGGTC
        maximum likelihood sequence       CTGCAAGGTCGAATCCGGTC
        maximum likelihood edit distance  0

evaluating sequence 24 of 100
        observed sequence                 GACCGGATTCGACCTTGCAG
        maximum likelihood sequence       GACCGGATTCGACCTTGCAG
        maximum likelihood edit distance  0

evaluating sequence 25 of 100
        observed sequence                 CCGGATTCGACCTTGCAG
        maximum likelihood sequence       CCGGATTCGACCTTGCAG
        maximum likelihood edit distance  0

evaluating sequence 26 of 100
        observed sequence                 CAAGGTCGAATCCGGTC
        maximum likelihood sequence       CAAGGTCGAATCCGGTC
        maximum likelihood edit distance  0

evaluating sequence 27 of 100
        observed sequence                 GACCGGATTCGACCTTGCAG
        maximum likelihood sequence       GACCGGATTCGACCTTGCAG
        maximum likelihood edit distance  0

evaluating sequence 28 of 100
        observed sequence                 CTGCAAGGTCGAATCCGGTC
        maximum likelihood sequence       CTGCAAGGTCGAATCCGGTC
        maximum likelihood edit distance  0

evaluating sequence 29 of 100
        observed sequence                 GACCGGATTCGACCTTGCAG
        maximum likelihood sequence       GACCGGATTCGACCTTGCAG
        maximum likelihood edit distance  0

evaluating sequence 30 of 100
        observed sequence                 GACCGGATTCGACCTTGCAG
        maximum likelihood sequence       GACCGGATTCGACCTTGCAG
        maximum likelihood edit distance  0

evaluating sequence 31 of 100
        observed sequence                 CTGCAAGGTCGAATCCGGTC
        maximum likelihood sequence       CTGCAAGGTCGAATCCGGTC
        maximum likelihood edit distance  0

evaluating sequence 32 of 100
        observed sequence                 CTGCAAGGTCGAATCCGGTC
        maximum likelihood sequence       CTGCAAGGTCGAATCCGGTC
        maximum likelihood edit distance  0

evaluating sequence 33 of 100
        observed sequence                 CTGCAAGGTCGAATCCGGTC
        maximum likelihood sequence       CTGCAAGGTCGAATCCGGTC
        maximum likelihood edit distance  0

evaluating sequence 34 of 100
        observed sequence                 GACCGGATTCGACCTTGCAG
        maximum likelihood sequence       GACCGGATTCGACCTTGCAG
        maximum likelihood edit distance  0

evaluating sequence 35 of 100
        observed sequence                 GACCGGATTCGACCTTGCAG
        maximum likelihood sequence       GACCGGATTCGACCTTGCAG
        maximum likelihood edit distance  0

evaluating sequence 36 of 100
        observed sequence                 CTGCAAGGTCGAATCCGGTC
        maximum likelihood sequence       CTGCAAGGTCGAATCCGGTC
        maximum likelihood edit distance  0

evaluating sequence 37 of 100
        observed sequence                 CCTGCAAGGTCGAATCCGGTC
        maximum likelihood sequence       CCTGCAAGGTCGAATCCGGTC
        maximum likelihood edit distance  0

evaluating sequence 38 of 100
        observed sequence                 GACCGGATTCGACCTTGCAG
        maximum likelihood sequence       GACCGGATTCGACCTTGCAG
        maximum likelihood edit distance  0

evaluating sequence 39 of 100
        observed sequence                 CCTGCAAGGTCGAATCCGGTC
        maximum likelihood sequence       CCTGCAAGGTCGAATCCGGTC
```

```
        maximum likelihood edit distance  0

evaluating sequence 40 of 100
        observed sequence                 CTGCAAGGTCGAATCCGGTC
        maximum likelihood sequence       CTGCAAGGTCGAATCCGGTC
        maximum likelihood edit distance  0

evaluating sequence 41 of 100
        observed sequence                 GGACCGGATTCGACCTTGCA
        maximum likelihood sequence       GGACCGGATTCGACCTTGCA
        maximum likelihood edit distance  0

evaluating sequence 42 of 100
        observed sequence                 CTGCAAGGTCGAATCCGGTC
        maximum likelihood sequence       CTGCAAGGTCGAATCCGGTC
        maximum likelihood edit distance  0

evaluating sequence 43 of 100
        observed sequence                 GACCGGATTCGACCTTGCAG
        maximum likelihood sequence       GACCGGATTCGACCTTGCAG
        maximum likelihood edit distance  0

evaluating sequence 44 of 100
        observed sequence                 CCTGCAAGGTCGAATCCGGTC
        maximum likelihood sequence       CCTGCAAGGTCGAATCCGGTC
        maximum likelihood edit distance  0

evaluating sequence 45 of 100
        observed sequence                 CTGCAAGGTCGAATCCGGTC
        maximum likelihood sequence       CTGCAAGGTCGAATCCGGTC
        maximum likelihood edit distance  0

evaluating sequence 46 of 100
        observed sequence                 GGACCGGATTCGACCTTGCAG
        maximum likelihood sequence       GGACCGGATTCGACCTTGCAG
        maximum likelihood edit distance  0

evaluating sequence 47 of 100
        observed sequence                 GACCGGATTCGACCTTGCAG
        maximum likelihood sequence       GACCGGATTCGACCTTGCAG
        maximum likelihood edit distance  0

evaluating sequence 48 of 100
        observed sequence                 ACCGGATTCGACCTTGCAG
        maximum likelihood sequence       ACCGGATTCGACCTTGCAG
        maximum likelihood edit distance  0

evaluating sequence 49 of 100
        observed sequence                 CTGCAAGGTCGAATCCGGTC
        maximum likelihood sequence       CTGCAAGGTCGAATCCGGTC
        maximum likelihood edit distance  0

evaluating sequence 50 of 100
        observed sequence                 GACCGGATTCGACCTTGCAG
        maximum likelihood sequence       GACCGGATTCGACCTTGCAG
        maximum likelihood edit distance  0

evaluating sequence 51 of 100
        observed sequence                 CTGCAAGGTCGAATCCGGTC
        maximum likelihood sequence       CTGCAAGGTCGAATCCGGTC
        maximum likelihood edit distance  0

evaluating sequence 52 of 100
        observed sequence                 TGCAAGGTCGAATCCGGTC
        maximum likelihood sequence       TGCAAGGTCGAATCCGGTC
        maximum likelihood edit distance  0

evaluating sequence 53 of 100
        observed sequence                 CCGGATTCGACCTTGCAG
        maximum likelihood sequence       CCGGATTCGACCTTGCAG
        maximum likelihood edit distance  0

evaluating sequence 54 of 100
        observed sequence                 GACCGGATTCGACCTTGCAG
        maximum likelihood sequence       GACCGGATTCGACCTTGCAG
        maximum likelihood edit distance  0

evaluating sequence 55 of 100
        observed sequence                 GACCGGATTCGACCTTGCAG
        maximum likelihood sequence       GACCGGATTCGACCTTGCAG
        maximum likelihood edit distance  0

evaluating sequence 56 of 100
        observed sequence                 CCTGCAAGGTCGAATCCGGTC
        maximum likelihood sequence       CCTGCAAGGTCGAATCCGGTC
        maximum likelihood edit distance  0

evaluating sequence 57 of 100
        observed sequence                 ACCGGATTCGACCTTGCAG
        maximum likelihood sequence       ACCGGATTCGACCTTGCAG
        maximum likelihood edit distance  0

evaluating sequence 58 of 100
        observed sequence                 CTGCAAGGTCGAATCCGGTC
        maximum likelihood sequence       CTGCAAGGTCGAATCCGGTC
        maximum likelihood edit distance  0

evaluating sequence 59 of 100
        observed sequence                 GACCGGATTCGACCTTGCAG
```

```
        maximum likelihood sequence       GACCGGATTCGACCTTGCAG
        maximum likelihood edit distance  0

evaluating sequence 60 of 100
        observed sequence                 CCGGATTCGACCTTGCAG
        maximum likelihood sequence       CCGGATTCGACCTTGCAG
        maximum likelihood edit distance  0

evaluating sequence 61 of 100
        observed sequence                 ACCGGATTCGACCTTGCAG
        maximum likelihood sequence       ACCGGATTCGACCTTGCAG
        maximum likelihood edit distance  0

evaluating sequence 62 of 100
        observed sequence                 CTGCAAGGTCGAATCCGGTC
        maximum likelihood sequence       CTGCAAGGTCGAATCCGGTC
        maximum likelihood edit distance  0

evaluating sequence 63 of 100
        observed sequence                 CTGCAAGGTCGAATCCGGTC
        maximum likelihood sequence       CTGCAAGGTCGAATCCGGTC
        maximum likelihood edit distance  0

evaluating sequence 64 of 100
        observed sequence                 GACCGGATTCGACCTTGCAG
        maximum likelihood sequence       GACCGGATTCGACCTTGCAG
        maximum likelihood edit distance  0

evaluating sequence 65 of 100
        observed sequence                 TGCAAGGTCGAATCCGGTC
        maximum likelihood sequence       TGCAAGGTCGAATCCGGTC
        maximum likelihood edit distance  0

evaluating sequence 66 of 100
        observed sequence                 CTGCAAGGTCGAATCCGGTC
        maximum likelihood sequence       CTGCAAGGTCGAATCCGGTC
        maximum likelihood edit distance  0

evaluating sequence 67 of 100
        observed sequence                 CTGCAAGGTCGAATCCGGTC
        maximum likelihood sequence       CTGCAAGGTCGAATCCGGTC
        maximum likelihood edit distance  0

evaluating sequence 68 of 100
        observed sequence                 GGACCGGATTCGACCTTGCA
        maximum likelihood sequence       GGACCGGATTCGACCTTGCA
        maximum likelihood edit distance  0

evaluating sequence 69 of 100
        observed sequence                 GCAAGGTCGAATCCGGTC
        maximum likelihood sequence       GCAAGGTCGAATCCGGTC
        maximum likelihood edit distance  0

evaluating sequence 70 of 100
        observed sequence                 CTGCAAGGTCGAATCCGGTC
        maximum likelihood sequence       CTGCAAGGTCGAATCCGGTC
        maximum likelihood edit distance  0

evaluating sequence 71 of 100
        observed sequence                 CCTGCAAGGTCGAATCCGGTC
        maximum likelihood sequence       CCTGCAAGGTCGAATCCGGTC
        maximum likelihood edit distance  0

evaluating sequence 72 of 100
        observed sequence                 GACCGGATTCGACCTTGCAG
        maximum likelihood sequence       GACCGGATTCGACCTTGCAG
        maximum likelihood edit distance  0

evaluating sequence 73 of 100
        observed sequence                 GACCGGATTCGACCTTGCAG
        maximum likelihood sequence       GACCGGATTCGACCTTGCAG
        maximum likelihood edit distance  0

evaluating sequence 74 of 100
        observed sequence                 CCTGCAAGGTCGAATCCGGTC
        maximum likelihood sequence       CCTGCAAGGTCGAATCCGGTC
        maximum likelihood edit distance  0

evaluating sequence 75 of 100
        observed sequence                 GACCGGATTCGACCTTGCAG
        maximum likelihood sequence       GACCGGATTCGACCTTGCAG
        maximum likelihood edit distance  0

evaluating sequence 76 of 100
        observed sequence                 GGACCGGATTCGACCTTGCAG
        maximum likelihood sequence       GGACCGGATTCGACCTTGCAG
        maximum likelihood edit distance  0

evaluating sequence 77 of 100
        observed sequence                 CTGCAAGGTCGAATCCGGTC
        maximum likelihood sequence       CTGCAAGGTCGAATCCGGTC
        maximum likelihood edit distance  0

evaluating sequence 78 of 100
        observed sequence                 GGACCGGATTCGACCTTGCAG
        maximum likelihood sequence       GGACCGGATTCGACCTTGCAG
        maximum likelihood edit distance  0

evaluating sequence 79 of 100
```

```
        observed sequence                 CTGCAAGGTCGAATCCGGTC
        maximum likelihood sequence       CTGCAAGGTCGAATCCGGTC
        maximum likelihood edit distance  0

evaluating sequence 80 of 100
        observed sequence                 CTGCAAGGTCGAATCCGGTC
        maximum likelihood sequence       CTGCAAGGTCGAATCCGGTC
        maximum likelihood edit distance  0

evaluating sequence 81 of 100
        observed sequence                 CCTGCAAGGTCGAATCCGGTC
        maximum likelihood sequence       CCTGCAAGGTCGAATCCGGTC
        maximum likelihood edit distance  0

evaluating sequence 82 of 100
        observed sequence                 GGACCGGATTCGACCTTGCAG
        maximum likelihood sequence       GGACCGGATTCGACCTTGCAG
        maximum likelihood edit distance  0

evaluating sequence 83 of 100
        observed sequence                 GACCGGATTCGACCTTGCAG
        maximum likelihood sequence       GACCGGATTCGACCTTGCAG
        maximum likelihood edit distance  0

evaluating sequence 84 of 100
        observed sequence                 CTGCAAGGTCGAATCCGGTC
        maximum likelihood sequence       CTGCAAGGTCGAATCCGGTC
        maximum likelihood edit distance  0

evaluating sequence 85 of 100
        observed sequence                 CCTGCAAGGTCGAATCCGGTC
        maximum likelihood sequence       CCTGCAAGGTCGAATCCGGTC
        maximum likelihood edit distance  0

evaluating sequence 86 of 100
        observed sequence                 CCTGCAAGGTCGAATCCGGTC
        maximum likelihood sequence       CCTGCAAGGTCGAATCCGGTC
        maximum likelihood edit distance  0

evaluating sequence 87 of 100
        observed sequence                 GACCGGATTCGACCTTGCAG
        maximum likelihood sequence       GACCGGATTCGACCTTGCAG
        maximum likelihood edit distance  0

evaluating sequence 88 of 100
        observed sequence                 CCGGATTCGACCTTGCAG
        maximum likelihood sequence       CCGGATTCGACCTTGCAG
        maximum likelihood edit distance  0

evaluating sequence 89 of 100
        observed sequence                 CTGCAAGGTCGAATCCGGTC
        maximum likelihood sequence       CTGCAAGGTCGAATCCGGTC
        maximum likelihood edit distance  0

evaluating sequence 90 of 100
        observed sequence                 GACCGGATTCGACCTTGCAG
        maximum likelihood sequence       GACCGGATTCGACCTTGCAG
        maximum likelihood edit distance  0

evaluating sequence 91 of 100
        observed sequence                 CCTGCAAGGTCGAATCCGGTC
        maximum likelihood sequence       CCTGCAAGGTCGAATCCGGTC
        maximum likelihood edit distance  0

evaluating sequence 92 of 100
        observed sequence                 CCTGCAAGGTCGAATCCGGT
        maximum likelihood sequence       CCTGCAAGGTCGAATCCGGT
        maximum likelihood edit distance  0

evaluating sequence 93 of 100
        observed sequence                 GGACCGGATTCGACCTTGCAG
        maximum likelihood sequence       GGACCGGATTCGACCTTGCAG
        maximum likelihood edit distance  0

evaluating sequence 94 of 100
        observed sequence                 GACCGGATTCGACCTTGCAG
        maximum likelihood sequence       GACCGGATTCGACCTTGCAG
        maximum likelihood edit distance  0

evaluating sequence 95 of 100
        observed sequence                 CTGCAAGGTCGAATCCGGTC
        maximum likelihood sequence       CTGCAAGGTCGAATCCGGTC
        maximum likelihood edit distance  0

evaluating sequence 96 of 100
        observed sequence                 CCTGCAAGGTCGAATCCGGTC
        maximum likelihood sequence       CCTGCAAGGTCGAATCCGGTC
        maximum likelihood edit distance  0

evaluating sequence 97 of 100
        observed sequence                 CCTGCAAGGTCGAATCCGGTC
        maximum likelihood sequence       CCTGCAAGGTCGAATCCGGTC
        maximum likelihood edit distance  0

evaluating sequence 98 of 100
        observed sequence                 CTGCAAGGTCGAATCCGGTC
        maximum likelihood sequence       CTGCAAGGTCGAATCCGGTC
        maximum likelihood edit distance  0
```

```
evaluating sequence 99 of 100
        observed sequence                 CTGCAAGGTCGAATCCGGT
        maximum likelihood sequence       CTGCAAGGTCGAATCCGGT
        maximum likelihood edit distance  0

evaluating sequence 100 of 100
        observed sequence                 GACCGGATTCGACCTTGCAG
        maximum likelihood sequence       GACCGGATTCGACCTTGCAG
        maximum likelihood edit distance  0

DATASET STATISTICS:
        assumed error rate    7.142857142857142%
        total bases observed  1993
        total edits accepted  0
        inferred error rate   0.0%
```

In [25]:
```julia
k = 17
canonical_kmers = collect(keys(Eisenia.count_canonical_kmers(maximum_likelihood_observations, k)))
stranded_kmer_graph = Eisenia.build_stranded_kmer_graph(canonical_kmers, maximum_likelihood_observations)
filename = reference_sequence_id * "." * replace(string(Dates.now()), ':' => '.') * ".svg"
Eisenia.plot_stranded_kmer_graph(stranded_kmer_graph, filename=filename)
HTML("""
<image src="$filename" width=50%>
""")
```

Out[25]:

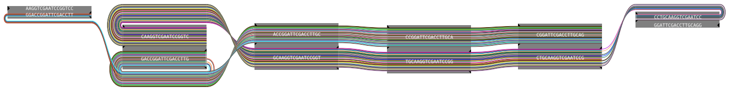

In [26]:
```julia
maximum_likelihood_observations = Eisenia.viterbi_maximum_likelihood_traversals(stranded_kmer_graph, verbosity="reads");
```

```
computing kmer counts...
computing kmer state likelihoods...
finding shortest paths between kmers...
finding viterbi maximum likelihood paths for observed sequences...

evaluating sequence 1 of 100
        observed sequence                 GACCGGATTCGACCTTGCAG
        maximum likelihood sequence       GACCGGATTCGACCTTGCAG
        maximum likelihood edit distance  0

evaluating sequence 2 of 100
        observed sequence                 CCTGCAAGGTCGAATCCGGTC
        maximum likelihood sequence       CCTGCAAGGTCGAATCCGGTC
        maximum likelihood edit distance  0

evaluating sequence 3 of 100
        observed sequence                 GACCGGATTCGACCTTGCAG
        maximum likelihood sequence       GACCGGATTCGACCTTGCAG
        maximum likelihood edit distance  0

evaluating sequence 4 of 100
        observed sequence                 GACCGGATTCGACCTTGCAG
        maximum likelihood sequence       GACCGGATTCGACCTTGCAG
        maximum likelihood edit distance  0

evaluating sequence 5 of 100
        observed sequence                 GACCGGATTCGACCTTGCAG
        maximum likelihood sequence       GACCGGATTCGACCTTGCAG
        maximum likelihood edit distance  0

evaluating sequence 6 of 100
        observed sequence                 GACCGGATTCGACCTTGCAG
        maximum likelihood sequence       GACCGGATTCGACCTTGCAG
        maximum likelihood edit distance  0

evaluating sequence 7 of 100
        observed sequence                 CTGCAAGGTCGAATCCGGTC
        maximum likelihood sequence       CTGCAAGGTCGAATCCGGTC
        maximum likelihood edit distance  0

evaluating sequence 8 of 100
        observed sequence                 CTGCAAGGTCGAATCCGGTC
        maximum likelihood sequence       CTGCAAGGTCGAATCCGGTC
        maximum likelihood edit distance  0

evaluating sequence 9 of 100
        observed sequence                 GACCGGATTCGACCTTGCAG
        maximum likelihood sequence       GACCGGATTCGACCTTGCAG
        maximum likelihood edit distance  0

evaluating sequence 10 of 100
        observed sequence                 CTGCAAGGTCGAATCCGGTC
        maximum likelihood sequence       CTGCAAGGTCGAATCCGGTC
        maximum likelihood edit distance  0

evaluating sequence 11 of 100
        observed sequence                 CCGGATTCGACCTTGCAG
        maximum likelihood sequence       CCGGATTCGACCTTGCAG
        maximum likelihood edit distance  0

evaluating sequence 12 of 100
        observed sequence                 CTGCAAGGTCGAATCCGGT
        maximum likelihood sequence       CTGCAAGGTCGAATCCGGT
        maximum likelihood edit distance  0

evaluating sequence 13 of 100
        observed sequence                 GACCGGATTCGACCTTGCAG
        maximum likelihood sequence       GACCGGATTCGACCTTGCAG
        maximum likelihood edit distance  0

evaluating sequence 14 of 100
        observed sequence                 TGCAAGGTCGAATCCGGTC
        maximum likelihood sequence       TGCAAGGTCGAATCCGGTC
        maximum likelihood edit distance  0

evaluating sequence 15 of 100
        observed sequence                 GACCGGATTCGACCTTGCAG
        maximum likelihood sequence       GACCGGATTCGACCTTGCAG
        maximum likelihood edit distance  0

evaluating sequence 16 of 100
        observed sequence                 ACCGGATTCGACCTTGCA
        maximum likelihood sequence       ACCGGATTCGACCTTGCA
        maximum likelihood edit distance  0

evaluating sequence 17 of 100
        observed sequence                 CCTGCAAGGTCGAATCCGGTC
        maximum likelihood sequence       CCTGCAAGGTCGAATCCGGTC
        maximum likelihood edit distance  0

evaluating sequence 18 of 100
        observed sequence                 CTGCAAGGTCGAATCCGGTC
        maximum likelihood sequence       CTGCAAGGTCGAATCCGGTC
        maximum likelihood edit distance  0

evaluating sequence 19 of 100
        observed sequence                 CTGCAAGGTCGAATCCGGTC
        maximum likelihood sequence       CTGCAAGGTCGAATCCGGTC
        maximum likelihood edit distance  0
```

```
evaluating sequence 20 of 100
        observed sequence                 GACCGGATTCGACCTTGCAG
        maximum likelihood sequence       GACCGGATTCGACCTTGCAG
        maximum likelihood edit distance  0

evaluating sequence 21 of 100
        observed sequence                 GACCGGATTCGACCTTGCAG
        maximum likelihood sequence       GACCGGATTCGACCTTGCAG
        maximum likelihood edit distance  0

evaluating sequence 22 of 100
        observed sequence                 GACCGGATTCGACCTTGCA
        maximum likelihood sequence       GACCGGATTCGACCTTGCA
        maximum likelihood edit distance  0

evaluating sequence 23 of 100
        observed sequence                 CTGCAAGGTCGAATCCGGTC
        maximum likelihood sequence       CTGCAAGGTCGAATCCGGTC
        maximum likelihood edit distance  0

evaluating sequence 24 of 100
        observed sequence                 GACCGGATTCGACCTTGCAG
        maximum likelihood sequence       GACCGGATTCGACCTTGCAG
        maximum likelihood edit distance  0

evaluating sequence 25 of 100
        observed sequence                 CCGGATTCGACCTTGCAG
        maximum likelihood sequence       CCGGATTCGACCTTGCAG
        maximum likelihood edit distance  0

evaluating sequence 26 of 100
        observed sequence                 CAAGGTCGAATCCGGTC
        maximum likelihood sequence       CAAGGTCGAATCCGGTC
        maximum likelihood edit distance  0

evaluating sequence 27 of 100
        observed sequence                 GACCGGATTCGACCTTGCAG
        maximum likelihood sequence       GACCGGATTCGACCTTGCAG
        maximum likelihood edit distance  0

evaluating sequence 28 of 100
        observed sequence                 CTGCAAGGTCGAATCCGGTC
        maximum likelihood sequence       CTGCAAGGTCGAATCCGGTC
        maximum likelihood edit distance  0

evaluating sequence 29 of 100
        observed sequence                 GACCGGATTCGACCTTGCAG
        maximum likelihood sequence       GACCGGATTCGACCTTGCAG
        maximum likelihood edit distance  0

evaluating sequence 30 of 100
        observed sequence                 GACCGGATTCGACCTTGCAG
        maximum likelihood sequence       GACCGGATTCGACCTTGCAG
        maximum likelihood edit distance  0

evaluating sequence 31 of 100
        observed sequence                 CTGCAAGGTCGAATCCGGTC
        maximum likelihood sequence       CTGCAAGGTCGAATCCGGTC
        maximum likelihood edit distance  0

evaluating sequence 32 of 100
        observed sequence                 CTGCAAGGTCGAATCCGGTC
        maximum likelihood sequence       CTGCAAGGTCGAATCCGGTC
        maximum likelihood edit distance  0

evaluating sequence 33 of 100
        observed sequence                 CTGCAAGGTCGAATCCGGTC
        maximum likelihood sequence       CTGCAAGGTCGAATCCGGTC
        maximum likelihood edit distance  0

evaluating sequence 34 of 100
        observed sequence                 GACCGGATTCGACCTTGCAG
        maximum likelihood sequence       GACCGGATTCGACCTTGCAG
        maximum likelihood edit distance  0

evaluating sequence 35 of 100
        observed sequence                 GACCGGATTCGACCTTGCAG
        maximum likelihood sequence       GACCGGATTCGACCTTGCAG
        maximum likelihood edit distance  0

evaluating sequence 36 of 100
        observed sequence                 CTGCAAGGTCGAATCCGGTC
        maximum likelihood sequence       CTGCAAGGTCGAATCCGGTC
        maximum likelihood edit distance  0

evaluating sequence 37 of 100
        observed sequence                 CCTGCAAGGTCGAATCCGGTC
        maximum likelihood sequence       CCTGCAAGGTCGAATCCGGTC
        maximum likelihood edit distance  0

evaluating sequence 38 of 100
        observed sequence                 GACCGGATTCGACCTTGCAG
        maximum likelihood sequence       GACCGGATTCGACCTTGCAG
        maximum likelihood edit distance  0

evaluating sequence 39 of 100
        observed sequence                 CCTGCAAGGTCGAATCCGGTC
        maximum likelihood sequence       CCTGCAAGGTCGAATCCGGTC
```

```
        maximum likelihood edit distance  0

evaluating sequence 40 of 100
        observed sequence                 CTGCAAGGTCGAATCCGGTC
        maximum likelihood sequence       CTGCAAGGTCGAATCCGGTC
        maximum likelihood edit distance  0

evaluating sequence 41 of 100
        observed sequence                 GGACCGGATTCGACCTTGCA
        maximum likelihood sequence       GGACCGGATTCGACCTTGCA
        maximum likelihood edit distance  0

evaluating sequence 42 of 100
        observed sequence                 CTGCAAGGTCGAATCCGGTC
        maximum likelihood sequence       CTGCAAGGTCGAATCCGGTC
        maximum likelihood edit distance  0

evaluating sequence 43 of 100
        observed sequence                 GACCGGATTCGACCTTGCAG
        maximum likelihood sequence       GACCGGATTCGACCTTGCAG
        maximum likelihood edit distance  0

evaluating sequence 44 of 100
        observed sequence                 CCTGCAAGGTCGAATCCGGTC
        maximum likelihood sequence       CCTGCAAGGTCGAATCCGGTC
        maximum likelihood edit distance  0

evaluating sequence 45 of 100
        observed sequence                 CTGCAAGGTCGAATCCGGTC
        maximum likelihood sequence       CTGCAAGGTCGAATCCGGTC
        maximum likelihood edit distance  0

evaluating sequence 46 of 100
        observed sequence                 GGACCGGATTCGACCTTGCAG
        maximum likelihood sequence       GGACCGGATTCGACCTTGCAG
        maximum likelihood edit distance  0

evaluating sequence 47 of 100
        observed sequence                 GACCGGATTCGACCTTGCAG
        maximum likelihood sequence       GACCGGATTCGACCTTGCAG
        maximum likelihood edit distance  0

evaluating sequence 48 of 100
        observed sequence                 ACCGGATTCGACCTTGCAG
        maximum likelihood sequence       ACCGGATTCGACCTTGCAG
        maximum likelihood edit distance  0

evaluating sequence 49 of 100
        observed sequence                 CTGCAAGGTCGAATCCGGTC
        maximum likelihood sequence       CTGCAAGGTCGAATCCGGTC
        maximum likelihood edit distance  0

evaluating sequence 50 of 100
        observed sequence                 GACCGGATTCGACCTTGCAG
        maximum likelihood sequence       GACCGGATTCGACCTTGCAG
        maximum likelihood edit distance  0

evaluating sequence 51 of 100
        observed sequence                 CTGCAAGGTCGAATCCGGTC
        maximum likelihood sequence       CTGCAAGGTCGAATCCGGTC
        maximum likelihood edit distance  0

evaluating sequence 52 of 100
        observed sequence                 TGCAAGGTCGAATCCGGTC
        maximum likelihood sequence       TGCAAGGTCGAATCCGGTC
        maximum likelihood edit distance  0

evaluating sequence 53 of 100
        observed sequence                 CCGGATTCGACCTTGCAG
        maximum likelihood sequence       CCGGATTCGACCTTGCAG
        maximum likelihood edit distance  0

evaluating sequence 54 of 100
        observed sequence                 GACCGGATTCGACCTTGCAG
        maximum likelihood sequence       GACCGGATTCGACCTTGCAG
        maximum likelihood edit distance  0

evaluating sequence 55 of 100
        observed sequence                 GACCGGATTCGACCTTGCAG
        maximum likelihood sequence       GACCGGATTCGACCTTGCAG
        maximum likelihood edit distance  0

evaluating sequence 56 of 100
        observed sequence                 CCTGCAAGGTCGAATCCGGTC
        maximum likelihood sequence       CCTGCAAGGTCGAATCCGGTC
        maximum likelihood edit distance  0

evaluating sequence 57 of 100
        observed sequence                 ACCGGATTCGACCTTGCAG
        maximum likelihood sequence       ACCGGATTCGACCTTGCAG
        maximum likelihood edit distance  0

evaluating sequence 58 of 100
        observed sequence                 CTGCAAGGTCGAATCCGGTC
        maximum likelihood sequence       CTGCAAGGTCGAATCCGGTC
        maximum likelihood edit distance  0

evaluating sequence 59 of 100
        observed sequence                 GACCGGATTCGACCTTGCAG
```

```
        maximum likelihood sequence       GACCGGATTCGACCTTGCAG
        maximum likelihood edit distance  0

evaluating sequence 60 of 100
        observed sequence                 CCGGATTCGACCTTGCAG
        maximum likelihood sequence       CCGGATTCGACCTTGCAG
        maximum likelihood edit distance  0

evaluating sequence 61 of 100
        observed sequence                 ACCGGATTCGACCTTGCAG
        maximum likelihood sequence       ACCGGATTCGACCTTGCAG
        maximum likelihood edit distance  0

evaluating sequence 62 of 100
        observed sequence                 CTGCAAGGTCGAATCCGGTC
        maximum likelihood sequence       CTGCAAGGTCGAATCCGGTC
        maximum likelihood edit distance  0

evaluating sequence 63 of 100
        observed sequence                 CTGCAAGGTCGAATCCGGTC
        maximum likelihood sequence       CTGCAAGGTCGAATCCGGTC
        maximum likelihood edit distance  0

evaluating sequence 64 of 100
        observed sequence                 GACCGGATTCGACCTTGCAG
        maximum likelihood sequence       GACCGGATTCGACCTTGCAG
        maximum likelihood edit distance  0

evaluating sequence 65 of 100
        observed sequence                 TGCAAGGTCGAATCCGGTC
        maximum likelihood sequence       TGCAAGGTCGAATCCGGTC
        maximum likelihood edit distance  0

evaluating sequence 66 of 100
        observed sequence                 CTGCAAGGTCGAATCCGGTC
        maximum likelihood sequence       CTGCAAGGTCGAATCCGGTC
        maximum likelihood edit distance  0

evaluating sequence 67 of 100
        observed sequence                 CTGCAAGGTCGAATCCGGTC
        maximum likelihood sequence       CTGCAAGGTCGAATCCGGTC
        maximum likelihood edit distance  0

evaluating sequence 68 of 100
        observed sequence                 GGACCGGATTCGACCTTGCA
        maximum likelihood sequence       GGACCGGATTCGACCTTGCA
        maximum likelihood edit distance  0

evaluating sequence 69 of 100
        observed sequence                 GCAAGGTCGAATCCGGTC
        maximum likelihood sequence       GCAAGGTCGAATCCGGTC
        maximum likelihood edit distance  0

evaluating sequence 70 of 100
        observed sequence                 CTGCAAGGTCGAATCCGGTC
        maximum likelihood sequence       CTGCAAGGTCGAATCCGGTC
        maximum likelihood edit distance  0

evaluating sequence 71 of 100
        observed sequence                 CCTGCAAGGTCGAATCCGGTC
        maximum likelihood sequence       CCTGCAAGGTCGAATCCGGTC
        maximum likelihood edit distance  0

evaluating sequence 72 of 100
        observed sequence                 GACCGGATTCGACCTTGCAG
        maximum likelihood sequence       GACCGGATTCGACCTTGCAG
        maximum likelihood edit distance  0

evaluating sequence 73 of 100
        observed sequence                 GACCGGATTCGACCTTGCAG
        maximum likelihood sequence       GACCGGATTCGACCTTGCAG
        maximum likelihood edit distance  0

evaluating sequence 74 of 100
        observed sequence                 CCTGCAAGGTCGAATCCGGTC
        maximum likelihood sequence       CCTGCAAGGTCGAATCCGGTC
        maximum likelihood edit distance  0

evaluating sequence 75 of 100
        observed sequence                 GACCGGATTCGACCTTGCAG
        maximum likelihood sequence       GACCGGATTCGACCTTGCAG
        maximum likelihood edit distance  0

evaluating sequence 76 of 100
        observed sequence                 GGACCGGATTCGACCTTGCAG
        maximum likelihood sequence       GGACCGGATTCGACCTTGCAG
        maximum likelihood edit distance  0

evaluating sequence 77 of 100
        observed sequence                 CTGCAAGGTCGAATCCGGTC
        maximum likelihood sequence       CTGCAAGGTCGAATCCGGTC
        maximum likelihood edit distance  0

evaluating sequence 78 of 100
        observed sequence                 GGACCGGATTCGACCTTGCAG
        maximum likelihood sequence       GGACCGGATTCGACCTTGCAG
        maximum likelihood edit distance  0

evaluating sequence 79 of 100
```

```
        observed sequence                 CTGCAAGGTCGAATCCGGTC
        maximum likelihood sequence       CTGCAAGGTCGAATCCGGTC
        maximum likelihood edit distance  0

evaluating sequence 80 of 100
        observed sequence                 CTGCAAGGTCGAATCCGGTC
        maximum likelihood sequence       CTGCAAGGTCGAATCCGGTC
        maximum likelihood edit distance  0

evaluating sequence 81 of 100
        observed sequence                 CCTGCAAGGTCGAATCCGGTC
        maximum likelihood sequence       CCTGCAAGGTCGAATCCGGTC
        maximum likelihood edit distance  0

evaluating sequence 82 of 100
        observed sequence                 GGACCGGATTCGACCTTGCAG
        maximum likelihood sequence       GGACCGGATTCGACCTTGCAG
        maximum likelihood edit distance  0

evaluating sequence 83 of 100
        observed sequence                 GACCGGATTCGACCTTGCAG
        maximum likelihood sequence       GACCGGATTCGACCTTGCAG
        maximum likelihood edit distance  0

evaluating sequence 84 of 100
        observed sequence                 CTGCAAGGTCGAATCCGGTC
        maximum likelihood sequence       CTGCAAGGTCGAATCCGGTC
        maximum likelihood edit distance  0

evaluating sequence 85 of 100
        observed sequence                 CCTGCAAGGTCGAATCCGGTC
        maximum likelihood sequence       CCTGCAAGGTCGAATCCGGTC
        maximum likelihood edit distance  0

evaluating sequence 86 of 100
        observed sequence                 CCTGCAAGGTCGAATCCGGTC
        maximum likelihood sequence       CCTGCAAGGTCGAATCCGGTC
        maximum likelihood edit distance  0

evaluating sequence 87 of 100
        observed sequence                 GACCGGATTCGACCTTGCAG
        maximum likelihood sequence       GACCGGATTCGACCTTGCAG
        maximum likelihood edit distance  0

evaluating sequence 88 of 100
        observed sequence                 CCGGATTCGACCTTGCAG
        maximum likelihood sequence       CCGGATTCGACCTTGCAG
        maximum likelihood edit distance  0

evaluating sequence 89 of 100
        observed sequence                 CTGCAAGGTCGAATCCGGTC
        maximum likelihood sequence       CTGCAAGGTCGAATCCGGTC
        maximum likelihood edit distance  0

evaluating sequence 90 of 100
        observed sequence                 GACCGGATTCGACCTTGCAG
        maximum likelihood sequence       GACCGGATTCGACCTTGCAG
        maximum likelihood edit distance  0

evaluating sequence 91 of 100
        observed sequence                 CCTGCAAGGTCGAATCCGGTC
        maximum likelihood sequence       CCTGCAAGGTCGAATCCGGTC
        maximum likelihood edit distance  0

evaluating sequence 92 of 100
        observed sequence                 CCTGCAAGGTCGAATCCGGT
        maximum likelihood sequence       CCTGCAAGGTCGAATCCGGT
        maximum likelihood edit distance  0

evaluating sequence 93 of 100
        observed sequence                 GGACCGGATTCGACCTTGCAG
        maximum likelihood sequence       GGACCGGATTCGACCTTGCAG
        maximum likelihood edit distance  0

evaluating sequence 94 of 100
        observed sequence                 GACCGGATTCGACCTTGCAG
        maximum likelihood sequence       GACCGGATTCGACCTTGCAG
        maximum likelihood edit distance  0

evaluating sequence 95 of 100
        observed sequence                 CTGCAAGGTCGAATCCGGTC
        maximum likelihood sequence       CTGCAAGGTCGAATCCGGTC
        maximum likelihood edit distance  0

evaluating sequence 96 of 100
        observed sequence                 CCTGCAAGGTCGAATCCGGTC
        maximum likelihood sequence       CCTGCAAGGTCGAATCCGGTC
        maximum likelihood edit distance  0

evaluating sequence 97 of 100
        observed sequence                 CCTGCAAGGTCGAATCCGGTC
        maximum likelihood sequence       CCTGCAAGGTCGAATCCGGTC
        maximum likelihood edit distance  0

evaluating sequence 98 of 100
        observed sequence                 CTGCAAGGTCGAATCCGGTC
        maximum likelihood sequence       CTGCAAGGTCGAATCCGGTC
        maximum likelihood edit distance  0
```

```
evaluating sequence 99 of 100
        observed sequence                 CTGCAAGGTCGAATCCGGT
        maximum likelihood sequence       CTGCAAGGTCGAATCCGGT
        maximum likelihood edit distance  0

evaluating sequence 100 of 100
        observed sequence                 GACCGGATTCGACCTTGCAG
        maximum likelihood sequence       GACCGGATTCGACCTTGCAG
        maximum likelihood edit distance  0

DATASET STATISTICS:
        assumed error rate    5.555555555555555%
        total bases observed  1993
        total edits accepted  0
        inferred error rate   0.0%
```

In [27]:
```julia
k = 19
canonical_kmers = collect(keys(Eisenia.count_canonical_kmers(maximum_likelihood_observations, k)))
stranded_kmer_graph = Eisenia.build_stranded_kmer_graph(canonical_kmers, maximum_likelihood_observations)
filename = reference_sequence_id * "." * replace(string(Dates.now()), ':' => '.') * ".svg"
Eisenia.plot_stranded_kmer_graph(stranded_kmer_graph, filename=filename)
HTML("""
<image src="$filename" width=50%>
""")
```

```
┌ Warning: skipping sequence shorter than k with id KA0V_7_7_7_11_13_17 & length 18
└ @ Eisenia /Users/Cameron/Desktop/Microbes/Eisenia/src/Eisenia.jl:1120
┌ Warning: skipping sequence shorter than k with id viPE_7_7_7_11_13_17 & length 18
└ @ Eisenia /Users/Cameron/Desktop/Microbes/Eisenia/src/Eisenia.jl:1120
┌ Warning: skipping sequence shorter than k with id PFiF_7_7_7_11_13_17 & length 18
└ @ Eisenia /Users/Cameron/Desktop/Microbes/Eisenia/src/Eisenia.jl:1120
┌ Warning: skipping sequence shorter than k with id 5Acz_7_7_7_11_13_17 & length 17
└ @ Eisenia /Users/Cameron/Desktop/Microbes/Eisenia/src/Eisenia.jl:1120
┌ Warning: skipping sequence shorter than k with id yg6g_7_7_7_11_13_17 & length 18
└ @ Eisenia /Users/Cameron/Desktop/Microbes/Eisenia/src/Eisenia.jl:1120
┌ Warning: skipping sequence shorter than k with id a0nt_7_7_7_11_13_17 & length 18
└ @ Eisenia /Users/Cameron/Desktop/Microbes/Eisenia/src/Eisenia.jl:1120
┌ Warning: skipping sequence shorter than k with id AMhH_7_7_7_11_13_17 & length 18
└ @ Eisenia /Users/Cameron/Desktop/Microbes/Eisenia/src/Eisenia.jl:1120
┌ Warning: skipping sequence shorter than k with id TGjD_7_7_7_11_13_17 & length 18
└ @ Eisenia /Users/Cameron/Desktop/Microbes/Eisenia/src/Eisenia.jl:1120
```

Out[27]:

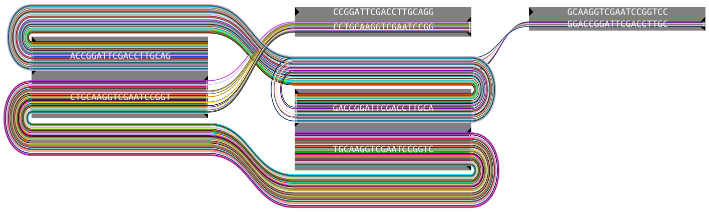


In [28]:
```julia
maximum_likelihood_observations = Eisenia.viterbi_maximum_likelihood_traversals(stranded_kmer_graph, verbosity="reads");
```

```
computing kmer counts...
computing kmer state likelihoods...
finding shortest paths between kmers...
finding viterbi maximum likelihood paths for observed sequences...

evaluating sequence 1 of 92
        observed sequence                 GACCGGATTCGACCTTGCAG
        maximum likelihood sequence       GACCGGATTCGACCTTGCAG
        maximum likelihood edit distance  0

evaluating sequence 2 of 92
        observed sequence                 CCTGCAAGGTCGAATCCGGTC
        maximum likelihood sequence       CCTGCAAGGTCGAATCCGGTC
        maximum likelihood edit distance  0

evaluating sequence 3 of 92
        observed sequence                 GACCGGATTCGACCTTGCAG
        maximum likelihood sequence       GACCGGATTCGACCTTGCAG
        maximum likelihood edit distance  0

evaluating sequence 4 of 92
        observed sequence                 GACCGGATTCGACCTTGCAG
        maximum likelihood sequence       GACCGGATTCGACCTTGCAG
        maximum likelihood edit distance  0

evaluating sequence 5 of 92
        observed sequence                 GACCGGATTCGACCTTGCAG
        maximum likelihood sequence       GACCGGATTCGACCTTGCAG
        maximum likelihood edit distance  0

evaluating sequence 6 of 92
        observed sequence                 GACCGGATTCGACCTTGCAG
        maximum likelihood sequence       GACCGGATTCGACCTTGCAG
        maximum likelihood edit distance  0

evaluating sequence 7 of 92
        observed sequence                 CTGCAAGGTCGAATCCGGTC
        maximum likelihood sequence       CTGCAAGGTCGAATCCGGTC
        maximum likelihood edit distance  0

evaluating sequence 8 of 92
        observed sequence                 CTGCAAGGTCGAATCCGGTC
        maximum likelihood sequence       CTGCAAGGTCGAATCCGGTC
        maximum likelihood edit distance  0

evaluating sequence 9 of 92
        observed sequence                 GACCGGATTCGACCTTGCAG
        maximum likelihood sequence       GACCGGATTCGACCTTGCAG
        maximum likelihood edit distance  0

evaluating sequence 10 of 92
        observed sequence                 CTGCAAGGTCGAATCCGGTC
        maximum likelihood sequence       CTGCAAGGTCGAATCCGGTC
        maximum likelihood edit distance  0

evaluating sequence 11 of 92
        observed sequence                 CTGCAAGGTCGAATCCGGT
        maximum likelihood sequence       CTGCAAGGTCGAATCCGGT
        maximum likelihood edit distance  0

evaluating sequence 12 of 92
        observed sequence                 GACCGGATTCGACCTTGCAG
        maximum likelihood sequence       GACCGGATTCGACCTTGCAG
        maximum likelihood edit distance  0

evaluating sequence 13 of 92
        observed sequence                 TGCAAGGTCGAATCCGGTC
        maximum likelihood sequence       TGCAAGGTCGAATCCGGTC
        maximum likelihood edit distance  0

evaluating sequence 14 of 92
        observed sequence                 GACCGGATTCGACCTTGCAG
        maximum likelihood sequence       GACCGGATTCGACCTTGCAG
        maximum likelihood edit distance  0

evaluating sequence 15 of 92
        observed sequence                 CCTGCAAGGTCGAATCCGGTC
        maximum likelihood sequence       CCTGCAAGGTCGAATCCGGTC
        maximum likelihood edit distance  0

evaluating sequence 16 of 92
        observed sequence                 CTGCAAGGTCGAATCCGGTC
        maximum likelihood sequence       CTGCAAGGTCGAATCCGGTC
        maximum likelihood edit distance  0

evaluating sequence 17 of 92
        observed sequence                 CTGCAAGGTCGAATCCGGTC
        maximum likelihood sequence       CTGCAAGGTCGAATCCGGTC
        maximum likelihood edit distance  0

evaluating sequence 18 of 92
        observed sequence                 GACCGGATTCGACCTTGCAG
        maximum likelihood sequence       GACCGGATTCGACCTTGCAG
        maximum likelihood edit distance  0

evaluating sequence 19 of 92
        observed sequence                 GACCGGATTCGACCTTGCAG
        maximum likelihood sequence       GACCGGATTCGACCTTGCAG
        maximum likelihood edit distance  0
```

```
evaluating sequence 20 of 92
        observed sequence                 GACCGGATTCGACCTTGCA
        maximum likelihood sequence       GACCGGATTCGACCTTGCA
        maximum likelihood edit distance  0

evaluating sequence 21 of 92
        observed sequence                 CTGCAAGGTCGAATCCGGTC
        maximum likelihood sequence       CTGCAAGGTCGAATCCGGTC
        maximum likelihood edit distance  0

evaluating sequence 22 of 92
        observed sequence                 GACCGGATTCGACCTTGCAG
        maximum likelihood sequence       GACCGGATTCGACCTTGCAG
        maximum likelihood edit distance  0

evaluating sequence 23 of 92
        observed sequence                 GACCGGATTCGACCTTGCAG
        maximum likelihood sequence       GACCGGATTCGACCTTGCAG
        maximum likelihood edit distance  0

evaluating sequence 24 of 92
        observed sequence                 CTGCAAGGTCGAATCCGGTC
        maximum likelihood sequence       CTGCAAGGTCGAATCCGGTC
        maximum likelihood edit distance  0

evaluating sequence 25 of 92
        observed sequence                 GACCGGATTCGACCTTGCAG
        maximum likelihood sequence       GACCGGATTCGACCTTGCAG
        maximum likelihood edit distance  0

evaluating sequence 26 of 92
        observed sequence                 GACCGGATTCGACCTTGCAG
        maximum likelihood sequence       GACCGGATTCGACCTTGCAG
        maximum likelihood edit distance  0

evaluating sequence 27 of 92
        observed sequence                 CTGCAAGGTCGAATCCGGTC
        maximum likelihood sequence       CTGCAAGGTCGAATCCGGTC
        maximum likelihood edit distance  0

evaluating sequence 28 of 92
        observed sequence                 CTGCAAGGTCGAATCCGGTC
        maximum likelihood sequence       CTGCAAGGTCGAATCCGGTC
        maximum likelihood edit distance  0

evaluating sequence 29 of 92
        observed sequence                 CTGCAAGGTCGAATCCGGTC
        maximum likelihood sequence       CTGCAAGGTCGAATCCGGTC
        maximum likelihood edit distance  0

evaluating sequence 30 of 92
        observed sequence                 GACCGGATTCGACCTTGCAG
        maximum likelihood sequence       GACCGGATTCGACCTTGCAG
        maximum likelihood edit distance  0

evaluating sequence 31 of 92
        observed sequence                 GACCGGATTCGACCTTGCAG
        maximum likelihood sequence       GACCGGATTCGACCTTGCAG
        maximum likelihood edit distance  0

evaluating sequence 32 of 92
        observed sequence                 CTGCAAGGTCGAATCCGGTC
        maximum likelihood sequence       CTGCAAGGTCGAATCCGGTC
        maximum likelihood edit distance  0

evaluating sequence 33 of 92
        observed sequence                 CCTGCAAGGTCGAATCCGGTC
        maximum likelihood sequence       CCTGCAAGGTCGAATCCGGTC
        maximum likelihood edit distance  0

evaluating sequence 34 of 92
        observed sequence                 GACCGGATTCGACCTTGCAG
        maximum likelihood sequence       GACCGGATTCGACCTTGCAG
        maximum likelihood edit distance  0

evaluating sequence 35 of 92
        observed sequence                 CCTGCAAGGTCGAATCCGGTC
        maximum likelihood sequence       CCTGCAAGGTCGAATCCGGTC
        maximum likelihood edit distance  0

evaluating sequence 36 of 92
        observed sequence                 CTGCAAGGTCGAATCCGGTC
        maximum likelihood sequence       CTGCAAGGTCGAATCCGGTC
        maximum likelihood edit distance  0

evaluating sequence 37 of 92
        observed sequence                 GGACCGGATTCGACCTTGCA
        maximum likelihood sequence       GGACCGGATTCGACCTTGCA
        maximum likelihood edit distance  0

evaluating sequence 38 of 92
        observed sequence                 CTGCAAGGTCGAATCCGGTC
        maximum likelihood sequence       CTGCAAGGTCGAATCCGGTC
        maximum likelihood edit distance  0

evaluating sequence 39 of 92
        observed sequence                 GACCGGATTCGACCTTGCAG
        maximum likelihood sequence       GACCGGATTCGACCTTGCAG
```

```
        maximum likelihood edit distance  0

evaluating sequence 40 of 92
        observed sequence                 CCTGCAAGGTCGAATCCGGTC
        maximum likelihood sequence       CCTGCAAGGTCGAATCCGGTC
        maximum likelihood edit distance  0

evaluating sequence 41 of 92
        observed sequence                 CTGCAAGGTCGAATCCGGTC
        maximum likelihood sequence       CTGCAAGGTCGAATCCGGTC
        maximum likelihood edit distance  0

evaluating sequence 42 of 92
        observed sequence                 GGACCGGATTCGACCTTGCAG
        maximum likelihood sequence       GGACCGGATTCGACCTTGCAG
        maximum likelihood edit distance  0

evaluating sequence 43 of 92
        observed sequence                 GACCGGATTCGACCTTGCAG
        maximum likelihood sequence       GACCGGATTCGACCTTGCAG
        maximum likelihood edit distance  0

evaluating sequence 44 of 92
        observed sequence                 ACCGGATTCGACCTTGCAG
        maximum likelihood sequence       ACCGGATTCGACCTTGCAG
        maximum likelihood edit distance  0

evaluating sequence 45 of 92
        observed sequence                 CTGCAAGGTCGAATCCGGTC
        maximum likelihood sequence       CTGCAAGGTCGAATCCGGTC
        maximum likelihood edit distance  0

evaluating sequence 46 of 92
        observed sequence                 GACCGGATTCGACCTTGCAG
        maximum likelihood sequence       GACCGGATTCGACCTTGCAG
        maximum likelihood edit distance  0

evaluating sequence 47 of 92
        observed sequence                 CTGCAAGGTCGAATCCGGTC
        maximum likelihood sequence       CTGCAAGGTCGAATCCGGTC
        maximum likelihood edit distance  0

evaluating sequence 48 of 92
        observed sequence                 TGCAAGGTCGAATCCGGTC
        maximum likelihood sequence       TGCAAGGTCGAATCCGGTC
        maximum likelihood edit distance  0

evaluating sequence 49 of 92
        observed sequence                 GACCGGATTCGACCTTGCAG
        maximum likelihood sequence       GACCGGATTCGACCTTGCAG
        maximum likelihood edit distance  0

evaluating sequence 50 of 92
        observed sequence                 GACCGGATTCGACCTTGCAG
        maximum likelihood sequence       GACCGGATTCGACCTTGCAG
        maximum likelihood edit distance  0

evaluating sequence 51 of 92
        observed sequence                 CCTGCAAGGTCGAATCCGGTC
        maximum likelihood sequence       CCTGCAAGGTCGAATCCGGTC
        maximum likelihood edit distance  0

evaluating sequence 52 of 92
        observed sequence                 ACCGGATTCGACCTTGCAG
        maximum likelihood sequence       ACCGGATTCGACCTTGCAG
        maximum likelihood edit distance  0

evaluating sequence 53 of 92
        observed sequence                 CTGCAAGGTCGAATCCGGTC
        maximum likelihood sequence       CTGCAAGGTCGAATCCGGTC
        maximum likelihood edit distance  0

evaluating sequence 54 of 92
        observed sequence                 GACCGGATTCGACCTTGCAG
        maximum likelihood sequence       GACCGGATTCGACCTTGCAG
        maximum likelihood edit distance  0

evaluating sequence 55 of 92
        observed sequence                 ACCGGATTCGACCTTGCAG
        maximum likelihood sequence       ACCGGATTCGACCTTGCAG
        maximum likelihood edit distance  0

evaluating sequence 56 of 92
        observed sequence                 CTGCAAGGTCGAATCCGGTC
        maximum likelihood sequence       CTGCAAGGTCGAATCCGGTC
        maximum likelihood edit distance  0

evaluating sequence 57 of 92
        observed sequence                 CTGCAAGGTCGAATCCGGTC
        maximum likelihood sequence       CTGCAAGGTCGAATCCGGTC
        maximum likelihood edit distance  0

evaluating sequence 58 of 92
        observed sequence                 GACCGGATTCGACCTTGCAG
        maximum likelihood sequence       GACCGGATTCGACCTTGCAG
        maximum likelihood edit distance  0

evaluating sequence 59 of 92
        observed sequence                 TGCAAGGTCGAATCCGGTC
```

```
        maximum likelihood sequence       TGCAAGGTCGAATCCGGTC
        maximum likelihood edit distance  0

evaluating sequence 60 of 92
        observed sequence                 CTGCAAGGTCGAATCCGGTC
        maximum likelihood sequence       CTGCAAGGTCGAATCCGGTC
        maximum likelihood edit distance  0

evaluating sequence 61 of 92
        observed sequence                 CTGCAAGGTCGAATCCGGTC
        maximum likelihood sequence       CTGCAAGGTCGAATCCGGTC
        maximum likelihood edit distance  0

evaluating sequence 62 of 92
        observed sequence                 GGACCGGATTCGACCTTGCA
        maximum likelihood sequence       GGACCGGATTCGACCTTGCA
        maximum likelihood edit distance  0

evaluating sequence 63 of 92
        observed sequence                 CTGCAAGGTCGAATCCGGTC
        maximum likelihood sequence       CTGCAAGGTCGAATCCGGTC
        maximum likelihood edit distance  0

evaluating sequence 64 of 92
        observed sequence                 CCTGCAAGGTCGAATCCGGTC
        maximum likelihood sequence       CCTGCAAGGTCGAATCCGGTC
        maximum likelihood edit distance  0

evaluating sequence 65 of 92
        observed sequence                 GACCGGATTCGACCTTGCAG
        maximum likelihood sequence       GACCGGATTCGACCTTGCAG
        maximum likelihood edit distance  0

evaluating sequence 66 of 92
        observed sequence                 GACCGGATTCGACCTTGCAG
        maximum likelihood sequence       GACCGGATTCGACCTTGCAG
        maximum likelihood edit distance  0

evaluating sequence 67 of 92
        observed sequence                 CCTGCAAGGTCGAATCCGGTC
        maximum likelihood sequence       CCTGCAAGGTCGAATCCGGTC
        maximum likelihood edit distance  0

evaluating sequence 68 of 92
        observed sequence                 GACCGGATTCGACCTTGCAG
        maximum likelihood sequence       GACCGGATTCGACCTTGCAG
        maximum likelihood edit distance  0

evaluating sequence 69 of 92
        observed sequence                 GGACCGGATTCGACCTTGCAG
        maximum likelihood sequence       GGACCGGATTCGACCTTGCAG
        maximum likelihood edit distance  0

evaluating sequence 70 of 92
        observed sequence                 CTGCAAGGTCGAATCCGGTC
        maximum likelihood sequence       CTGCAAGGTCGAATCCGGTC
        maximum likelihood edit distance  0

evaluating sequence 71 of 92
        observed sequence                 GGACCGGATTCGACCTTGCAG
        maximum likelihood sequence       GGACCGGATTCGACCTTGCAG
        maximum likelihood edit distance  0

evaluating sequence 72 of 92
        observed sequence                 CTGCAAGGTCGAATCCGGTC
        maximum likelihood sequence       CTGCAAGGTCGAATCCGGTC
        maximum likelihood edit distance  0

evaluating sequence 73 of 92
        observed sequence                 CTGCAAGGTCGAATCCGGTC
        maximum likelihood sequence       CTGCAAGGTCGAATCCGGTC
        maximum likelihood edit distance  0

evaluating sequence 74 of 92
        observed sequence                 CCTGCAAGGTCGAATCCGGTC
        maximum likelihood sequence       CCTGCAAGGTCGAATCCGGTC
        maximum likelihood edit distance  0

evaluating sequence 75 of 92
        observed sequence                 GGACCGGATTCGACCTTGCAG
        maximum likelihood sequence       GGACCGGATTCGACCTTGCAG
        maximum likelihood edit distance  0

evaluating sequence 76 of 92
        observed sequence                 GACCGGATTCGACCTTGCAG
        maximum likelihood sequence       GACCGGATTCGACCTTGCAG
        maximum likelihood edit distance  0

evaluating sequence 77 of 92
        observed sequence                 CTGCAAGGTCGAATCCGGTC
        maximum likelihood sequence       CTGCAAGGTCGAATCCGGTC
        maximum likelihood edit distance  0

evaluating sequence 78 of 92
        observed sequence                 CCTGCAAGGTCGAATCCGGTC
        maximum likelihood sequence       CCTGCAAGGTCGAATCCGGTC
        maximum likelihood edit distance  0

evaluating sequence 79 of 92
```

```
        observed sequence                 CCTGCAAGGTCGAATCCGGTC
        maximum likelihood sequence       CCTGCAAGGTCGAATCCGGTC
        maximum likelihood edit distance  0

evaluating sequence 80 of 92
        observed sequence                 GACCGGATTCGACCTTGCAG
        maximum likelihood sequence       GACCGGATTCGACCTTGCAG
        maximum likelihood edit distance  0

evaluating sequence 81 of 92
        observed sequence                 CTGCAAGGTCGAATCCGGTC
        maximum likelihood sequence       CTGCAAGGTCGAATCCGGTC
        maximum likelihood edit distance  0

evaluating sequence 82 of 92
        observed sequence                 GACCGGATTCGACCTTGCAG
        maximum likelihood sequence       GACCGGATTCGACCTTGCAG
        maximum likelihood edit distance  0

evaluating sequence 83 of 92
        observed sequence                 CCTGCAAGGTCGAATCCGGTC
        maximum likelihood sequence       CCTGCAAGGTCGAATCCGGTC
        maximum likelihood edit distance  0

evaluating sequence 84 of 92
        observed sequence                 CCTGCAAGGTCGAATCCGGT
        maximum likelihood sequence       CCTGCAAGGTCGAATCCGGT
        maximum likelihood edit distance  0

evaluating sequence 85 of 92
        observed sequence                 GGACCGGATTCGACCTTGCAG
        maximum likelihood sequence       GGACCGGATTCGACCTTGCAG
        maximum likelihood edit distance  0

evaluating sequence 86 of 92
        observed sequence                 GACCGGATTCGACCTTGCAG
        maximum likelihood sequence       GACCGGATTCGACCTTGCAG
        maximum likelihood edit distance  0

evaluating sequence 87 of 92
        observed sequence                 CTGCAAGGTCGAATCCGGTC
        maximum likelihood sequence       CTGCAAGGTCGAATCCGGTC
        maximum likelihood edit distance  0

evaluating sequence 88 of 92
        observed sequence                 CCTGCAAGGTCGAATCCGGTC
        maximum likelihood sequence       CCTGCAAGGTCGAATCCGGTC
        maximum likelihood edit distance  0

evaluating sequence 89 of 92
        observed sequence                 CCTGCAAGGTCGAATCCGGTC
        maximum likelihood sequence       CCTGCAAGGTCGAATCCGGTC
        maximum likelihood edit distance  0

evaluating sequence 90 of 92
        observed sequence                 CTGCAAGGTCGAATCCGGTC
        maximum likelihood sequence       CTGCAAGGTCGAATCCGGTC
        maximum likelihood edit distance  0

evaluating sequence 91 of 92
        observed sequence                 CTGCAAGGTCGAATCCGGT
        maximum likelihood sequence       CTGCAAGGTCGAATCCGGT
        maximum likelihood edit distance  0

evaluating sequence 92 of 92
        observed sequence                 GACCGGATTCGACCTTGCAG
        maximum likelihood sequence       GACCGGATTCGACCTTGCAG
        maximum likelihood edit distance  0

DATASET STATISTICS:
        assumed error rate    5.0%
        total bases observed  1850
        total edits accepted  0
        inferred error rate   0.0%
```

# L50

In [29]:

```
L = 50
Random.seed!(L)
reference_sequence = randdnaseq(L)
reference_sequence_id = randstring(Int(round(log10(length(L)))+3))
reference_FASTA_record = FASTA.Record(reference_sequence_id, reference_sequence)
```

Out[29]:

```
BioSequences.FASTA.Record:
   identifier: j96
  description: <missing>
     sequence: TGGAACCAGGATCATGCTACGGCGCGTAATCTACCACGA…
```

In [30]:

```
error_rate = 0.15
n_sequences = 100
observations = [Eisenia.observe(reference_FASTA_record, error_rate=error_rate) for i in 1:n_sequences]
```

```
Out[30]: 100-element Array{BioSequences.FASTA.Record,1}:
 BioSequences.FASTA.Record:
   identifier: nxKL8
  description: <missing>
     sequence: TGGAACTAGGATCATGCTACGGCGCGAATCTACCAGATG…
 BioSequences.FASTA.Record:
   identifier: h0jcA
  description: <missing>
     sequence: TGGAACCAGATCATGCTACGGCGCGTATTTACCACGATG…
 BioSequences.FASTA.Record:
   identifier: RKMGo
  description: <missing>
     sequence: TGCTAGAAGCACGTGGTAGATTACGCGCTAGCATGATCC…
 BioSequences.FASTA.Record:
   identifier: deuPI
  description: <missing>
     sequence: TGGAACCAAGTCTCAGTGCTTACGGCGCGTAATCTACCC…
 BioSequences.FASTA.Record:
   identifier: HKzzv
  description: <missing>
     sequence: TGGAACCAGATCATCTACGGCGCGTATTCTACCACGACC…
 BioSequences.FASTA.Record:
   identifier: cQcBT
  description: <missing>
     sequence: GGCTAGAAGCACGTGGTAGATTACGCGCTCGGTAGCATG…
 BioSequences.FASTA.Record:
   identifier: CUBmJ
  description: <missing>
     sequence: TGCTAAGAAGCATCTGGTAGATTATGCGCCGTAGCATGA…
 BioSequences.FASTA.Record:
   identifier: dOAzV
  description: <missing>
     sequence: TGGAGCCATGATCATGCTAGGCGGTAATCTACACGGGCT…
 BioSequences.FASTA.Record:
   identifier: mmQ99
  description: <missing>
     sequence: TGCTAAAAGCACTACGTAGGTGATTACGCGCCGTAGCAT…
 BioSequences.FASTA.Record:
   identifier: 2qtSf
  description: <missing>
     sequence: TGCTAGAAGCATCTGGTCGATTCGCGCCGTAGCATGATC…
 BioSequences.FASTA.Record:
   identifier: noSRT
  description: <missing>
     sequence: TGGAACCAGCGATAGTGCTACGTGCGTAATCTACCACGA…
 BioSequences.FASTA.Record:
   identifier: GfF8L
  description: <missing>
     sequence: TGGAACCGGATCATAGCGTACGGCGAGTAATCTACCACG…
 BioSequences.FASTA.Record:
   identifier: Mwne4
  description: <missing>
     sequence: TGTAGAAGGCAAGTGGTAGAATTACGCGCCATAGCATGA…
 ⋮
 BioSequences.FASTA.Record:
   identifier: WuscQ
  description: <missing>
     sequence: TGCTAGATCATCGTGGCTAGATAAAGCGCCGTAGCATGA…
 BioSequences.FASTA.Record:
   identifier: sqr0C
  description: <missing>
     sequence: TGGAACCAGATCAGCACGGGGCGTATATCTACCACGATC…
 BioSequences.FASTA.Record:
   identifier: rFxJB
  description: <missing>
     sequence: TGGAACCAGGATATGCTACGGCGCGTAATCTACCACGAT…
 BioSequences.FASTA.Record:
   identifier: 5Rj2k
  description: <missing>
     sequence: TGCTAGAAGCATATCGTGTAGATTACCAGCCGTAGATGT…
 BioSequences.FASTA.Record:
   identifier: g0COT
  description: <missing>
     sequence: TGCTAGAACGCATCGTGGTAGCTTACGCGCCGAGCATGA…
 BioSequences.FASTA.Record:
   identifier: KvngM
  description: <missing>
     sequence: TGGAAACAGATCATACAGACGCGTATCTAACCACGAAGC…
 BioSequences.FASTA.Record:
   identifier: k4dEC
  description: <missing>
     sequence: TTGCTAAAGCATCGTGGTAATTCGCGCGTAGCATGATCC…
 BioSequences.FASTA.Record:
   identifier: EudvV
  description: <missing>
     sequence: TGCACCAGGATACATGCTAGCGGCCGCGTAATCTACCAC…
 BioSequences.FASTA.Record:
   identifier: N58zx
  description: <missing>
     sequence: TTGCTAGAAGCAATCGTGGTAGATTACGCCCGGTAGCAT…
 BioSequences.FASTA.Record:
   identifier: Kiap5
  description: <missing>
     sequence: TGGATACAGGAATCATGCTGCGGCAGCGTATCTACCACG…
 BioSequences.FASTA.Record:
   identifier: TBsym
  description: <missing>
     sequence: TGCTAGAAGCATCGTGGTAGATTACGCGCGTAGCATGAT…
 BioSequences.FASTA.Record:
```

```
 identifier: fzWjN
description: <missing>
   sequence: GTGGAACCAGAGTCATGCTCCGCGCGTAATCTCCCACGT…
```

#### L50 starting @ K=7

In [31]:
```
k = 7
canonical_kmers = collect(keys(Eisenia.count_canonical_kmers(observations, k)))
stranded_kmer_graph = Eisenia.build_stranded_kmer_graph(canonical_kmers, observations)
filename = reference_sequence_id * "." * replace(string(Dates.now()), ':' => '.') * ".svg"
Eisenia.plot_stranded_kmer_graph(stranded_kmer_graph, filename=filename)
HTML("""
<image src="$filename" width=50%>
""")
```

Out[31]:

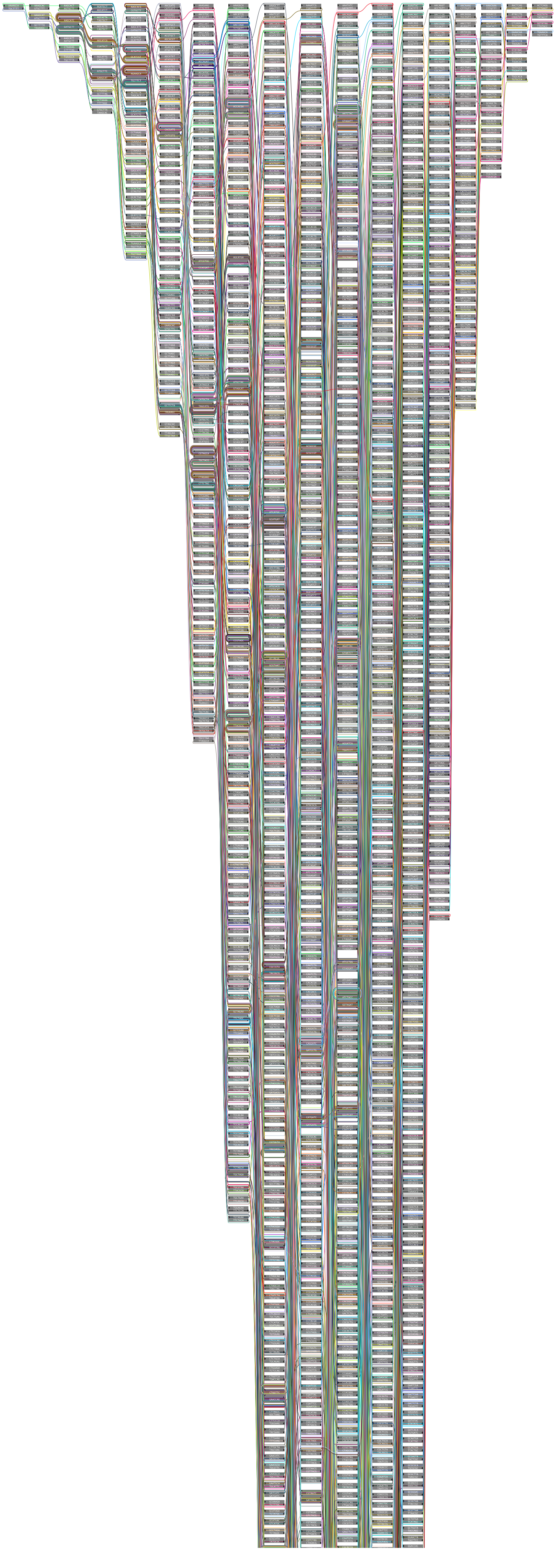

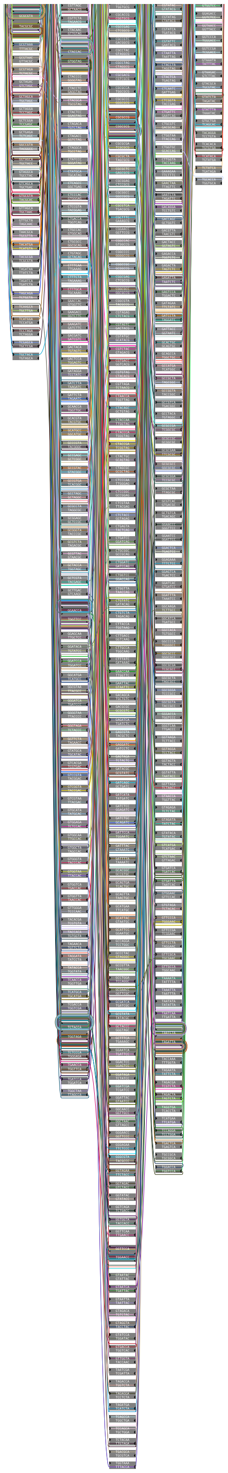

```
In [32]: maximum_likelihood_observations = Eisenia.viterbi_maximum_likelihood_traversals(stranded_kmer_graph, verbosity="reads");
```

```
computing kmer counts...
computing kmer state likelihoods...
finding shortest paths between kmers...
finding viterbi maximum likelihood paths for observed sequences...

evaluating sequence 1 of 100
        observed sequence                 TGGAACTAGGATCATGCTACGGCGCGAATCTACCAGATGCTTCTCCA
        maximum likelihood sequence       TGGAACCAGGATCATGCTACGGCGCGTAATCTACCACGATGCTTCTAGCA
        maximum likelihood edit distance  5

evaluating sequence 2 of 100
        observed sequence                 TGGAACCAGATCATGCTACGGCGCGTATTTACCACGATGTCTAGCA
        maximum likelihood sequence       TGGAACCAGGATCATGCTACGGCGCGTAATCTACCACGATGCTTCTAGCA
        maximum likelihood edit distance  5

evaluating sequence 3 of 100
        observed sequence                 TGCTAGAAGCACGTGGTAGATTACGCGCTAGCATGATCCTAGTCCA
        maximum likelihood sequence       TGCTAGAAGCATCGTGGTAGATTACGCGCCGTAGCATGATCCTGGTTCCA
        maximum likelihood edit distance  5

evaluating sequence 4 of 100
        observed sequence                 TGGAACCAAGTCTCAGTGCTTACGGCGCGTAATCTACCCGATGCTTCCGAGCA
        maximum likelihood sequence       TGGAACCAGGATCATGCTACGGCGCGTAATCTACCACGATGCTTCTAGCA
        maximum likelihood edit distance  9

evaluating sequence 5 of 100
        observed sequence                 TGGAACCAGATCATCTACGGCGCGTATTCTACCACGACCGCTGTCTAGCA
        maximum likelihood sequence       TGGAACCAGGATCATGCTACGGCGCGTAATCTACCACGATGCTTCTAGCA
        maximum likelihood edit distance  7

evaluating sequence 6 of 100
        observed sequence                 GGCTAGAAGCACGTGGTAGATTACGCGCTCGGTAGCATGTAGTCTCCTGAGTTCCA
        maximum likelihood sequence       GGCTAGAAGCATCGTGGTAGATTACGCGCCGTAGCATGATCCTGGTTCCA
        maximum likelihood edit distance  11

evaluating sequence 7 of 100
        observed sequence                 TGCTAAGAAGCATCTGGTAGATTATGCGCCGTAGCATGATCTGGTTCCA
        maximum likelihood sequence       TTGCTAGAAGCATCGTGGTAGATTACGCGCCGTAGCATGATCCTGGTTCCA
        maximum likelihood edit distance  4

evaluating sequence 8 of 100
        observed sequence                 TGGAGCCATGATCATGCTAGGCGGTAATCTACACGGGCTTCTAGCA
        maximum likelihood sequence       TGGAACCAGGATCATGCTACGGCGCGTAATCTACCACGATGCTTCTAGCA
        maximum likelihood edit distance  7

evaluating sequence 9 of 100
        observed sequence                 TGCTAAAAGCACTACGTAGGTGATTACGCGCCGTAGCATGATCACTGTGGTCCA
        maximum likelihood sequence       TGCTAGAAGCATCGTGGTAGATTACGCGCCGTAGCATGATCCTGGTTCCA
        maximum likelihood edit distance  12

evaluating sequence 10 of 100
        observed sequence                 TGCTAGAAGCATCTGGTCGATTCGCGCCGTAGCATGATCTGGTTCCA
        maximum likelihood sequence       TGCTAGAAGCATCGTGGTAGATTACGCGCCGTAGCATGATCCTGGTTCCA
        maximum likelihood edit distance  4

evaluating sequence 11 of 100
        observed sequence                 TGGAACCAGCGATAGTGCTACGTGCGTAATCTACCACGATTTTAGTCA
        maximum likelihood sequence       TGGAACCAGGATCATGCTACGGCGCGTAATCTACCACGATGCTTCTAGCA
        maximum likelihood edit distance  10

evaluating sequence 12 of 100
        observed sequence                 TGGAACCGGATCATAGCGTACGGCGAGTAATCTACCACGATGCTTCTAGCG
        maximum likelihood sequence       TGGAACCAGGATCATGCTACGGCGCGTAATCTACCACGATGCTTCTAGCA
        maximum likelihood edit distance  7

evaluating sequence 13 of 100
        observed sequence                 TGTAGAAGGCAAGTGGTAGAATTACGCGCCATAGCATGATCCATGGTTCA
        maximum likelihood sequence       TGCTAGAAGCATCGTGGTAGATTACGCGCCGTAGCATGATCCTGGTTCA
        maximum likelihood edit distance  8

evaluating sequence 14 of 100
        observed sequence                 TGCTAGAAGCATGGTGGTAGATTACGCGCCATAGCATGATCCTGGTTCCA
        maximum likelihood sequence       TGCTAGAAGCATCGTGGTAGATTACGCGCCGTAGCATGATCCTGGTTCCA
        maximum likelihood edit distance  2

evaluating sequence 15 of 100
        observed sequence                 GACCAGGATCATGCTAAGGCGCGTAATCTACTCAGCATGCTTCTAGCA
        maximum likelihood sequence       GACCAGGATCATGCTACGGCGCGTAATCTACCACGATGCTTCTAGCA
        maximum likelihood edit distance  5

evaluating sequence 16 of 100
        observed sequence                 TGCTAGAAGCATCTGGAGATTTACGCCCGTAGCATGACCCTGGTTCCTA
        maximum likelihood sequence       TGCTAGAAGCATCGTGGTAGATTACGCGCCGTAGCATGATCCTGGTTCCA
        maximum likelihood edit distance  7

evaluating sequence 17 of 100
        observed sequence                 TGCTTAGAAGCATCGGGGTAGATTACGCGCCGTGAGCATGATCCTGGTTCA
        maximum likelihood sequence       TGCTAGAAGCATCGTGGTAGATTACGCGCCGTAGCATGATCCTGGTTCA
        maximum likelihood edit distance  4

evaluating sequence 18 of 100
        observed sequence                 TGCTAGAAGCATCGGGATAGATTACGCCCGTAGCAGATCCTGGGGCAA
        maximum likelihood sequence       TGCTAGAAGCATCGTGGTAGATTACGCGCCGTAGCATGATCCTGGTTCCA
        maximum likelihood edit distance  7

evaluating sequence 19 of 100
        observed sequence                 TGCTGAAGCATCGTGGTAGATTACGCGCCTAGCATGATCCTGGTTCCA
        maximum likelihood sequence       GCTAGAAGCATCGTGGTAGATTACGCGCCGTAGCATGATCCTGGTTCCA
        maximum likelihood edit distance  2
```

```
evaluating sequence 20 of 100
        observed sequence                TGTAGAAGCAATCGTGGTAGATTACGCGCCGTAGCATGATCTCGCCA
        maximum likelihood sequence      TGTAGAAGCATCGTGGTAGATTACGCGCCGTAGCATGATCCTGGTTCCA
        maximum likelihood edit distance  5

evaluating sequence 21 of 100
        observed sequence                TGGAACCAGGATCATGCTAACGGTGCGTAAGCTACCCACGATGCTTCTTTGCA
        maximum likelihood sequence      TGGAACCAGGATCATGCTACGGCGCGTAATCTACCACGATGCTTCTAGCA
        maximum likelihood edit distance  6

evaluating sequence 22 of 100
        observed sequence                TGAAACAAAGATCAATGCGTACGGCGCGTTAATCTATGCACGATGCGCCTAGCA
        maximum likelihood sequence      TGGAACCAGGATCATGCTACGGCGCGTAATCTACCACGATGCTTCTAGCA
        maximum likelihood edit distance  12

evaluating sequence 23 of 100
        observed sequence                TGCTGAAGCAGTCGTGGTAGATTACGCACTGCAGATCCTGGATCCA
        maximum likelihood sequence      GCTAGAAGCATCGTGGTAGATTACGCGCCGTAGCATGATCCTGGTTCCA
        maximum likelihood edit distance  9

evaluating sequence 24 of 100
        observed sequence                TGGAACCAGAGATCATGCTACGGCGGTAATCTCCACGATCTTAGCCA
        maximum likelihood sequence      TGGAACCAGGATCATGCTACGGCGCGTAATCTACCACGATGCTTCTAGCA
        maximum likelihood edit distance  7

evaluating sequence 25 of 100
        observed sequence                GTGCTCGAAGCATCGTTGTAGATTACGGGCCGCTAGCATATCCTGGTTCCA
        maximum likelihood sequence      TGCTAGAAGCATCGTGGTAGATTACGCGCCGTAGCATGATCCTGGTTCCA
        maximum likelihood edit distance  8

evaluating sequence 26 of 100
        observed sequence                TGGTACCAGGATCTGCTACGGCCGTAATCTCCACGTGCTTAGCA
        maximum likelihood sequence      TGGAACCAGGATCATGCTACGGCGCGTAATCTACCACGATGCTTCTAGCA
        maximum likelihood edit distance  7

evaluating sequence 27 of 100
        observed sequence                TGGGACCAGGATCATGCTACGGCCGCGTAATCTACCCGATGCTTCTAGCA
        maximum likelihood sequence      TGGAACCAGGATCATGCTACGGCGCGTAATCTACCACGATGCTTCTAGCA
        maximum likelihood edit distance  3

evaluating sequence 28 of 100
        observed sequence                TGCTAGAGGATCGTGTGTGATTACGCGCCGTGCATGACCTGGTCCA
        maximum likelihood sequence      TGCTAGAAGCATCGTGGTAGATTACGCGCCGTAGCATGATCCTGGTTCCA
        maximum likelihood edit distance  8

evaluating sequence 29 of 100
        observed sequence                TGCGTAGAAGCATCGTGGTAGGATTTACGCGCCGGTAGCATTCCTGGTTCCA
        maximum likelihood sequence      TGCTAGAAGCATCGTGGTAGATTACGCGCCGTAGCATGATCCTGGTTCCA
        maximum likelihood edit distance  7

evaluating sequence 30 of 100
        observed sequence                CGATGAGCATCGGGTAGATTTCCGCGCCGTAGCAGATCCTGATTCCA
        maximum likelihood sequence      CTAGAAGCATCGTGGTAGATTACGCGCCGTAGCATGATCCTGGTTCCA
        maximum likelihood edit distance  8

evaluating sequence 31 of 100
        observed sequence                TGGAAACAGCGATCTCTACGGCGCGTAAAATACCACGATGCTTCTAGCA
        maximum likelihood sequence      TGGAACCAGGATCATGCTACGGCGCGTAATCTACCACGATGCTTCTAGCA
        maximum likelihood edit distance  6

evaluating sequence 32 of 100
        observed sequence                TGGAACCAGGATATGCATACGGCGCAGTAATCTACCACGTGCTTCTAAGCA
        maximum likelihood sequence      TGGAACCAGGATCATGCTACGGCGCGTAATCTACCACGATGCTTCTAGCA
        maximum likelihood edit distance  7

evaluating sequence 33 of 100
        observed sequence                TGGAACCAGGATCATTGCTCCGGCGCGTAATCTACCACGCATGCTTCTAGCA
        maximum likelihood sequence      TGGAACCAGGATCATGCTACGGCGCGTAATCTACCACGATGCTTCTAGCA
        maximum likelihood edit distance  4

evaluating sequence 34 of 100
        observed sequence                TGGAATCAGATCTTGCTACGGCGCGTCACATCTACCACGGGTCAAGCA
        maximum likelihood sequence      TGGAACCAGGATCATGCTACGGCGCGTAATCTACCACGATGCTTCTAGCA
        maximum likelihood edit distance  11

evaluating sequence 35 of 100
        observed sequence                TGCTAGAAGCACGTGGTAGATTACGCACCAGTAGCTGTATCCTGTGTGCCA
        maximum likelihood sequence      TGCTAGAAGCATCGTGGTAGATTACGCGCCGTAGCATGATCCTGGTTCCA
        maximum likelihood edit distance  10

evaluating sequence 36 of 100
        observed sequence                TGGATCCAGCAATCTGCTACGGCCACAGCAATCTACCACGATATTTCTAGCA
        maximum likelihood sequence      TGGAACCAGGATCATGCTACGGCGCGTAATCTACCACGATGCTTCTAGCA
        maximum likelihood edit distance  11

evaluating sequence 37 of 100
        observed sequence                TGGAGTCCAGGATCAGGCTAACGGCGCGTAGTCTACCACGGTCTTCTAGCCA
        maximum likelihood sequence      TGGAACCAGGATCATGCTACGGCGCGTAATCTACCACGATGCTTCTAGCA
        maximum likelihood edit distance  9

evaluating sequence 38 of 100
        observed sequence                TGGAACCAGGATCATTCCACGGCGCGTAAATCTACCACGATGCTTCTAGCA
        maximum likelihood sequence      TGGAACCAGGATCATGCTACGGCGCGTAATCTACCACGATGCTTCTAGCA
        maximum likelihood edit distance  3

evaluating sequence 39 of 100
        observed sequence                TGCTAGAGCTCGTGGCTAATTACGCGACGTTCATGATCCTGGTTCCA
        maximum likelihood sequence      GCTAGAAGCATCGTGGTAGATTACGCGCCGTAGCATGATCCTGGTTCCA
```

```
        maximum likelihood edit distance  8

evaluating sequence 40 of 100
        observed sequence                 TGGGAACTAAGATCACTGCTACGGCGCGTAATCTTGCCACGATGCGTCTAGCA
        maximum likelihood sequence       TGGAACCAGGATCATGCTACGGCGCGTAATCTACCACGATGCTTCTAGCA
        maximum likelihood edit distance  10

evaluating sequence 41 of 100
        observed sequence                 TGGAACCTAGGATATGCTATGGCTGCGTATCTACCACGATGGCTTCAGCA
        maximum likelihood sequence       TGGAACCAGGATCATGCTACGGCGCGTAATCTACCACGATGCTTCTAGCA
        maximum likelihood edit distance  9

evaluating sequence 42 of 100
        observed sequence                 AGGACAGGATCATGCTACGGCGCGTAATCAACCAGACTGCTTCTAGC
        maximum likelihood sequence       GAACCAGGATCATGCTACGGCGCGTAATCTACCACGATGCTTCTAGC
        maximum likelihood edit distance  6

evaluating sequence 43 of 100
        observed sequence                 TCGGACCAGCATCATGCTACGGAGCGTAAATCTACCACGATGCTTCTCAGCCA
        maximum likelihood sequence       TGGAACCAGGATCATGCTACGGCGCGTAATCTACCACGATGCTTCTAGCA
        maximum likelihood edit distance  8

evaluating sequence 44 of 100
        observed sequence                 TGCTAGAATCATCGTGGTAGATTACGGGTCCGTAGCAGATCCTGGGTCCTA
        maximum likelihood sequence       TGCTAGAAGCATCGTGGTAGATTACGCGCCGTAGCATGATCCTGGTTCCA
        maximum likelihood edit distance  8

evaluating sequence 45 of 100
        observed sequence                 TGCTAGAGCATGTGGTAGATTAGCGCCGTCAGCATGATCCCGGTACCA
        maximum likelihood sequence       GCTAGAAGCATCGTGGTAGATTACGCGCCGTAGCATGATCCTGGTTCCA
        maximum likelihood edit distance  6

evaluating sequence 46 of 100
        observed sequence                 TCGGAACCACGGCATCATGCTAACGTCGCGTAATCTACCACTATGCTGTCAGGCA
        maximum likelihood sequence       TGGAACCAGGATCATGCTACGGCGCGTAATCTACCACGATGCTTCTAGCA
        maximum likelihood edit distance  13

evaluating sequence 47 of 100
        observed sequence                 TGGACCAGGATCATGCTCGGCGCGTAACTGCTACCACGATGACGCAGCG
        maximum likelihood sequence       GGAACCAGGATCATGCTACGGCGCGTAATCTACCACGATGCTTCTAGCA
        maximum likelihood edit distance  12

evaluating sequence 48 of 100
        observed sequence                 GGCTAGAATGCATCGTGGTATATTACGACGCGTAGGCATGATCCTGGCTTCCA
        maximum likelihood sequence       TGCTAGAAGCATCGTGGTAGATTACGCGCCGTAGCATGATCCTGGTTCCA
        maximum likelihood edit distance  10

evaluating sequence 49 of 100
        observed sequence                 TGCTGAAGATCGGGTAGATTAGCGCGCCGTAGCATGATCCTGGTTCA
        maximum likelihood sequence       GCTAGAAGCATCGTGGTAGATTACGCGCCGTAGCATGATCCTGGTTCA
        maximum likelihood edit distance  4

evaluating sequence 50 of 100
        observed sequence                 TGGAACCAGTATATGCTACGGCGCGTAATCGTGACCACGATGCTTCCTAGCA
        maximum likelihood sequence       TGGAACCAGGATCATGCTACGGCGCGTAATCTACCACGATGCTTCTAGCA
        maximum likelihood edit distance  7

evaluating sequence 51 of 100
        observed sequence                 CGCTCGAAGCATCGTGGTAGATTTCGCGCCGTGGCATGCCTGTTTACC
        maximum likelihood sequence       TGCTAGAAGCATCGTGGTAGATTACGCGCCGTAGCATGATCCTGGTTCC
        maximum likelihood edit distance  9

evaluating sequence 52 of 100
        observed sequence                 TGCAGAGCATCGTGGTAGATTACGAGCCGTATCATGATCCTGGCCA
        maximum likelihood sequence       GTAGAAGCATCGTGGTAGATTACGCGCCGTAGCATGATCCTGGTTCCA
        maximum likelihood edit distance  6

evaluating sequence 53 of 100
        observed sequence                 TGGAATCGGGATCAGTCTAACGGCGCGTAATACCACGATGGCTTCTAGA
        maximum likelihood sequence       TGGAACCAGGATCATGCTACGGCGCGTAATCTACCACGATGCTTCTAGA
        maximum likelihood edit distance  8

evaluating sequence 54 of 100
        observed sequence                 TGGAACCAGGATCATTGTAGGCGCTTAATCTCCCAACGATGCTTCAGCA
        maximum likelihood sequence       TGGAACCAGGATCATGCTACGGCGCGTAATCTACCACGATGCTTCTAGCA
        maximum likelihood edit distance  7

evaluating sequence 55 of 100
        observed sequence                 GTGGAACCAGCTCACTCCTATCGCGCGTAAATTACCACGATGCTTTAGCA
        maximum likelihood sequence       GTGGAACCAGGATCATGCTACGGCGCGTAATCTACCACGATGCTTCTAGCA
        maximum likelihood edit distance  10

evaluating sequence 56 of 100
        observed sequence                 TTGAACTAGGATCATGCACGGCGCGTAATCTACCACCATGCTTCTAGCA
        maximum likelihood sequence       TGGAACCAGGATCATGCTACGGCGCGTAATCTACCACGATGCTTCTAGCA
        maximum likelihood edit distance  4

evaluating sequence 57 of 100
        observed sequence                 GCTAGAATGCAGCGGGTAGAATGGGGCCGTACATGACTCATGGATCCA
        maximum likelihood sequence       TGCTAGAAGCATCGTGGTAGATTACGCGCCGTAGCATGATCCTGGTTCCA
        maximum likelihood edit distance  13

evaluating sequence 58 of 100
        observed sequence                 TGCTAGAAAGAACCGTGGTAAGACTACGCGCTCGTAGCATGATCCTGGTTCCA
        maximum likelihood sequence       TGCTAGAAGCATCGTGGTAGATTACGCGCCGTAGCATGATCCTGGTTCCA
        maximum likelihood edit distance  6

evaluating sequence 59 of 100
        observed sequence                 TGGAACCAAGGATCCATGCTACGGCGCGTAATCTACCAAGATGCCTTTAGCA
```

```
        maximum likelihood sequence       TGGAACCAGGATCATGCTACGGCGCGTAATCTACCACGATGCTTCTAGCA
        maximum likelihood edit distance  5

evaluating sequence 60 of 100
        observed sequence                 TCCTAGAAGCATCGTGGGAGATTACGCGCCGTAGCATGATCACTGGTTCCA
        maximum likelihood sequence       TGCTAGAAGCATCGTGGTAGATTACGCGCCGTAGCATGATCCTGGTTCCA
        maximum likelihood edit distance  3

evaluating sequence 61 of 100
        observed sequence                 AGTAGAAGCATGGTGGTAGCATTATCGGCGCCGTAGCATGATCCAGGCTTCCCA
        maximum likelihood sequence       GCTAGAAGCATCGTGGTAGATTACGCGCCGTAGCATGATCCTGGTTCCA
        maximum likelihood edit distance  11

evaluating sequence 62 of 100
        observed sequence                 TGACCAGGATCATGCTACGGCGCGTATACCAGAGCTTTCAAGCA
        maximum likelihood sequence       TGACCAGGATCATGCTACGGCGCGTAATCTACCACGATGCTTCTAGCA
        maximum likelihood edit distance  7

evaluating sequence 63 of 100
        observed sequence                 TGGACGAGGATCGTCTACGGCGCGTAATCTCCACGATGCATTCTAGA
        maximum likelihood sequence       GGAACCAGGATCATGCTACGGCGCGTAATCTACCACGATGCTTCTAGA
        maximum likelihood edit distance  8

evaluating sequence 64 of 100
        observed sequence                 TGGAACCAGGACTCAATCTACGGCGGGACTCTACCACGAGCTTCGAGCA
        maximum likelihood sequence       TGGAACCAGGATCATGCTACGGCGCGTAATCTACCACGATGCTTCTAGCA
        maximum likelihood edit distance  9

evaluating sequence 65 of 100
        observed sequence                 TGGACACTAGGAGTCATGCTCACGGCGCGTAATCTACCACTGAAGCATCTAGCA
        maximum likelihood sequence       GTGGAACCAGGATCATGCTACGGCGCGTAATCTACCACGATGCTTCTAGCA
        maximum likelihood edit distance  10

evaluating sequence 66 of 100
        observed sequence                 TGGAACCCGGATTATGCTACGCGCGGAATCTGACCAGATGCTTCTAGCA
        maximum likelihood sequence       TGGAACCAGGATCATGCTACGGCGCGTAATCTACCACGATGCTTCTAGCA
        maximum likelihood edit distance  7

evaluating sequence 67 of 100
        observed sequence                 TGGGAACGAGGATCATGTAGGGCGCGTAATCTACCGACGATGCTATCTAGA
        maximum likelihood sequence       TGGAACCAGGATCATGCTACGGCGCGTAATCTACCACGATGCTTCTAGA
        maximum likelihood edit distance  10

evaluating sequence 68 of 100
        observed sequence                 GCTAGAAGCATCGTCGTAGCATTACGCGCCGTACATGATCCTGGTTCCA
        maximum likelihood sequence       GCTAGAAGCATCGTGGTAGATTACGCGCCGTAGCATGATCCTGGTTCCA
        maximum likelihood edit distance  4

evaluating sequence 69 of 100
        observed sequence                 TGCCTAAAGCATTGTGGAGATCACAGCGCAGTAGCAACCTGGTACCA
        maximum likelihood sequence       TGCTAGAAGCATCGTGGTAGATTACGCGCCGTAGCATGATCCTGGTTCCA
        maximum likelihood edit distance  11

evaluating sequence 70 of 100
        observed sequence                 TTGAACCAGCATCAGGTATAACGCCGCATTAATCTACCACGATGCCTTCTAGCA
        maximum likelihood sequence       TGGAACCAGGATCATGCTACGGCGCGTAATCTACCACGATGCTTCTAGCA
        maximum likelihood edit distance  11

evaluating sequence 71 of 100
        observed sequence                 TTGCTAGAAGCACGTGGTAGATTAGCTGCCGTGGCAGTGATCCTGGTTCGCA
        maximum likelihood sequence       TTGCTAGAAGCATCGTGGTAGATTACGCGCCGTAGCATGATCCTGGTTCCA
        maximum likelihood edit distance  8

evaluating sequence 72 of 100
        observed sequence                 TGGAATAGGATCATGCTACGAGCGCGTATCTACCACGTGCTTCTAGC
        maximum likelihood sequence       GGAACCAGGATCATGCTACGGCGCGTAATCTACCACGATGCTTCTAGC
        maximum likelihood edit distance  5

evaluating sequence 73 of 100
        observed sequence                 TGCTAGAACCATCCGTAGGTAGATACGCGCCGTAGCATAGATTCCTGTTCCTA
        maximum likelihood sequence       TGCTAGAAGCATCGTGGTAGATTACGCGCCGTAGCATGATCCTGGTTCCA
        maximum likelihood edit distance  11

evaluating sequence 74 of 100
        observed sequence                 TGCTAGAAGCATCGAGGTAGACTATACGCGCCGTAGCATGATCCTGGTTCA
        maximum likelihood sequence       TGCTAGAAGCATCGTGGTAGATTACGCGCCGTAGCATGATCCTGGTTCA
        maximum likelihood edit distance  5

evaluating sequence 75 of 100
        observed sequence                 TGCTAAAGCATCGGAGTAGATTACGCCGCCGCTAGCATGAGTCCTGGTCTCCA
        maximum likelihood sequence       GCTAGAAGCATCGTGGTAGATTACGCGCCGTAGCATGATCCTGGTTCCA
        maximum likelihood edit distance  10

evaluating sequence 76 of 100
        observed sequence                 TGGAAGCAGGATCATGCTACGCGCGTAATACTACCAAAGCTTCGTAGCA
        maximum likelihood sequence       TGGAACCAGGATCATGCTACGGCGCGTAATCTACCACGATGCTTCTAGCA
        maximum likelihood edit distance  9

evaluating sequence 77 of 100
        observed sequence                 TGCTAGAAGCATCGTCGGTAGATTACGGCGCCCGTGCATGATCCTAGTGTCCA
        maximum likelihood sequence       TGCTAGAAGCATCGTGGTAGATTACGCGCCGTAGCATGATCCTGGTTCCA
        maximum likelihood edit distance  8

evaluating sequence 78 of 100
        observed sequence                 GCAGAAGCACGTTGGTAGTTCGCGCCGCTAGCATGATCCTGGTTGCCA
        maximum likelihood sequence       GTAGAAGCATCGTGGTAGATTACGCGCCGTAGCATGATCCTGGTTCCA
        maximum likelihood edit distance  9

evaluating sequence 79 of 100
```

```
        observed sequence                 TGCTTGAAGCATCGTGGTAGTCTACGCGCGTCATGATCCTGGTTCCA
        maximum likelihood sequence       TGCTAGAAGCATCGTGGTAGATTACGCGCCGTAGCATGATCCTGGTTCCA
        maximum likelihood edit distance  6

evaluating sequence 80 of 100
        observed sequence                 TGCCTAGAATTCTATCATGGTAGATTACGCGCGTAGCATGGAGTCCTGGTTCTA
        maximum likelihood sequence       TGCTAGAAGCATCGTGGTAGATTACGCGCCGTAGCATGATCCTGGTTCCA
        maximum likelihood edit distance  13

evaluating sequence 81 of 100
        observed sequence                 TGAACCGGATCATGCTACGGGGTAATCTACCAGATGCTTCAGCA
        maximum likelihood sequence       GAACCAGGATCATGCTACGGCGCGTAATCTACCACGATGCTTCTAGCA
        maximum likelihood edit distance  5

evaluating sequence 82 of 100
        observed sequence                 TGGAACCAGGATCATGCTACAGCGCGTTTCTCCCAGATGCTTCTAGCA
        maximum likelihood sequence       TGGAACCAGGATCATGCTACGGCGCGTAATCTACCACGATGCTTCTAGCA
        maximum likelihood edit distance  5

evaluating sequence 83 of 100
        observed sequence                 TGGTAACCAGGATCATGCGTTACGGCGCGTAATCTACCACATGTCTACGCA
        maximum likelihood sequence       GTGGAACCAGGATCATGCTACGGCGCGTAATCTACCACGATGCTTCTAGCA
        maximum likelihood edit distance  9

evaluating sequence 84 of 100
        observed sequence                 TGGAACCAGGATCATGCATACGGCGCGCAATTTACCACGATGCTTTTAGCA
        maximum likelihood sequence       TGGAACCAGGATCATGCTACGGCGCGTAATCTACCACGATGCTTCTAGCA
        maximum likelihood edit distance  5

evaluating sequence 85 of 100
        observed sequence                 TGGACCAGCGATCATGCCAGGCGCGTTAATCTACCTACGATGTTCTAGCA
        maximum likelihood sequence       TGGAACCAGGATCATGCTACGGCGCGTAATCTACCACGATGCTTCTAGCA
        maximum likelihood edit distance  10

evaluating sequence 86 of 100
        observed sequence                 TGCTAGAAGCATCGTGGTCGATTACGCGCCGTTAGGCATGAACCTGTTTCCA
        maximum likelihood sequence       TGCTAGAAGCATCGTGGTAGATTACGCGCCGTAGCATGATCCTGGTTCCA
        maximum likelihood edit distance  5

evaluating sequence 87 of 100
        observed sequence                 TGGAACCAGGATCGATGTATACGGCTGCGTATTCTACCACGATCGCTTCTACA
        maximum likelihood sequence       TGGAACCAGGATCATGCTACGGCGCGTAATCTACCACGATGCTTCTAGCA
        maximum likelihood edit distance  10

evaluating sequence 88 of 100
        observed sequence                 TGCGAACCAGGATACATGCTACGGCGCGTAATCCTACCACGATGCTTCTAGA
        maximum likelihood sequence       TGGAACCAGGATCATGCTACGGCGCGTAATCTACCACGATGCTTCTAGA
        maximum likelihood edit distance  5

evaluating sequence 89 of 100
        observed sequence                 TGCTAGATCATCGTGGCTAGATAAAGCGCCGTAGCATGATCCTGGATTCCA
        maximum likelihood sequence       TGCTAGAAGCATCGTGGTAGATTACGCGCCGTAGCATGATCCTGGTTCCA
        maximum likelihood edit distance  8

evaluating sequence 90 of 100
        observed sequence                 TGGAACCAGATCAGCACGGGGCGTATATCTACCACGATCGCTTTCTAGCA
        maximum likelihood sequence       TGGAACCAGGATCATGCTACGGCGCGTAATCTACCACGATGCTTCTAGCA
        maximum likelihood edit distance  7

evaluating sequence 91 of 100
        observed sequence                 TGGAACCAGGATATGCTACGGCGCGTAATCTACCACGATTGGTCTAGCA
        maximum likelihood sequence       TGGAACCAGGATCATGCTACGGCGCGTAATCTACCACGATGCTTCTAGCA
        maximum likelihood edit distance  4

evaluating sequence 92 of 100
        observed sequence                 TGCTAGAAGCATATCGTGTAGATTACCAGCCGTAGATGTATCCTGGTTCCA
        maximum likelihood sequence       TGCTAGAAGCATCGTGGTAGATTACGCGCCGTAGCATGATCCTGGTTCCA
        maximum likelihood edit distance  8

evaluating sequence 93 of 100
        observed sequence                 TGCTAGAACGCATCGTGGTAGCTTACGCGCCGAGCATGATCCTGGTTCCA
        maximum likelihood sequence       TGCTAGAAGCATCGTGGTAGATTACGCGCCGTAGCATGATCCTGGTTCCA
        maximum likelihood edit distance  4

evaluating sequence 94 of 100
        observed sequence                 TGGAAACAGATCATACAGACGCGTATCTAACCACGAAGCTATCTAGCA
        maximum likelihood sequence       TGGAACCAGGATCATGCTACGGCGCGTAATCTACCACGATGCTTCTAGCA
        maximum likelihood edit distance  11

evaluating sequence 95 of 100
        observed sequence                 TTGCTAAAGCATCGTGGTAATTCGCGCGTAGCATGATCCTGGTTCCCA
        maximum likelihood sequence       TGCTAGAAGCATCGTGGTAGATTACGCGCCGTAGCATGATCCTGGTTCCA
        maximum likelihood edit distance  5

evaluating sequence 96 of 100
        observed sequence                 TGCACCAGGATACATGCTAGCGGCCGCGTAATCTACCACATGCTTCTAGCA
        maximum likelihood sequence       GGAACCAGGATCATGCTACGGCGCGTAATCTACCACGATGCTTCTAGCA
        maximum likelihood edit distance  8

evaluating sequence 97 of 100
        observed sequence                 TTGCTAGAAGCAATCGTGGTAGATTACGCCCGGTAGCATGATCCTGGTTCCA
        maximum likelihood sequence       TTGCTAGAAGCATCGTGGTAGATTACGCGCCGTAGCATGATCCTGGTTCCA
        maximum likelihood edit distance  3

evaluating sequence 98 of 100
        observed sequence                 TGGATACAGGAATCATGCTGCGGCAGCGTATCTACCACGATGCTTCTAGA
        maximum likelihood sequence       TGGAACCAGGATCATGCTACGGCGCGTAATCTACCACGATGCTTCTAGA
        maximum likelihood edit distance  7
```

```
evaluating sequence 99 of 100
        observed sequence                 TGCTAGAAGCATCGTGGTAGATTACGCGCGTAGCATGATCCTGGTTCCCA
        maximum likelihood sequence       TGCTAGAAGCATCGTGGTAGATTACGCGCCGTAGCATGATCCTGGTTCCA
        maximum likelihood edit distance  2

evaluating sequence 100 of 100
        observed sequence                 GTGGAACCAGAGTCATGCTCCGCGCGTAATCTCCCACGTGCTTCTAGA
        maximum likelihood sequence       GTGGAACCAGGATCATGCTACGGCGCGTAATCTACCACGATGCTTCTAGA
        maximum likelihood edit distance  6

DATASET STATISTICS:
        assumed error rate    12.5%
        total bases observed  4984
        total edits accepted  733
        inferred error rate   14.707062600321027%
```

In [33]:

```
canonical_kmers = collect(keys(Eisenia.count_canonical_kmers(maximum_likelihood_observations, k)))
stranded_kmer_graph = Eisenia.build_stranded_kmer_graph(canonical_kmers, maximum_likelihood_observations)
filename = reference_sequence_id * "." * replace(string(Dates.now()), ':' => '.') * ".svg"
Eisenia.plot_stranded_kmer_graph(stranded_kmer_graph, filename=filename)
HTML("""
<image src="$filename" width=50%>
""")
```

Out[33]:

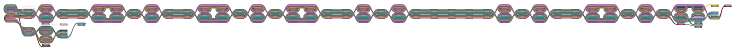

In [34]:

```
maximum_likelihood_observations = Eisenia.viterbi_maximum_likelihood_traversals(stranded_kmer_graph, verbosity="reads");
```

```
computing kmer counts...
computing kmer state likelihoods...
finding shortest paths between kmers...
finding viterbi maximum likelihood paths for observed sequences...

evaluating sequence 1 of 100
        observed sequence                 TGGAACCAGGATCATGCTACGGCGCGTAATCTACCACGATGCTTCTAGCA
        maximum likelihood sequence       TGGAACCAGGATCATGCTACGGCGCGTAATCTACCACGATGCTTCTAGCA
        maximum likelihood edit distance  0

evaluating sequence 2 of 100
        observed sequence                 TGGAACCAGGATCATGCTACGGCGCGTAATCTACCACGATGCTTCTAGCA
        maximum likelihood sequence       TGGAACCAGGATCATGCTACGGCGCGTAATCTACCACGATGCTTCTAGCA
        maximum likelihood edit distance  0

evaluating sequence 3 of 100
        observed sequence                 TGCTAGAAGCATCGTGGTAGATTACGCGCCGTAGCATGATCCTGGTTCCA
        maximum likelihood sequence       TGCTAGAAGCATCGTGGTAGATTACGCGCCGTAGCATGATCCTGGTTCCA
        maximum likelihood edit distance  0

evaluating sequence 4 of 100
        observed sequence                 TGGAACCAGGATCATGCTACGGCGCGTAATCTACCACGATGCTTCTAGCA
        maximum likelihood sequence       TGGAACCAGGATCATGCTACGGCGCGTAATCTACCACGATGCTTCTAGCA
        maximum likelihood edit distance  0

evaluating sequence 5 of 100
        observed sequence                 TGGAACCAGGATCATGCTACGGCGCGTAATCTACCACGATGCTTCTAGCA
        maximum likelihood sequence       TGGAACCAGGATCATGCTACGGCGCGTAATCTACCACGATGCTTCTAGCA
        maximum likelihood edit distance  0

evaluating sequence 6 of 100
        observed sequence                 GGCTAGAAGCATCGTGGTAGATTACGCGCCGTAGCATGATCCTGGTTCCA
        maximum likelihood sequence       TGCTAGAAGCATCGTGGTAGATTACGCGCCGTAGCATGATCCTGGTTCCA
        maximum likelihood edit distance  1

evaluating sequence 7 of 100
        observed sequence                 TTGCTAGAAGCATCGTGGTAGATTACGCGCCGTAGCATGATCCTGGTTCCA
        maximum likelihood sequence       TTGCTAGAAGCATCGTGGTAGATTACGCGCCGTAGCATGATCCTGGTTCCA
        maximum likelihood edit distance  0

evaluating sequence 8 of 100
        observed sequence                 TGGAACCAGGATCATGCTACGGCGCGTAATCTACCACGATGCTTCTAGCA
        maximum likelihood sequence       TGGAACCAGGATCATGCTACGGCGCGTAATCTACCACGATGCTTCTAGCA
        maximum likelihood edit distance  0

evaluating sequence 9 of 100
        observed sequence                 TGCTAGAAGCATCGTGGTAGATTACGCGCCGTAGCATGATCCTGGTTCCA
        maximum likelihood sequence       TGCTAGAAGCATCGTGGTAGATTACGCGCCGTAGCATGATCCTGGTTCCA
        maximum likelihood edit distance  0

evaluating sequence 10 of 100
        observed sequence                 TGCTAGAAGCATCGTGGTAGATTACGCGCCGTAGCATGATCCTGGTTCCA
        maximum likelihood sequence       TGCTAGAAGCATCGTGGTAGATTACGCGCCGTAGCATGATCCTGGTTCCA
        maximum likelihood edit distance  0

evaluating sequence 11 of 100
        observed sequence                 TGGAACCAGGATCATGCTACGGCGCGTAATCTACCACGATGCTTCTAGCA
        maximum likelihood sequence       TGGAACCAGGATCATGCTACGGCGCGTAATCTACCACGATGCTTCTAGCA
        maximum likelihood edit distance  0

evaluating sequence 12 of 100
        observed sequence                 TGGAACCAGGATCATGCTACGGCGCGTAATCTACCACGATGCTTCTAGCA
        maximum likelihood sequence       TGGAACCAGGATCATGCTACGGCGCGTAATCTACCACGATGCTTCTAGCA
        maximum likelihood edit distance  0

evaluating sequence 13 of 100
        observed sequence                 TGCTAGAAGCATCGTGGTAGATTACGCGCCGTAGCATGATCCTGGTTCA
        maximum likelihood sequence       TGCTAGAAGCATCGTGGTAGATTACGCGCCGTAGCATGATCCTGGTTCC
        maximum likelihood edit distance  1

evaluating sequence 14 of 100
        observed sequence                 TGCTAGAAGCATCGTGGTAGATTACGCGCCGTAGCATGATCCTGGTTCCA
        maximum likelihood sequence       TGCTAGAAGCATCGTGGTAGATTACGCGCCGTAGCATGATCCTGGTTCCA
        maximum likelihood edit distance  0

evaluating sequence 15 of 100
        observed sequence                 GACCAGGATCATGCTACGGCGCGTAATCTACCACGATGCTTCTAGCA
        maximum likelihood sequence       AACCAGGATCATGCTACGGCGCGTAATCTACCACGATGCTTCTAGCA
        maximum likelihood edit distance  1

evaluating sequence 16 of 100
        observed sequence                 TGCTAGAAGCATCGTGGTAGATTACGCGCCGTAGCATGATCCTGGTTCCA
        maximum likelihood sequence       TGCTAGAAGCATCGTGGTAGATTACGCGCCGTAGCATGATCCTGGTTCCA
        maximum likelihood edit distance  0

evaluating sequence 17 of 100
        observed sequence                 TGCTAGAAGCATCGTGGTAGATTACGCGCCGTAGCATGATCCTGGTTCA
        maximum likelihood sequence       TGCTAGAAGCATCGTGGTAGATTACGCGCCGTAGCATGATCCTGGTTCC
        maximum likelihood edit distance  1

evaluating sequence 18 of 100
        observed sequence                 TGCTAGAAGCATCGTGGTAGATTACGCGCCGTAGCATGATCCTGGTTCCA
        maximum likelihood sequence       TGCTAGAAGCATCGTGGTAGATTACGCGCCGTAGCATGATCCTGGTTCCA
        maximum likelihood edit distance  0

evaluating sequence 19 of 100
        observed sequence                 GCTAGAAGCATCGTGGTAGATTACGCGCCGTAGCATGATCCTGGTTCCA
        maximum likelihood sequence       GCTAGAAGCATCGTGGTAGATTACGCGCCGTAGCATGATCCTGGTTCCA
        maximum likelihood edit distance  0
```

```
evaluating sequence 20 of 100
        observed sequence                TGTAGAAGCATCGTGGTAGATTACGCGCCGTAGCATGATCCTGGTTCCA
        maximum likelihood sequence      GCTAGAAGCATCGTGGTAGATTACGCGCCGTAGCATGATCCTGGTTCCA
        maximum likelihood edit distance  2

evaluating sequence 21 of 100
        observed sequence                TGGAACCAGGATCATGCTACGGCGCGTAATCTACCACGATGCTTCTAGCA
        maximum likelihood sequence      TGGAACCAGGATCATGCTACGGCGCGTAATCTACCACGATGCTTCTAGCA
        maximum likelihood edit distance  0

evaluating sequence 22 of 100
        observed sequence                TGGAACCAGGATCATGCTACGGCGCGTAATCTACCACGATGCTTCTAGCA
        maximum likelihood sequence      TGGAACCAGGATCATGCTACGGCGCGTAATCTACCACGATGCTTCTAGCA
        maximum likelihood edit distance  0

evaluating sequence 23 of 100
        observed sequence                GCTAGAAGCATCGTGGTAGATTACGCGCCGTAGCATGATCCTGGTTCCA
        maximum likelihood sequence      GCTAGAAGCATCGTGGTAGATTACGCGCCGTAGCATGATCCTGGTTCCA
        maximum likelihood edit distance  0

evaluating sequence 24 of 100
        observed sequence                TGGAACCAGGATCATGCTACGGCGCGTAATCTACCACGATGCTTCTAGCA
        maximum likelihood sequence      TGGAACCAGGATCATGCTACGGCGCGTAATCTACCACGATGCTTCTAGCA
        maximum likelihood edit distance  0

evaluating sequence 25 of 100
        observed sequence                TGCTAGAAGCATCGTGGTAGATTACGCGCCGTAGCATGATCCTGGTTCCA
        maximum likelihood sequence      TGCTAGAAGCATCGTGGTAGATTACGCGCCGTAGCATGATCCTGGTTCCA
        maximum likelihood edit distance  0

evaluating sequence 26 of 100
        observed sequence                TGGAACCAGGATCATGCTACGGCGCGTAATCTACCACGATGCTTCTAGCA
        maximum likelihood sequence      TGGAACCAGGATCATGCTACGGCGCGTAATCTACCACGATGCTTCTAGCA
        maximum likelihood edit distance  0

evaluating sequence 27 of 100
        observed sequence                TGGAACCAGGATCATGCTACGGCGCGTAATCTACCACGATGCTTCTAGCA
        maximum likelihood sequence      TGGAACCAGGATCATGCTACGGCGCGTAATCTACCACGATGCTTCTAGCA
        maximum likelihood edit distance  0

evaluating sequence 28 of 100
        observed sequence                TGCTAGAAGCATCGTGGTAGATTACGCGCCGTAGCATGATCCTGGTTCCA
        maximum likelihood sequence      TGCTAGAAGCATCGTGGTAGATTACGCGCCGTAGCATGATCCTGGTTCCA
        maximum likelihood edit distance  0

evaluating sequence 29 of 100
        observed sequence                TGCTAGAAGCATCGTGGTAGATTACGCGCCGTAGCATGATCCTGGTTCCA
        maximum likelihood sequence      TGCTAGAAGCATCGTGGTAGATTACGCGCCGTAGCATGATCCTGGTTCCA
        maximum likelihood edit distance  0

evaluating sequence 30 of 100
        observed sequence                CTAGAAGCATCGTGGTAGATTACGCGCCGTAGCATGATCCTGGTTCCA
        maximum likelihood sequence      CTAGAAGCATCGTGGTAGATTACGCGCCGTAGCATGATCCTGGTTCCA
        maximum likelihood edit distance  0

evaluating sequence 31 of 100
        observed sequence                TGGAACCAGGATCATGCTACGGCGCGTAATCTACCACGATGCTTCTAGCA
        maximum likelihood sequence      TGGAACCAGGATCATGCTACGGCGCGTAATCTACCACGATGCTTCTAGCA
        maximum likelihood edit distance  0

evaluating sequence 32 of 100
        observed sequence                TGGAACCAGGATCATGCTACGGCGCGTAATCTACCACGATGCTTCTAGCA
        maximum likelihood sequence      TGGAACCAGGATCATGCTACGGCGCGTAATCTACCACGATGCTTCTAGCA
        maximum likelihood edit distance  0

evaluating sequence 33 of 100
        observed sequence                TGGAACCAGGATCATGCTACGGCGCGTAATCTACCACGATGCTTCTAGCA
        maximum likelihood sequence      TGGAACCAGGATCATGCTACGGCGCGTAATCTACCACGATGCTTCTAGCA
        maximum likelihood edit distance  0

evaluating sequence 34 of 100
        observed sequence                TGGAACCAGGATCATGCTACGGCGCGTAATCTACCACGATGCTTCTAGCA
        maximum likelihood sequence      TGGAACCAGGATCATGCTACGGCGCGTAATCTACCACGATGCTTCTAGCA
        maximum likelihood edit distance  0

evaluating sequence 35 of 100
        observed sequence                TGCTAGAAGCATCGTGGTAGATTACGCGCCGTAGCATGATCCTGGTTCCA
        maximum likelihood sequence      TGCTAGAAGCATCGTGGTAGATTACGCGCCGTAGCATGATCCTGGTTCCA
        maximum likelihood edit distance  0

evaluating sequence 36 of 100
        observed sequence                TGGAACCAGGATCATGCTACGGCGCGTAATCTACCACGATGCTTCTAGCA
        maximum likelihood sequence      TGGAACCAGGATCATGCTACGGCGCGTAATCTACCACGATGCTTCTAGCA
        maximum likelihood edit distance  0

evaluating sequence 37 of 100
        observed sequence                TGGAACCAGGATCATGCTACGGCGCGTAATCTACCACGATGCTTCTAGCA
        maximum likelihood sequence      TGGAACCAGGATCATGCTACGGCGCGTAATCTACCACGATGCTTCTAGCA
        maximum likelihood edit distance  0

evaluating sequence 38 of 100
        observed sequence                TGGAACCAGGATCATGCTACGGCGCGTAATCTACCACGATGCTTCTAGCA
        maximum likelihood sequence      TGGAACCAGGATCATGCTACGGCGCGTAATCTACCACGATGCTTCTAGCA
        maximum likelihood edit distance  0

evaluating sequence 39 of 100
        observed sequence                GCTAGAAGCATCGTGGTAGATTACGCGCCGTAGCATGATCCTGGTTCCA
        maximum likelihood sequence      GCTAGAAGCATCGTGGTAGATTACGCGCCGTAGCATGATCCTGGTTCCA
```

```
        maximum likelihood edit distance  0

evaluating sequence 40 of 100
        observed sequence                 TGGAACCAGGATCATGCTACGGCGCGTAATCTACCACGATGCTTCTAGCA
        maximum likelihood sequence       TGGAACCAGGATCATGCTACGGCGCGTAATCTACCACGATGCTTCTAGCA
        maximum likelihood edit distance  0

evaluating sequence 41 of 100
        observed sequence                 TGGAACCAGGATCATGCTACGGCGCGTAATCTACCACGATGCTTCTAGCA
        maximum likelihood sequence       TGGAACCAGGATCATGCTACGGCGCGTAATCTACCACGATGCTTCTAGCA
        maximum likelihood edit distance  0

evaluating sequence 42 of 100
        observed sequence                 GAACCAGGATCATGCTACGGCGCGTAATCTACCACGATGCTTCTAGC
        maximum likelihood sequence       GAACCAGGATCATGCTACGGCGCGTAATCTACCACGATGCTTCTAGC
        maximum likelihood edit distance  0

evaluating sequence 43 of 100
        observed sequence                 TGGAACCAGGATCATGCTACGGCGCGTAATCTACCACGATGCTTCTAGCA
        maximum likelihood sequence       TGGAACCAGGATCATGCTACGGCGCGTAATCTACCACGATGCTTCTAGCA
        maximum likelihood edit distance  0

evaluating sequence 44 of 100
        observed sequence                 TGCTAGAAGCATCGTGGTAGATTACGCGCCGTAGCATGATCCTGGTTCCA
        maximum likelihood sequence       TGCTAGAAGCATCGTGGTAGATTACGCGCCGTAGCATGATCCTGGTTCCA
        maximum likelihood edit distance  0

evaluating sequence 45 of 100
        observed sequence                 GCTAGAAGCATCGTGGTAGATTACGCGCCGTAGCATGATCCTGGTTCCA
        maximum likelihood sequence       GCTAGAAGCATCGTGGTAGATTACGCGCCGTAGCATGATCCTGGTTCCA
        maximum likelihood edit distance  0

evaluating sequence 46 of 100
        observed sequence                 TGGAACCAGGATCATGCTACGGCGCGTAATCTACCACGATGCTTCTAGCA
        maximum likelihood sequence       TGGAACCAGGATCATGCTACGGCGCGTAATCTACCACGATGCTTCTAGCA
        maximum likelihood edit distance  0

evaluating sequence 47 of 100
        observed sequence                 GGAACCAGGATCATGCTACGGCGCGTAATCTACCACGATGCTTCTAGCA
        maximum likelihood sequence       GGAACCAGGATCATGCTACGGCGCGTAATCTACCACGATGCTTCTAGCA
        maximum likelihood edit distance  0

evaluating sequence 48 of 100
        observed sequence                 TGCTAGAAGCATCGTGGTAGATTACGCGCCGTAGCATGATCCTGGTTCCA
        maximum likelihood sequence       TGCTAGAAGCATCGTGGTAGATTACGCGCCGTAGCATGATCCTGGTTCCA
        maximum likelihood edit distance  0

evaluating sequence 49 of 100
        observed sequence                 GCTAGAAGCATCGTGGTAGATTACGCGCCGTAGCATGATCCTGGTTCA
        maximum likelihood sequence       GCTAGAAGCATCGTGGTAGATTACGCGCCGTAGCATGATCCTGGTTCC
        maximum likelihood edit distance  1

evaluating sequence 50 of 100
        observed sequence                 TGGAACCAGGATCATGCTACGGCGCGTAATCTACCACGATGCTTCTAGCA
        maximum likelihood sequence       TGGAACCAGGATCATGCTACGGCGCGTAATCTACCACGATGCTTCTAGCA
        maximum likelihood edit distance  0

evaluating sequence 51 of 100
        observed sequence                 TGCTAGAAGCATCGTGGTAGATTACGCGCCGTAGCATGATCCTGGTTCC
        maximum likelihood sequence       TGCTAGAAGCATCGTGGTAGATTACGCGCCGTAGCATGATCCTGGTTCC
        maximum likelihood edit distance  0

evaluating sequence 52 of 100
        observed sequence                 GTAGAAGCATCGTGGTAGATTACGCGCCGTAGCATGATCCTGGTTCCA
        maximum likelihood sequence       CTAGAAGCATCGTGGTAGATTACGCGCCGTAGCATGATCCTGGTTCCA
        maximum likelihood edit distance  1

evaluating sequence 53 of 100
        observed sequence                 TGGAACCAGGATCATGCTACGGCGCGTAATCTACCACGATGCTTCTAGA
        maximum likelihood sequence       TGGAACCAGGATCATGCTACGGCGCGTAATCTACCACGATGCTTCTAGC
        maximum likelihood edit distance  1

evaluating sequence 54 of 100
        observed sequence                 TGGAACCAGGATCATGCTACGGCGCGTAATCTACCACGATGCTTCTAGCA
        maximum likelihood sequence       TGGAACCAGGATCATGCTACGGCGCGTAATCTACCACGATGCTTCTAGCA
        maximum likelihood edit distance  0

evaluating sequence 55 of 100
        observed sequence                 GTGGAACCAGGATCATGCTACGGCGCGTAATCTACCACGATGCTTCTAGCA
        maximum likelihood sequence       GTGGAACCAGGATCATGCTACGGCGCGTAATCTACCACGATGCTTCTAGCA
        maximum likelihood edit distance  0

evaluating sequence 56 of 100
        observed sequence                 TGGAACCAGGATCATGCTACGGCGCGTAATCTACCACGATGCTTCTAGCA
        maximum likelihood sequence       TGGAACCAGGATCATGCTACGGCGCGTAATCTACCACGATGCTTCTAGCA
        maximum likelihood edit distance  0

evaluating sequence 57 of 100
        observed sequence                 TGCTAGAAGCATCGTGGTAGATTACGCGCCGTAGCATGATCCTGGTTCCA
        maximum likelihood sequence       TGCTAGAAGCATCGTGGTAGATTACGCGCCGTAGCATGATCCTGGTTCCA
        maximum likelihood edit distance  0

evaluating sequence 58 of 100
        observed sequence                 TGCTAGAAGCATCGTGGTAGATTACGCGCCGTAGCATGATCCTGGTTCCA
        maximum likelihood sequence       TGCTAGAAGCATCGTGGTAGATTACGCGCCGTAGCATGATCCTGGTTCCA
        maximum likelihood edit distance  0

evaluating sequence 59 of 100
        observed sequence                 TGGAACCAGGATCATGCTACGGCGCGTAATCTACCACGATGCTTCTAGCA
```

```
        maximum likelihood sequence       TGGAACCAGGATCATGCTACGGCGCGTAATCTACCACGATGCTTCTAGCA
        maximum likelihood edit distance  0

evaluating sequence 60 of 100
        observed sequence                 TGCTAGAAGCATCGTGGTAGATTACGCGCCGTAGCATGATCCTGGTTCCA
        maximum likelihood sequence       TGCTAGAAGCATCGTGGTAGATTACGCGCCGTAGCATGATCCTGGTTCCA
        maximum likelihood edit distance  0

evaluating sequence 61 of 100
        observed sequence                 GCTAGAAGCATCGTGGTAGATTACGCGCCGTAGCATGATCCTGGTTCCA
        maximum likelihood sequence       GCTAGAAGCATCGTGGTAGATTACGCGCCGTAGCATGATCCTGGTTCCA
        maximum likelihood edit distance  0

evaluating sequence 62 of 100
        observed sequence                 TGACCAGGATCATGCTACGGCGCGTAATCTACCACGATGCTTCTAGCA
        maximum likelihood sequence       GAACCAGGATCATGCTACGGCGCGTAATCTACCACGATGCTTCTAGCA
        maximum likelihood edit distance  2

evaluating sequence 63 of 100
        observed sequence                 GGAACCAGGATCATGCTACGGCGCGTAATCTACCACGATGCTTCTAGA
        maximum likelihood sequence       GGAACCAGGATCATGCTACGGCGCGTAATCTACCACGATGCTTCTAGC
        maximum likelihood edit distance  1

evaluating sequence 64 of 100
        observed sequence                 TGGAACCAGGATCATGCTACGGCGCGTAATCTACCACGATGCTTCTAGCA
        maximum likelihood sequence       TGGAACCAGGATCATGCTACGGCGCGTAATCTACCACGATGCTTCTAGCA
        maximum likelihood edit distance  0

evaluating sequence 65 of 100
        observed sequence                 GTGGAACCAGGATCATGCTACGGCGCGTAATCTACCACGATGCTTCTAGCA
        maximum likelihood sequence       GTGGAACCAGGATCATGCTACGGCGCGTAATCTACCACGATGCTTCTAGCA
        maximum likelihood edit distance  0

evaluating sequence 66 of 100
        observed sequence                 TGGAACCAGGATCATGCTACGGCGCGTAATCTACCACGATGCTTCTAGCA
        maximum likelihood sequence       TGGAACCAGGATCATGCTACGGCGCGTAATCTACCACGATGCTTCTAGCA
        maximum likelihood edit distance  0

evaluating sequence 67 of 100
        observed sequence                 TGGAACCAGGATCATGCTACGGCGCGTAATCTACCACGATGCTTCTAGA
        maximum likelihood sequence       TGGAACCAGGATCATGCTACGGCGCGTAATCTACCACGATGCTTCTAGC
        maximum likelihood edit distance  1

evaluating sequence 68 of 100
        observed sequence                 GCTAGAAGCATCGTGGTAGATTACGCGCCGTAGCATGATCCTGGTTCCA
        maximum likelihood sequence       GCTAGAAGCATCGTGGTAGATTACGCGCCGTAGCATGATCCTGGTTCCA
        maximum likelihood edit distance  0

evaluating sequence 69 of 100
        observed sequence                 TGCTAGAAGCATCGTGGTAGATTACGCGCCGTAGCATGATCCTGGTTCCA
        maximum likelihood sequence       TGCTAGAAGCATCGTGGTAGATTACGCGCCGTAGCATGATCCTGGTTCCA
        maximum likelihood edit distance  0

evaluating sequence 70 of 100
        observed sequence                 TGGAACCAGGATCATGCTACGGCGCGTAATCTACCACGATGCTTCTAGCA
        maximum likelihood sequence       TGGAACCAGGATCATGCTACGGCGCGTAATCTACCACGATGCTTCTAGCA
        maximum likelihood edit distance  0

evaluating sequence 71 of 100
        observed sequence                 TTGCTAGAAGCATCGTGGTAGATTACGCGCCGTAGCATGATCCTGGTTCCA
        maximum likelihood sequence       TTGCTAGAAGCATCGTGGTAGATTACGCGCCGTAGCATGATCCTGGTTCCA
        maximum likelihood edit distance  0

evaluating sequence 72 of 100
        observed sequence                 GGAACCAGGATCATGCTACGGCGCGTAATCTACCACGATGCTTCTAGC
        maximum likelihood sequence       GGAACCAGGATCATGCTACGGCGCGTAATCTACCACGATGCTTCTAGC
        maximum likelihood edit distance  0

evaluating sequence 73 of 100
        observed sequence                 TGCTAGAAGCATCGTGGTAGATTACGCGCCGTAGCATGATCCTGGTTCCA
        maximum likelihood sequence       TGCTAGAAGCATCGTGGTAGATTACGCGCCGTAGCATGATCCTGGTTCCA
        maximum likelihood edit distance  0

evaluating sequence 74 of 100
        observed sequence                 TGCTAGAAGCATCGTGGTAGATTACGCGCCGTAGCATGATCCTGGTTCA
        maximum likelihood sequence       TGCTAGAAGCATCGTGGTAGATTACGCGCCGTAGCATGATCCTGGTTCC
        maximum likelihood edit distance  1

evaluating sequence 75 of 100
        observed sequence                 GCTAGAAGCATCGTGGTAGATTACGCGCCGTAGCATGATCCTGGTTCCA
        maximum likelihood sequence       GCTAGAAGCATCGTGGTAGATTACGCGCCGTAGCATGATCCTGGTTCCA
        maximum likelihood edit distance  0

evaluating sequence 76 of 100
        observed sequence                 TGGAACCAGGATCATGCTACGGCGCGTAATCTACCACGATGCTTCTAGCA
        maximum likelihood sequence       TGGAACCAGGATCATGCTACGGCGCGTAATCTACCACGATGCTTCTAGCA
        maximum likelihood edit distance  0

evaluating sequence 77 of 100
        observed sequence                 TGCTAGAAGCATCGTGGTAGATTACGCGCCGTAGCATGATCCTGGTTCCA
        maximum likelihood sequence       TGCTAGAAGCATCGTGGTAGATTACGCGCCGTAGCATGATCCTGGTTCCA
        maximum likelihood edit distance  0

evaluating sequence 78 of 100
        observed sequence                 GTAGAAGCATCGTGGTAGATTACGCGCCGTAGCATGATCCTGGTTCCA
        maximum likelihood sequence       CTAGAAGCATCGTGGTAGATTACGCGCCGTAGCATGATCCTGGTTCCA
        maximum likelihood edit distance  1

evaluating sequence 79 of 100
```

```
        observed sequence                 TGCTAGAAGCATCGTGGTAGATTACGCGCCGTAGCATGATCCTGGTTCCA
        maximum likelihood sequence       TGCTAGAAGCATCGTGGTAGATTACGCGCCGTAGCATGATCCTGGTTCCA
        maximum likelihood edit distance  0

evaluating sequence 80 of 100
        observed sequence                 TGCTAGAAGCATCGTGGTAGATTACGCGCCGTAGCATGATCCTGGTTCCA
        maximum likelihood sequence       TGCTAGAAGCATCGTGGTAGATTACGCGCCGTAGCATGATCCTGGTTCCA
        maximum likelihood edit distance  0

evaluating sequence 81 of 100
        observed sequence                 GAACCAGGATCATGCTACGGCGCGTAATCTACCACGATGCTTCTAGCA
        maximum likelihood sequence       GAACCAGGATCATGCTACGGCGCGTAATCTACCACGATGCTTCTAGCA
        maximum likelihood edit distance  0

evaluating sequence 82 of 100
        observed sequence                 TGGAACCAGGATCATGCTACGGCGCGTAATCTACCACGATGCTTCTAGCA
        maximum likelihood sequence       TGGAACCAGGATCATGCTACGGCGCGTAATCTACCACGATGCTTCTAGCA
        maximum likelihood edit distance  0

evaluating sequence 83 of 100
        observed sequence                 GTGGAACCAGGATCATGCTACGGCGCGTAATCTACCACGATGCTTCTAGCA
        maximum likelihood sequence       GTGGAACCAGGATCATGCTACGGCGCGTAATCTACCACGATGCTTCTAGCA
        maximum likelihood edit distance  0

evaluating sequence 84 of 100
        observed sequence                 TGGAACCAGGATCATGCTACGGCGCGTAATCTACCACGATGCTTCTAGCA
        maximum likelihood sequence       TGGAACCAGGATCATGCTACGGCGCGTAATCTACCACGATGCTTCTAGCA
        maximum likelihood edit distance  0

evaluating sequence 85 of 100
        observed sequence                 TGGAACCAGGATCATGCTACGGCGCGTAATCTACCACGATGCTTCTAGCA
        maximum likelihood sequence       TGGAACCAGGATCATGCTACGGCGCGTAATCTACCACGATGCTTCTAGCA
        maximum likelihood edit distance  0

evaluating sequence 86 of 100
        observed sequence                 TGCTAGAAGCATCGTGGTAGATTACGCGCCGTAGCATGATCCTGGTTCCA
        maximum likelihood sequence       TGCTAGAAGCATCGTGGTAGATTACGCGCCGTAGCATGATCCTGGTTCCA
        maximum likelihood edit distance  0

evaluating sequence 87 of 100
        observed sequence                 TGGAACCAGGATCATGCTACGGCGCGTAATCTACCACGATGCTTCTAGCA
        maximum likelihood sequence       TGGAACCAGGATCATGCTACGGCGCGTAATCTACCACGATGCTTCTAGCA
        maximum likelihood edit distance  0

evaluating sequence 88 of 100
        observed sequence                 TGGAACCAGGATCATGCTACGGCGCGTAATCTACCACGATGCTTCTAGA
        maximum likelihood sequence       TGGAACCAGGATCATGCTACGGCGCGTAATCTACCACGATGCTTCTAGC
        maximum likelihood edit distance  1

evaluating sequence 89 of 100
        observed sequence                 TGCTAGAAGCATCGTGGTAGATTACGCGCCGTAGCATGATCCTGGTTCCA
        maximum likelihood sequence       TGCTAGAAGCATCGTGGTAGATTACGCGCCGTAGCATGATCCTGGTTCCA
        maximum likelihood edit distance  0

evaluating sequence 90 of 100
        observed sequence                 TGGAACCAGGATCATGCTACGGCGCGTAATCTACCACGATGCTTCTAGCA
        maximum likelihood sequence       TGGAACCAGGATCATGCTACGGCGCGTAATCTACCACGATGCTTCTAGCA
        maximum likelihood edit distance  0

evaluating sequence 91 of 100
        observed sequence                 TGGAACCAGGATCATGCTACGGCGCGTAATCTACCACGATGCTTCTAGCA
        maximum likelihood sequence       TGGAACCAGGATCATGCTACGGCGCGTAATCTACCACGATGCTTCTAGCA
        maximum likelihood edit distance  0

evaluating sequence 92 of 100
        observed sequence                 TGCTAGAAGCATCGTGGTAGATTACGCGCCGTAGCATGATCCTGGTTCCA
        maximum likelihood sequence       TGCTAGAAGCATCGTGGTAGATTACGCGCCGTAGCATGATCCTGGTTCCA
        maximum likelihood edit distance  0

evaluating sequence 93 of 100
        observed sequence                 TGCTAGAAGCATCGTGGTAGATTACGCGCCGTAGCATGATCCTGGTTCCA
        maximum likelihood sequence       TGCTAGAAGCATCGTGGTAGATTACGCGCCGTAGCATGATCCTGGTTCCA
        maximum likelihood edit distance  0

evaluating sequence 94 of 100
        observed sequence                 TGGAACCAGGATCATGCTACGGCGCGTAATCTACCACGATGCTTCTAGCA
        maximum likelihood sequence       TGGAACCAGGATCATGCTACGGCGCGTAATCTACCACGATGCTTCTAGCA
        maximum likelihood edit distance  0

evaluating sequence 95 of 100
        observed sequence                 TGCTAGAAGCATCGTGGTAGATTACGCGCCGTAGCATGATCCTGGTTCCA
        maximum likelihood sequence       TGCTAGAAGCATCGTGGTAGATTACGCGCCGTAGCATGATCCTGGTTCCA
        maximum likelihood edit distance  0

evaluating sequence 96 of 100
        observed sequence                 GGAACCAGGATCATGCTACGGCGCGTAATCTACCACGATGCTTCTAGCA
        maximum likelihood sequence       GGAACCAGGATCATGCTACGGCGCGTAATCTACCACGATGCTTCTAGCA
        maximum likelihood edit distance  0

evaluating sequence 97 of 100
        observed sequence                 TTGCTAGAAGCATCGTGGTAGATTACGCGCCGTAGCATGATCCTGGTTCCA
        maximum likelihood sequence       TTGCTAGAAGCATCGTGGTAGATTACGCGCCGTAGCATGATCCTGGTTCCA
        maximum likelihood edit distance  0

evaluating sequence 98 of 100
        observed sequence                 TGGAACCAGGATCATGCTACGGCGCGTAATCTACCACGATGCTTCTAGA
        maximum likelihood sequence       TGGAACCAGGATCATGCTACGGCGCGTAATCTACCACGATGCTTCTAGC
        maximum likelihood edit distance  1
```

```
evaluating sequence 99 of 100
        observed sequence                 TGCTAGAAGCATCGTGGTAGATTACGCGCCGTAGCATGATCCTGGTTCCA
        maximum likelihood sequence       TGCTAGAAGCATCGTGGTAGATTACGCGCCGTAGCATGATCCTGGTTCCA
        maximum likelihood edit distance  0

evaluating sequence 100 of 100
        observed sequence                 GTGGAACCAGGATCATGCTACGGCGCGTAATCTACCACGATGCTTCTAGA
        maximum likelihood sequence       GTGGAACCAGGATCATGCTACGGCGCGTAATCTACCACGATGCTTCTAGC
        maximum likelihood edit distance  1

DATASET STATISTICS:
        assumed error rate    12.5%
        total bases observed  4966
        total edits accepted  18
        inferred error rate   0.3624647603705195%
```

In [35]:

```
canonical_kmers = collect(keys(Eisenia.count_canonical_kmers(maximum_likelihood_observations, k)))
stranded_kmer_graph = Eisenia.build_stranded_kmer_graph(canonical_kmers, maximum_likelihood_observations)
filename = reference_sequence_id * "." * replace(string(Dates.now()), ':' => '.') * ".svg"
Eisenia.plot_stranded_kmer_graph(stranded_kmer_graph, filename=filename)
HTML("""
<image src="$filename" width=50%>
""")
```

Out[35]:

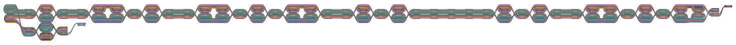

In [36]:

```
maximum_likelihood_observations = Eisenia.viterbi_maximum_likelihood_traversals(stranded_kmer_graph, verbosity="reads");
```

```
computing kmer counts...
computing kmer state likelihoods...
finding shortest paths between kmers...
finding viterbi maximum likelihood paths for observed sequences...

evaluating sequence 1 of 100
        observed sequence                 TGGAACCAGGATCATGCTACGGCGCGTAATCTACCACGATGCTTCTAGCA
        maximum likelihood sequence       TGGAACCAGGATCATGCTACGGCGCGTAATCTACCACGATGCTTCTAGCA
        maximum likelihood edit distance  0

evaluating sequence 2 of 100
        observed sequence                 TGGAACCAGGATCATGCTACGGCGCGTAATCTACCACGATGCTTCTAGCA
        maximum likelihood sequence       TGGAACCAGGATCATGCTACGGCGCGTAATCTACCACGATGCTTCTAGCA
        maximum likelihood edit distance  0

evaluating sequence 3 of 100
        observed sequence                 TGCTAGAAGCATCGTGGTAGATTACGCGCCGTAGCATGATCCTGGTTCCA
        maximum likelihood sequence       TGCTAGAAGCATCGTGGTAGATTACGCGCCGTAGCATGATCCTGGTTCCA
        maximum likelihood edit distance  0

evaluating sequence 4 of 100
        observed sequence                 TGGAACCAGGATCATGCTACGGCGCGTAATCTACCACGATGCTTCTAGCA
        maximum likelihood sequence       TGGAACCAGGATCATGCTACGGCGCGTAATCTACCACGATGCTTCTAGCA
        maximum likelihood edit distance  0

evaluating sequence 5 of 100
        observed sequence                 TGGAACCAGGATCATGCTACGGCGCGTAATCTACCACGATGCTTCTAGCA
        maximum likelihood sequence       TGGAACCAGGATCATGCTACGGCGCGTAATCTACCACGATGCTTCTAGCA
        maximum likelihood edit distance  0

evaluating sequence 6 of 100
        observed sequence                 TGCTAGAAGCATCGTGGTAGATTACGCGCCGTAGCATGATCCTGGTTCCA
        maximum likelihood sequence       TGCTAGAAGCATCGTGGTAGATTACGCGCCGTAGCATGATCCTGGTTCCA
        maximum likelihood edit distance  0

evaluating sequence 7 of 100
        observed sequence                 TTGCTAGAAGCATCGTGGTAGATTACGCGCCGTAGCATGATCCTGGTTCCA
        maximum likelihood sequence       TTGCTAGAAGCATCGTGGTAGATTACGCGCCGTAGCATGATCCTGGTTCCA
        maximum likelihood edit distance  0

evaluating sequence 8 of 100
        observed sequence                 TGGAACCAGGATCATGCTACGGCGCGTAATCTACCACGATGCTTCTAGCA
        maximum likelihood sequence       TGGAACCAGGATCATGCTACGGCGCGTAATCTACCACGATGCTTCTAGCA
        maximum likelihood edit distance  0

evaluating sequence 9 of 100
        observed sequence                 TGCTAGAAGCATCGTGGTAGATTACGCGCCGTAGCATGATCCTGGTTCCA
        maximum likelihood sequence       TGCTAGAAGCATCGTGGTAGATTACGCGCCGTAGCATGATCCTGGTTCCA
        maximum likelihood edit distance  0

evaluating sequence 10 of 100
        observed sequence                 TGCTAGAAGCATCGTGGTAGATTACGCGCCGTAGCATGATCCTGGTTCCA
        maximum likelihood sequence       TGCTAGAAGCATCGTGGTAGATTACGCGCCGTAGCATGATCCTGGTTCCA
        maximum likelihood edit distance  0

evaluating sequence 11 of 100
        observed sequence                 TGGAACCAGGATCATGCTACGGCGCGTAATCTACCACGATGCTTCTAGCA
        maximum likelihood sequence       TGGAACCAGGATCATGCTACGGCGCGTAATCTACCACGATGCTTCTAGCA
        maximum likelihood edit distance  0

evaluating sequence 12 of 100
        observed sequence                 TGGAACCAGGATCATGCTACGGCGCGTAATCTACCACGATGCTTCTAGCA
        maximum likelihood sequence       TGGAACCAGGATCATGCTACGGCGCGTAATCTACCACGATGCTTCTAGCA
        maximum likelihood edit distance  0

evaluating sequence 13 of 100
        observed sequence                 TGCTAGAAGCATCGTGGTAGATTACGCGCCGTAGCATGATCCTGGTTCC
        maximum likelihood sequence       TGCTAGAAGCATCGTGGTAGATTACGCGCCGTAGCATGATCCTGGTTCC
        maximum likelihood edit distance  0

evaluating sequence 14 of 100
        observed sequence                 TGCTAGAAGCATCGTGGTAGATTACGCGCCGTAGCATGATCCTGGTTCCA
        maximum likelihood sequence       TGCTAGAAGCATCGTGGTAGATTACGCGCCGTAGCATGATCCTGGTTCCA
        maximum likelihood edit distance  0

evaluating sequence 15 of 100
        observed sequence                 AACCAGGATCATGCTACGGCGCGTAATCTACCACGATGCTTCTAGCA
        maximum likelihood sequence       AACCAGGATCATGCTACGGCGCGTAATCTACCACGATGCTTCTAGCA
        maximum likelihood edit distance  0

evaluating sequence 16 of 100
        observed sequence                 TGCTAGAAGCATCGTGGTAGATTACGCGCCGTAGCATGATCCTGGTTCCA
        maximum likelihood sequence       TGCTAGAAGCATCGTGGTAGATTACGCGCCGTAGCATGATCCTGGTTCCA
        maximum likelihood edit distance  0

evaluating sequence 17 of 100
        observed sequence                 TGCTAGAAGCATCGTGGTAGATTACGCGCCGTAGCATGATCCTGGTTCC
        maximum likelihood sequence       TGCTAGAAGCATCGTGGTAGATTACGCGCCGTAGCATGATCCTGGTTCC
        maximum likelihood edit distance  0

evaluating sequence 18 of 100
        observed sequence                 TGCTAGAAGCATCGTGGTAGATTACGCGCCGTAGCATGATCCTGGTTCCA
        maximum likelihood sequence       TGCTAGAAGCATCGTGGTAGATTACGCGCCGTAGCATGATCCTGGTTCCA
        maximum likelihood edit distance  0

evaluating sequence 19 of 100
        observed sequence                 GCTAGAAGCATCGTGGTAGATTACGCGCCGTAGCATGATCCTGGTTCCA
        maximum likelihood sequence       GCTAGAAGCATCGTGGTAGATTACGCGCCGTAGCATGATCCTGGTTCCA
        maximum likelihood edit distance  0
```

```
evaluating sequence 20 of 100
        observed sequence               GCTAGAAGCATCGTGGTAGATTACGCGCCGTAGCATGATCCTGGTTCCA
        maximum likelihood sequence     GCTAGAAGCATCGTGGTAGATTACGCGCCGTAGCATGATCCTGGTTCCA
        maximum likelihood edit distance  0

evaluating sequence 21 of 100
        observed sequence               TGGAACCAGGATCATGCTACGGCGCGTAATCTACCACGATGCTTCTAGCA
        maximum likelihood sequence     TGGAACCAGGATCATGCTACGGCGCGTAATCTACCACGATGCTTCTAGCA
        maximum likelihood edit distance  0

evaluating sequence 22 of 100
        observed sequence               TGGAACCAGGATCATGCTACGGCGCGTAATCTACCACGATGCTTCTAGCA
        maximum likelihood sequence     TGGAACCAGGATCATGCTACGGCGCGTAATCTACCACGATGCTTCTAGCA
        maximum likelihood edit distance  0

evaluating sequence 23 of 100
        observed sequence               GCTAGAAGCATCGTGGTAGATTACGCGCCGTAGCATGATCCTGGTTCCA
        maximum likelihood sequence     GCTAGAAGCATCGTGGTAGATTACGCGCCGTAGCATGATCCTGGTTCCA
        maximum likelihood edit distance  0

evaluating sequence 24 of 100
        observed sequence               TGGAACCAGGATCATGCTACGGCGCGTAATCTACCACGATGCTTCTAGCA
        maximum likelihood sequence     TGGAACCAGGATCATGCTACGGCGCGTAATCTACCACGATGCTTCTAGCA
        maximum likelihood edit distance  0

evaluating sequence 25 of 100
        observed sequence               TGCTAGAAGCATCGTGGTAGATTACGCGCCGTAGCATGATCCTGGTTCCA
        maximum likelihood sequence     TGCTAGAAGCATCGTGGTAGATTACGCGCCGTAGCATGATCCTGGTTCCA
        maximum likelihood edit distance  0

evaluating sequence 26 of 100
        observed sequence               TGGAACCAGGATCATGCTACGGCGCGTAATCTACCACGATGCTTCTAGCA
        maximum likelihood sequence     TGGAACCAGGATCATGCTACGGCGCGTAATCTACCACGATGCTTCTAGCA
        maximum likelihood edit distance  0

evaluating sequence 27 of 100
        observed sequence               TGGAACCAGGATCATGCTACGGCGCGTAATCTACCACGATGCTTCTAGCA
        maximum likelihood sequence     TGGAACCAGGATCATGCTACGGCGCGTAATCTACCACGATGCTTCTAGCA
        maximum likelihood edit distance  0

evaluating sequence 28 of 100
        observed sequence               TGCTAGAAGCATCGTGGTAGATTACGCGCCGTAGCATGATCCTGGTTCCA
        maximum likelihood sequence     TGCTAGAAGCATCGTGGTAGATTACGCGCCGTAGCATGATCCTGGTTCCA
        maximum likelihood edit distance  0

evaluating sequence 29 of 100
        observed sequence               TGCTAGAAGCATCGTGGTAGATTACGCGCCGTAGCATGATCCTGGTTCCA
        maximum likelihood sequence     TGCTAGAAGCATCGTGGTAGATTACGCGCCGTAGCATGATCCTGGTTCCA
        maximum likelihood edit distance  0

evaluating sequence 30 of 100
        observed sequence               CTAGAAGCATCGTGGTAGATTACGCGCCGTAGCATGATCCTGGTTCCA
        maximum likelihood sequence     CTAGAAGCATCGTGGTAGATTACGCGCCGTAGCATGATCCTGGTTCCA
        maximum likelihood edit distance  0

evaluating sequence 31 of 100
        observed sequence               TGGAACCAGGATCATGCTACGGCGCGTAATCTACCACGATGCTTCTAGCA
        maximum likelihood sequence     TGGAACCAGGATCATGCTACGGCGCGTAATCTACCACGATGCTTCTAGCA
        maximum likelihood edit distance  0

evaluating sequence 32 of 100
        observed sequence               TGGAACCAGGATCATGCTACGGCGCGTAATCTACCACGATGCTTCTAGCA
        maximum likelihood sequence     TGGAACCAGGATCATGCTACGGCGCGTAATCTACCACGATGCTTCTAGCA
        maximum likelihood edit distance  0

evaluating sequence 33 of 100
        observed sequence               TGGAACCAGGATCATGCTACGGCGCGTAATCTACCACGATGCTTCTAGCA
        maximum likelihood sequence     TGGAACCAGGATCATGCTACGGCGCGTAATCTACCACGATGCTTCTAGCA
        maximum likelihood edit distance  0

evaluating sequence 34 of 100
        observed sequence               TGGAACCAGGATCATGCTACGGCGCGTAATCTACCACGATGCTTCTAGCA
        maximum likelihood sequence     TGGAACCAGGATCATGCTACGGCGCGTAATCTACCACGATGCTTCTAGCA
        maximum likelihood edit distance  0

evaluating sequence 35 of 100
        observed sequence               TGCTAGAAGCATCGTGGTAGATTACGCGCCGTAGCATGATCCTGGTTCCA
        maximum likelihood sequence     TGCTAGAAGCATCGTGGTAGATTACGCGCCGTAGCATGATCCTGGTTCCA
        maximum likelihood edit distance  0

evaluating sequence 36 of 100
        observed sequence               TGGAACCAGGATCATGCTACGGCGCGTAATCTACCACGATGCTTCTAGCA
        maximum likelihood sequence     TGGAACCAGGATCATGCTACGGCGCGTAATCTACCACGATGCTTCTAGCA
        maximum likelihood edit distance  0

evaluating sequence 37 of 100
        observed sequence               TGGAACCAGGATCATGCTACGGCGCGTAATCTACCACGATGCTTCTAGCA
        maximum likelihood sequence     TGGAACCAGGATCATGCTACGGCGCGTAATCTACCACGATGCTTCTAGCA
        maximum likelihood edit distance  0

evaluating sequence 38 of 100
        observed sequence               TGGAACCAGGATCATGCTACGGCGCGTAATCTACCACGATGCTTCTAGCA
        maximum likelihood sequence     TGGAACCAGGATCATGCTACGGCGCGTAATCTACCACGATGCTTCTAGCA
        maximum likelihood edit distance  0

evaluating sequence 39 of 100
        observed sequence               GCTAGAAGCATCGTGGTAGATTACGCGCCGTAGCATGATCCTGGTTCCA
        maximum likelihood sequence     GCTAGAAGCATCGTGGTAGATTACGCGCCGTAGCATGATCCTGGTTCCA
```

```
        maximum likelihood edit distance  0

evaluating sequence 40 of 100
        observed sequence                 TGGAACCAGGATCATGCTACGGCGCGTAATCTACCACGATGCTTCTAGCA
        maximum likelihood sequence       TGGAACCAGGATCATGCTACGGCGCGTAATCTACCACGATGCTTCTAGCA
        maximum likelihood edit distance  0

evaluating sequence 41 of 100
        observed sequence                 TGGAACCAGGATCATGCTACGGCGCGTAATCTACCACGATGCTTCTAGCA
        maximum likelihood sequence       TGGAACCAGGATCATGCTACGGCGCGTAATCTACCACGATGCTTCTAGCA
        maximum likelihood edit distance  0

evaluating sequence 42 of 100
        observed sequence                 GAACCAGGATCATGCTACGGCGCGTAATCTACCACGATGCTTCTAGC
        maximum likelihood sequence       GAACCAGGATCATGCTACGGCGCGTAATCTACCACGATGCTTCTAGC
        maximum likelihood edit distance  0

evaluating sequence 43 of 100
        observed sequence                 TGGAACCAGGATCATGCTACGGCGCGTAATCTACCACGATGCTTCTAGCA
        maximum likelihood sequence       TGGAACCAGGATCATGCTACGGCGCGTAATCTACCACGATGCTTCTAGCA
        maximum likelihood edit distance  0

evaluating sequence 44 of 100
        observed sequence                 TGCTAGAAGCATCGTGGTAGATTACGCGCCGTAGCATGATCCTGGTTCCA
        maximum likelihood sequence       TGCTAGAAGCATCGTGGTAGATTACGCGCCGTAGCATGATCCTGGTTCCA
        maximum likelihood edit distance  0

evaluating sequence 45 of 100
        observed sequence                 GCTAGAAGCATCGTGGTAGATTACGCGCCGTAGCATGATCCTGGTTCCA
        maximum likelihood sequence       GCTAGAAGCATCGTGGTAGATTACGCGCCGTAGCATGATCCTGGTTCCA
        maximum likelihood edit distance  0

evaluating sequence 46 of 100
        observed sequence                 TGGAACCAGGATCATGCTACGGCGCGTAATCTACCACGATGCTTCTAGCA
        maximum likelihood sequence       TGGAACCAGGATCATGCTACGGCGCGTAATCTACCACGATGCTTCTAGCA
        maximum likelihood edit distance  0

evaluating sequence 47 of 100
        observed sequence                 GGAACCAGGATCATGCTACGGCGCGTAATCTACCACGATGCTTCTAGCA
        maximum likelihood sequence       GGAACCAGGATCATGCTACGGCGCGTAATCTACCACGATGCTTCTAGCA
        maximum likelihood edit distance  0

evaluating sequence 48 of 100
        observed sequence                 TGCTAGAAGCATCGTGGTAGATTACGCGCCGTAGCATGATCCTGGTTCCA
        maximum likelihood sequence       TGCTAGAAGCATCGTGGTAGATTACGCGCCGTAGCATGATCCTGGTTCCA
        maximum likelihood edit distance  0

evaluating sequence 49 of 100
        observed sequence                 GCTAGAAGCATCGTGGTAGATTACGCGCCGTAGCATGATCCTGGTTCC
        maximum likelihood sequence       GCTAGAAGCATCGTGGTAGATTACGCGCCGTAGCATGATCCTGGTTCC
        maximum likelihood edit distance  0

evaluating sequence 50 of 100
        observed sequence                 TGGAACCAGGATCATGCTACGGCGCGTAATCTACCACGATGCTTCTAGCA
        maximum likelihood sequence       TGGAACCAGGATCATGCTACGGCGCGTAATCTACCACGATGCTTCTAGCA
        maximum likelihood edit distance  0

evaluating sequence 51 of 100
        observed sequence                 TGCTAGAAGCATCGTGGTAGATTACGCGCCGTAGCATGATCCTGGTTCC
        maximum likelihood sequence       TGCTAGAAGCATCGTGGTAGATTACGCGCCGTAGCATGATCCTGGTTCC
        maximum likelihood edit distance  0

evaluating sequence 52 of 100
        observed sequence                 CTAGAAGCATCGTGGTAGATTACGCGCCGTAGCATGATCCTGGTTCCA
        maximum likelihood sequence       CTAGAAGCATCGTGGTAGATTACGCGCCGTAGCATGATCCTGGTTCCA
        maximum likelihood edit distance  0

evaluating sequence 53 of 100
        observed sequence                 TGGAACCAGGATCATGCTACGGCGCGTAATCTACCACGATGCTTCTAGC
        maximum likelihood sequence       TGGAACCAGGATCATGCTACGGCGCGTAATCTACCACGATGCTTCTAGC
        maximum likelihood edit distance  0

evaluating sequence 54 of 100
        observed sequence                 TGGAACCAGGATCATGCTACGGCGCGTAATCTACCACGATGCTTCTAGCA
        maximum likelihood sequence       TGGAACCAGGATCATGCTACGGCGCGTAATCTACCACGATGCTTCTAGCA
        maximum likelihood edit distance  0

evaluating sequence 55 of 100
        observed sequence                 GTGGAACCAGGATCATGCTACGGCGCGTAATCTACCACGATGCTTCTAGCA
        maximum likelihood sequence       GTGGAACCAGGATCATGCTACGGCGCGTAATCTACCACGATGCTTCTAGCA
        maximum likelihood edit distance  0

evaluating sequence 56 of 100
        observed sequence                 TGGAACCAGGATCATGCTACGGCGCGTAATCTACCACGATGCTTCTAGCA
        maximum likelihood sequence       TGGAACCAGGATCATGCTACGGCGCGTAATCTACCACGATGCTTCTAGCA
        maximum likelihood edit distance  0

evaluating sequence 57 of 100
        observed sequence                 TGCTAGAAGCATCGTGGTAGATTACGCGCCGTAGCATGATCCTGGTTCCA
        maximum likelihood sequence       TGCTAGAAGCATCGTGGTAGATTACGCGCCGTAGCATGATCCTGGTTCCA
        maximum likelihood edit distance  0

evaluating sequence 58 of 100
        observed sequence                 TGCTAGAAGCATCGTGGTAGATTACGCGCCGTAGCATGATCCTGGTTCCA
        maximum likelihood sequence       TGCTAGAAGCATCGTGGTAGATTACGCGCCGTAGCATGATCCTGGTTCCA
        maximum likelihood edit distance  0

evaluating sequence 59 of 100
        observed sequence                 TGGAACCAGGATCATGCTACGGCGCGTAATCTACCACGATGCTTCTAGCA
```

```
        maximum likelihood sequence       TGGAACCAGGATCATGCTACGGCGCGTAATCTACCACGATGCTTCTAGCA
        maximum likelihood edit distance  0

evaluating sequence 60 of 100
        observed sequence                 TGCTAGAAGCATCGTGGTAGATTACGCGCCGTAGCATGATCCTGGTTCCA
        maximum likelihood sequence       TGCTAGAAGCATCGTGGTAGATTACGCGCCGTAGCATGATCCTGGTTCCA
        maximum likelihood edit distance  0

evaluating sequence 61 of 100
        observed sequence                 GCTAGAAGCATCGTGGTAGATTACGCGCCGTAGCATGATCCTGGTTCCA
        maximum likelihood sequence       GCTAGAAGCATCGTGGTAGATTACGCGCCGTAGCATGATCCTGGTTCCA
        maximum likelihood edit distance  0

evaluating sequence 62 of 100
        observed sequence                 GAACCAGGATCATGCTACGGCGCGTAATCTACCACGATGCTTCTAGCA
        maximum likelihood sequence       GAACCAGGATCATGCTACGGCGCGTAATCTACCACGATGCTTCTAGCA
        maximum likelihood edit distance  0

evaluating sequence 63 of 100
        observed sequence                 GGAACCAGGATCATGCTACGGCGCGTAATCTACCACGATGCTTCTAGC
        maximum likelihood sequence       GGAACCAGGATCATGCTACGGCGCGTAATCTACCACGATGCTTCTAGC
        maximum likelihood edit distance  0

evaluating sequence 64 of 100
        observed sequence                 TGGAACCAGGATCATGCTACGGCGCGTAATCTACCACGATGCTTCTAGCA
        maximum likelihood sequence       TGGAACCAGGATCATGCTACGGCGCGTAATCTACCACGATGCTTCTAGCA
        maximum likelihood edit distance  0

evaluating sequence 65 of 100
        observed sequence                 GTGGAACCAGGATCATGCTACGGCGCGTAATCTACCACGATGCTTCTAGCA
        maximum likelihood sequence       GTGGAACCAGGATCATGCTACGGCGCGTAATCTACCACGATGCTTCTAGCA
        maximum likelihood edit distance  0

evaluating sequence 66 of 100
        observed sequence                 TGGAACCAGGATCATGCTACGGCGCGTAATCTACCACGATGCTTCTAGCA
        maximum likelihood sequence       TGGAACCAGGATCATGCTACGGCGCGTAATCTACCACGATGCTTCTAGCA
        maximum likelihood edit distance  0

evaluating sequence 67 of 100
        observed sequence                 TGGAACCAGGATCATGCTACGGCGCGTAATCTACCACGATGCTTCTAGC
        maximum likelihood sequence       TGGAACCAGGATCATGCTACGGCGCGTAATCTACCACGATGCTTCTAGC
        maximum likelihood edit distance  0

evaluating sequence 68 of 100
        observed sequence                 GCTAGAAGCATCGTGGTAGATTACGCGCCGTAGCATGATCCTGGTTCCA
        maximum likelihood sequence       GCTAGAAGCATCGTGGTAGATTACGCGCCGTAGCATGATCCTGGTTCCA
        maximum likelihood edit distance  0

evaluating sequence 69 of 100
        observed sequence                 TGCTAGAAGCATCGTGGTAGATTACGCGCCGTAGCATGATCCTGGTTCCA
        maximum likelihood sequence       TGCTAGAAGCATCGTGGTAGATTACGCGCCGTAGCATGATCCTGGTTCCA
        maximum likelihood edit distance  0

evaluating sequence 70 of 100
        observed sequence                 TGGAACCAGGATCATGCTACGGCGCGTAATCTACCACGATGCTTCTAGCA
        maximum likelihood sequence       TGGAACCAGGATCATGCTACGGCGCGTAATCTACCACGATGCTTCTAGCA
        maximum likelihood edit distance  0

evaluating sequence 71 of 100
        observed sequence                 TTGCTAGAAGCATCGTGGTAGATTACGCGCCGTAGCATGATCCTGGTTCCA
        maximum likelihood sequence       TTGCTAGAAGCATCGTGGTAGATTACGCGCCGTAGCATGATCCTGGTTCCA
        maximum likelihood edit distance  0

evaluating sequence 72 of 100
        observed sequence                 GGAACCAGGATCATGCTACGGCGCGTAATCTACCACGATGCTTCTAGC
        maximum likelihood sequence       GGAACCAGGATCATGCTACGGCGCGTAATCTACCACGATGCTTCTAGC
        maximum likelihood edit distance  0

evaluating sequence 73 of 100
        observed sequence                 TGCTAGAAGCATCGTGGTAGATTACGCGCCGTAGCATGATCCTGGTTCCA
        maximum likelihood sequence       TGCTAGAAGCATCGTGGTAGATTACGCGCCGTAGCATGATCCTGGTTCCA
        maximum likelihood edit distance  0

evaluating sequence 74 of 100
        observed sequence                 TGCTAGAAGCATCGTGGTAGATTACGCGCCGTAGCATGATCCTGGTTCC
        maximum likelihood sequence       TGCTAGAAGCATCGTGGTAGATTACGCGCCGTAGCATGATCCTGGTTCC
        maximum likelihood edit distance  0

evaluating sequence 75 of 100
        observed sequence                 GCTAGAAGCATCGTGGTAGATTACGCGCCGTAGCATGATCCTGGTTCCA
        maximum likelihood sequence       GCTAGAAGCATCGTGGTAGATTACGCGCCGTAGCATGATCCTGGTTCCA
        maximum likelihood edit distance  0

evaluating sequence 76 of 100
        observed sequence                 TGGAACCAGGATCATGCTACGGCGCGTAATCTACCACGATGCTTCTAGCA
        maximum likelihood sequence       TGGAACCAGGATCATGCTACGGCGCGTAATCTACCACGATGCTTCTAGCA
        maximum likelihood edit distance  0

evaluating sequence 77 of 100
        observed sequence                 TGCTAGAAGCATCGTGGTAGATTACGCGCCGTAGCATGATCCTGGTTCCA
        maximum likelihood sequence       TGCTAGAAGCATCGTGGTAGATTACGCGCCGTAGCATGATCCTGGTTCCA
        maximum likelihood edit distance  0

evaluating sequence 78 of 100
        observed sequence                 CTAGAAGCATCGTGGTAGATTACGCGCCGTAGCATGATCCTGGTTCCA
        maximum likelihood sequence       CTAGAAGCATCGTGGTAGATTACGCGCCGTAGCATGATCCTGGTTCCA
        maximum likelihood edit distance  0

evaluating sequence 79 of 100
```

```
        observed sequence                 TGCTAGAAGCATCGTGGTAGATTACGCGCCGTAGCATGATCCTGGTTCCA
        maximum likelihood sequence       TGCTAGAAGCATCGTGGTAGATTACGCGCCGTAGCATGATCCTGGTTCCA
        maximum likelihood edit distance  0

evaluating sequence 80 of 100
        observed sequence                 TGCTAGAAGCATCGTGGTAGATTACGCGCCGTAGCATGATCCTGGTTCCA
        maximum likelihood sequence       TGCTAGAAGCATCGTGGTAGATTACGCGCCGTAGCATGATCCTGGTTCCA
        maximum likelihood edit distance  0

evaluating sequence 81 of 100
        observed sequence                 GAACCAGGATCATGCTACGGCGCGTAATCTACCACGATGCTTCTAGCA
        maximum likelihood sequence       GAACCAGGATCATGCTACGGCGCGTAATCTACCACGATGCTTCTAGCA
        maximum likelihood edit distance  0

evaluating sequence 82 of 100
        observed sequence                 TGGAACCAGGATCATGCTACGGCGCGTAATCTACCACGATGCTTCTAGCA
        maximum likelihood sequence       TGGAACCAGGATCATGCTACGGCGCGTAATCTACCACGATGCTTCTAGCA
        maximum likelihood edit distance  0

evaluating sequence 83 of 100
        observed sequence                 GTGGAACCAGGATCATGCTACGGCGCGTAATCTACCACGATGCTTCTAGCA
        maximum likelihood sequence       GTGGAACCAGGATCATGCTACGGCGCGTAATCTACCACGATGCTTCTAGCA
        maximum likelihood edit distance  0

evaluating sequence 84 of 100
        observed sequence                 TGGAACCAGGATCATGCTACGGCGCGTAATCTACCACGATGCTTCTAGCA
        maximum likelihood sequence       TGGAACCAGGATCATGCTACGGCGCGTAATCTACCACGATGCTTCTAGCA
        maximum likelihood edit distance  0

evaluating sequence 85 of 100
        observed sequence                 TGGAACCAGGATCATGCTACGGCGCGTAATCTACCACGATGCTTCTAGCA
        maximum likelihood sequence       TGGAACCAGGATCATGCTACGGCGCGTAATCTACCACGATGCTTCTAGCA
        maximum likelihood edit distance  0

evaluating sequence 86 of 100
        observed sequence                 TGCTAGAAGCATCGTGGTAGATTACGCGCCGTAGCATGATCCTGGTTCCA
        maximum likelihood sequence       TGCTAGAAGCATCGTGGTAGATTACGCGCCGTAGCATGATCCTGGTTCCA
        maximum likelihood edit distance  0

evaluating sequence 87 of 100
        observed sequence                 TGGAACCAGGATCATGCTACGGCGCGTAATCTACCACGATGCTTCTAGCA
        maximum likelihood sequence       TGGAACCAGGATCATGCTACGGCGCGTAATCTACCACGATGCTTCTAGCA
        maximum likelihood edit distance  0

evaluating sequence 88 of 100
        observed sequence                 TGGAACCAGGATCATGCTACGGCGCGTAATCTACCACGATGCTTCTAGC
        maximum likelihood sequence       TGGAACCAGGATCATGCTACGGCGCGTAATCTACCACGATGCTTCTAGC
        maximum likelihood edit distance  0

evaluating sequence 89 of 100
        observed sequence                 TGCTAGAAGCATCGTGGTAGATTACGCGCCGTAGCATGATCCTGGTTCCA
        maximum likelihood sequence       TGCTAGAAGCATCGTGGTAGATTACGCGCCGTAGCATGATCCTGGTTCCA
        maximum likelihood edit distance  0

evaluating sequence 90 of 100
        observed sequence                 TGGAACCAGGATCATGCTACGGCGCGTAATCTACCACGATGCTTCTAGCA
        maximum likelihood sequence       TGGAACCAGGATCATGCTACGGCGCGTAATCTACCACGATGCTTCTAGCA
        maximum likelihood edit distance  0

evaluating sequence 91 of 100
        observed sequence                 TGGAACCAGGATCATGCTACGGCGCGTAATCTACCACGATGCTTCTAGCA
        maximum likelihood sequence       TGGAACCAGGATCATGCTACGGCGCGTAATCTACCACGATGCTTCTAGCA
        maximum likelihood edit distance  0

evaluating sequence 92 of 100
        observed sequence                 TGCTAGAAGCATCGTGGTAGATTACGCGCCGTAGCATGATCCTGGTTCCA
        maximum likelihood sequence       TGCTAGAAGCATCGTGGTAGATTACGCGCCGTAGCATGATCCTGGTTCCA
        maximum likelihood edit distance  0

evaluating sequence 93 of 100
        observed sequence                 TGCTAGAAGCATCGTGGTAGATTACGCGCCGTAGCATGATCCTGGTTCCA
        maximum likelihood sequence       TGCTAGAAGCATCGTGGTAGATTACGCGCCGTAGCATGATCCTGGTTCCA
        maximum likelihood edit distance  0

evaluating sequence 94 of 100
        observed sequence                 TGGAACCAGGATCATGCTACGGCGCGTAATCTACCACGATGCTTCTAGCA
        maximum likelihood sequence       TGGAACCAGGATCATGCTACGGCGCGTAATCTACCACGATGCTTCTAGCA
        maximum likelihood edit distance  0

evaluating sequence 95 of 100
        observed sequence                 TGCTAGAAGCATCGTGGTAGATTACGCGCCGTAGCATGATCCTGGTTCCA
        maximum likelihood sequence       TGCTAGAAGCATCGTGGTAGATTACGCGCCGTAGCATGATCCTGGTTCCA
        maximum likelihood edit distance  0

evaluating sequence 96 of 100
        observed sequence                 GGAACCAGGATCATGCTACGGCGCGTAATCTACCACGATGCTTCTAGCA
        maximum likelihood sequence       GGAACCAGGATCATGCTACGGCGCGTAATCTACCACGATGCTTCTAGCA
        maximum likelihood edit distance  0

evaluating sequence 97 of 100
        observed sequence                 TTGCTAGAAGCATCGTGGTAGATTACGCGCCGTAGCATGATCCTGGTTCCA
        maximum likelihood sequence       TTGCTAGAAGCATCGTGGTAGATTACGCGCCGTAGCATGATCCTGGTTCCA
        maximum likelihood edit distance  0

evaluating sequence 98 of 100
        observed sequence                 TGGAACCAGGATCATGCTACGGCGCGTAATCTACCACGATGCTTCTAGC
        maximum likelihood sequence       TGGAACCAGGATCATGCTACGGCGCGTAATCTACCACGATGCTTCTAGC
        maximum likelihood edit distance  0
```

```
evaluating sequence 99 of 100
        observed sequence                 TGCTAGAAGCATCGTGGTAGATTACGCGCCGTAGCATGATCCTGGTTCCA
        maximum likelihood sequence       TGCTAGAAGCATCGTGGTAGATTACGCGCCGTAGCATGATCCTGGTTCCA
        maximum likelihood edit distance  0

evaluating sequence 100 of 100
        observed sequence                 GTGGAACCAGGATCATGCTACGGCGCGTAATCTACCACGATGCTTCTAGC
        maximum likelihood sequence       GTGGAACCAGGATCATGCTACGGCGCGTAATCTACCACGATGCTTCTAGC
        maximum likelihood edit distance  0

DATASET STATISTICS:
        assumed error rate    12.5%
        total bases observed  4966
        total edits accepted  0
        inferred error rate   0.0%
```

In [37]:

```
k = 11
canonical_kmers = collect(keys(Eisenia.count_canonical_kmers(maximum_likelihood_observations, k)))
stranded_kmer_graph = Eisenia.build_stranded_kmer_graph(canonical_kmers, maximum_likelihood_observations)
filename = reference_sequence_id * "." * replace(string(Dates.now()), ':' => '.') * ".svg"
Eisenia.plot_stranded_kmer_graph(stranded_kmer_graph, filename=filename)
HTML("""
<image src="$filename" width=50%>
""")
```

Out[37]:

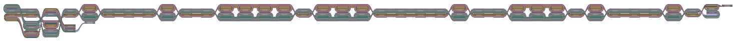

In [38]:

```
maximum_likelihood_observations = Eisenia.viterbi_maximum_likelihood_traversals(stranded_kmer_graph, verbosity="reads");
```

```
computing kmer counts...
computing kmer state likelihoods...
finding shortest paths between kmers...
finding viterbi maximum likelihood paths for observed sequences...

evaluating sequence 1 of 100
        observed sequence                 TGGAACCAGGATCATGCTACGGCGCGTAATCTACCACGATGCTTCTAGCA
        maximum likelihood sequence       TGGAACCAGGATCATGCTACGGCGCGTAATCTACCACGATGCTTCTAGCA
        maximum likelihood edit distance  0

evaluating sequence 2 of 100
        observed sequence                 TGGAACCAGGATCATGCTACGGCGCGTAATCTACCACGATGCTTCTAGCA
        maximum likelihood sequence       TGGAACCAGGATCATGCTACGGCGCGTAATCTACCACGATGCTTCTAGCA
        maximum likelihood edit distance  0

evaluating sequence 3 of 100
        observed sequence                 TGCTAGAAGCATCGTGGTAGATTACGCGCCGTAGCATGATCCTGGTTCCA
        maximum likelihood sequence       TGCTAGAAGCATCGTGGTAGATTACGCGCCGTAGCATGATCCTGGTTCCA
        maximum likelihood edit distance  0

evaluating sequence 4 of 100
        observed sequence                 TGGAACCAGGATCATGCTACGGCGCGTAATCTACCACGATGCTTCTAGCA
        maximum likelihood sequence       TGGAACCAGGATCATGCTACGGCGCGTAATCTACCACGATGCTTCTAGCA
        maximum likelihood edit distance  0

evaluating sequence 5 of 100
        observed sequence                 TGGAACCAGGATCATGCTACGGCGCGTAATCTACCACGATGCTTCTAGCA
        maximum likelihood sequence       TGGAACCAGGATCATGCTACGGCGCGTAATCTACCACGATGCTTCTAGCA
        maximum likelihood edit distance  0

evaluating sequence 6 of 100
        observed sequence                 TGCTAGAAGCATCGTGGTAGATTACGCGCCGTAGCATGATCCTGGTTCCA
        maximum likelihood sequence       TGCTAGAAGCATCGTGGTAGATTACGCGCCGTAGCATGATCCTGGTTCCA
        maximum likelihood edit distance  0

evaluating sequence 7 of 100
        observed sequence                 TTGCTAGAAGCATCGTGGTAGATTACGCGCCGTAGCATGATCCTGGTTCCA
        maximum likelihood sequence       TTGCTAGAAGCATCGTGGTAGATTACGCGCCGTAGCATGATCCTGGTTCCA
        maximum likelihood edit distance  0

evaluating sequence 8 of 100
        observed sequence                 TGGAACCAGGATCATGCTACGGCGCGTAATCTACCACGATGCTTCTAGCA
        maximum likelihood sequence       TGGAACCAGGATCATGCTACGGCGCGTAATCTACCACGATGCTTCTAGCA
        maximum likelihood edit distance  0

evaluating sequence 9 of 100
        observed sequence                 TGCTAGAAGCATCGTGGTAGATTACGCGCCGTAGCATGATCCTGGTTCCA
        maximum likelihood sequence       TGCTAGAAGCATCGTGGTAGATTACGCGCCGTAGCATGATCCTGGTTCCA
        maximum likelihood edit distance  0

evaluating sequence 10 of 100
        observed sequence                 TGCTAGAAGCATCGTGGTAGATTACGCGCCGTAGCATGATCCTGGTTCCA
        maximum likelihood sequence       TGCTAGAAGCATCGTGGTAGATTACGCGCCGTAGCATGATCCTGGTTCCA
        maximum likelihood edit distance  0

evaluating sequence 11 of 100
        observed sequence                 TGGAACCAGGATCATGCTACGGCGCGTAATCTACCACGATGCTTCTAGCA
        maximum likelihood sequence       TGGAACCAGGATCATGCTACGGCGCGTAATCTACCACGATGCTTCTAGCA
        maximum likelihood edit distance  0

evaluating sequence 12 of 100
        observed sequence                 TGGAACCAGGATCATGCTACGGCGCGTAATCTACCACGATGCTTCTAGCA
        maximum likelihood sequence       TGGAACCAGGATCATGCTACGGCGCGTAATCTACCACGATGCTTCTAGCA
        maximum likelihood edit distance  0

evaluating sequence 13 of 100
        observed sequence                 TGCTAGAAGCATCGTGGTAGATTACGCGCCGTAGCATGATCCTGGTTCC
        maximum likelihood sequence       TGCTAGAAGCATCGTGGTAGATTACGCGCCGTAGCATGATCCTGGTTCC
        maximum likelihood edit distance  0

evaluating sequence 14 of 100
        observed sequence                 TGCTAGAAGCATCGTGGTAGATTACGCGCCGTAGCATGATCCTGGTTCCA
        maximum likelihood sequence       TGCTAGAAGCATCGTGGTAGATTACGCGCCGTAGCATGATCCTGGTTCCA
        maximum likelihood edit distance  0

evaluating sequence 15 of 100
        observed sequence                 AACCAGGATCATGCTACGGCGCGTAATCTACCACGATGCTTCTAGCA
        maximum likelihood sequence       AACCAGGATCATGCTACGGCGCGTAATCTACCACGATGCTTCTAGCA
        maximum likelihood edit distance  0

evaluating sequence 16 of 100
        observed sequence                 TGCTAGAAGCATCGTGGTAGATTACGCGCCGTAGCATGATCCTGGTTCCA
        maximum likelihood sequence       TGCTAGAAGCATCGTGGTAGATTACGCGCCGTAGCATGATCCTGGTTCCA
        maximum likelihood edit distance  0

evaluating sequence 17 of 100
        observed sequence                 TGCTAGAAGCATCGTGGTAGATTACGCGCCGTAGCATGATCCTGGTTCC
        maximum likelihood sequence       TGCTAGAAGCATCGTGGTAGATTACGCGCCGTAGCATGATCCTGGTTCC
        maximum likelihood edit distance  0

evaluating sequence 18 of 100
        observed sequence                 TGCTAGAAGCATCGTGGTAGATTACGCGCCGTAGCATGATCCTGGTTCCA
        maximum likelihood sequence       TGCTAGAAGCATCGTGGTAGATTACGCGCCGTAGCATGATCCTGGTTCCA
        maximum likelihood edit distance  0

evaluating sequence 19 of 100
        observed sequence                 GCTAGAAGCATCGTGGTAGATTACGCGCCGTAGCATGATCCTGGTTCCA
        maximum likelihood sequence       GCTAGAAGCATCGTGGTAGATTACGCGCCGTAGCATGATCCTGGTTCCA
        maximum likelihood edit distance  0
```

```
evaluating sequence 20 of 100
        observed sequence                GCTAGAAGCATCGTGGTAGATTACGCGCCGTAGCATGATCCTGGTTCCA
        maximum likelihood sequence      GCTAGAAGCATCGTGGTAGATTACGCGCCGTAGCATGATCCTGGTTCCA
        maximum likelihood edit distance  0

evaluating sequence 21 of 100
        observed sequence                TGGAACCAGGATCATGCTACGGCGCGTAATCTACCACGATGCTTCTAGCA
        maximum likelihood sequence      TGGAACCAGGATCATGCTACGGCGCGTAATCTACCACGATGCTTCTAGCA
        maximum likelihood edit distance  0

evaluating sequence 22 of 100
        observed sequence                TGGAACCAGGATCATGCTACGGCGCGTAATCTACCACGATGCTTCTAGCA
        maximum likelihood sequence      TGGAACCAGGATCATGCTACGGCGCGTAATCTACCACGATGCTTCTAGCA
        maximum likelihood edit distance  0

evaluating sequence 23 of 100
        observed sequence                GCTAGAAGCATCGTGGTAGATTACGCGCCGTAGCATGATCCTGGTTCCA
        maximum likelihood sequence      GCTAGAAGCATCGTGGTAGATTACGCGCCGTAGCATGATCCTGGTTCCA
        maximum likelihood edit distance  0

evaluating sequence 24 of 100
        observed sequence                TGGAACCAGGATCATGCTACGGCGCGTAATCTACCACGATGCTTCTAGCA
        maximum likelihood sequence      TGGAACCAGGATCATGCTACGGCGCGTAATCTACCACGATGCTTCTAGCA
        maximum likelihood edit distance  0

evaluating sequence 25 of 100
        observed sequence                TGCTAGAAGCATCGTGGTAGATTACGCGCCGTAGCATGATCCTGGTTCCA
        maximum likelihood sequence      TGCTAGAAGCATCGTGGTAGATTACGCGCCGTAGCATGATCCTGGTTCCA
        maximum likelihood edit distance  0

evaluating sequence 26 of 100
        observed sequence                TGGAACCAGGATCATGCTACGGCGCGTAATCTACCACGATGCTTCTAGCA
        maximum likelihood sequence      TGGAACCAGGATCATGCTACGGCGCGTAATCTACCACGATGCTTCTAGCA
        maximum likelihood edit distance  0

evaluating sequence 27 of 100
        observed sequence                TGGAACCAGGATCATGCTACGGCGCGTAATCTACCACGATGCTTCTAGCA
        maximum likelihood sequence      TGGAACCAGGATCATGCTACGGCGCGTAATCTACCACGATGCTTCTAGCA
        maximum likelihood edit distance  0

evaluating sequence 28 of 100
        observed sequence                TGCTAGAAGCATCGTGGTAGATTACGCGCCGTAGCATGATCCTGGTTCCA
        maximum likelihood sequence      TGCTAGAAGCATCGTGGTAGATTACGCGCCGTAGCATGATCCTGGTTCCA
        maximum likelihood edit distance  0

evaluating sequence 29 of 100
        observed sequence                TGCTAGAAGCATCGTGGTAGATTACGCGCCGTAGCATGATCCTGGTTCCA
        maximum likelihood sequence      TGCTAGAAGCATCGTGGTAGATTACGCGCCGTAGCATGATCCTGGTTCCA
        maximum likelihood edit distance  0

evaluating sequence 30 of 100
        observed sequence                CTAGAAGCATCGTGGTAGATTACGCGCCGTAGCATGATCCTGGTTCCA
        maximum likelihood sequence      CTAGAAGCATCGTGGTAGATTACGCGCCGTAGCATGATCCTGGTTCCA
        maximum likelihood edit distance  0

evaluating sequence 31 of 100
        observed sequence                TGGAACCAGGATCATGCTACGGCGCGTAATCTACCACGATGCTTCTAGCA
        maximum likelihood sequence      TGGAACCAGGATCATGCTACGGCGCGTAATCTACCACGATGCTTCTAGCA
        maximum likelihood edit distance  0

evaluating sequence 32 of 100
        observed sequence                TGGAACCAGGATCATGCTACGGCGCGTAATCTACCACGATGCTTCTAGCA
        maximum likelihood sequence      TGGAACCAGGATCATGCTACGGCGCGTAATCTACCACGATGCTTCTAGCA
        maximum likelihood edit distance  0

evaluating sequence 33 of 100
        observed sequence                TGGAACCAGGATCATGCTACGGCGCGTAATCTACCACGATGCTTCTAGCA
        maximum likelihood sequence      TGGAACCAGGATCATGCTACGGCGCGTAATCTACCACGATGCTTCTAGCA
        maximum likelihood edit distance  0

evaluating sequence 34 of 100
        observed sequence                TGGAACCAGGATCATGCTACGGCGCGTAATCTACCACGATGCTTCTAGCA
        maximum likelihood sequence      TGGAACCAGGATCATGCTACGGCGCGTAATCTACCACGATGCTTCTAGCA
        maximum likelihood edit distance  0

evaluating sequence 35 of 100
        observed sequence                TGCTAGAAGCATCGTGGTAGATTACGCGCCGTAGCATGATCCTGGTTCCA
        maximum likelihood sequence      TGCTAGAAGCATCGTGGTAGATTACGCGCCGTAGCATGATCCTGGTTCCA
        maximum likelihood edit distance  0

evaluating sequence 36 of 100
        observed sequence                TGGAACCAGGATCATGCTACGGCGCGTAATCTACCACGATGCTTCTAGCA
        maximum likelihood sequence      TGGAACCAGGATCATGCTACGGCGCGTAATCTACCACGATGCTTCTAGCA
        maximum likelihood edit distance  0

evaluating sequence 37 of 100
        observed sequence                TGGAACCAGGATCATGCTACGGCGCGTAATCTACCACGATGCTTCTAGCA
        maximum likelihood sequence      TGGAACCAGGATCATGCTACGGCGCGTAATCTACCACGATGCTTCTAGCA
        maximum likelihood edit distance  0

evaluating sequence 38 of 100
        observed sequence                TGGAACCAGGATCATGCTACGGCGCGTAATCTACCACGATGCTTCTAGCA
        maximum likelihood sequence      TGGAACCAGGATCATGCTACGGCGCGTAATCTACCACGATGCTTCTAGCA
        maximum likelihood edit distance  0

evaluating sequence 39 of 100
        observed sequence                GCTAGAAGCATCGTGGTAGATTACGCGCCGTAGCATGATCCTGGTTCCA
        maximum likelihood sequence      GCTAGAAGCATCGTGGTAGATTACGCGCCGTAGCATGATCCTGGTTCCA
```

```
        maximum likelihood edit distance  0

evaluating sequence 40 of 100
        observed sequence                 TGGAACCAGGATCATGCTACGGCGCGTAATCTACCACGATGCTTCTAGCA
        maximum likelihood sequence       TGGAACCAGGATCATGCTACGGCGCGTAATCTACCACGATGCTTCTAGCA
        maximum likelihood edit distance  0

evaluating sequence 41 of 100
        observed sequence                 TGGAACCAGGATCATGCTACGGCGCGTAATCTACCACGATGCTTCTAGCA
        maximum likelihood sequence       TGGAACCAGGATCATGCTACGGCGCGTAATCTACCACGATGCTTCTAGCA
        maximum likelihood edit distance  0

evaluating sequence 42 of 100
        observed sequence                 GAACCAGGATCATGCTACGGCGCGTAATCTACCACGATGCTTCTAGC
        maximum likelihood sequence       GAACCAGGATCATGCTACGGCGCGTAATCTACCACGATGCTTCTAGC
        maximum likelihood edit distance  0

evaluating sequence 43 of 100
        observed sequence                 TGGAACCAGGATCATGCTACGGCGCGTAATCTACCACGATGCTTCTAGCA
        maximum likelihood sequence       TGGAACCAGGATCATGCTACGGCGCGTAATCTACCACGATGCTTCTAGCA
        maximum likelihood edit distance  0

evaluating sequence 44 of 100
        observed sequence                 TGCTAGAAGCATCGTGGTAGATTACGCGCCGTAGCATGATCCTGGTTCCA
        maximum likelihood sequence       TGCTAGAAGCATCGTGGTAGATTACGCGCCGTAGCATGATCCTGGTTCCA
        maximum likelihood edit distance  0

evaluating sequence 45 of 100
        observed sequence                 GCTAGAAGCATCGTGGTAGATTACGCGCCGTAGCATGATCCTGGTTCCA
        maximum likelihood sequence       GCTAGAAGCATCGTGGTAGATTACGCGCCGTAGCATGATCCTGGTTCCA
        maximum likelihood edit distance  0

evaluating sequence 46 of 100
        observed sequence                 TGGAACCAGGATCATGCTACGGCGCGTAATCTACCACGATGCTTCTAGCA
        maximum likelihood sequence       TGGAACCAGGATCATGCTACGGCGCGTAATCTACCACGATGCTTCTAGCA
        maximum likelihood edit distance  0

evaluating sequence 47 of 100
        observed sequence                 GGAACCAGGATCATGCTACGGCGCGTAATCTACCACGATGCTTCTAGCA
        maximum likelihood sequence       GGAACCAGGATCATGCTACGGCGCGTAATCTACCACGATGCTTCTAGCA
        maximum likelihood edit distance  0

evaluating sequence 48 of 100
        observed sequence                 TGCTAGAAGCATCGTGGTAGATTACGCGCCGTAGCATGATCCTGGTTCCA
        maximum likelihood sequence       TGCTAGAAGCATCGTGGTAGATTACGCGCCGTAGCATGATCCTGGTTCCA
        maximum likelihood edit distance  0

evaluating sequence 49 of 100
        observed sequence                 GCTAGAAGCATCGTGGTAGATTACGCGCCGTAGCATGATCCTGGTTCC
        maximum likelihood sequence       GCTAGAAGCATCGTGGTAGATTACGCGCCGTAGCATGATCCTGGTTCC
        maximum likelihood edit distance  0

evaluating sequence 50 of 100
        observed sequence                 TGGAACCAGGATCATGCTACGGCGCGTAATCTACCACGATGCTTCTAGCA
        maximum likelihood sequence       TGGAACCAGGATCATGCTACGGCGCGTAATCTACCACGATGCTTCTAGCA
        maximum likelihood edit distance  0

evaluating sequence 51 of 100
        observed sequence                 TGCTAGAAGCATCGTGGTAGATTACGCGCCGTAGCATGATCCTGGTTCC
        maximum likelihood sequence       TGCTAGAAGCATCGTGGTAGATTACGCGCCGTAGCATGATCCTGGTTCC
        maximum likelihood edit distance  0

evaluating sequence 52 of 100
        observed sequence                 CTAGAAGCATCGTGGTAGATTACGCGCCGTAGCATGATCCTGGTTCCA
        maximum likelihood sequence       CTAGAAGCATCGTGGTAGATTACGCGCCGTAGCATGATCCTGGTTCCA
        maximum likelihood edit distance  0

evaluating sequence 53 of 100
        observed sequence                 TGGAACCAGGATCATGCTACGGCGCGTAATCTACCACGATGCTTCTAGC
        maximum likelihood sequence       TGGAACCAGGATCATGCTACGGCGCGTAATCTACCACGATGCTTCTAGC
        maximum likelihood edit distance  0

evaluating sequence 54 of 100
        observed sequence                 TGGAACCAGGATCATGCTACGGCGCGTAATCTACCACGATGCTTCTAGCA
        maximum likelihood sequence       TGGAACCAGGATCATGCTACGGCGCGTAATCTACCACGATGCTTCTAGCA
        maximum likelihood edit distance  0

evaluating sequence 55 of 100
        observed sequence                 GTGGAACCAGGATCATGCTACGGCGCGTAATCTACCACGATGCTTCTAGCA
        maximum likelihood sequence       GTGGAACCAGGATCATGCTACGGCGCGTAATCTACCACGATGCTTCTAGCA
        maximum likelihood edit distance  0

evaluating sequence 56 of 100
        observed sequence                 TGGAACCAGGATCATGCTACGGCGCGTAATCTACCACGATGCTTCTAGCA
        maximum likelihood sequence       TGGAACCAGGATCATGCTACGGCGCGTAATCTACCACGATGCTTCTAGCA
        maximum likelihood edit distance  0

evaluating sequence 57 of 100
        observed sequence                 TGCTAGAAGCATCGTGGTAGATTACGCGCCGTAGCATGATCCTGGTTCCA
        maximum likelihood sequence       TGCTAGAAGCATCGTGGTAGATTACGCGCCGTAGCATGATCCTGGTTCCA
        maximum likelihood edit distance  0

evaluating sequence 58 of 100
        observed sequence                 TGCTAGAAGCATCGTGGTAGATTACGCGCCGTAGCATGATCCTGGTTCCA
        maximum likelihood sequence       TGCTAGAAGCATCGTGGTAGATTACGCGCCGTAGCATGATCCTGGTTCCA
        maximum likelihood edit distance  0

evaluating sequence 59 of 100
        observed sequence                 TGGAACCAGGATCATGCTACGGCGCGTAATCTACCACGATGCTTCTAGCA
```

```
        maximum likelihood sequence       TGGAACCAGGATCATGCTACGGCGCGTAATCTACCACGATGCTTCTAGCA
        maximum likelihood edit distance  0

evaluating sequence 60 of 100
        observed sequence                 TGCTAGAAGCATCGTGGTAGATTACGCGCCGTAGCATGATCCTGGTTCCA
        maximum likelihood sequence       TGCTAGAAGCATCGTGGTAGATTACGCGCCGTAGCATGATCCTGGTTCCA
        maximum likelihood edit distance  0

evaluating sequence 61 of 100
        observed sequence                 GCTAGAAGCATCGTGGTAGATTACGCGCCGTAGCATGATCCTGGTTCCA
        maximum likelihood sequence       GCTAGAAGCATCGTGGTAGATTACGCGCCGTAGCATGATCCTGGTTCCA
        maximum likelihood edit distance  0

evaluating sequence 62 of 100
        observed sequence                 GAACCAGGATCATGCTACGGCGCGTAATCTACCACGATGCTTCTAGCA
        maximum likelihood sequence       GAACCAGGATCATGCTACGGCGCGTAATCTACCACGATGCTTCTAGCA
        maximum likelihood edit distance  0

evaluating sequence 63 of 100
        observed sequence                 GGAACCAGGATCATGCTACGGCGCGTAATCTACCACGATGCTTCTAGC
        maximum likelihood sequence       GGAACCAGGATCATGCTACGGCGCGTAATCTACCACGATGCTTCTAGC
        maximum likelihood edit distance  0

evaluating sequence 64 of 100
        observed sequence                 TGGAACCAGGATCATGCTACGGCGCGTAATCTACCACGATGCTTCTAGCA
        maximum likelihood sequence       TGGAACCAGGATCATGCTACGGCGCGTAATCTACCACGATGCTTCTAGCA
        maximum likelihood edit distance  0

evaluating sequence 65 of 100
        observed sequence                 GTGGAACCAGGATCATGCTACGGCGCGTAATCTACCACGATGCTTCTAGCA
        maximum likelihood sequence       GTGGAACCAGGATCATGCTACGGCGCGTAATCTACCACGATGCTTCTAGCA
        maximum likelihood edit distance  0

evaluating sequence 66 of 100
        observed sequence                 TGGAACCAGGATCATGCTACGGCGCGTAATCTACCACGATGCTTCTAGCA
        maximum likelihood sequence       TGGAACCAGGATCATGCTACGGCGCGTAATCTACCACGATGCTTCTAGCA
        maximum likelihood edit distance  0

evaluating sequence 67 of 100
        observed sequence                 TGGAACCAGGATCATGCTACGGCGCGTAATCTACCACGATGCTTCTAGC
        maximum likelihood sequence       TGGAACCAGGATCATGCTACGGCGCGTAATCTACCACGATGCTTCTAGC
        maximum likelihood edit distance  0

evaluating sequence 68 of 100
        observed sequence                 GCTAGAAGCATCGTGGTAGATTACGCGCCGTAGCATGATCCTGGTTCCA
        maximum likelihood sequence       GCTAGAAGCATCGTGGTAGATTACGCGCCGTAGCATGATCCTGGTTCCA
        maximum likelihood edit distance  0

evaluating sequence 69 of 100
        observed sequence                 TGCTAGAAGCATCGTGGTAGATTACGCGCCGTAGCATGATCCTGGTTCCA
        maximum likelihood sequence       TGCTAGAAGCATCGTGGTAGATTACGCGCCGTAGCATGATCCTGGTTCCA
        maximum likelihood edit distance  0

evaluating sequence 70 of 100
        observed sequence                 TGGAACCAGGATCATGCTACGGCGCGTAATCTACCACGATGCTTCTAGCA
        maximum likelihood sequence       TGGAACCAGGATCATGCTACGGCGCGTAATCTACCACGATGCTTCTAGCA
        maximum likelihood edit distance  0

evaluating sequence 71 of 100
        observed sequence                 TTGCTAGAAGCATCGTGGTAGATTACGCGCCGTAGCATGATCCTGGTTCCA
        maximum likelihood sequence       TTGCTAGAAGCATCGTGGTAGATTACGCGCCGTAGCATGATCCTGGTTCCA
        maximum likelihood edit distance  0

evaluating sequence 72 of 100
        observed sequence                 GGAACCAGGATCATGCTACGGCGCGTAATCTACCACGATGCTTCTAGC
        maximum likelihood sequence       GGAACCAGGATCATGCTACGGCGCGTAATCTACCACGATGCTTCTAGC
        maximum likelihood edit distance  0

evaluating sequence 73 of 100
        observed sequence                 TGCTAGAAGCATCGTGGTAGATTACGCGCCGTAGCATGATCCTGGTTCCA
        maximum likelihood sequence       TGCTAGAAGCATCGTGGTAGATTACGCGCCGTAGCATGATCCTGGTTCCA
        maximum likelihood edit distance  0

evaluating sequence 74 of 100
        observed sequence                 TGCTAGAAGCATCGTGGTAGATTACGCGCCGTAGCATGATCCTGGTTCC
        maximum likelihood sequence       TGCTAGAAGCATCGTGGTAGATTACGCGCCGTAGCATGATCCTGGTTCC
        maximum likelihood edit distance  0

evaluating sequence 75 of 100
        observed sequence                 GCTAGAAGCATCGTGGTAGATTACGCGCCGTAGCATGATCCTGGTTCCA
        maximum likelihood sequence       GCTAGAAGCATCGTGGTAGATTACGCGCCGTAGCATGATCCTGGTTCCA
        maximum likelihood edit distance  0

evaluating sequence 76 of 100
        observed sequence                 TGGAACCAGGATCATGCTACGGCGCGTAATCTACCACGATGCTTCTAGCA
        maximum likelihood sequence       TGGAACCAGGATCATGCTACGGCGCGTAATCTACCACGATGCTTCTAGCA
        maximum likelihood edit distance  0

evaluating sequence 77 of 100
        observed sequence                 TGCTAGAAGCATCGTGGTAGATTACGCGCCGTAGCATGATCCTGGTTCCA
        maximum likelihood sequence       TGCTAGAAGCATCGTGGTAGATTACGCGCCGTAGCATGATCCTGGTTCCA
        maximum likelihood edit distance  0

evaluating sequence 78 of 100
        observed sequence                 CTAGAAGCATCGTGGTAGATTACGCGCCGTAGCATGATCCTGGTTCCA
        maximum likelihood sequence       CTAGAAGCATCGTGGTAGATTACGCGCCGTAGCATGATCCTGGTTCCA
        maximum likelihood edit distance  0

evaluating sequence 79 of 100
```

```
        observed sequence                 TGCTAGAAGCATCGTGGTAGATTACGCGCCGTAGCATGATCCTGGTTCCA
        maximum likelihood sequence       TGCTAGAAGCATCGTGGTAGATTACGCGCCGTAGCATGATCCTGGTTCCA
        maximum likelihood edit distance  0

evaluating sequence 80 of 100
        observed sequence                 TGCTAGAAGCATCGTGGTAGATTACGCGCCGTAGCATGATCCTGGTTCCA
        maximum likelihood sequence       TGCTAGAAGCATCGTGGTAGATTACGCGCCGTAGCATGATCCTGGTTCCA
        maximum likelihood edit distance  0

evaluating sequence 81 of 100
        observed sequence                 GAACCAGGATCATGCTACGGCGCGTAATCTACCACGATGCTTCTAGCA
        maximum likelihood sequence       GAACCAGGATCATGCTACGGCGCGTAATCTACCACGATGCTTCTAGCA
        maximum likelihood edit distance  0

evaluating sequence 82 of 100
        observed sequence                 TGGAACCAGGATCATGCTACGGCGCGTAATCTACCACGATGCTTCTAGCA
        maximum likelihood sequence       TGGAACCAGGATCATGCTACGGCGCGTAATCTACCACGATGCTTCTAGCA
        maximum likelihood edit distance  0

evaluating sequence 83 of 100
        observed sequence                 GTGGAACCAGGATCATGCTACGGCGCGTAATCTACCACGATGCTTCTAGCA
        maximum likelihood sequence       GTGGAACCAGGATCATGCTACGGCGCGTAATCTACCACGATGCTTCTAGCA
        maximum likelihood edit distance  0

evaluating sequence 84 of 100
        observed sequence                 TGGAACCAGGATCATGCTACGGCGCGTAATCTACCACGATGCTTCTAGCA
        maximum likelihood sequence       TGGAACCAGGATCATGCTACGGCGCGTAATCTACCACGATGCTTCTAGCA
        maximum likelihood edit distance  0

evaluating sequence 85 of 100
        observed sequence                 TGGAACCAGGATCATGCTACGGCGCGTAATCTACCACGATGCTTCTAGCA
        maximum likelihood sequence       TGGAACCAGGATCATGCTACGGCGCGTAATCTACCACGATGCTTCTAGCA
        maximum likelihood edit distance  0

evaluating sequence 86 of 100
        observed sequence                 TGCTAGAAGCATCGTGGTAGATTACGCGCCGTAGCATGATCCTGGTTCCA
        maximum likelihood sequence       TGCTAGAAGCATCGTGGTAGATTACGCGCCGTAGCATGATCCTGGTTCCA
        maximum likelihood edit distance  0

evaluating sequence 87 of 100
        observed sequence                 TGGAACCAGGATCATGCTACGGCGCGTAATCTACCACGATGCTTCTAGCA
        maximum likelihood sequence       TGGAACCAGGATCATGCTACGGCGCGTAATCTACCACGATGCTTCTAGCA
        maximum likelihood edit distance  0

evaluating sequence 88 of 100
        observed sequence                 TGGAACCAGGATCATGCTACGGCGCGTAATCTACCACGATGCTTCTAGC
        maximum likelihood sequence       TGGAACCAGGATCATGCTACGGCGCGTAATCTACCACGATGCTTCTAGC
        maximum likelihood edit distance  0

evaluating sequence 89 of 100
        observed sequence                 TGCTAGAAGCATCGTGGTAGATTACGCGCCGTAGCATGATCCTGGTTCCA
        maximum likelihood sequence       TGCTAGAAGCATCGTGGTAGATTACGCGCCGTAGCATGATCCTGGTTCCA
        maximum likelihood edit distance  0

evaluating sequence 90 of 100
        observed sequence                 TGGAACCAGGATCATGCTACGGCGCGTAATCTACCACGATGCTTCTAGCA
        maximum likelihood sequence       TGGAACCAGGATCATGCTACGGCGCGTAATCTACCACGATGCTTCTAGCA
        maximum likelihood edit distance  0

evaluating sequence 91 of 100
        observed sequence                 TGGAACCAGGATCATGCTACGGCGCGTAATCTACCACGATGCTTCTAGCA
        maximum likelihood sequence       TGGAACCAGGATCATGCTACGGCGCGTAATCTACCACGATGCTTCTAGCA
        maximum likelihood edit distance  0

evaluating sequence 92 of 100
        observed sequence                 TGCTAGAAGCATCGTGGTAGATTACGCGCCGTAGCATGATCCTGGTTCCA
        maximum likelihood sequence       TGCTAGAAGCATCGTGGTAGATTACGCGCCGTAGCATGATCCTGGTTCCA
        maximum likelihood edit distance  0

evaluating sequence 93 of 100
        observed sequence                 TGCTAGAAGCATCGTGGTAGATTACGCGCCGTAGCATGATCCTGGTTCCA
        maximum likelihood sequence       TGCTAGAAGCATCGTGGTAGATTACGCGCCGTAGCATGATCCTGGTTCCA
        maximum likelihood edit distance  0

evaluating sequence 94 of 100
        observed sequence                 TGGAACCAGGATCATGCTACGGCGCGTAATCTACCACGATGCTTCTAGCA
        maximum likelihood sequence       TGGAACCAGGATCATGCTACGGCGCGTAATCTACCACGATGCTTCTAGCA
        maximum likelihood edit distance  0

evaluating sequence 95 of 100
        observed sequence                 TGCTAGAAGCATCGTGGTAGATTACGCGCCGTAGCATGATCCTGGTTCCA
        maximum likelihood sequence       TGCTAGAAGCATCGTGGTAGATTACGCGCCGTAGCATGATCCTGGTTCCA
        maximum likelihood edit distance  0

evaluating sequence 96 of 100
        observed sequence                 GGAACCAGGATCATGCTACGGCGCGTAATCTACCACGATGCTTCTAGCA
        maximum likelihood sequence       GGAACCAGGATCATGCTACGGCGCGTAATCTACCACGATGCTTCTAGCA
        maximum likelihood edit distance  0

evaluating sequence 97 of 100
        observed sequence                 TTGCTAGAAGCATCGTGGTAGATTACGCGCCGTAGCATGATCCTGGTTCCA
        maximum likelihood sequence       TTGCTAGAAGCATCGTGGTAGATTACGCGCCGTAGCATGATCCTGGTTCCA
        maximum likelihood edit distance  0

evaluating sequence 98 of 100
        observed sequence                 TGGAACCAGGATCATGCTACGGCGCGTAATCTACCACGATGCTTCTAGC
        maximum likelihood sequence       TGGAACCAGGATCATGCTACGGCGCGTAATCTACCACGATGCTTCTAGC
        maximum likelihood edit distance  0
```

```
evaluating sequence 99 of 100
	observed sequence                 TGCTAGAAGCATCGTGGTAGATTACGCGCCGTAGCATGATCCTGGTTCCA
	maximum likelihood sequence       TGCTAGAAGCATCGTGGTAGATTACGCGCCGTAGCATGATCCTGGTTCCA
	maximum likelihood edit distance  0

evaluating sequence 100 of 100
	observed sequence                 GTGGAACCAGGATCATGCTACGGCGCGTAATCTACCACGATGCTTCTAGC
	maximum likelihood sequence       GTGGAACCAGGATCATGCTACGGCGCGTAATCTACCACGATGCTTCTAGC
	maximum likelihood edit distance  0

DATASET STATISTICS:
	assumed error rate    8.333333333333332%
	total bases observed  4966
	total edits accepted  0
	inferred error rate   0.0%
```
